# Elucidating immune-related gene transcriptional programs via factorization of large-scale RNA-profiles

**DOI:** 10.1101/2024.05.10.593433

**Authors:** Shan He, Matthew M. Gubin, Hind Rafei, Rafet Basar, Merve Dede, Xianli Jiang, Qingnan Liang, Yukun Tan, Kunhee Kim, Maura L. Gillison, Katayoun Rezvani, Weiyi Peng, Cara Haymaker, Sharia Hernandez, Luisa M. Solis, Vakul Mohanty, Ken Chen

**Affiliations:** Department of Bioinformatics and Computational Biology, The University of Texas MD Anderson Cancer Center, Houston, Texas 77030, USA; Department of Immunology, The University of Texas MD Anderson Cancer Center, Houston, Texas 77030, USA; Department of Stem Cell Transplantation and Cellular Therapy, The University of Texas MD Anderson Cancer Center, Houston, Texas 77030, USA; Department of Thoracic/Head and Neck Medical Oncology, University of Texas MD Anderson Cancer Center, Houston, Texas 77030, USA; Department of Biology and Biochemistry, The University of Houston, Houston, TX, USA; Department of Translational Molecular Pathology, The University of Texas MD Anderson Cancer Center, Houston, TX, USA

**Keywords:** Immunology, ImmuneSigDB, Gene sets, TCGA, Immunotherapy, Spatial Transcriptomics

## Abstract

Recent developments in immunotherapy, including immune checkpoint blockade (ICB) and adoptive cell therapy, have encountered challenges such as immune-related adverse events and resistance, especially in solid tumors. To advance the field, a deeper understanding of the molecular mechanisms behind treatment responses and resistance is essential. However, the lack of functionally characterized immune-related gene sets has limited data-driven immunological research. To address this gap, we adopted non-negative matrix factorization on 83 human bulk RNA-seq datasets and constructed 28 immune-specific gene sets. After rigorous immunologist-led manual annotations and orthogonal validations across immunological contexts and functional omics data, we demonstrated that these gene sets can be applied to refine pan-cancer immune subtypes, improve ICB response prediction and functionally annotate spatial transcriptomic data. These functional gene sets, informing diverse immune states, will advance our understanding of immunology and cancer research.

## Introduction

Despite recent breakthroughs in immunotherapy such as immune checkpoint blockade (ICB)^1^ and adoptive cell therapy (ACT)^2^, only a limited fraction of patients benefits from these biomedical advancements. Challenges related to immune-related adverse events and resistance remain and hinder positive outcomes, especially in solid tumors^3^. It is essential for the field of immunotherapy to understand molecular mechanisms underlying the treatment resistance and responses better predict treatment outcomes and develop novel therapeutics. A systematic way to address these challenges is through statistical analysis of transcriptional programs of human patient samples^4^, which can uncover novel gene expression patterns associated with immunotherapeutic responses. Interpretation of transcriptional programs and their functional significance, however, relies on utilizing previously established gene signature knowledgebases^5–8^ such as Molecular Signature Database (MSigDB) Hallmarks (Hallmark)^9^, Kyoto Encyclopedia of Genes and Genomes (KEGG)^10^, Gene Ontology (GO)^11^, etc. Unfortunately, these knowledgebases are largely cell-type agnostic and lack granularity or immune specificity^7^. Such biases severely hamper data interpretation in immunological studies utilizing high-throughput RNA profiling, resulting in many missed opportunities. To further advance cancer immunotherapy, there is an urgent need to discover and expand immune-related gene signatures/sets (irGSs) in current knowledgebases to comprehensively represent diverse immune-specific transcriptional states and to apply them to interpret molecular functions.

A closer inspection of the above-mentioned knowledgebases reveals important limitations that are inherent to each source. For example, Hallmark^9^ contains only 7 immune-related pathways, with an average of 160 member genes each. The large gene sets in this database can potentially inflate biological significance, without genuine and specific relevance to the designated functions. On the other hand, KEGG^10^ contains mostly metabolism related pathways that are not specific to immune systems. The GO^11^ terms, with its hierarchical nature, result in collective enrichment of general terms with overlapping functions, diminishing statistical power to detect specific terms. Moreover, although there are some immune-related pathways in the above-mentioned knowledgebases, they are too broadly defined. For example, in KEGG, the term T cell receptor (TCR) signaling does not distinguish receptor formation by T cells from antigen presentation by antigen presenting cells, two complimentary functions in TCR signaling. In Hallmark, “inflammatory response” is too broad to inform detailed molecular programs or mechanisms. Therefore, there is a need to define more specific gene sets and refined annotations to achieve higher granularity.

To address this knowledge gap, some immune specific knowledgebases have been constructed. In particular, ImmuneSigDB ^12^, consisting of over 5,000 irGSs, was derived from RNA-expression datasets. These RNA expression datasets are very rich, derived from nearly 400 noncancer-specific immunological experiments, containing samples challenged with pathogen infections, cytokines or immunological perturbations of different kinds and magnitudes, thus possessing rich immunological states. Some immunological states may be shared by a variety of conditions. For example, the severe immune dysregulation and inflammation^13,14^, observed in sepsis samples may inform immune cell proliferation, inflammatory signaling, and cellular dysfunction and exhaustion in tumor immune microenvironment (TIME).

However, the ImmuneSigDB study, conducted nearly a decade ago, applied empirical approaches that may have resulted in some limitations. First, the study examined sets of genes that are differentially expressed (DEGs) across different comparisons of interest, and these comparison groups were subjectively decided by researchers as important and immune-relevant^12^. Second, differential gene expression analysis tends to output large numbers of DEGs (200±70, max=1992), with 50% of the gene set size ranges from 190 to 196 member genes. Third, the derived irGSs tend to have limited translational implications as they can only be used to differentiate major states (such as cell lineages) instead of more granular substates. Fourth, given that these comparison groups are defined based on empirical knowledge, ImmuneSigDB irGSs are likely incomplete. Lastly, the ImmuneSigDB irGSs are simply annotated using the GEO ID of the dataset and the comparison groups, thus containing limited annotations of immune functions, hampering results interpretation.

In this paper, we aim to construct concise and well-annotated immune-specific gene sets using Non-negative Matrix Factorization (NMF)^15^, a powerful factor analysis technique ^16–20^ that has not yet been applied to ImmuneSigDB data. By decomposing gene expression of samples from ImmuneSigDB into composite programs, followed by addition statistical analysis to consolidate and annotate these programs, we have identified 28 novel irGSs describing various immunological processes. These irGSs can be applied to study and characterize various immune-related transcriptional states such as those in play in TIMEs. We have systematically validated gene set annotations on 12 different datasets and explored their translational implications on 15 different datasets (see **Methods Data Sources**). These datasets include various modalities, CITE-seq, BulkRNAseq, scRNAseq and spatial transcriptomics, and were profiled from a variety of immunological contexts, including but not limited to cytokine perturbation experiments, infectious disease, and treatment naïve or immunotherapy treated cancer patients. Through these exploratory analyses, we showed that these gene sets can better characterize pan-cancer tumor-immune subtypes, separate ICB response in melanoma, liver and lung cancer and reveal spatial heterogeneity in ovarian, intestinal and breast cancer samples.

## Results

### 28 gene sets covering diverse immunological functions in the TIME

The ImmuneSigDB datasets comprise 83 human bulk gene expression profiles with 1826 RNA samples from studies published in leading immunology journals (**Figure S1a**). The datasets included in the ImmuneSigDB were derived from patients affected by severe infections and sepsis or from cell lines challenged with experimental manipulations ^12^: gamma delta T cells activated by *IL2*^21^, and *CD14*+ monocytes^22^ cultured with IFN-gamma and stimulated with toll-like receptor 2 ligand. Such a large collection of datasets entails transcriptional profiles from diverse cell states, making them suitable to derive generalizable irGSs.

The datasets encompass gene expression data obtained from human and mouse samples (**Figure S1b**) that originated from samples deriving from different cell types (**Figure S1c**). Among datasets profiled from *Homo sapiens*, the number of genes and samples profiled varies among datasets (**Figure S1d and S1e**). We partitioned the 83 ImmuneSigDB human datasets into 47 lymphoid-derived and 36 myeloid-derived datasets.

To obtain unique irGSs, we carried out the analytic workflow, illustrated in **Figure 1a** and detailed in **Figure S1f** and **methods**, separately for the lymphoid and the myeloid data. We applied NMF on each qualifying dataset (see **Methods**) and obtained the top 50 genes from each latent factor based on the NMF loadings, producing a total of 529 gene sets. To ensure generalizability, gene sets derived from one dataset were kept only if they overlap at least 20% with at least one gene set derived from another dataset. To ensure non-redundancy, we excluded gene sets with over 20% overlap with gene sets from the same dataset, resulting in 115 gene sets. To further reduce redundancy, we employed the algorithm proposed by Tirosh et al^16^, whereby gene sets are progressively merged into meta programs (MP) based on Jaccard distances (see **Methods**).

**Figure 1.**
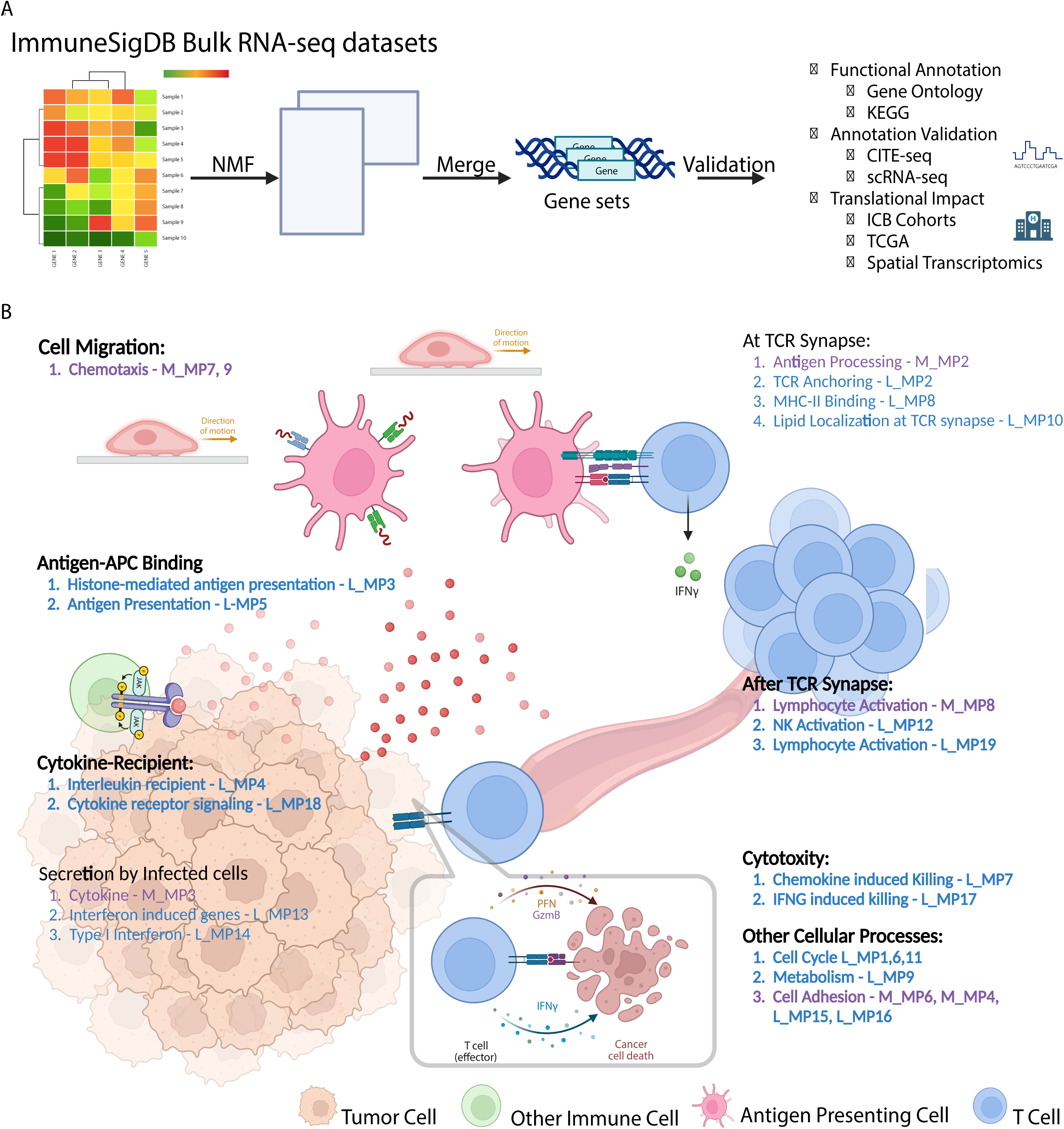
Analy&c workflow and constructed gene sets. **(A)** Gene set construction analysis workflow: We downloaded 83 immunology relevant human studies and performed NMF on each of the qualifying dataset. We curated robust NMF programs and merged them into meta gene sets. We performed over-representation enrichment analysis for each meta gene set using KEGG pathways, Hallmarks, and biological process terms from GO. The name and annotation of each meta gene set was determined based on counts and False Discovery Rate (FDR) adjusted p-values of the highly enriched terms. We seek relevant multimodal datasets to functionally name and validate each gene set, and further examined the translational utilities of these gene sets. **(B)** The roles of the constructed gene sets in central immunity: constructed gene sets overlaid near the most relevant processes during central immune activities in tumor-immune microenvironment. Created with BioRender.com.

Using the workflow described above, we derived 19 immune-related MPs (irMPs) from lymphoid samples (L_MP1-19) and 9 irMPs from myeloid samples (M_MP1-9) (**Table 1**). To annotate each irMP with immune-relevant functions, we performed over-representation analysis (ORA) using biological processes from GO, KEGG and Hallmark and sorted the enrichment results first by counts of core enrichment then by ascending adjusted p-values. Enrichment results and constituent genes for each L_MP and M_MP were presented in **Table S1-S4**. We named the irMPs based on top enriched terms and refined the annotations by going over the constituent genes and the stringDB^23^ protein-protein interaction network, followed by meticulous scrutiny by multiple immunologists (Matthew Gubin Ph.D.; Hind Rafei M.D.; Rafet Basar M.D.; Katy Rezvani M.D/Ph.D.; Weiyi Peng M.D/Ph.D.) from MD Anderson Cancer Center. Rationales for each gene set annotation have been documented in **Supplementary Notes**.

**Table 1.**
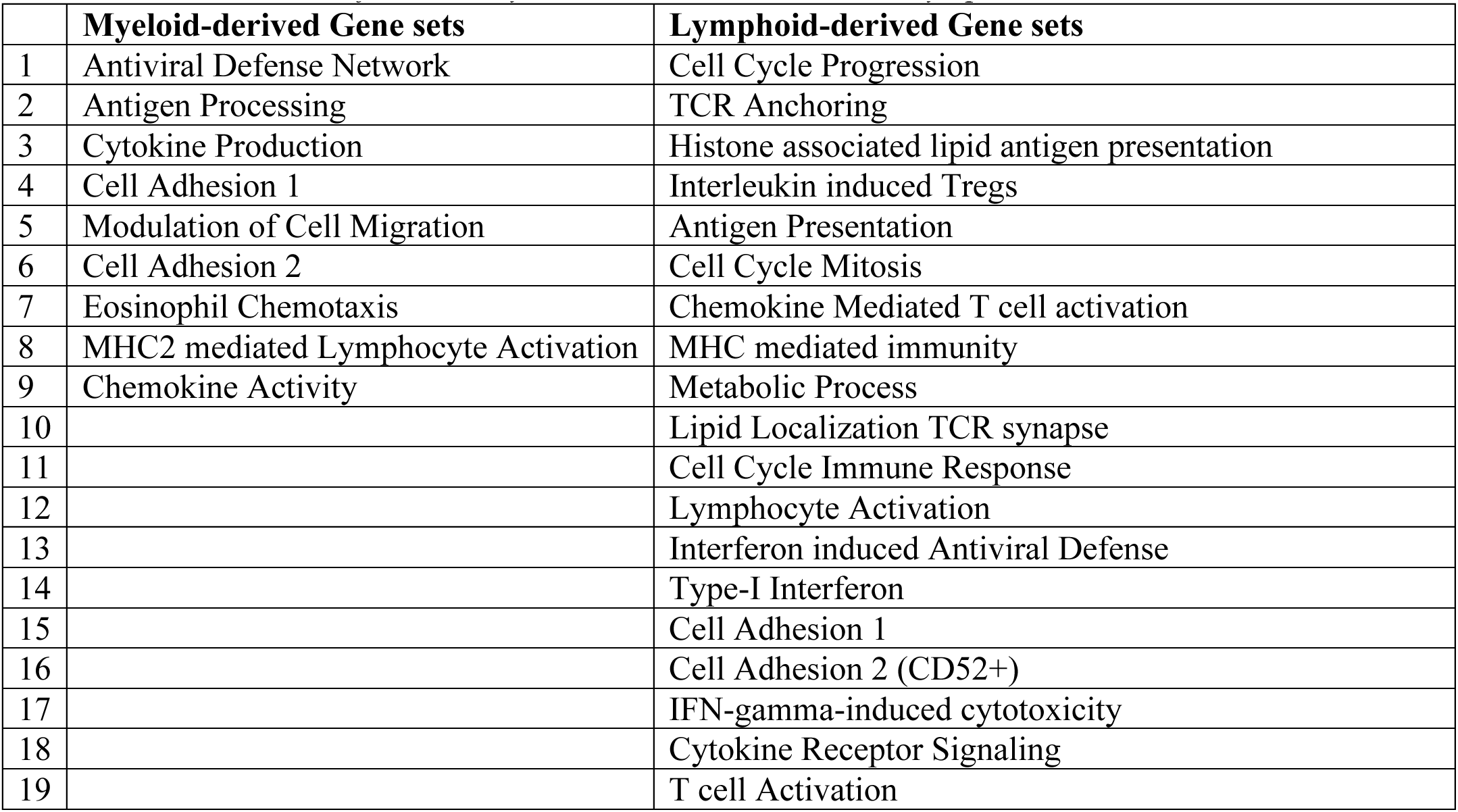
Annotations for the 9 Myeloid-derived irMPs and 19 Lymphoid-derived irMPs.

Overall, the L_MPs are highly specific to immune functions, consisting of pathways such as lymphocyte activation, antigen processing, TCR anchoring and cytotoxic functions. M_MPs are less diverse, including pathways related to cell migration, adhesion, and antiviral response (**Figure 1b and Table 1**). A few MPs are related to core cellular processes, such as cell cycle and metabolism.

To access the novelty of irMPs, we compared and quantified their constituent genes overlap (see **Methods**) with 231 manually curated Spectra immune gene sets^24^, 5,000 ImmuneSigDB pathways, 50 Hallmark pathways and 29 immune-related KEGG (irKEGG). We observed significant distinction (FDR-adjusted p-value < 0.05) in 99.5%, 98%, 99%, and 87% of the comparison with Spectra gene sets (**Figure S2**), ImmuneSigDB (**Figures S3-S4**), Hallmark (**Figures S5**), and irKEGGs (**Figure S6**), respectively.

### Functional characterization of the irMPs

To validate the functional annotation of irMPs and demonstrate their general applicability, we compiled 12 publicly accessible gene expression datasets encompassing a diverse range of immune contexts, captured through various sequencing techniques (see **Methods Data Sources**). These datasets were derived from well-designed immunological experiments with well-defined phenotypes, providing independent and orthogonal evidence required to validate annotations of irMPs.

Compared to RNA expression, protein expression is often a more direct indicator of biological activity and reflects real physiological processes. We, therefore, leveraged two CITE-seq datasets^25,26^, in which the whole transcriptome and ∼200 surface antibodies were simultaneously profiled in the peripheral blood mononuclear cells (PBMC) of healthy human and COVID-19 patients, respectively. irMPs were scored for each cell in the RNA level using single sample gene set enrichment analysis (ssGSEA)^27^ (see **Methods**). For each irMP of interest, cells were dichotomized into MP-high or MP-low (see **Methods**) and the abundance of proteins indicative of the function described by the irMP was compared between MP-high and MP-low groups to show consistency between the irMP and proteomic measurement. For irMP without relevant proteins in the CITE-seq data, we defined highly relevant phenotypes in appropriate scRNA-seq data or TCGA pan-cancer data and checked for gene set enrichment in a pre-defined phenotype. In addition, we utilized protein-protein interaction network constructed by the *StringDB* database^28^ to elucidate the activity cascade of each irMP (**Figure S7-8**).

Interestingly, the irMPs we obtained can be broadly classified into categories that align with the sequential progression of a typical immune response: cytokine/chemokine production, cell adhesion, cell chemotaxis, antigen processing, antigen presentation, lymphocyte activation, and cytotoxicity (**Figure 1b**), indicating the key importance of these functional components and the rich molecular programs that have been under-explored.

Below we describe the function of each irMP and evidence from orthogonal validations.

#### 1. Cytokine/Chemokine production

There are two cytokine/chemokine related irMPs from myeloid data (M_MP3, M_MP9) and two (L_MP7, L_MP18) from lymphoid data. To validate their annotations, we examined scRNAseq data profiled from two severe COVID-19 patients, with symptoms consistent with inflammatory cytokine storms (ICS)^29^. If the irMPs are in fact related to pathogenic cytokine/chemokine activities, they should be enriched in cells deriving from patients rather than health controls. As myeloid cells were concluded to be the source of ICS ^29^, we extracted myeloid cells and performed gene set enrichment analysis (GSEA) on fold-changes of genes expression between ICS and healthy controls. Indeed, we found that all four irMPs were significantly enriched in ICS patients relative to healthy controls (**Figure 2a**).

**Figure 2.**
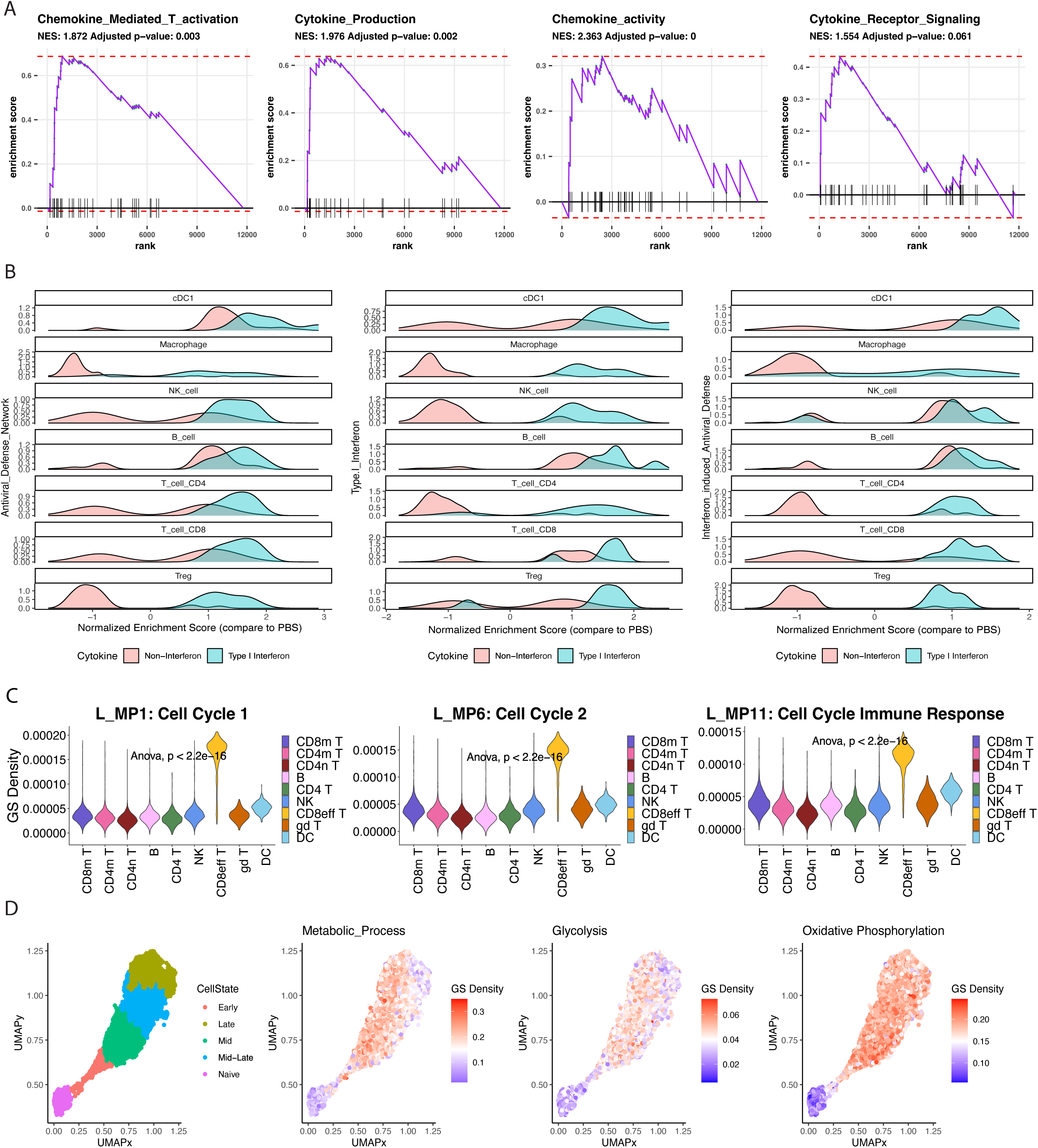
Func&onal characteriza&on of the generic immune gene sets. **(A)** Barcode Enrichment Plot of 4 cytokine/chemokine release irMPs; All genes from the data are ranked along x-axis according to enrichment score (y-axis). There is a significant upregulation in all 4 irMPs in samples with severe COVID-19 patients with cytokine storm, and the ranked position of each gene within a signature is shown in the x axis. **(B)** Density plot comparing the normalized enrichment score of three antiviral irMPs in different immune cell types when stimulated with interferon vs non-interferon. **(C)** Violin plot comparing cell cycle MP activities calculated using GS Density across different cell types identified from COVID-19 single cell atlas. Anova test was performed for multi-group comparison **(D)** UMAPs labeled with T cell activation stages, L_MP9 activity, Glycolysis and Oxidative Phosphorylation activities from Hallmark using GS Density.

Upon further examining the PPI network constructed from the genes in M_MP9 (**Figure S7.9**), we found that M_MP9 has a central module of chemokine genes (*CXCL10, CXCL9, CCL4, CCL2, CCRL2, CCL7, CCL8*), connected with APC-specific Human Leukocyte Antigen (HLA) class II genes (*HLA-DPA1, HLA-DMA, HLA-DRA*) by *CD74*, a chaperone responsible for antigen presentation^30^. This further supports the hypothesis that M_MP9 captures the signaling cascade via chemokine secretion leading to the activation of APCs.

#### 2. Interferon-signaling pathways

There is one antiviral response irMP (M_MP1) from myeloid data and two interferon-mediated viral defense irMPs (L_MP13, L_MP14) from lymphoid data. To validate these annotations, we used cytokine dictionary curated by Cui et.al^31^, in which the authors have profiled scRNA-seq on 17 different immune cell types extracted from lymph nodes of mice treated with 86 different cytokines *in vivo* or phosphate buffered saline (PBS) as control. As these gene sets are associated with interferon signaling, they should be induced upon stimulation by interferon rather than other cytokines. For each cytokine treatment in each cell-type, we performed differential expression relative to corresponding PBS controls followed by GSEA. Across cell-types, we observed that all three irMPs show a general trend of significantly higher enrichment under interferon treatments (IFN-α1, IFN-β, IFN-ε, IFN-κ, IFN-γ, IFN-λ2) relative to other cytokine treatments (**Figure 2b**), indicating that these three irMPs are all signaling pathways downstream of interferon stimulation.

#### 3. Cell Adhesion/Cell Migration

There are two cell adhesion (M_MP4, M_MP6) and two chemotaxis related (M_MP5, M_MP7) irMPs from myeloid data, and two cell adhesion irMPs (L_MP15 and L_MP16) from lymphoid data. Cell adhesion and migration are essential steps in the immune response, as they facilitate downstream immune cell trafficking, activation, and effector function^32^. In general, the three steps in cell adhesion are rolling, weak, and strong adhesion^33^. Rolling involves selectin molecules (L-selectin, E-selectin, P-selectin) loosely moving immune cells on endothelial surface. Weak adhesion occurs when integrin molecules (CD49d, CD29, CD11a, CD11b, CD11c) weakly bind to ligands (ICAM) on endothelial cells, slowing down the immune cell movement. Strong adhesion occurs when integrin molecules firmly anchor the immune cells to the endothelial surface, allowing them to extravasate through the endothelium and into surrounding tissues. If the irMPs are associated with cell-adhesion and chemotaxis, their activity should be concordant with the expression of relevant protein markers discussed above.

To validate the functional annotation of these irMPs, we dichotomized cells in the PBMC CITE-seq data^34^ by their RNA-based MP activity levels and examined the abundance levels of relevant proteins (see **Methods**). We found that, among myeloid cells, the activity of M_MP4 and M_MP6 are associated with the abundance of cell adhesion proteins related to both rolling and weak adhesion (**Figure S9a**). In particular, myeloid cells with higher M_MP4 have the highest CD62L abundance, suggesting that M_MP4 is driven by L-selectin, a cell adhesion molecule on the surfaces of leukocytes. In contrast, the activity levels of M_MP5 and M_MP7 that are associated with cellular migration are associated with lower abundance of cell adhesion marker proteins (**Figure S9a)**, consistent with their contrasting functional annotation. Among lymphoid cells, we found that L_MP15 shows strong association with integrins (CD49d, CD29, CD11a, CD11b, CD11c), and L_MP16 with selectins (CD62P, CD62L) (**Figure S9b**). These two irMPs differ from myeloid-derived cell adhesion irMPs in that L_MP15 and L_MP16 are enriched in HLA genes, suggesting possible cell adhesion through antigen presentation.

#### 4. Core cellular process

Besides highly immune-specific gene sets, there are also gene sets related to core cellular processes. In particular, there are three irMPs (L_MP1, 6, 11) annotated as cell cycle irMPs from lymphoid data. To validate their functions, we used a COVID-19 single cell gene expression atlas^35^, where author identified a cluster of proliferative CD8+ effector T cells. Since proliferative lymphocytes undergo clonal expansion, we expect cell cycle pathways to be upregulated in this cluster.^28^ Indeed, we observed significantly higher gene set activities^26^ for all three cell-cycle irMPs in the proliferative lymphocyte cluster (**Figure 2c**) than in the other clusters.

There is one metabolism-related gene set (L_MP9), enriched in glycolysis genes (*ENO1, PGK1, ALDOA, PGAM1, TPI1, STMN1, LDHA, HMMR, ENO2,* and *PPIA*). To confirm its relevancy to metabolism, we utilized a study in which authors perform in vitro stimulation of naïve CD8+ T cells and characterized metabolic programs at different activation stages^36^, showing activation of T-cells is accompanied with increased metabolism. We observed that the activity of L_MP9 increases as T cells become more activated, which also coincided with the trend of glycolysis and oxidative phosphorylation (**Figure 2d**).

Besides the above-mentioned gene sets that represent shared functions in both myeloid and lymphoid compartments, we have also discovered irMPs that are highly specific to a cell type or a contact interface between two immune cells.

#### 5. Leukocyte Activation

Two of the lymphoid irMPs (L_MP12, L_MP19) were annotated as Lymphocyte activation and T cell activation, respectively. The activation of these pathways should coincide with high expression of protein markers reflective of lymphocyte activation. As COVID-19 induces immune cell activation, we leveraged the COVID-19 PBMC CITE-seq data^26^. We extracted NK and CD8 T cells based on SingleR^37^ annotation (see **Methods**) and observed that NK/T cells with higher L_MP12 activity have higher abundance of CD16, CD56, KLRG1, CD8 and CD69 (**Figure 3a**), which are known to be expressed by activated T and NK cells. It is also known that CD8 can be expressed by cytotoxic NK cells^38^. Cells with higher L_MP19 activity have higher abundance of CD3, CD4, CD8, and CD69 (**Figure 3a**), which are known to be expressed by activated T cells. In addition, L_MP12 and L_MP19 appear to be associated with checkpoint proteins, TIGIT, CD152 (CTLA4) and CD279 (PD-1), which also indicate activation and regulation of immune homeostasis^39^.

**Figure 3.**
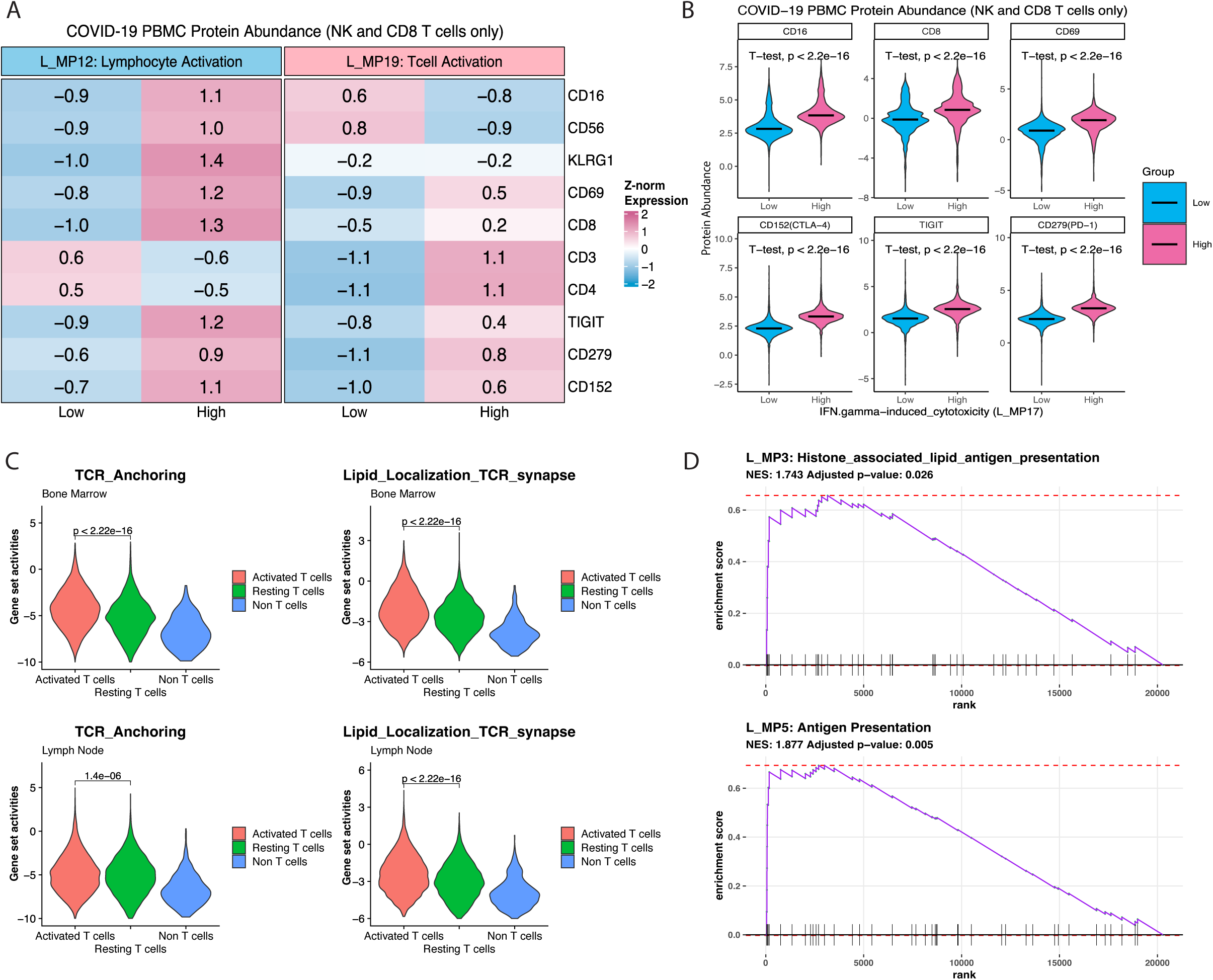
Func&onal characteriza&on for the specific irMPs. **(A)** COVID-19 PBMC cells annotated as NK or CD8+ T cells were dichotomized into two groups (High vs Low) based on the activity levels of 2 lymphoid-derived gene sets: L_MP12 and 19, estimated respectively from the RNA expression data. Shown in the heatmap table (and colored accordingly) are the average Z-transformed protein abundance of surface proteins: CD16, CD56, KLRG1, CD8, CD3, CD4, CD69, TIGIT, CD279, and CD152 in each of the respective cell groups. (**B**) Shown in violin plots are the abundance of 6 surface proteins: CD16, CD8, CD69, CD152, CD279, TIGIT in two dichotomized COVID-19 PBMC cells (annotated as NK or CD8+ T cells) groups with respectively low and high L_MP17 activity levels, determined from the RNA expression data. In all the cases, dichotomization was performed using Gaussian Mixture Models. Activities between two groups were compared using T test with significance level 0.05. **(C)** Violin plot comparing two TCR-related irMPs across different T cell activation states using ssGSEA. **(D)** Barcode Enrichment Plot of L_MP3 and L_MP5; All genes from the TCGA are ranked along x-axis according to enrichment score (y-axis). There is a significant upregulation in both L_MP3 and L_MP5 in samples with a high persistent tumor mutational burden, and the ranked position of each gene within a signature is shown in the x axis.

#### 6. Interferon-induced cytotoxicity pathway

One lymphoid irMP (L_MP17) was annotated as IFN-gamma induced cytotoxicity. As COVID-19 induces cytotoxicity, we leveraged the COVID-19 PBMC CITE-seq data^26^. Extracting NK and CD8 T cells based on SingleR annotation, we observed that cells with higher L_MP17 activity have significantly higher abundance of CD8, CD16, and CD69 (FDR adjusted p-value<0.05), which are protein markers for cytotoxic CD8 T cells and cytotoxic NK cells (**Figure 3b)**. In addition, higher abundance of TIGIT, CD152 (CTLA4) and CD279 (PD-1) in L_MP17 high cells indicate activation, required for achieving cytotoxicity and regulating immune response to avoid over-activation^39^. The protein-protein interaction network for this MP suggests the following activity cascade (**Figure S7.17**): Activated NK cells (*NKG7, GNLY*) secrete *IFNG*, inducing the activation of granzyme-producing (*GZMH, GZMB, GZMK, GZMA*) cytotoxic macrophages (*LYZ*). Moreover, downstream targets of leukocyte activation are also captured by L_MP17, specifically *NFKBIA, KLF6* and Nuclear Receptor genes (*NR4A2, NR4A3*), which work harmoniously to ensure T cell homeostasis and proper proliferation^40,41^.

To further validate the cytotoxicity nature of this irMP, we used a COVID-19 single cell atlas data^35^. We observed that lymphoid cells with higher L_MP17 activities correspond to cells with higher cytotoxicity scores (**Figure S9c)**, calculated using GSDensity with the following genes: *PRF1, GZMA, GZMB, GZMH,* and *GNLY* ^35^. In addition to cytotoxicity, L_MP17 scored high in the CD8 effector T cell cluster, a cluster defined as highly proliferating T cells in the paper, suggesting that L_MP17 describes not only a cytotoxic but also a proliferative cell state. Indeed, immediately connected to the granzyme modules (**Figure S7.17**) are *TUBA4A* and *TUBA1C*, which are known to be involved in forming and stabilizing microtubules in proliferating T cells during various stages of cell cycling, contributing to spindle formation during mitosis and ensuring proper chromosome segregation and accurate cell division^42^.

#### 7. T cell receptors (TCR)

Two lymphoid irMPs (L_MP2, L_MP10) were annotated as TCR-anchoring and lipid localization at TCR synapse, respectively. L_MP2 shows significant enrichment in antigen receptor-mediated signaling pathways (**Table S1.2**). In addition, L_MP2 is enriched in genes related to T-cell surface glycoprotein and transmembrane adapter (*CD247, CD2, CD28, CD3D, TRAT1, CD52, HLA-E*) and genes that have key roles in T-cell antigen receptor-linked signal transduction pathways (*ICOS, SLAMF1, LCK, LAT*)^43–46^ (**Figure S7.2**). As we expect TCR signaling to be upregulated when T cells are activated, we examined scRNA-seq from T cells isolated from bone marrow (BM) and lymph nodes (LN) of healthy donors that were cultured in resting condition vs activated by anti-CD3/anti-CD28 antibodies^47^. As expected, T-cells showed higher L_MP2 and L_MP10 activities, indicating cell-type specificity (**Figure 3c**). Further, the activities of these gene sets were significantly higher in activated T-cells indicating their relevance to TCR engagement upon T cell activation (**Figure 3c**).

#### 8. Antigen Presentation and Processing pathway

Two of the 19 lymphoid irMPs (L_MP3 and L_MP5) are related to antigen-presentation mediating immunity. We named L_MP3 as Histone-associated lipid antigen presentation and L_MP5 as Antigen presentation. L_MP3 is enriched in chromatin organization histones genes with lipid antigen presentation genes connected by a V(D)J recombination gene RAG1. Histone genes play an important factor in V(D)J recombination and formation of antigen receptors on lymphocytes^48^. The PPI network of L_MP3 (**Figure S7.3**) shows a HISTONE gene module and a *CD1* gene module connected by *RAG1*. L_MP5 is enriched in genes responsible for cell cycle regulation (*DLGAP5, BUB1B, CCNB1*) and genome stability (*TUBB, HMGB4, HNRNPA2B1, PTTG1, HMGB1*), each of which is an important element for V(D)J recombination fidelity^49^ (*RAG1* and *RAG2*) (**Figure S7.5**).

To validate the antigen presentation function of L_MP3 and L_MP5, we used pan-cancer mutational and gene-expression data from TCGA^50^. Persistent tumor mutational burden (pTMB), defined as the number of single copy and multiple copy mutations^51^, informs immune activation. Tumors with higher pTMB burdens are more likely to be visible to the immune system and are associated with sustained anti-tumor immune responses and improved response to ICB^51^. These tumors should therefore have higher activity of antigen presentation pathways. We performed differential gene expression analysis (DGE) between TCGA samples with high vs. low pTMB, regardless of cancer type and stage (see **Methods**). GSEA with 19 lymphoid irMPs was performed. Both L_MP3 and L_MP5 are significantly upregulated among samples with high pTMB (**Figure 3d**), confirming the antigen presentation function of these two irMPs. Notably, L_MP3 and L_MP5 are the only two irMPs (out of the 19) that are significantly enriched among high pTMB samples in the full GSEA profiles (FDR adjusted p-value <0.05) (**Figure S9d).**

#### 9. pMHC-TCR contact interface

Among myeloid irMPs, M_MP2 was annotated as Antigen Processing. **Figure S10a** depicts a typical antigen uptake by an Antigen Presenting Cell (APC), such as, dendritic cell (DC), in which an immature DC (iDC) transforms into a mature DC (mDC) upon detecting pathogen-associated molecular patterns (PAMPs). This activation process involves the DC recognizing PAMPs, leading to upregulation of major histocompatibility complex (MHC) molecules for antigen presentation and the co-stimulatory molecules *CD80*/*CD86*, crucial for T-cell activation^52,53^. Simultaneously, there is a change in cytoskeleton organization, notably the F-actin, to facilitate processing and presentation of the peptide-MHC (p-MHC) complexes on their surface for downstream T-cell activation.

Concordant with **Figure S10a**, we observed that APCs (Dendritic cells and Macrophages, identified by *SingleR*^37^) with higher ssGSEA in M_MP2 also have significantly (p-value<0.05) higher abundance in MHC proteins in the COVID-19 CITE-seq data (**Figure S10a**), such as HLA-F, HLA-A-B-C, and HLA-DR, suggesting APC activation. In addition, these cells with higher M_MP2 activity also have significantly (p-value<0.05) higher abundance in CD11a/ CD18, which facilitate downstream T cell binding, and in CD86, a co-stimulatory molecule for T cells activation.

Among lymphoid irMPs, L_MP8 was annotated as MHC mediated immunity. **Figure S10b** depicts a typical interaction between a mDC and a T cell at the TCR synapse: Upon recognition of the p-MHC complex by TCR, the T cell upregulates *CD69*, indicating that the T cell has been successfully engaged and activated. Concordant with **Figure 3f**, we observed that cells with higher M_MP2 activity also have significantly (p-value<0.05) higher abundance in MHC molecules, CD69 and TCRab from COVID-19 CITE-seq data, indicating successful TCR engagement at the immunological synapse.

Among myeloid irMP, M_MP8 was annotated as MHC-II-mediated lymphocyte activation because it is enriched in MHC-II markers (*HLA-DRB4, HLA-DQB1, HLA-DMA, HLA-DMB*) and T cell receptor complex (*TRAC* and *TRBC1*), suggesting MHC-II dependent CD4 T cell activation (as MHC-II presents to helper T cells)^54^. To confirm if this gene set is describing functions at the interface between macrophages and CD4 T cells, we extracted macrophages and CD4 T cells based on SingleR annotation and observed that cells with high activity in M_MP8 have higher protein abundance in M2-like macrophages (CD163, CD206), T cell activation (CD45RO, TCR) and MHC-II (HLA-DR) (**Figure S10c**), suggesting CD4 T cell activation via MHC-II on M2-like macrophages.

#### 10. Interleukin-induced Tregs

There is one lymphoid irMP (L_MP4) annotated as interleukin induced Tregs because it is enriched in Treg activation markers (*CTLA4, IL2RA, TIGIT, TNFRSF9, IL1R2*). **Figure S10d** depicts one of the mechanisms for Treg survival: binding between *IL2* and *CD25 (IL2RA)* promotes the growth and suppressive functions of Tregs, upregulating *CD39*^55,56^, eliciting a more suppressive Treg phenotype, defined by upregulation of *TNFRSF9* and checkpoint transcripts^56^. Using COVID-19 CITE-seq data, we extracted all Tregs based on *SingleR* annotation and found that Tregs with higher L_MP4 activity do have significantly (p-value<0.05) higher abundance in IL2 receptors (IL2RA, IL2RB), CD39, CTLA4, TIGIT, and TNFRS9 (**Figure S10d**), which are all protein markers over-expressed in effector Tregs, suggesting possible interleukin-induced Treg activation.

### irMPs redefine immune subtypes in TCGA pan-cancer data

In TCGA Pan-cancer immunity study, tumor samples were previously categorized into six major immune subtypes (C1: Wound Healing, C2: IFN-gamma dominant, C3: Inflammatory, C4: Lymphocyte Depleted, C5: Immunologically Quiet and C6: TGF-beta dominant), based on 160 immune signatures^57^. We performed Uniform Manifold Approximation and Projection (UMAP) based on ssGSEA^27^ scores with the 28 irMPs (**Figure S11a**). Since C4 and C6 are comprised of less than 10% and 2% of TCGA samples, respectively, we did not see a clear separation of these two clusters, but C4 is mostly distributed in the bottom left and C6 is mostly distributed in the top right of the UMAP (**Figure S11a**). Nevertheless, the UMAP well assigned the tumor samples into C1+C2, C3, or C5 subtypes with significant (log rank p-value<0.0001) differential overall survival between C1+C2 and C3+C5 (**Figure 4a-b**), but the overall survival difference between C3 and C5 is insignificant (p-value=0.623), despite having significantly (p-value<2.2e-16) different levels of lymphocyte infiltration. Through k-mean clustering with k=6 based on the same UMAP, we re-clustered the same set of TCGA samples into 6 new subtypes (**Figure 4c**), each of which is enriched in different types of cancer (**Figure 4d**).

**Figure 4.**
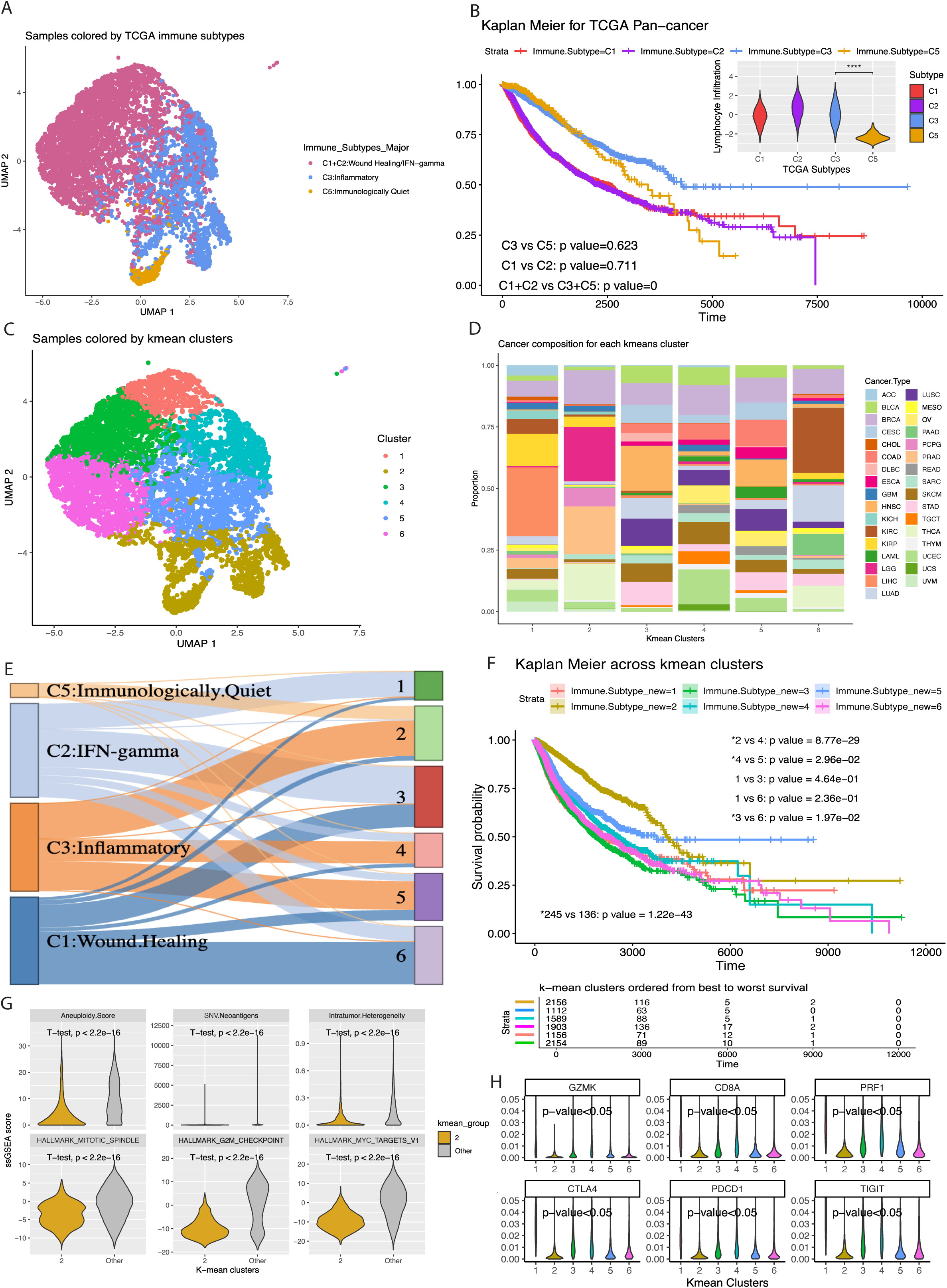
irMPs redefine immune subtypes in TCGA pan-cancer data. **(A)** UMAP of subsets of TCGA samples (n=7768) using single sample gene-set enrichment (ssGSEA) scores calculated from the 28 irMPs. Each dot is a TCGA sample colored by its immune subtypes defined by Thorsson et al. **(B)** Kaplan Meier Curves for Overall Survival between TCGA immune subtypes. Log-rank test is performed between comparisons of interests, with corresponding lymphocyte infiltration scores compared on the top right (p-value is calculated with one-way anova test). **(C)** UMAP of all TCGA samples (n=10128) using ssGSEA scores calculated from the 28 irMPs. Each dot is a TCGA sample colored by clusters identified using k-mean clustering. **(D)** Stacked plot showing the proportions of cancer types across k-mean clusters. **(E)** Sankey River plot showing the shuffling of each TCGA immune subtype into each of the k-mean clusters. **(F)** Kaplan Meier Overall Survival curves across the different k-mean clusters. Log-rank test is performed between pairs of interests (2 vs 5, 2 vs 4, 4 vs 5). Risk table is shown below with clusters ordered from best survival (top) to worst survival (bolom). **(G)** Violin plot comparing tumor intrinsic characteristics between cluster 2 and the rest of the clusters by scoring programs with ssGSEA. **(H)** Violin plot comparing the gene expression levels of 3 classical activation and exhaustion markers across the k-mean clusters.

Upon re-clustering, C3 and C5 are resolved into new clusters 2, 4 and 5 (**Figure 4e**) that well delineate patients by their overall survival (**Figure 4f**), with cluster 2 having the best overall survival. To further study the molecular characteristics of cluster 2, we assessed selected TCGA signature scores on tumor characteristics^57^ (see **Methods**) across our clusters and found that cluster 2 has significantly (p-value<0.05) lower scores, compared to other k-mean clusters in terms of aneuploidy score, neoantigen loads, intratumor heterogeneity and hallmark pathways indicative of proliferation (mitotic spindle, G2M checkpoint and MYC targets V1) (**Figure 4g**), suggesting that cluster 2 contains more indolent tumors, corroborated with the observed low cell proliferation. In addition, Cluster 2 has significantly lower expression in cytotoxic markers (*GZMK, PRF1,* and *CD8A*) and exhaustion markers (*CTLA4, PDCD1,* and *TIGIT*) (**Figure 4h**), suggesting less immunogenic environment. Cluster 2 is also enriched in Lower Grade Glioma, Prostate Adenocarcinoma, and Thyroid Carcinoma (**Figure 4d**), which are often lower grade cancers with indolent to slower growth^58^ and are less likely to invade nearby tissues or metastasize to distant organs. These tumors generally have better prognosis^59^ compared to higher-grade cancers. Therefore, the superior survival of cluster 2 is mainly driven by its simple tumor composition, low proliferation, and low immunogenicity, showing that our irMPs can also discern tumor intrinsic differences via their associated immunological states. Cluster 4 and 5 are both characterized by low tumor proliferation and high tumor development (**Figure S11b**). In term of immune landscape, cluster 4 has significantly (p-value<0.05) higher immune infiltration (Th1 cells and Gamma-Delta T cells) than cluster 2 or 5 and cluster 5 has significantly (p-value<0.05) higher level of activated NK cells compared to cluster 2 or 4 (**Figure S11b-c**).

The k-mean algorithm also redistributed C1 and C2 into cluster 1, 3 and 6 (**Figure 4e**). Specifically, we identified a transitional subtype between C1 and C2 (cluster 3 in **Figure 4c**) ^30,31^ characterized by in between expression levels of characteristic IFNG-subtype (C2) markers (*IFNG, JAK1, STAT1*)^60^ and classical wound-healing (C1) markers (*S100A1, E2F1, APC*)^61^ (**Figure S11d**). Cluster 3 has significantly (p-value<0.05) worse overall survival compared to cluster 1 and 6, respectively (**Figure S11e**), and it is characterized by high tumor proliferation, significantly (p-value<0.05) higher tumor glycolysis and oxidative phosphorylation than cluster 6 (**Figure S11b**). Cluster 1 and 6 are both characterized by high tumor proliferation (**Figure S11b**). In term of immune landscape, cluster 1 has significantly (p-value<0.05) higher infiltration of CD8 T cells and M1 Macrophages (**Figure S11c**) than the other clusters, suggesting high level of immune activation, which is related to the observed high expression of exhaustion markers^62^ (**Figure S11f**). Cluster 6 is characterized by significantly (p-value<0.05) lower hallmark immune pathway scores than other clusters (**Figure S11b**).

In summary, our irGSs redefine TCGA subtypes with significantly distinct survival patterns, unveiling not only the intrinsic traits of tumors such as their proliferation and metabolic profiles, but also the dynamic immune activities within the tumor microenvironment. Compared with the immune archetypes defined by Combes et.al^63^, we are defining differential functional states rather than cell type compositions. **Figure S11b-c** contain detailed information on hallmark pathway scores and the relative abundance of immune cell infiltration for each cluster.

### irMPs better distinguish immune checkpoint blockade (ICB) response

We investigated if the irMP expression at baseline could be used to differentiate ICB responders and non-responders. We utilized four ICB melanoma cohorts^64–67^, details in **Table 2**. To account for differences in sequencing depth and coverage, we calculated the Fragments Per Kilobase of transcript per Million mapped reads (FPKM) for each of the four RNAseq data and obtained 104 RNA samples at baseline. We calculated ssGSEA score for each RNA expression sample in the integrated ICB cohort using (1) 50 Hallmark pathways^9^, (2) 29 immune-relevant KEGG (irKEGG) pathways^10^, (3) lymphoid irMPs (L_MPs), (4) myeloid irMPs (M_MPs), or (5) lymphoid-myeloid combined irMPs. We randomly divided 104 baseline samples into 70% training (n_train_ =73) and 30% testing (n_test_=21). To avoid over-fitting, we fitted five generalized linear models (GLM) with LASSO regularization adjusting for the above-mentioned gene set variables. Using the best model selected from 10-fold cross-validation, we fitted the model on the test data and evaluated model performance by comparing classification accuracy, which is the average of sensitivity and specificity. To deal with randomness in data splitting, we performed the procedure 1000 times and computed the range for classification accuracy from each of the five models (see **Methods** and **Figure 5a**). Our gene sets resulted in the highest mean accuracy of 0.71, followed by Hallmark (0.69) and irKEGG (0.62), with most of the gene sets being negatively associated with response (**Figure S12a**). The classification accuracy is comparable (p-value=0.13) between Hallmark and combined irMPs (**Figure 5a**). However, Hallmark contains 50 gene sets with relatively large (146±67) gene set sizes, whereas the combined irMPs has only 28 gene sets with 50 genes each, possessing comparable predictive power using fewer parameters. We also compared the classification accuracies with the 18-gene tumor inflammation signature (TIS)^68^, which has been approved for use as an investigational use-only (IUO) criteria to stratify patients based on potential to respond to ICB, and the classification accuracy with TIS only on the test data is 52.83%.

**Figure 5.**
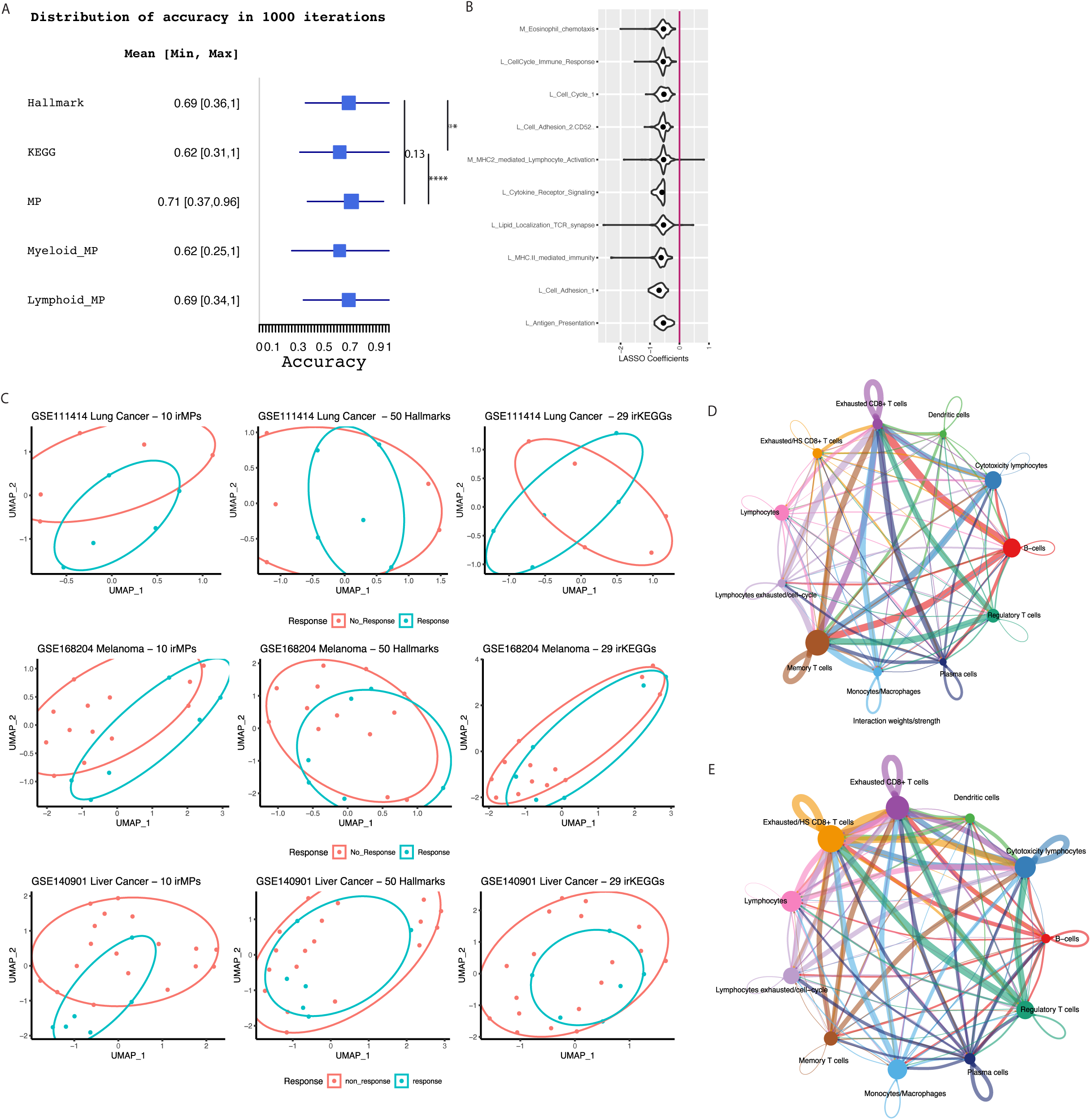
irMPs beKer separate ICB responses. **(A)** Distribution of ICB classification accuracy in each of the five model across 1000 iterations. Distribution of accuracies was compared using T test with significance level 0.05. **(B)** The average filed coefficients for top 10 selected irMPs across 1000 iterations. **(C)** UMAP derived from 10 most selected irMPs activities (len), 50 Hallmarks (middle), and 29 irKEGGs (right) in lung (top), melanoma (middle) and liver (bolom) cancer. Color represents different responses. Baseline cell-cell communications for responders **(D)** and non-responders **(E)** inferred from cell chat. Nodes are the 11 cell types annotated in the original single cell data, edges represent significant interactions (adjusted p-value<0.05) and the thickness of the edges represent the probability of interactions.

**Table 2.** Number of samples contained in each of the melanoma ICB cohort.

| GEO ID | Disease | Treatment | Pre (n) | Post (n) | Responder (n) | Non-responder (n) |
| --- | --- | --- | --- | --- | --- | --- |
| 91061 | Metastatic Melanoma | Anti PD-1 | 51 | 58 | 60 | 49 |
| 115821 | Metastatic Melanoma | Anti PD-1 | 9 | 18 | 1 | 26 |
| 78220 | Metastatic Melanoma | Anti PD-1 | 25 | 0 | 13 | 12 |
| 145996 | Metastatic Melanoma | Anti PD-1 | 14 | 0 | 9 | 5 |
| Total (n) |  |  | 99 | 76 | 83 | 92 |

To corroborate if our gene sets can better separate ICB response with baseline gene expression only, we extracted the top 10 most selected irMPs across the 1000 LASSO models (**Figure 5b**) and calculated their activities at baseline using BulkRNAseq profiled from lung cancer^69^, melanoma^70^, and liver cancer^71^. We again observed that these 10 irMPs can better separate ICB response in comparison to irKEGGs or Hallmarks (**Figure 5c**). These data indicate that the irMPs can be useful to develop tumor type agnostic general prognostic signatures to predict ICB response.

To understand what these irMPs explain, we calculated the ssGSEA for exhaustion, cytotoxicity, and tissue resident memory (T_RM_) using their respective marker gene expression (full marker list detailed in **Methods**). Computing the correlation between irMPs activity and these signature expressions at baseline (**Figure S12b**), we observed significant (FDR adjusted p-value<0.05) positive correlation between most of the irMPs and exhaustion, cytotoxicity, and T_RM_, suggesting that at baseline non-responders show phenotype of activation induced exhaustion, a result of prior antigen stimulation. To confirm our observation, we deconvoluted each of the bulk RNA sample into exhausted (T_EX_), tissue resident memory (T_RM_), tissue effector memory (T_EM_), naïve (T_N_), and effector T cells (T_EFF_) using CibersortX^72^ and pan-cancer T cell signature matrix^73^ as reference (see **Methods**). Indeed, non-responders show higher T_EX_ and T_RM_ and lower T_EM_ and T_N_ at baseline **(Figure S12c)**. However, when we fit the model with only exhaustion, cytotoxicity and T_RM_ score, the classification accuracy is only 54%, suggesting that irMPs have captured addition factors that dictate response to ICB in addition abundance of T-cell states.

As cellular communication plays an important role in orchestrating the immune response^74,75^, we hypothesize that irMPs have also captured differential cellular interactions between responders and non-responders. Utilizing an independent scRNA-seq dataset^76^ from baseline melanoma patients later treated with anti-PD1, we converted the data into pseudo-bulk expression (see **Methods**), calculated activity for selected irMPs (top 10 most frequently selected irMPs from 1000 LASSO regressions in the previous analysis), Hallmarks and irKEGGs pathways, and projected the 12 samples onto a UMAP. Once again, we observed that irMPs more effectively separated out responders and non-responders (**Figure S12d**) and these 10 irMPs are also associated with exhaustion, cytotoxicity and T_RM_ (**Figure S12e**). We then performed CellChat^77^ analysis (see **Methods**) to infer cell-cell interactions based on ligand-receptor expression and observed that responders are enriched in interactions surrounding memory T cells, while non-responders are enriched in interactions surrounding exhausted T cells at baseline (**Figure 5d**). To offer additional evidence of these differential interactions, we used scrublet^78^ to identify doublets and double-annotated the doublets (see **Methods**). Although doublets in droplet-based scRNA-seq data may result from multiple sources, they have been shown to reflect true physical interactions between cell types^79^. Responders were characterized by a limited repertoire of doublet phenotypes with a marked enrichment for doublets with memory T-cells. In contrast, non-responders presented with a more varied doublet repertoire with frequent doublets involving various exhausted lymphocytes (**Figure S12f**). These trends mirror results of the cell-cell communication analysis (**Figure 5d-e**). To confirm whether our irMPs have captured these interactions, we calculated the activities of the 10 important irMPs in each type of doublets at baseline. We observed that the top 10 selected irMPs all have high activities in doublets that are enriched in non-responders at baseline (**Figure S13**). Taken together, the irMPs may capture interactions between exhausted lymphocytes and other cell types at baseline, which have been associated with poor response in anti-PD1 immunotherapy ^80–82^.

### irMPs granularly segment spatial transcriptomics data

irMPs can potentially enhance segmentation and phenotyping of spatially resolved transcriptomic (SRT) data. For illustration, we obtained 10x Visium data from a FFPE human breast tissue section^83^. The H&E slide from the tissue section was segmented by pathologists into major cell types (**Figure 6a**). We calculated the activity scores of the irMPs from the SRT sample and observed that irMPs consistently isolate out the tumor regions (dark blue), aligning with pathologist’s annotation (**Figure 6b, Figure S14**). Beside identifying tumor regions, our irMPs can also be characterized by the infiltrated immunological patterns on the H&E image. We used CellTrek ^84^ to project single cells from a reference breast cancer patient profiled by scRNA-seq in Tumor Immune Single-cell Hub 2 (TISCH2)^85^ onto the spatial spots ^84^ (see **Methods**)^83^. We then visualized the average irMP activity score for each projected cell type in **Figure 6c** using a clustered dot plot and observed that clustering of irMPs was largely driven by lineages (3 clusters from top to bottom): myeloid, lymphoid, and cycling. On one hand, some irMPs are strongly lineage specific. For example, the "TCR anchoring" score is prominent among T cells, nearly absent in other cell types (**Figure 6c**). On the other hand, some irMPs also provide functional insight across the TIME, not limited to a cell type. For example, "IFN-gamma induced cytotoxicity" exhibits the highest score in T cells and a notable score in macrophages (**Figure 6c**), suggesting a co-enforcement feedback loop between activated T/NK cells and cytotoxic macrophages mediated by potent interferon gamma, a hub gene in the PPI network (**Figure S7.17**), which aligns with the annotation of this irMP. Benchmarking with immune-related KEGG pathways and Hallmarks, we observed that Hallmark pathways are mostly specific to cancer and endothelial regions (**Figure S15a**) and irKEGG pathways are myeloid-specific (**Figure S15b**). To examine the generalizability of our results, we repeated above experiments on an FFPE human ovarian tumor tissue and an FFPE human intestine tumor tissue using ovarian cancer patients^86^ and colorectal cancer^87^ patients as single cell references, respectively. We obtained coherent results (**Figure S16**). Together, we found the irMPs can be applied to not only segment spatial transcriptomic data aligning with standard pathological segmentation but also reveal granular spot-level cell type information and multicellular processes in TIME.

**Figure 6.**
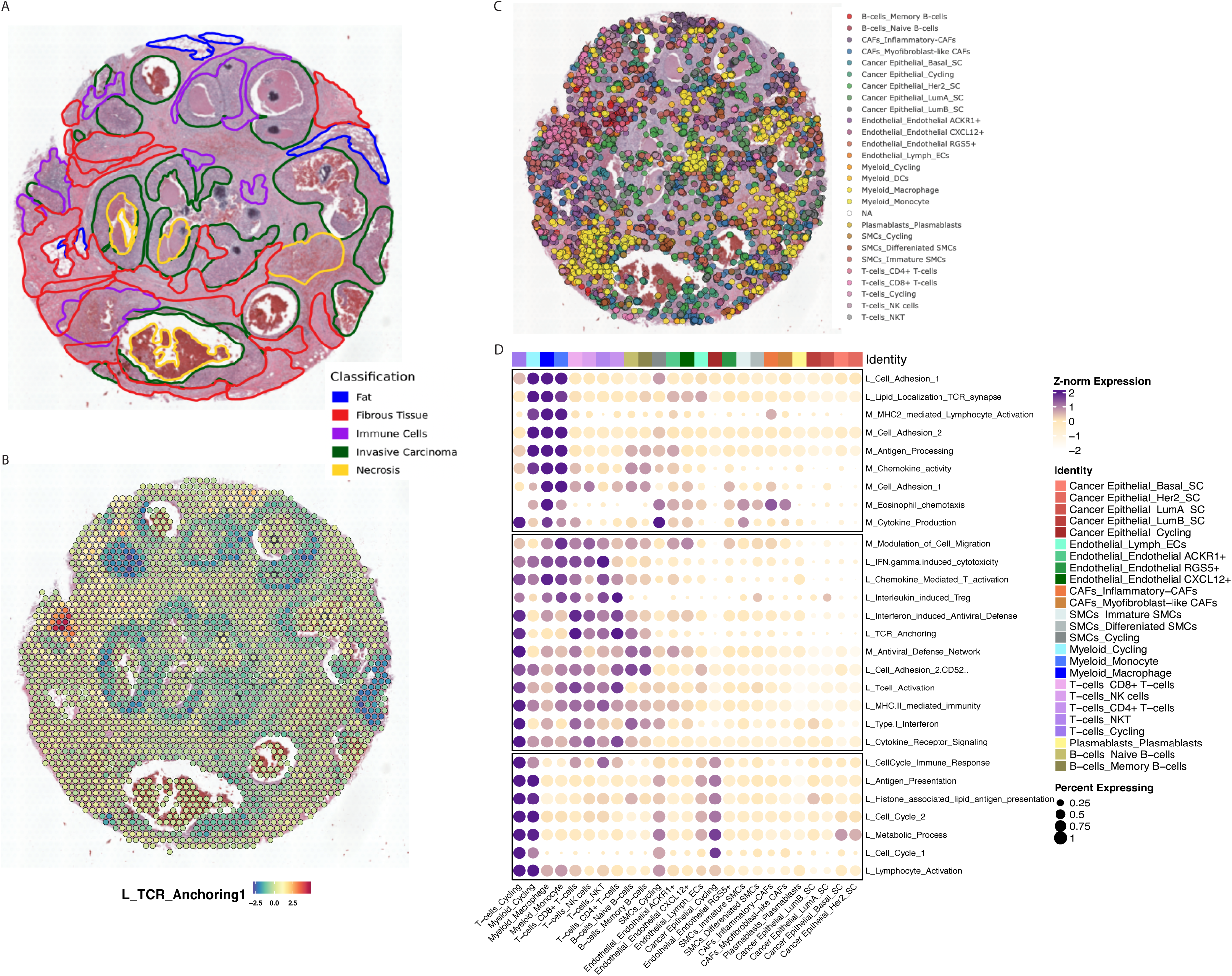
irMPs granularly segment spa&al transcriptomics data. **(A)** Pathologist-annotated 10x Genomics-acquired FFPE human breast tissue spatial image with colored-boundaries separating different cell types **(B)** Module score overlaid on spatial spots for L_MP2 (TCR anchoring). **(C)** CellTrek projected single cells onto the breast tumor tissue spatial slide. Color represents different cell types (Important compartments are Myeloid (yellow), Lymphoid (pink/red) and Tumor (green) **(D)** Dot plot comparing the z-transformed expression level of 28 irMPs across different celltrek-annotated cell types. Color represents the average expression level, and the dot size represents the percentage of cells expressing the irMP activity.

To further assess how gene sets explain spatial heterogeneity and identify niches with distinct functions, we referred back to **Figure 6d** and observed that TCR_anchoring shows high activities in regions dominated by B and T cells. We further plotted the activity of TCR_anchoring on intestine, breast, and ovarian cancer Visium data and observed that, in each of the three tumor slides, there is a region with TCR_anchoring activity (**Figure 7a-c left panel**) that is significantly higher than the rest of the spots (**Figure 7a-c right panel**). To further define these regions, a board-certified pathologist (Sharia. H) performed pathological review from respectively matched H&E images and concluded that the identified regions corresponded to a tertiary lymphoid structure (TLS) and two lymphoid aggregates (LA), respectively (**Figure 7a-c middle panel**). TLS has recently gained attention in cancer research as their presence in the TIME often reflect active and local immune response against the tumor, contributing to favorable prognosis^88,89^. LA, on the other hand, is often considered the precursor to TLS^89^. Therefore, the strong association between TCR_anchoring scores and TLS/LA regions demonstrated the potential of applying our irMPs to delineate spatial heterogeneity and identify functionally active regions.

**Figure 7.**
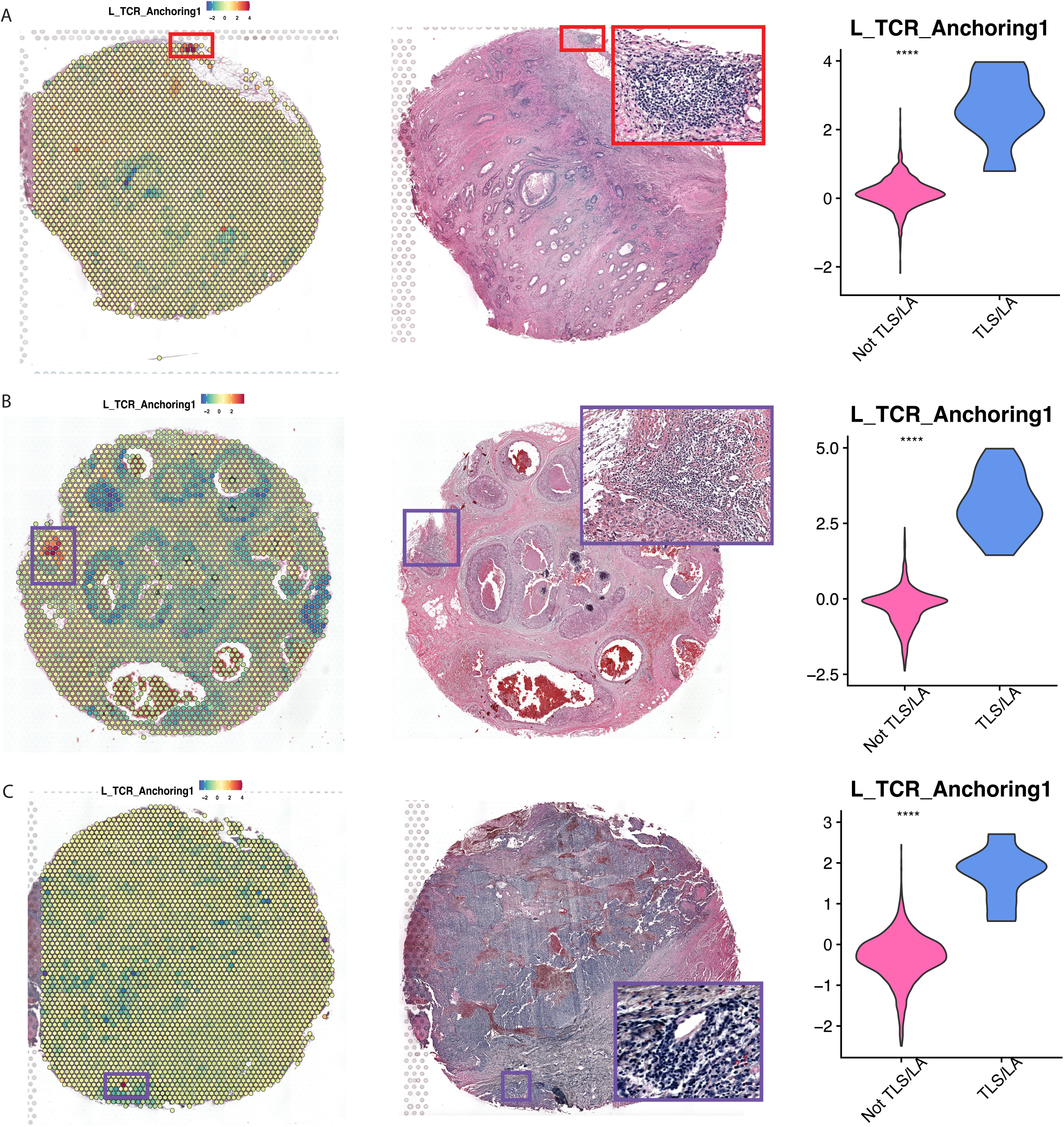
L_MP2 associates with lymphoid aggregates and ter&ary lymphoid structure. Gene set activity for L_MP2 (len), H&E image with pathologist-confirmed lymphoid aggregates (middle), and statistical test for gene set activities comparing TLS/LA spots vs other spots (right) for **(A)** intestine cancer, **(B)** breast cancer, and **(C)** ovarian cancer. (red boxes are confirmed TLS, purple boxes are LA with insufficient evidence to be TLS)

## Discussion

Our study aims to address the lack of objectively defined immune-specific gene sets in cancer immunology research. We developed these gene sets based on different studies representing immune cell involvement from bulk samples challenged with infections and immune perturbations. We further utilized CITE-seq and other orthogonal data to validate the immunological functions of the novel immune-related meta programs (irMPs) we identified in our study. Most importantly, we demonstrated that these transcriptional programs, derived from high-throughput experiments, can provide valuable insights into various aspects of cancer immunological research. Specifically, (1) irMPs improved tumor immune subtype clustering and identified new subtypes with distinct survival outcomes, enhancing our understanding of immune heterogeneity, (2) irMPs demonstrated higher accuracy in separating ICB response, highlighting their potential for predicting ICB outcomes by providing insights into baseline cytotoxicity and cellular communications and (3) irMPs offered enhanced granularity in delineating tumor-immune boundaries and niches in spatial transcriptomic data. These findings suggest promising translational applications in diverse immunological contexts, as these irMPs are originated from non-cancerous experiments, implying common immunological mechanisms shared by cancer and other pathophysiological conditions.

Our work also shows the importance of cross-lineage coordination and the interconnectivity between immune and tumor compartment. By leveraging BulkRNAseq data, we have discovered transcriptional programs that span across multiple cell lineages. These observations are important as they underpin the intricate web of cellular communications, orchestrating functional phenotypes crucial for mounting effective immune responses. By unraveling these shared transcriptional signatures across diverse cell populations, this research enhances our fundamental understanding of signaling cascades within the immune system. Communication between tumor and immune cells was demonstrated in **Figure 4f-h**, when samples grouped based on immune activities also show differences in tumor-specific characteristics. There is a complex bidirectional dialogue between immune cells and cancer cells, thus the activities of irMPs provides valuable insights into the nature of the tumor, its aggressiveness, and even its susceptibility to treatments like immunotherapy. Therefore, studying immune transcriptional programs offers a window into understanding the broader dynamics of tumor biology and devising strategies to harness the immune system’s potential in combating cancer.

Our work serves as a parallel effort to the Cross-tissue immune atlas by Teichmann group, in which they leveraged scRNAseq dataset and developed CellTypist^91^ to classify immune cells into 101 distinct populations with context-dependent functional states across various human healthy tissues. In a similar vein, our work leveraged BulkRNAseq and concentrate on elucidating immune activities in diseased samples. Future studies are warranted to (1) analyze the immune landscape differences between healthy and diseased microenvironment and (2) leverage scRNAseq to derive cell type specific transcriptional programs.

Algorithms designed to enhance our comprehension of molecular-level functional states are crucial, offering abundant information unattainable through individual laboratory experiments. By integrating extensive data sources, we often capture a broader spectrum of functional states under varying conditions, offering insights into numerous biological inquiries. For instance, recent research^92^ linking metabolic fitness to the resistance of Chimeric Antigen Receptor-engineered Natural Killer (CAR-NK) cells underscores the importance of comprehending cellular characteristics in deciphering therapeutic resistance. In this study, we demonstrated the NMF can be used to efficiently output functional gene sets with translational implications. Overall, our study constructed 28 immune-specific gene sets, taking a significant step toward advancing our comprehension of TIME. This knowledge can empower cancer immunologists to gain deeper insights into immune checkpoint blockade response and cancer survival, potentially expediting the development of novel immunotherapies and biomarkers, and ultimately improving patient outcomes.

## Limitations

In this study, we constructed gene sets using extensive bulk RNA data, offering valuable insights into the molecular landscape. We divided the data by lymphoid and myeloid lineages. While that may have enhanced the discovery of lineage-specific irGSs, it may bias against discovery of irGSs shared between the lineages. Another inherent limitation in our work, as well as studies of similar nature^9,12,16,93,94^ is the challenge of annotating the functional roles of each gene sets. To address this limitation and enhance the reliability of our findings, we emphasize the importance of employing various functional assays for validation. One promising avenue is the utilization of targeted perturb-seq experiments, which can provide experimental evidence to corroborate our gene set annotations. As a matter of fact, there are already *in silico* experiments to predict transcriptional profiles from combinatorial perturbations^95^, and such predictions, if successfully validated in wet lab, can further bolster our functional annotations. Another promising avenue is to apply these gene sets on scRNA data derived in different contexts, thereby enhancing annotations with higher granularity and context-dependent information. By applying gene sets to single cell data, we can precisely assess their functional relevance in distinct cellular contexts and unveil novel insights into the regulation of biological processes at the single-cell level.

## Author Contributions

Writing – Original Draft: S.H

Conceptualization: S.H, V.M, K.C and K.R

Methodology: S.H, V.M, H.R, and R.B

Visualization: S.H

Writing – Review & Editing: S.H, V.M, K.C, M.D, C.H, MM.G, and W.P

Suggestions: X.J, Q.L, C.H, Y.T, K.K, C.H, K.K, K.R, Sharia.H, L.S and ML.G

Supervision: V.M and K.C.

## Acknowledgements

This project has been made possible in part by grant U01CA247760 and U01CA281902 to KC and the Cancer Center Support Grant P30 CA016672 to PP from National Cancer Institute. This project was also partially supported by the MD Anderson Moonshot programs; MMG is a Cancer Prevention and Research Institute of Texas (CPRIT) Scholar in Cancer Research and is supported by CPRIT (Recruitment of First-Time Tenure-Track Faculty Members; RR190017). We also thanks Dr. Chloé Villani for their helpful comments.

## STAR Methods

### RESOURCE AVAILABILITY

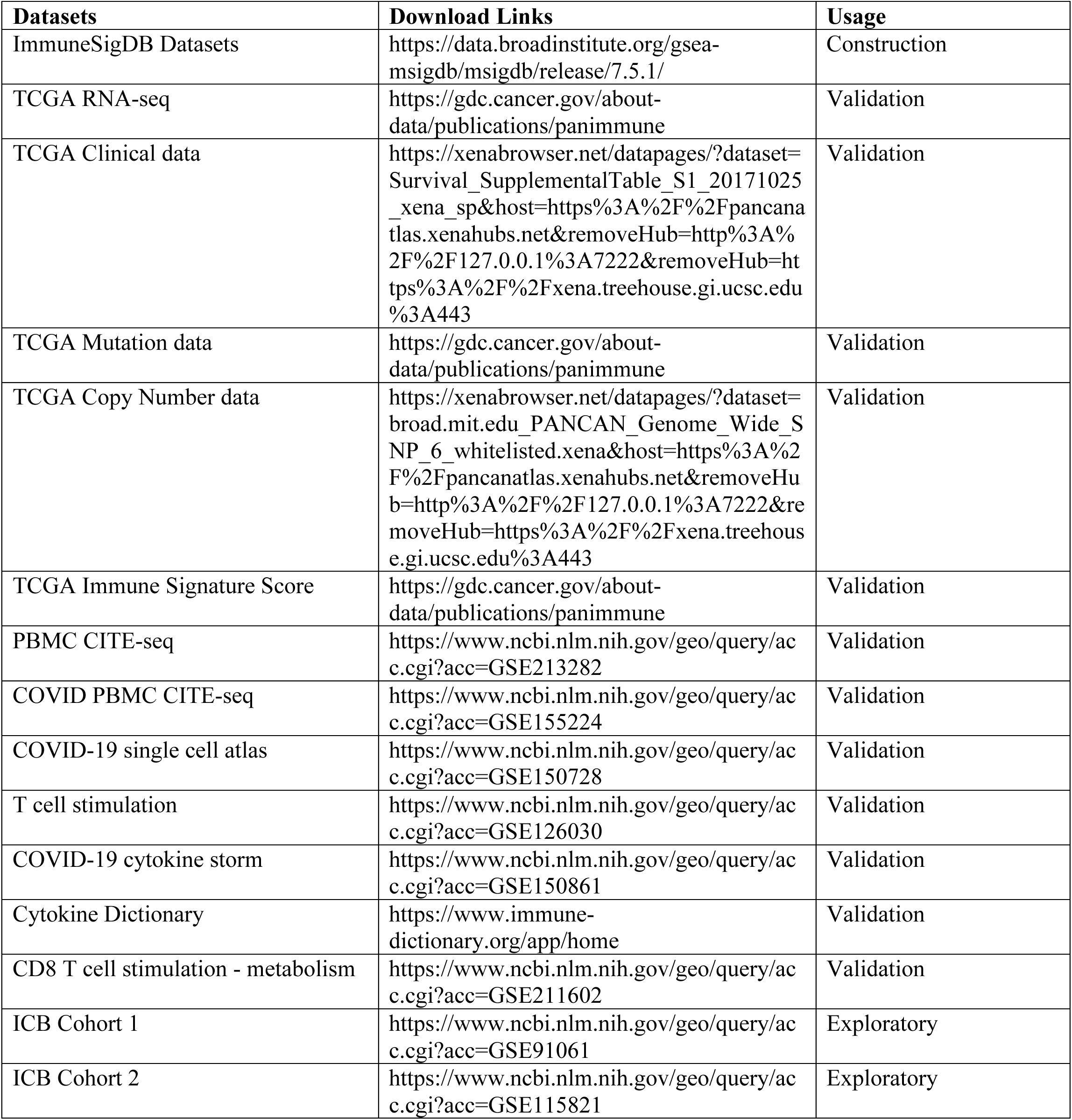
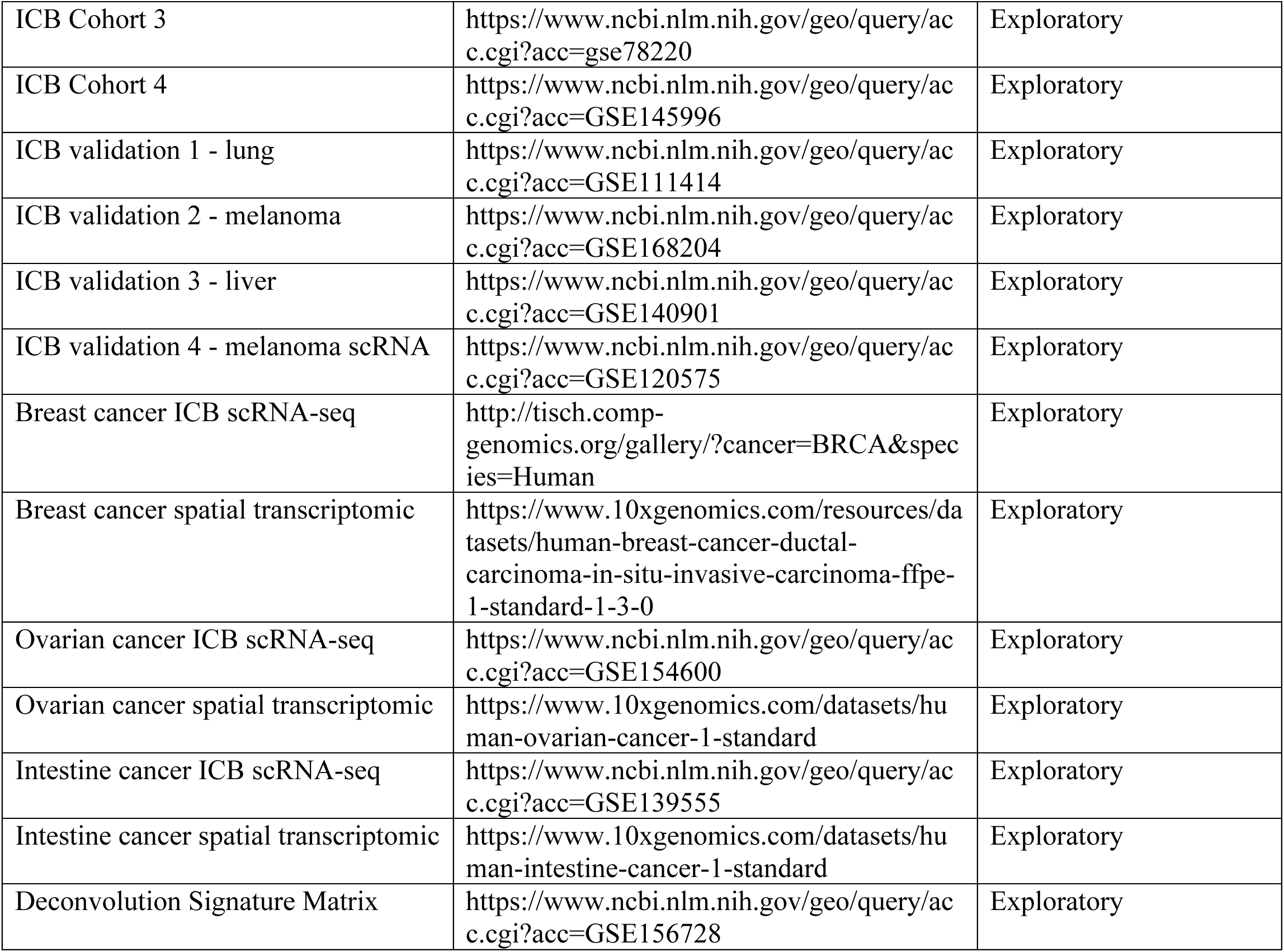

| **Datasets** | **Download Links** | **Usage** |
| --- | --- | --- |
| ImmuneSigDB Datasets | https://data.broadinstitute.org/gsea-msigdb/msigdb/release/7.5.1/ | Construction |
| TCGA RNA-seq | https://gdc.cancer.gov/about-data/publications/panimmune | Validation |
| TCGA Clinical data | https://xenabrowser.net/datapages/?dataset=Survival_SupplementalTable_S1_20171025_xena_sp&host=https%3A%2F%2Fpancanatlas.xenahubs.net&removeHub=http%3A%2F%2F127.0.0.1%3A7222&removeHub=https%3A%2F%2Fxena.treehouse.gi.ucsc.edu%3A443 | Validation |
| TCGA Mutation data | https://gdc.cancer.gov/about-data/publications/panimmune | Validation |
| TCGA Copy Number data | https://xenabrowser.net/datapages/?dataset=broad.mit.edu_PANCAN_Genome_Wide_SNP_6_whitelisted.xena&host=https%3A%2F%2Fpancanatlas.xenahubs.net&removeHub=http%3A%2F%2F127.0.0.1%3A7222&removeHub=https%3A%2F%2Fxena.treehouse.gi.ucsc.edu%3A443 | Validation |
| TCGA Immune Signature Score | https://gdc.cancer.gov/about-data/publications/panimmune | Validation |
| PBMC CITE-seq | https://www.ncbi.nlm.nih.gov/geo/query/acc.cgi?acc=GSE213282 | Validation |
| COVID PBMC CITE-seq | https://www.ncbi.nlm.nih.gov/geo/query/acc.cgi?acc=GSE155224 | Validation |
| COVID-19 single cell atlas | https://www.ncbi.nlm.nih.gov/geo/query/acc.cgi?acc=GSE150728 | Validation |
| T cell stimulation | https://www.ncbi.nlm.nih.gov/geo/query/acc.cgi?acc=GSE126030 | Validation |
| COVID-19 cytokine storm | https://www.ncbi.nlm.nih.gov/geo/query/acc.cgi?acc=GSE150861 | Validation |
| Cytokine Dictionary | https://www.immune-dictionary.org/app/home | Validation |
| CD8 T cell stimulation - metabolism | https://www.ncbi.nlm.nih.gov/geo/query/acc.cgi?acc=GSE211602 | Validation |
| ICB Cohort 1 | https://www.ncbi.nlm.nih.gov/geo/query/acc.cgi?acc=GSE91061 | Exploratory |
| ICB Cohort 2 | https://www.ncbi.nlm.nih.gov/geo/query/acc.cgi?acc=GSE115821 | Exploratory |
| ICB Cohort 3 | https://www.ncbi.nlm.nih.gov/geo/query/acc.cgi?acc=gse78220 | Exploratory |
| ICB Cohort 4 | https://www.ncbi.nlm.nih.gov/geo/query/acc.cgi?acc=GSE145996 | Exploratory |
| ICB validation 1 - lung | https://www.ncbi.nlm.nih.gov/geo/query/acc.cgi?acc=GSE111414 | Exploratory |
| ICB validation 2 - melanoma | https://www.ncbi.nlm.nih.gov/geo/query/acc.cgi?acc=GSE168204 | Exploratory |
| ICB validation 3 - liver | https://www.ncbi.nlm.nih.gov/geo/query/acc.cgi?acc=GSE140901 | Exploratory |
| ICB validation 4 - melanoma scRNA | https://www.ncbi.nlm.nih.gov/geo/query/acc.cgi?acc=GSE120575 | Exploratory |
| Breast cancer ICB scRNA-seq | http://tisch.comp-genomics.org/gallery/?cancer=BRCA&species=Human | Exploratory |
| Breast cancer spatial transcriptomic | https://www.10xgenomics.com/resources/datasets/human-breast-cancer-ductal-carcinoma-in-situ-invasive-carcinoma-ffpe-1-standard-1-3-0 | Exploratory |
| Ovarian cancer ICB scRNA-seq | https://www.ncbi.nlm.nih.gov/geo/query/acc.cgi?acc=GSE154600 | Exploratory |
| Ovarian cancer spatial transcriptomic | https://www.10xgenomics.com/datasets/human-ovarian-cancer-1-standard | Exploratory |
| Intestine cancer ICB scRNA-seq | https://www.ncbi.nlm.nih.gov/geo/query/acc.cgi?acc=GSE139555 | Exploratory |
| Intestine cancer spatial transcriptomic | https://www.10xgenomics.com/datasets/human-intestine-cancer-1-standard | Exploratory |
| Deconvolution Signature Matrix | https://www.ncbi.nlm.nih.gov/geo/query/acc.cgi?acc=GSE156728 | Exploratory |

#### Code Availability

Codes to generate the figures and tables presented in the paper can be accessed through https://github.com/chloehe1129/immune-hallmark

### METHOD DETAILS

#### Bulk RNA-seq Data Curation

We referenced the 389 immunology relevant studies identified in the ImmuneSigDB publication^12^ and curated the corresponding available bulk RNA-seq datasets deposited in NCBI Gene Expression Omnibus (GEO)^96^. Bulk RNA-seq datasets sourced from human samples with more than 4 individuals were kept for downstream analysis. Since we focused on immune pathways represented in these datasets, mitochondria genes and ribosomal genes were removed. In addition, we grouped the datasets based on broad immune lineages^97^. As a result, 47 datasets sourced from lymphoid lineages and 36 datasets sourced from myeloid lineages were kept for further analysis.

#### “Data-driven” Statistical Analysis

We performed the following analysis for each of the 47 lymphoid dataset and 36 myeloid datasets. Non-negative matrix factorization (NMF)^15^ is a common approach to establish genes that have coordinated expression or consistently opposite expression pattern that underpin the data. We performed NMF on each of the qualifying dataset with number of latent factors K = 4, 5, 6, and each NMF program is composed of genes with top 50 loadings from one latent factor. We column-combined the loadings of 529 NMF programs and normalized them. To ensure that the programs are not only non-redundant but also generalizable, we curated robust NMF programs^16^, defined as programs that (1) would reoccur across datasets (have at least 20% overlapping genes with at least one NMF program derived from outside of the current data) and (2) is non-redundant within the same dataset (have less than 20% overlapping genes with all NMF programs derived from the current dataset). To further reduce redundancy, we iteratively merged and updated the robust NMF programs into meta programs (MP) based on algorithm proposed by Tirosh et al^16^: we initiated the process by selecting two robust NMF programs with the greatest gene overlap. Combining these two robust NMF programs, we generated a fresh gene set consisting of 50 genes, comprising of both the common genes present in both programs and selected genes unique to each program. The selected genes were chosen from all the unique genes arranged in descending order of their loadings to ensuring that the gene set was populated to its full capacity of 50 genes. We then selected another robust NMF programs with the highest overlap with this fresh gene set and repeated the same process until the overlap is < 5, in which we restarted the process by selecting two robust NMF programs with the greatest gene overlap.

#### Functional Annotation

We performed over-representation enrichment analysis (ORA)^98^ for each MP using KEGG^10^ pathways, Hallmarks^9^, and biological process terms from GO^11^. ORA was performed with “clusterProfiler” package^99^. Annotation for each MP was determined by highly enriched terms based on core enrichment count and False Discovery Rate (FDR) adjusted p-value < 0.05^100^. The annotations were then manually reviewed and refined by immunologists from MD Anderson Cancer Center.

#### Protein-protein interaction network for each MP

We overlaid the genes from each gene set onto protein-protein interaction (PPI) network from string database^23^ using *STRINGdb* package in R.

#### Novelty Validation

We calculated the Jaccard distance between our gene sets with Hallmark, KEGG and GO to validate the novelty of our gene sets.

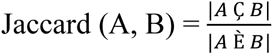

Significance of each pairwise Jaccard distance was computed by randomly choosing gene set of the same length 1000 times and p-value was calculated as the number of times a permutated jaccard distance is larger than the actual jaccard distance divided by 1000. Multiple testing was adjusted using FDR adjusted p-values.

#### Cell type annotation for CITE-seq

We performed cell type annotation using *SingleR*^37^ package in R with CHETAH^101^ as reference.

#### Cell dichotomization in CITE-seq

We dichotomized cells into two groups based on their RNA activity levels of each irMP of interest using Gaussian Mixture Models^102^ with G=2 using *mclust* package in R and compared the relevant protein abundance between the irMP-high and the irMP-low groups with two sample t-test.

#### Gene set scoring

We performed ssGSEA using *ssgsea* function in the *corto* package in R with min.size=0. We performed GSDensity when scoring cell cycle and metabolism activities using *gsdensity* package in R.

#### Antigen-related pathways validation

Persistent tumor mutational burden (pTMB) was calculated for each sample in TCGA, as the total number of single copy mutations and multiple copy mutations^51^. We grouped samples based on pTMB, dichotomized at mean. We performed Differential gene expression (DGE) analysis between samples with high and low pTMB using function DESeq2^103^. Results from DGE analysis with the whole transcriptome were further analyzed in Gene set enrichment analysis (GSEA) with the 19 lymphoid MPs using function *GSEA*^104^ from the *clusterProfiler* package.

#### Cytotoxicity pathway validation

We calculated the IFN.gamma-induced cytotoxicity (L_MP17) score and exhaustion score based on GS Density^105^. Exhaustion was calculated using the expression of the following genes: *TIGIT, TOX, PDCD1, CD160, CD244, CTLA4, BTLA, HAVCR2, and LAG3*.

#### TCGA immune subtypes refinement

We calculated the single sample gene set enrichment score (ssGSEA score)^27^ for each of the MP across all patient samples in TCGA pan-cancer data and projected the samples onto a two-dimensional space through UMAP using *umap* function. We colored each sample with corresponding TCGA immune subtype information. In addition, we fitted Kaplan Meier (KM) survival curves for each of the four TCGA immune subtypes. We also performed k-mean clustering with *kmean* function using the ssGSEA matrix with the value of K=6 selected by low within-cluster sum of square and fitted KM survival curves for each identified cluster.

#### ICB response model

We calculated ssGSEA score for each patient from the combined ICB studies^106–108^ using (1) 50 MSigDB Hallmark pathways, (2) 29 immune-related KEGG pathways (3) lymphoid-derived irMPs, (4) myeloid-derived irMPs and (5) lymphoid-myeloid combined irMPs. We fitted five generalized linear model with LASSO regularization using *glmnet* with binary response variable adjusting for each set of the above-mentioned gene sets using 70% of the full data as training data. In cases where ICB response is defined in more than two categories, we grouped complete and partial responder into responders and stable and progressive into non-responders. Using the best model selected from 10-fold cross-validation, we fitted the model on the test data and evaluated model performance by comparing classification accuracy. Classification accuracy is defined as the average of sensitivity and specificity. To deal with randomness in data splitting, we performed the procedure 1000 times and computed a range of classification accuracy from each of the five models.

#### Marker genes for immune signatures

Exhaustion: *PDCD1, TIGIT, HAVCR2, CTLA4, LAG3*

Cytotoxicity: *NKG7, CCL4, CST7, PRF1, GZMA, GZMB, IFNG, CCL3*

Tissue resident memory: *ITGAE, ZNF683, ITGA1, CD69, CXCR6, CXCL13, PDCD1*

Immune signatures were scored based on ssGSEA

#### Pseudo-bulk aggregation

scRNA-seq data was converted into pseudo-bulk gene expression by using the function *AggregateExpression* with group.by = "sample".

#### Cell type deconvolution

We used breast cancer infiltrated T cells subtypes annotated by Zheng et al^73^ to deconvolute the bulk RNA samples using CIBERSORTx^109^ with the following configuration: B-mode batch correction and 50 permutations for significance calculation.

#### Cell Chat Analysis

We performed cell chat analysis using the *CellChat* package in R.

#### Doublet detection and annotation

We used *scrublet* to calculate a doublet score for each cell and identified doublets with top 5% doublet score. We used *FindAllMarkers* with adjusted p-value<0.05 and log fold change>0.25 to define differentially expressed genes for each cell type and calculate the cell type ssGSEA for each doublet. We double-annotated each doublet with the top two positive scoring cell types.

#### Single cell level annotation on spatial transcriptomic

We calculated the irMPs score on each spatial spot using *AddModuleScore* function in R. We performed *CellTrek* to achieve single-cell level spatial cell type mapping on breast cancer spatial transcriptomic (ST) data with an independent scRNAseq (SC) data as reference. Default arguments were used for *celltrek* function in R.

#### scRNAseq data processing

All single cell RNA sequencing data were processed using standard Seurat procedure implemented by *Seurat* package in R. We performed the following procedures for all scRNAseq data: *NormalizeData(), FindVariableGenes(), ScaleData(), RunPCA(), FindNeighbors(),FindClusters()* and *RunUMAP()* with 10 PCs.

#### Differential gene expression and pathway analysis

For single cell data, we used *FindAllMarkers()* to identify differentially expressed genes. For BulkRNAseq data, we used *Deseq2* to identify differentially expressed genes. Pathway enrichment analysis was performed using *clusterProfiler* package in R. For gene set enrichment, statistical significance was determined by a rank-based test, in which the enrichment score (ES) reflects the degree to which genes in a set are over-represented at the extremes of a ranked list, utilizing a Kolmogorov–Smirnov-like statistic. Then the statistical significance of the ES is determined through a permutation test based on phenotypic labels, generating a null distribution for comparison to ascertain the gene set’s dependency on these labels.

### QUANTIFICATION AND STATISTICAL ANALYSIS

All statistical analysis were performed in R version 4.2.1 and Seurat version 4.3.0. Differences between continuous distributions were tested using Standardized t test in case of two variables or ANOVA in cases of more than two variables. Multiple testing correction for false discovery was performed using False Discovery Rate (FDR) with function *p. adjust*, and statistical significance is defined at FDR adjusted p-value < 0.05.

#### Lead Contact and Materials Availability

Further information and requests for resources and reagents should be directed to and will be fulfilled by the Lead Contact, Ken Chen. This study did not generate new unique reagents

