## Supplemental material for "Elucidating immune-related gene transcriptional programs via factorization of large-scale RNA-profiles"

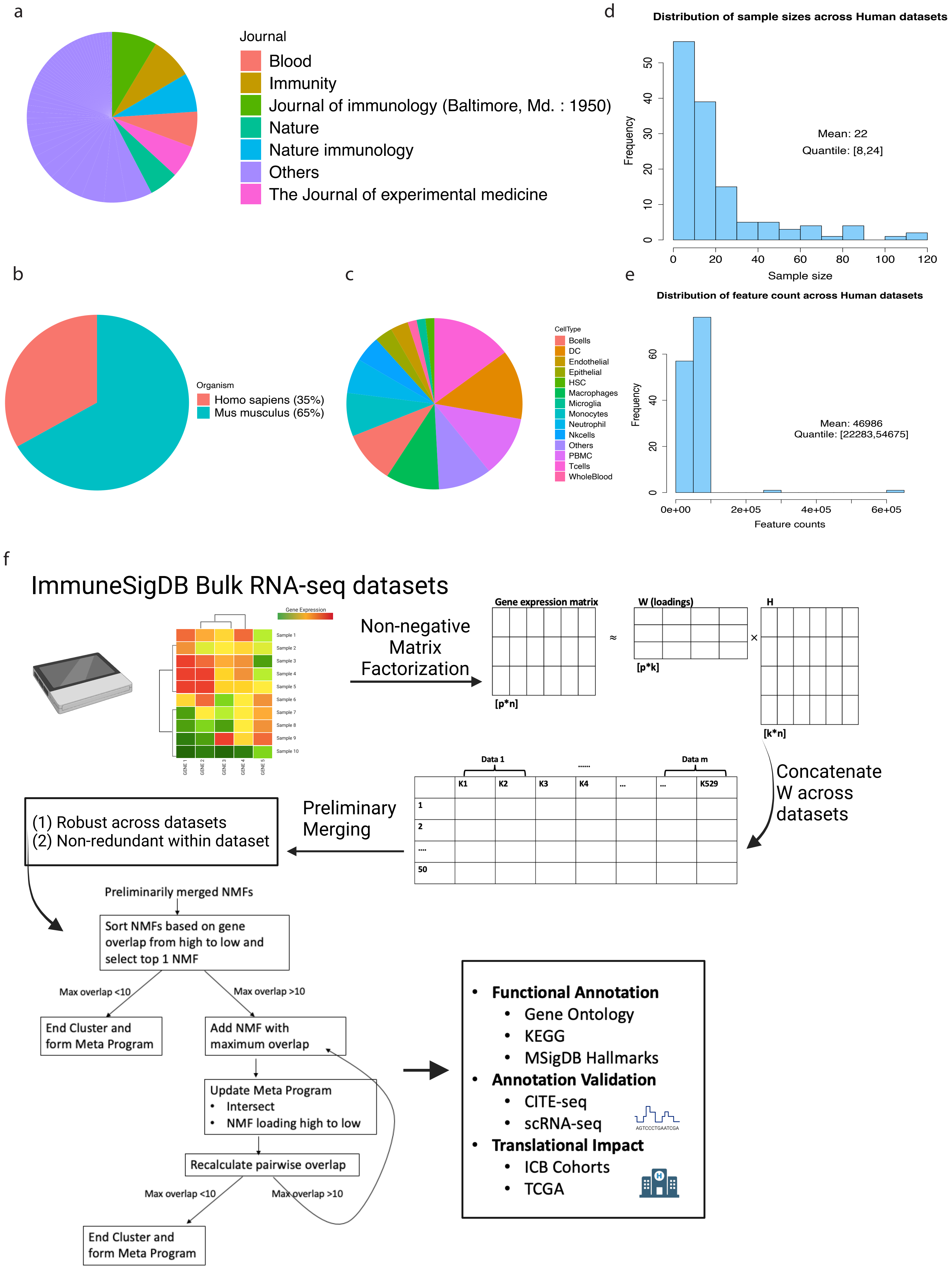

a
Journal
Blood
Immunity
Journal of immunology (Baltimore, Md. : 1950)
Nature
Nature immunology
Others
The Journal of experimental medicine
d
Distribution of sample sizes across Human datasets
Frequency
Sample size
Mean: 22
Quantile: [8,24]
0 20 40 60 80 100 120
0 10 20 30 40 50
b
Organism
Homo sapiens (35%)
Mus musculus (65%)
c
CellType
Bcells
DC
Endothelial
Epithelial
HSC
Macrophages
Microglia
Monocytes
Neutrophil
Nkcells
Others
PBMC
Tcells
WholeBlood
e
Distribution of feature count across Human datasets
Frequency
Feature counts
Mean: 46986
Quantile: [22283,54675]
0e+00 2e+05 4e+05 6e+05
0 20 40 60
f
ImmuneSigDB Bulk RNA-seq datasets
Gene Expression
Sample 1
Sample 2
Sample 3
Sample 4
Sample 5
Sample 6
Sample 7
Sample 8
Sample 9
Sample 10
GENE 1
GENE 2
GENE 3
GENE 4
GENE 5
Non-negative Matrix Factorization
Gene expression matrix
[p*n]
≈
W (loadings)
[p*k]
×
H
[k*n]
Concatenate W across datasets
Data 1
......
Data m
K1
K2
K3
K4
...
...
K529
1
2
....
50
Preliminary Merging
(1) Robust across datasets
(2) Non-redundant within dataset
Preliminarily merged NMFs
Sort NMFs based on gene overlap from high to low and select top 1 NMF
Max overlap <10
Max overlap >10
End Cluster and form Meta Program
Add NMF with maximum overlap
Update Meta Program
• Intersect
• NMF loading high to low
Recalculate pairwise overlap
Max overlap <10
Max overlap >10
End Cluster and form Meta Program
• Functional Annotation
• Gene Ontology
• KEGG
• MSigDB Hallmarks
• Annotation Validation
• CITE-seq
• scRNA-seq
AGTCCCTGAATCGA
• Translational Impact
• ICB Cohorts
• TCGA

**Figure S1. [Analysis workflow], Related to Figure 1.**
**(a) ImmuneSigDB datasets compartments: (a)** Pie chart summarizing the source journal where the corresponding datasets have been published. **(b)** Pie chart summarizing the proportion of species that the datasets are derived from. **(c)** Pie chart summarizing the proportion of sorted cell types from human datasets. **(d)** Distribution of sample sizes across human datasets. **(e)** Distribution of feature counts across human datasets. **(f) Gene set construction analysis workflow:** We downloaded 389 immunology relevant studies and performed NMF on each of the qualifying human dataset. We curated robust NMF programs and merged them into meta gene set. We performed over-representation enrichment analysis for each meta gene set using KEGG pathways, Hallmarks, and biological process terms from GO. Annotation for each meta gene set was determined by highly enriched terms based on count and False Discovery Rate (FDR) adjusted p-value. We sought relevant datasets to validate the annotations and propose translational utilities of these gene sets.

Overlap between Lymphoid MPs and spectra pathways

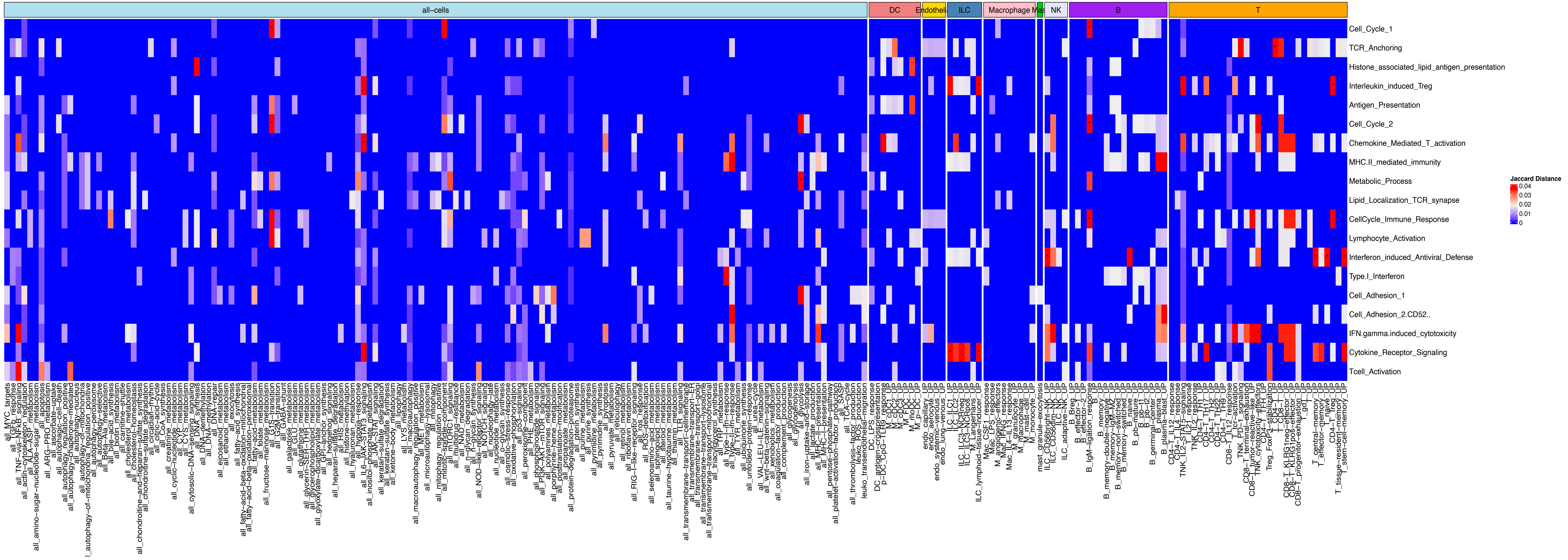


Overlap between Myeloid MPs and spectra pathways

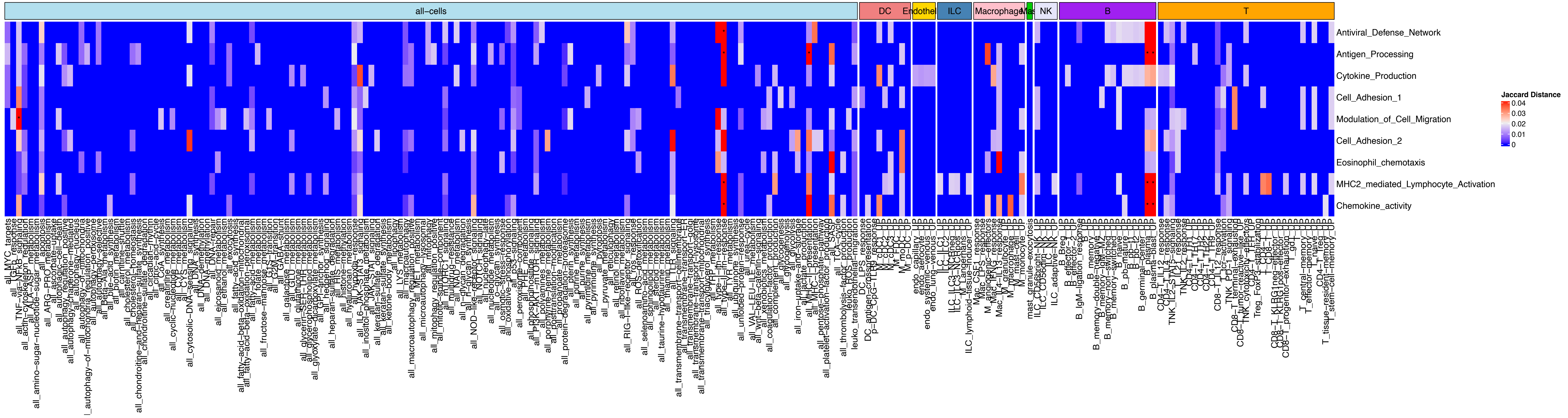

**Figure S2. [Gene sets comparison with Spectra], Related to Figure 1.**
Pairwise jaccard distance between Lymphoid irMPs (top)/Myeloid irMPs (bottom) and 231 spectra gene sets. Columns are separated into gene sets based on their cell type assignment. Pairwise comparisons with adjusted p-value<0.05 are overlaid with asterisks.

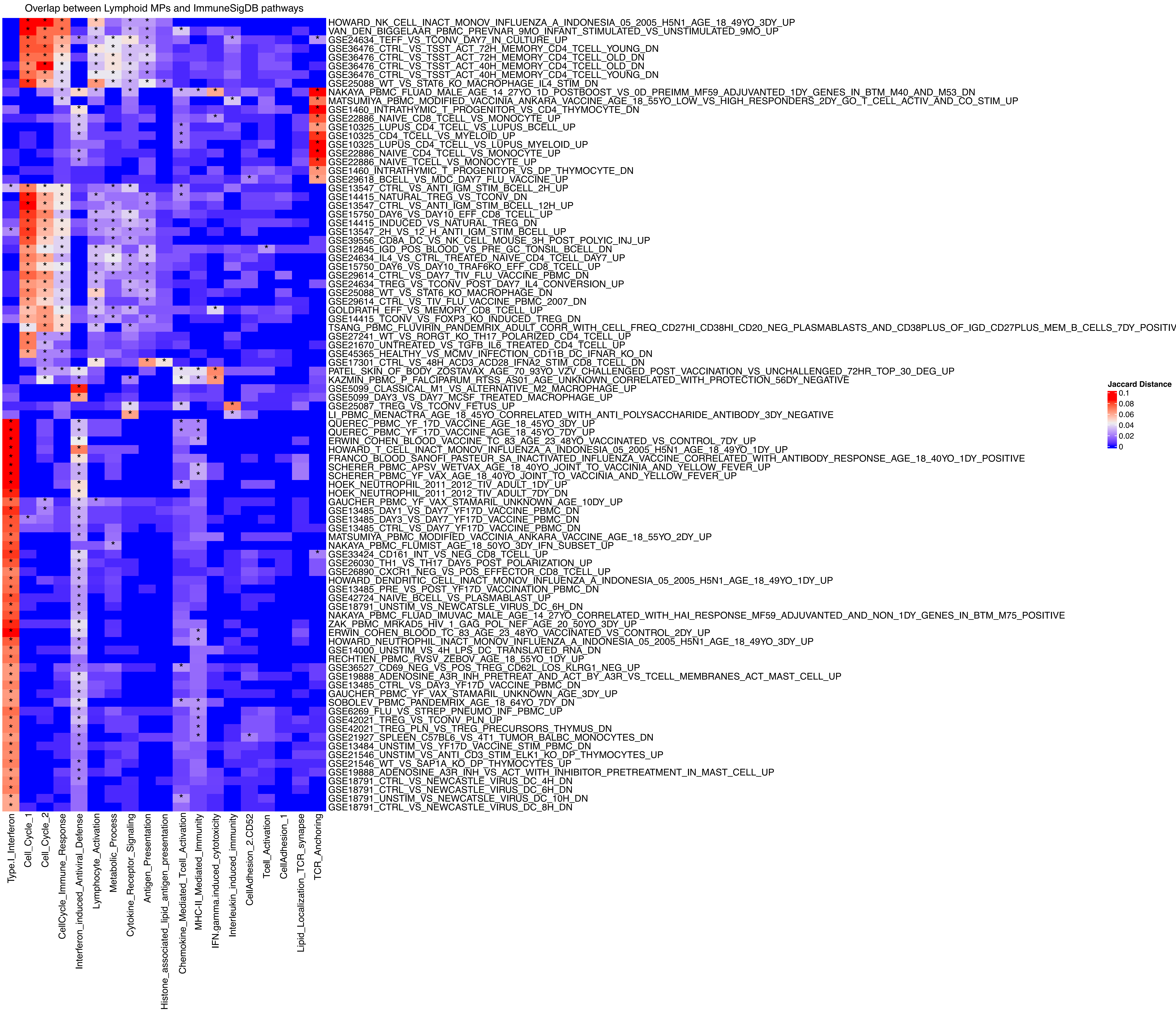

Overlap between Lymphoid MPs and ImmuneSigDB pathways
HOWARD_NK_CELL_INACT_MONOV_INFLUENZA_A_INDONESIA_05_2005_H5N1_AGE_18_49YO_3DY_UP
VAN_DEN_BIGGELAAR_PBMC_PREVNAR_9MO_INFANT_STIMULATED_VS_UNSTIMULATED_9MO_UP
GSE24634_TEFF_VS_TCONV_DAY7_IN_CULTURE_UP
GSE36476_CTRL_VS_TSST_ACT_72H_MEMORY_CD4_TCELL_YOUNG_DN
GSE36476_CTRL_VS_TSST_ACT_72H_MEMORY_CD4_TCELL_OLD_DN
GSE36476_CTRL_VS_TSST_ACT_40H_MEMORY_CD4_TCELL_OLD_DN
GSE36476_CTRL_VS_TSST_ACT_40H_MEMORY_CD4_TCELL_YOUNG_DN
GSE25088_WT_VS_STAT6_KO_MACROPHAGE_IL4_STIM_DN
NAKAYA_PBMC_FLUAD_MALE_AGE_14_27YO_1D_POSTBOOST_VS_0D_PREIMM_MF59_ADJUVANTED_1DY_GENES_IN_BTM_M40_AND_M53_DN
MATSUMIYA_PBMC_MODIFIED_VACCINIA_ANKARA_VACCINE_AGE_18_55YO_LOW_VS_HIGH_RESPONDERS_2DY_GO_T_CELL_ACTIV_AND_CO_STIM_UP
GSE1460_INTRATHYMIC_T_PROGENITOR_VS_CD4_THYMOCYTE_DN
GSE22886_NAIVE_CD8_TCELL_VS_MONOCYTE_UP
GSE10325_LUPUS_CD4_TCELL_VS_LUPUS_BCELL_UP
GSE10325_CD4_TCELL_VS_MYELOID_UP
GSE10325_LUPUS_CD4_TCELL_VS_LUPUS_MYELOID_UP
GSE22886_NAIVE_CD4_TCELL_VS_MONOCYTE_UP
GSE22886_NAIVE_TCELL_VS_MONOCYTE_UP
GSE1460_INTRATHYMIC_T_PROGENITOR_VS_DP_THYMOCYTE_DN
GSE29618_BCELL_VS_MDC_DAY7_FLU_VACCINE_UP
GSE13547_CTRL_VS_ANTI_IGM_STIM_BCELL_2H_UP
GSE14415_NATURAL_TREG_VS_TCONV_DN
GSE13547_CTRL_VS_ANTI_IGM_STIM_BCELL_12H_UP
GSE15750_DAY6_VS_DAY10_EFF_CD8_TCELL_UP
GSE14415_INDUCED_VS_NATURAL_TREG_DN
GSE13547_2H_VS_12_H_ANTI_IGM_STIM_BCELL_UP
GSE39556_CD8A_DC_VS_NK_CELL_MOUSE_3H_POST_POLYIC_INJ_UP
GSE12845_IGD_POS_BLOOD_VS_PRE_GC_TONSIL_BCELL_DN
GSE24634_IL4_VS_CTRL_TREATED_NAIVE_CD4_TCELL_DAY7_UP
GSE15750_DAY6_VS_DAY10_TRAF6KO_EFF_CD8_TCELL_UP
GSE29614_CTRL_VS_DAY7_TIV_FLU_VACCINE_PBMC_DN
GSE24634_TREG_VS_TCONV_POST_DAY7_IL4_CONVERSION_UP
GSE25088_WT_VS_STAT6_KO_MACROPHAGE_DN
GSE29614_CTRL_VS_TIV_FLU_VACCINE_PBMC_2007_DN
GOLDRATH_EFF_VS_MEMORY_CD8_TCELL_UP
GSE14415_TCONV_VS_FOXP3_KO_INDUCED_TREG_DN
TSANG_PBMC_FLUVIRIN_PANDEMRIX_ADULT_CORR_WITH_CELL_FREQ_CD27HI_CD38HI_CD20_NEG_PLASMABLASTS_AND_CD38PLUS_OF_IGD_CD27PLUS_MEM_B_CELLS_7DY_POSITIV
GSE27241_WT_VS_RORGT_KO_TH17_POLARIZED_CD4_TCELL_UP
GSE21670_UNTREATED_VS_TGFB_IL6_TREATED_CD4_TCELL_UP
GSE45365_HEALTHY_VS_MCMV_INFECTION_CD11B_DC_IFNAR_KO_DN
GSE17301_CTRL_VS_48H_ACD3_ACD28_IFNA2_STIM_CD8_TCELL_DN
PATEL_SKIN_OF_BODY_ZOSTAVAX_AGE_70_93YO_VZV_CHALLENGED_POST_VACCINATION_VS_UNCHALLENGED_72HR_TOP_30_DEG_UP
KAZMIN_PBMC_P_FALCIPARUM_RTSS_AS01_AGE_UNKNOWN_CORRELATED_WITH_PROTECTION_56DY_NEGATIVE
GSE5099_CLASSICAL_M1_VS_ALTERNATIVE_M2_MACROPHAGE_UP
GSE5099_DAY3_VS_DAY7_MCSF_TREATED_MACROPHAGE_UP
GSE25087_TREG_VS_TCONV_FETUS_UP
LI_PBMC_MENACTRA_AGE_18_45YO_CORRELATED_WITH_ANTI_POLYSACCHARIDE_ANTIBODY_3DY_NEGATIVE
QUEREC_PBMC_YF_17D_VACCINE_AGE_18_45YO_3DY_UP
QUEREC_PBMC_YF_17D_VACCINE_AGE_18_45YO_7DY_UP
ERWIN_COHEN_BLOOD_VACCINE_TC_83_AGE_23_48YO_VACCINATED_VS_CONTROL_7DY_UP
HOWARD_T_CELL_INACT_MONOV_INFLUENZA_A_INDONESIA_05_2005_H5N1_AGE_18_49YO_1DY_UP
FRANCO_BLOOD_SANOFI_PASTEUR_SA_INACTIVATED_INFLUENZA_VACCINE_CORRELATED_WITH_ANTIBODY_RESPONSE_AGE_18_40YO_1DY_POSITIVE
SCHERER_PBMC_APSV_WETVAX_AGE_18_40YO_JOINT_TO_VACCINIA_AND_YELLOW_FEVER_UP
SCHERER_PBMC_YF_VAX_AGE_18_40YO_JOINT_TO_VACCINIA_AND_YELLOW_FEVER_UP
HOEK_NEUTROPHIL_2011_2012_TIV_ADULT_1DY_UP
HOEK_NEUTROPHIL_2011_2012_TIV_ADULT_7DY_DN
GAUCHER_PBMC_YF_VAX_STAMARIL_UNKNOWN_AGE_10DY_UP
GSE13485_DAY1_VS_DAY7_YF17D_VACCINE_PBMC_DN
GSE13485_DAY3_VS_DAY7_YF17D_VACCINE_PBMC_DN
GSE13485_CTRL_VS_DAY7_YF17D_VACCINE_PBMC_DN
MATSUMIYA_PBMC_MODIFIED_VACCINIA_ANKARA_VACCINE_AGE_18_55YO_2DY_UP
NAKAYA_PBMC_FLUMIST_AGE_18_50YO_3DY_IFN_SUBSET_UP
GSE33424_CD161_INT_VS_NEG_CD8_TCELL_UP
GSE26030_TH1_VS_TH17_DAY5_POST_POLARIZATION_UP
GSE26890_CXCR1_NEG_VS_POS_EFFECTOR_CD8_TCELL_UP
HOWARD_DENDRITIC_CELL_INACT_MONOV_INFLUENZA_A_INDONESIA_05_2005_H5N1_AGE_18_49YO_1DY_UP
GSE13485_PRE_VS_POST_YF17D_VACCINATION_PBMC_DN
GSE42724_NAIVE_BCELL_VS_PLASMABLAST_UP
GSE18791_UNSTIM_VS_NEWCATSLE_VIRUS_DC_6H_DN
NAKAYA_PBMC_FLUAD_IMUVAC_MALE_AGE_14_27YO_CORRELATED_WITH_HAI_RESPONSE_MF59_ADJUVANTED_AND_NON_1DY_GENES_IN_BTM_M75_POSITIVE
ZAK_PBMC_MRKAD5_HIV_1_GAG_POL_NEF_AGE_20_50YO_3DY_UP
ERWIN_COHEN_BLOOD_TC_83_AGE_23_48YO_VACCINATED_VS_CONTROL_2DY_UP
HOWARD_NEUTROPHIL_INACT_MONOV_INFLUENZA_A_INDONESIA_05_2005_H5N1_AGE_18_49YO_3DY_UP
GSE14000_UNSTIM_VS_4H_LPS_DC_TRANSLATED_RNA_DN
RECHTIEN_PBMC_RVSV_ZEBOV_AGE_18_55YO_1DY_UP
GSE36527_CD69_NEG_VS_POS_TREG_CD62L_LOS_KLRG1_NEG_UP
GSE19888_ADENOSINE_A3R_INH_PRETREAT_AND_ACT_BY_A3R_VS_TCELL_MEMBRANES_ACT_MAST_CELL_UP
GSE13485_CTRL_VS_DAY3_YF17D_VACCINE_PBMC_DN
GAUCHER_PBMC_YF_VAX_STAMARIL_UNKNOWN_AGE_3DY_UP
SOBOLEV_PBMC_PANDEMRIX_AGE_18_64YO_7DY_DN
GSE6269_FLU_VS_STREP_PNEUMO_INF_PBMC_UP
GSE42021_TREG_VS_TCONV_PLN_UP
GSE42021_TREG_PLN_VS_TREG_PRECURSORS_THYMUS_DN
GSE21927_SPLEEN_C57BL6_VS_4T1_TUMOR_BALBC_MONOCYTES_DN
GSE13484_UNSTIM_VS_YF17D_VACCINE_STIM_PBMC_DN
GSE21546_UNSTIM_VS_ANTI_CD3_STIM_ELK1_KO_DP_THYMOCYTES_UP
GSE21546_WT_VS_SAP1A_KO_DP_THYMOCYTES_UP
GSE19888_ADENOSINE_A3R_INH_VS_ACT_WITH_INHIBITOR_PRETREATMENT_IN_MAST_CELL_UP
GSE18791_CTRL_VS_NEWCASTLE_VIRUS_DC_4H_DN
GSE18791_CTRL_VS_NEWCASTLE_VIRUS_DC_6H_DN
GSE18791_UNSTIM_VS_NEWCATSLE_VIRUS_DC_10H_DN
GSE18791_CTRL_VS_NEWCASTLE_VIRUS_DC_8H_DN
Type.I_Interferon
Cell_Cycle_1
Cell_Cycle_2
CellCycle_Immune_Response
Interferon_induced_Antiviral_Defense
Lymphocyte_Activation
Metabolic_Process
Cytokine_Receptor_Signaling
Antigen_Presentation
Histone_associated_lipid_antigen_presentation
Chemokine_Mediated_Tcell_Activation
MHC-II_Mediated_Immunity
IFN.gamma.induced_cytotoxicity
Interleukin_induced_immunity
CellAdhesion_2.CD52
Tcell_Activation
CellAdhesion_1
Lipid_Localization_TCR_synapse
TCR_Anchoring
Jaccard Distance
0.1
0.08
0.06
0.04
0.02
0

**Figure S3. [Gene sets comparison with ImmuneSigDB], Related to Figure 1.**
Pairwise jaccard distance between Lymphoid irMPs and selected ImmuneSigDB gene sets. Pairwise comparisons with adjusted p-value<0.05 are overlaid with asterisks. We select ImmuneSigDB based on the following criteria (1) at least one significant overlap with Lymphoid irMPs and (2) such overlap > 0.06.

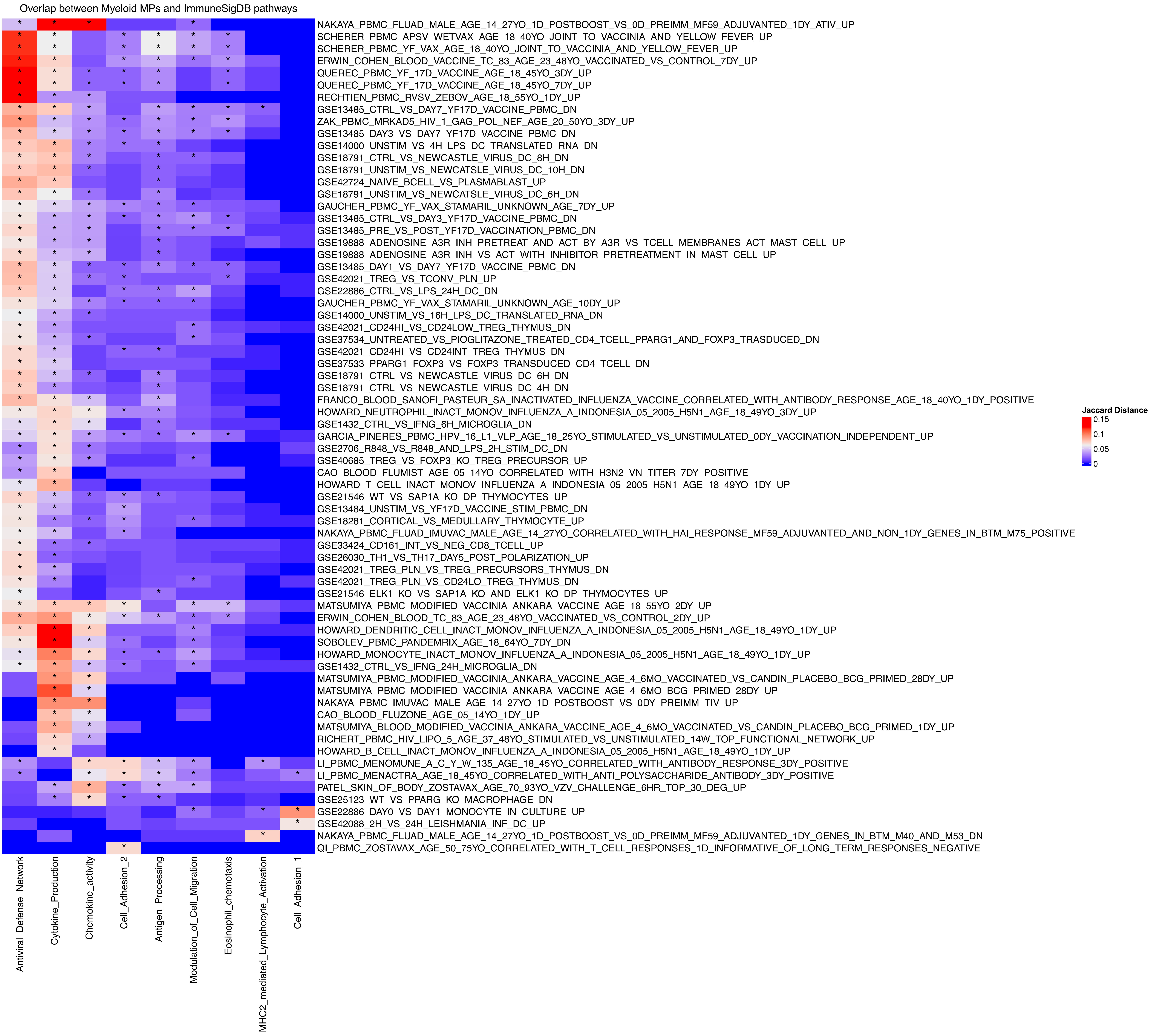

Overlap between Myeloid MPs and ImmuneSigDB pathways
NAKAYA_PBMC_FLUAD_MALE_AGE_14_27YO_1D_POSTBOOST_VS_0D_PREIMM_MF59_ADJUVANTED_1DY_ATIV_UP
SCHERER_PBMC_APSV_WETVAX_AGE_18_40YO_JOINT_TO_VACCINIA_AND_YELLOW_FEVER_UP
SCHERER_PBMC_YF_VAX_AGE_18_40YO_JOINT_TO_VACCINIA_AND_YELLOW_FEVER_UP
ERWIN_COHEN_BLOOD_VACCINE_TC_83_AGE_23_48YO_VACCINATED_VS_CONTROL_7DY_UP
QUEREC_PBMC_YF_17D_VACCINE_AGE_18_45YO_3DY_UP
QUEREC_PBMC_YF_17D_VACCINE_AGE_18_45YO_7DY_UP
RECHTIEN_PBMC_RVSV_ZEBOV_AGE_18_55YO_1DY_UP
GSE13485_CTRL_VS_DAY7_YF17D_VACCINE_PBMC_DN
ZAK_PBMC_MRKAD5_HIV_1_GAG_POL_NEF_AGE_20_50YO_3DY_UP
GSE13485_DAY3_VS_DAY7_YF17D_VACCINE_PBMC_DN
GSE14000_UNSTIM_VS_4H_LPS_DC_TRANSLATED_RNA_DN
GSE18791_CTRL_VS_NEWCASTLE_VIRUS_DC_8H_DN
GSE18791_UNSTIM_VS_NEWCATSLE_VIRUS_DC_10H_DN
GSE42724_NAIVE_BCELL_VS_PLASMABLAST_UP
GSE18791_UNSTIM_VS_NEWCATSLE_VIRUS_DC_6H_DN
GAUCHER_PBMC_YF_VAX_STAMARIL_UNKNOWN_AGE_7DY_UP
GSE13485_CTRL_VS_DAY3_YF17D_VACCINE_PBMC_DN
GSE13485_PRE_VS_POST_YF17D_VACCINATION_PBMC_DN
GSE19888_ADENOSINE_A3R_INH_PRETREAT_AND_ACT_BY_A3R_VS_TCELL_MEMBRANES_ACT_MAST_CELL_UP
GSE19888_ADENOSINE_A3R_INH_VS_ACT_WITH_INHIBITOR_PRETREATMENT_IN_MAST_CELL_UP
GSE13485_DAY1_VS_DAY7_YF17D_VACCINE_PBMC_DN
GSE42021_TREG_VS_TCONV_PLN_UP
GSE22886_CTRL_VS_LPS_24H_DC_DN
GAUCHER_PBMC_YF_VAX_STAMARIL_UNKNOWN_AGE_10DY_UP
GSE14000_UNSTIM_VS_16H_LPS_DC_TRANSLATED_RNA_DN
GSE42021_CD24HI_VS_CD24LOW_TREG_THYMUS_DN
GSE37534_UNTREATED_VS_PIOGLITAZONE_TREATED_CD4_TCELL_PPARG1_AND_FOXP3_TRASDUCED_DN
GSE42021_CD24HI_VS_CD24INT_TREG_THYMUS_DN
GSE37533_PPARG1_FOXP3_VS_FOXP3_TRANSDUCED_CD4_TCELL_DN
GSE18791_CTRL_VS_NEWCASTLE_VIRUS_DC_6H_DN
GSE18791_CTRL_VS_NEWCASTLE_VIRUS_DC_4H_DN
FRANCO_BLOOD_SANOFI_PASTEUR_SA_INACTIVATED_INFLUENZA_VACCINE_CORRELATED_WITH_ANTIBODY_RESPONSE_AGE_18_40YO_1DY_POSITIVE
HOWARD_NEUTROPHIL_INACT_MONOV_INFLUENZA_A_INDONESIA_05_2005_H5N1_AGE_18_49YO_3DY_UP
GSE1432_CTRL_VS_IFNG_6H_MICROGLIA_DN
GARCIA_PINERES_PBMC_HPV_16_L1_VLP_AGE_18_25YO_STIMULATED_VS_UNSTIMULATED_0DY_VACCINATION_INDEPENDENT_UP
GSE2706_R848_VS_R848_AND_LPS_2H_STIM_DC_DN
GSE40685_TREG_VS_FOXP3_KO_TREG_PRECURSOR_UP
CAO_BLOOD_FLUMIST_AGE_05_14YO_CORRELATED_WITH_H3N2_VN_TITER_7DY_POSITIVE
HOWARD_T_CELL_INACT_MONOV_INFLUENZA_A_INDONESIA_05_2005_H5N1_AGE_18_49YO_1DY_UP
GSE21546_WT_VS_SAP1A_KO_DP_THYMOCYTES_UP
GSE13484_UNSTIM_VS_YF17D_VACCINE_STIM_PBMC_DN
GSE18281_CORTICAL_VS_MEDULLARY_THYMOCYTE_UP
NAKAYA_PBMC_FLUAD_IMUVAC_MALE_AGE_14_27YO_CORRELATED_WITH_HAI_RESPONSE_MF59_ADJUVANTED_AND_NON_1DY_GENES_IN_BTM_M75_POSITIVE
GSE33424_CD161_INT_VS_NEG_CD8_TCELL_UP
GSE26030_TH1_VS_TH17_DAY5_POST_POLARIZATION_UP
GSE42021_TREG_PLN_VS_TREG_PRECURSORS_THYMUS_DN
GSE42021_TREG_PLN_VS_CD24LO_TREG_THYMUS_DN
GSE21546_ELK1_KO_VS_SAP1A_KO_AND_ELK1_KO_DP_THYMOCYTES_UP
MATSUMIYA_PBMC_MODIFIED_VACCINIA_ANKARA_VACCINE_AGE_18_55YO_2DY_UP
ERWIN_COHEN_BLOOD_TC_83_AGE_23_48YO_VACCINATED_VS_CONTROL_2DY_UP
HOWARD_DENDRITIC_CELL_INACT_MONOV_INFLUENZA_A_INDONESIA_05_2005_H5N1_AGE_18_49YO_1DY_UP
SOBOLEV_PBMC_PANDEMRIX_AGE_18_64YO_7DY_DN
HOWARD_MONOCYTE_INACT_MONOV_INFLUENZA_A_INDONESIA_05_2005_H5N1_AGE_18_49YO_1DY_UP
GSE1432_CTRL_VS_IFNG_24H_MICROGLIA_DN
MATSUMIYA_PBMC_MODIFIED_VACCINIA_ANKARA_VACCINE_AGE_4_6MO_VACCINATED_VS_CANDIN_PLACEBO_BCG_PRIMED_28DY_UP
MATSUMIYA_PBMC_MODIFIED_VACCINIA_ANKARA_VACCINE_AGE_4_6MO_BCG_PRIMED_28DY_UP
NAKAYA_PBMC_IMUVAC_MALE_AGE_14_27YO_1D_POSTBOOST_VS_0DY_PREIMM_TIV_UP
CAO_BLOOD_FLUZONE_AGE_05_14YO_1DY_UP
MATSUMIYA_BLOOD_MODIFIED_VACCINIA_ANKARA_VACCINE_AGE_4_6MO_VACCINATED_VS_CANDIN_PLACEBO_BCG_PRIMED_1DY_UP
RICHERT_PBMC_HIV_LIPO_5_AGE_37_48YO_STIMULATED_VS_UNSTIMULATED_14W_TOP_FUNCTIONAL_NETWORK_UP
HOWARD_B_CELL_INACT_MONOV_INFLUENZA_A_INDONESIA_05_2005_H5N1_AGE_18_49YO_1DY_UP
LI_PBMC_MENOMUNE_A_C_Y_W_135_AGE_18_45YO_CORRELATED_WITH_ANTIBODY_RESPONSE_3DY_POSITIVE
LI_PBMC_MENACTRA_AGE_18_45YO_CORRELATED_WITH_ANTI_POLYSACCHARIDE_ANTIBODY_3DY_POSITIVE
PATEL_SKIN_OF_BODY_ZOSTAVAX_AGE_70_93YO_VZV_CHALLENGE_6HR_TOP_30_DEG_UP
GSE25123_WT_VS_PPARG_KO_MACROPHAGE_DN
GSE22886_DAY0_VS_DAY1_MONOCYTE_IN_CULTURE_UP
GSE42088_2H_VS_24H_LEISHMANIA_INF_DC_UP
NAKAYA_PBMC_FLUAD_MALE_AGE_14_27YO_1D_POSTBOOST_VS_0D_PREIMM_MF59_ADJUVANTED_1DY_GENES_IN_BTM_M40_AND_M53_DN
QI_PBMC_ZOSTAVAX_AGE_50_75YO_CORRELATED_WITH_T_CELL_RESPONSES_1D_INFORMATIVE_OF_LONG_TERM_RESPONSES_NEGATIVE
Antiviral_Defense_Network
Cytokine_Production
Chemokine_activity
Cell_Adhesion_2
Antigen_Processing
Modulation_of_Cell_Migration
Eosinophil_chemotaxis
MHC2_mediated_Lymphocyte_Activation
Cell_Adhesion_1
Jaccard Distance
0.15
0.1
0.05
0

**Figure S4. [Gene sets comparison with ImmuneSigDB], Related to Figure 1.**

Pairwise jaccard distance between Myeloid irMPs and selected ImmuneSigDB gene sets. Pairwise comparisons with adjusted p-value<0.05 are overlaid with asterisks. We select ImmuneSigDB based on the following criteria (1) at least one significant overlap with Myeloid irMPs and (2) such overlap > 0.06.

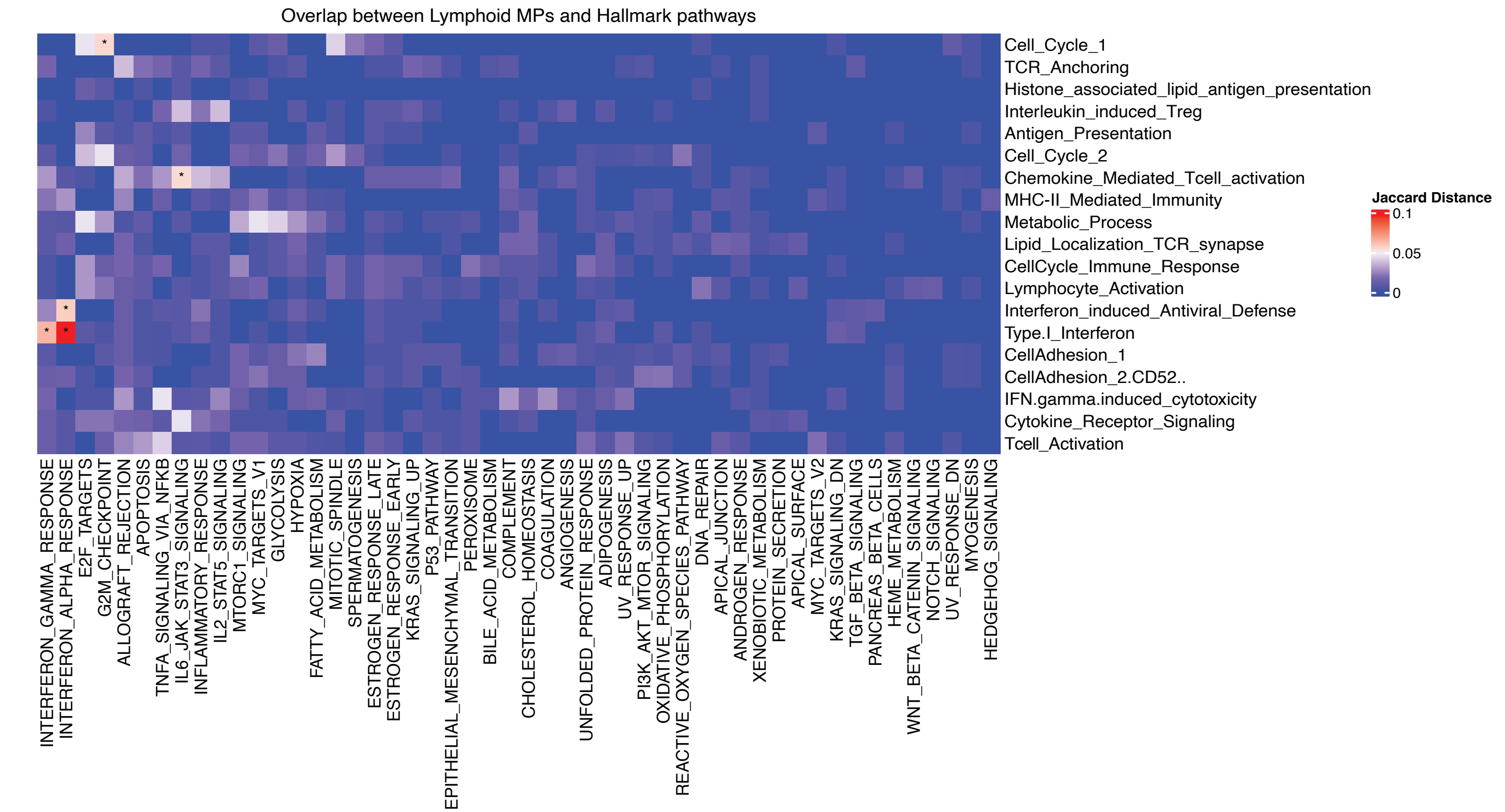
Overlap between Lymphoid MPs and Hallmark pathways
Cell_Cycle_1
TCR_Anchoring
Histone_associated_lipid_antigen_presentation
Interleukin_induced_Treg
Antigen_Presentation
Cell_Cycle_2
Chemokine_Mediated_Tcell_activation
MHC-II_Mediated_Immunity
Metabolic_Process
Lipid_Localization_TCR_synapse
CellCycle_Immune_Response
Lymphocyte_Activation
Interferon_induced_Antiviral_Defense
Type.I_Interferon
CellAdhesion_1
CellAdhesion_2.CD52..
IFN.gamma.induced_cytotoxicity
Cytokine_Receptor_Signaling
Tcell_Activation
Jaccard Distance
0.1
0.05
0
INTERFERON_GAMMA_RESPONSE
INTERFERON_ALPHA_RESPONSE
E2F_TARGETS
G2M_CHECKPOINT
ALLOGRAFT_REJECTION
APOPTOSIS
TNFA_SIGNALING_VIA_NFKB
IL6_JAK_STAT3_SIGNALING
INFLAMMATORY_RESPONSE
IL2_STAT5_SIGNALING
MTORC1_SIGNALING
MYC_TARGETS_V1
GLYCOLYSIS
HYPOXIA
FATTY_ACID_METABOLISM
MITOTIC_SPINDLE
SPERMATOGENESIS
ESTROGEN_RESPONSE_LATE
ESTROGEN_RESPONSE_EARLY
KRAS_SIGNALING_UP
P53_PATHWAY
EPITHELIAL_MESENCHYMAL_TRANSITION
PEROXISOME
BILE_ACID_METABOLISM
COMPLEMENT
CHOLESTEROL_HOMEOSTASIS
COAGULATION
ANGIOGENESIS
UNFOLDED_PROTEIN_RESPONSE
ADIPOGENESIS
UV_RESPONSE_UP
PI3K_AKT_MTOR_SIGNALING
OXIDATIVE_PHOSPHORYLATION
REACTIVE_OXYGEN_SPECIES_PATHWAY
DNA_REPAIR
APICAL_JUNCTION
ANDROGEN_RESPONSE
XENOBIOTIC_METABOLISM
PROTEIN_SECRETION
APICAL_SURFACE
MYC_TARGETS_V2
KRAS_SIGNALING_DN
TGF_BETA_SIGNALING
PANCREAS_BETA_CELLS
HEME_METABOLISM
WNT_BETA_CATENIN_SIGNALING
NOTCH_SIGNALING
UV_RESPONSE_DN
MYOGENESIS
HEDGEHOG_SIGNALING

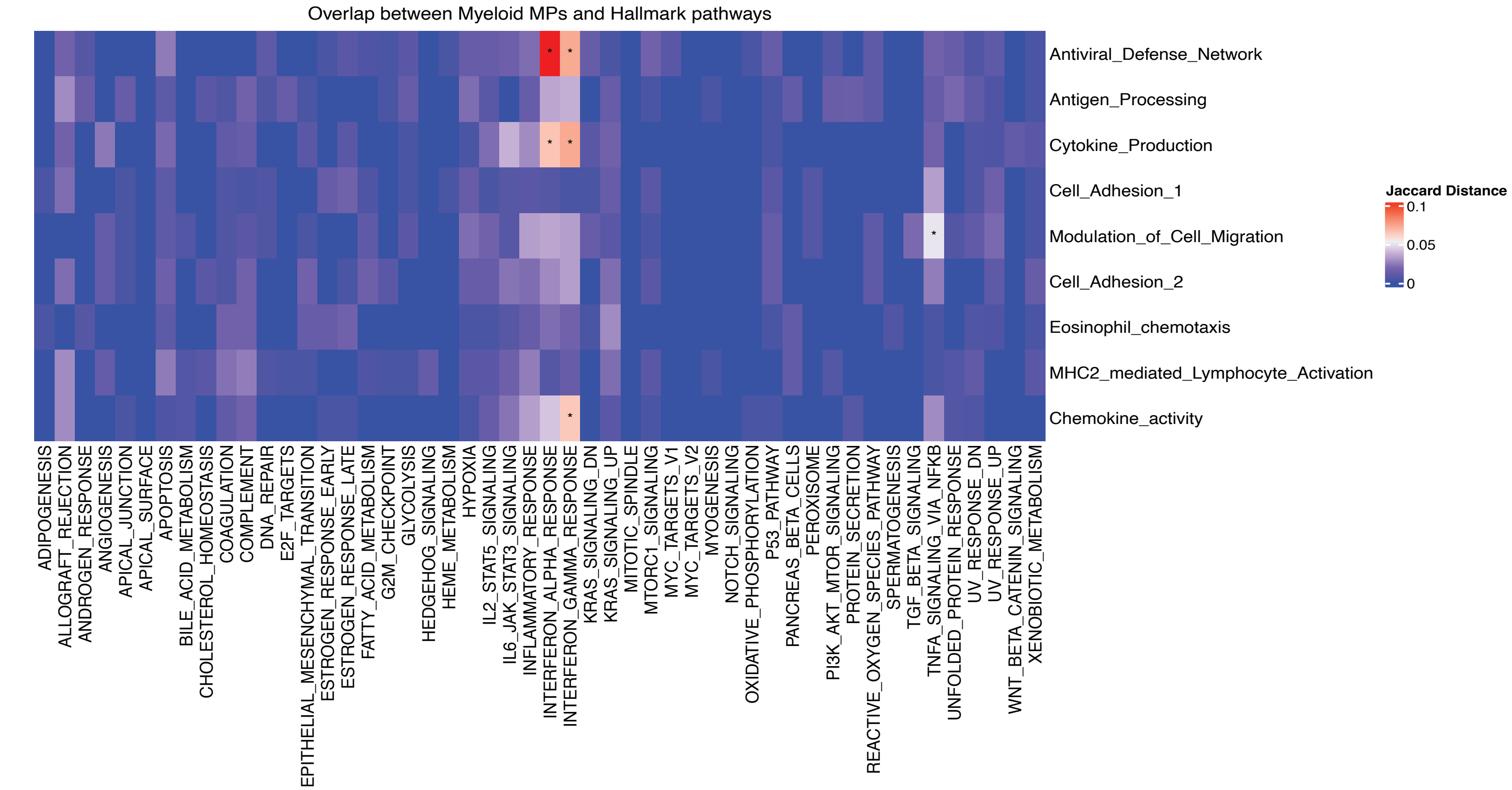
Overlap between Myeloid MPs and Hallmark pathways
Antiviral_Defense_Network
Antigen_Processing
Cytokine_Production
Cell_Adhesion_1
Modulation_of_Cell_Migration
Cell_Adhesion_2
Eosinophil_chemotaxis
MHC2_mediated_Lymphocyte_Activation
Chemokine_activity
Jaccard Distance
0.1
0.05
0
ADIPOGENESIS
ALLOGRAFT_REJECTION
ANDROGEN_RESPONSE
ANGIOGENESIS
APICAL_JUNCTION
APICAL_SURFACE
APOPTOSIS
BILE_ACID_METABOLISM
CHOLESTEROL_HOMEOSTASIS
COAGULATION
COMPLEMENT
DNA_REPAIR
E2F_TARGETS
EPITHELIAL_MESENCHYMAL_TRANSITION
ESTROGEN_RESPONSE_EARLY
ESTROGEN_RESPONSE_LATE
FATTY_ACID_METABOLISM
G2M_CHECKPOINT
GLYCOLYSIS
HEDGEHOG_SIGNALING
HEME_METABOLISM
HYPOXIA
IL2_STAT5_SIGNALING
IL6_JAK_STAT3_SIGNALING
INFLAMMATORY_RESPONSE
INTERFERON_ALPHA_RESPONSE
INTERFERON_GAMMA_RESPONSE
KRAS_SIGNALING_DN
KRAS_SIGNALING_UP
MITOTIC_SPINDLE
MTORC1_SIGNALING
MYC_TARGETS_V1
MYC_TARGETS_V2
MYOGENESIS
NOTCH_SIGNALING
OXIDATIVE_PHOSPHORYLATION
P53_PATHWAY
PANCREAS_BETA_CELLS
PEROXISOME
PI3K_AKT_MTOR_SIGNALING
PROTEIN_SECRETION
REACTIVE_OXYGEN_SPECIES_PATHWAY
SPERMATOGENESIS
TGF_BETA_SIGNALING
TNFA_SIGNALING_VIA_NFKB
UNFOLDED_PROTEIN_RESPONSE
UV_RESPONSE_DN
UV_RESPONSE_UP
WNT_BETA_CATENIN_SIGNALING
XENOBIOTIC_METABOLISM

**Figure S5. [Gene sets comparison with Hallmark], Related to Figure 1.**

Pairwise jaccard distance between Lymphoid irMPs (top)/Myeloid irMPs (bottom) and 50 Hallmark pathways. Pairwise comparisons with adjusted p-value<0.05 are overlaid with asterisks.

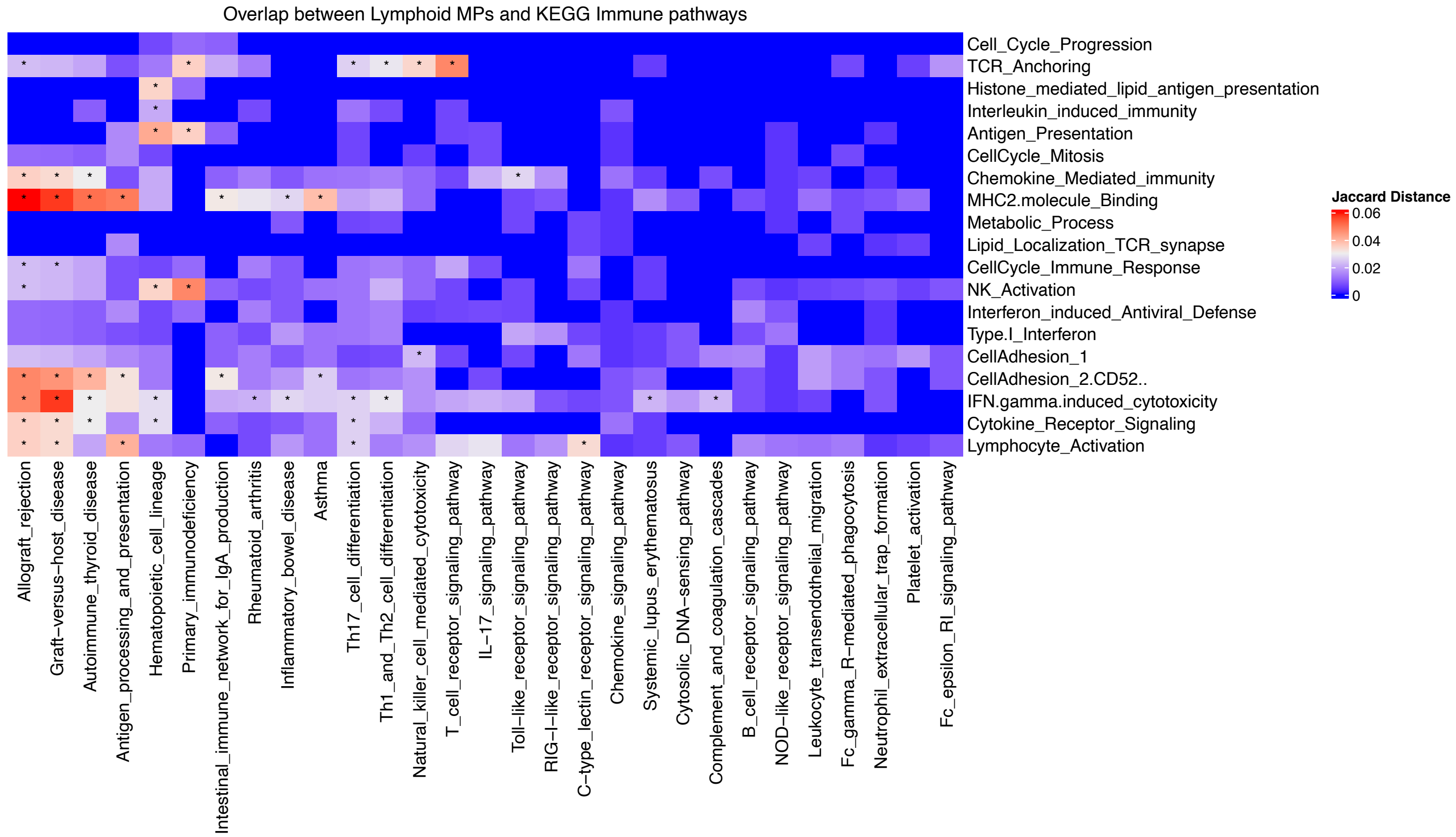

Overlap between Lymphoid MPs and KEGG Immune pathways
Jaccard Distance
0.06
0.04
0.02
0
Cell_Cycle_Progression
TCR_Anchoring
Histone_mediated_lipid_antigen_presentation
Interleukin_induced_immunity
Antigen_Presentation
CellCycle_Mitosis
Chemokine_Mediated_immunity
MHC2.molecule_Binding
Metabolic_Process
Lipid_Localization_TCR_synapse
CellCycle_Immune_Response
NK_Activation
Interferon_induced_Antiviral_Defense
Type.I_Interferon
CellAdhesion_1
CellAdhesion_2.CD52..
IFN.gamma.induced_cytotoxicity
Cytokine_Receptor_Signaling
Lymphocyte_Activation
Allograft_rejection
Graft−versus−host_disease
Autoimmune_thyroid_disease
Antigen_processing_and_presentation
Hematopoietic_cell_lineage
Primary_immunodeficiency
Intestinal_immune_network_for_IgA_production
Rheumatoid_arthritis
Inflammatory_bowel_disease
Asthma
Th17_cell_differentiation
Th1_and_Th2_cell_differentiation
Natural_killer_cell_mediated_cytotoxicity
T_cell_receptor_signaling_pathway
IL−17_signaling_pathway
Toll−like_receptor_signaling_pathway
RIG−I−like_receptor_signaling_pathway
C−type_lectin_receptor_signaling_pathway
Chemokine_signaling_pathway
Systemic_lupus_erythematosus
Cytosolic_DNA−sensing_pathway
Complement_and_coagulation_cascades
B_cell_receptor_signaling_pathway
NOD−like_receptor_signaling_pathway
Leukocyte_transendothelial_migration
Fc_gamma_R−mediated_phagocytosis
Neutrophil_extracellular_trap_formation
Platelet_activation
Fc_epsilon_RI_signaling_pathway


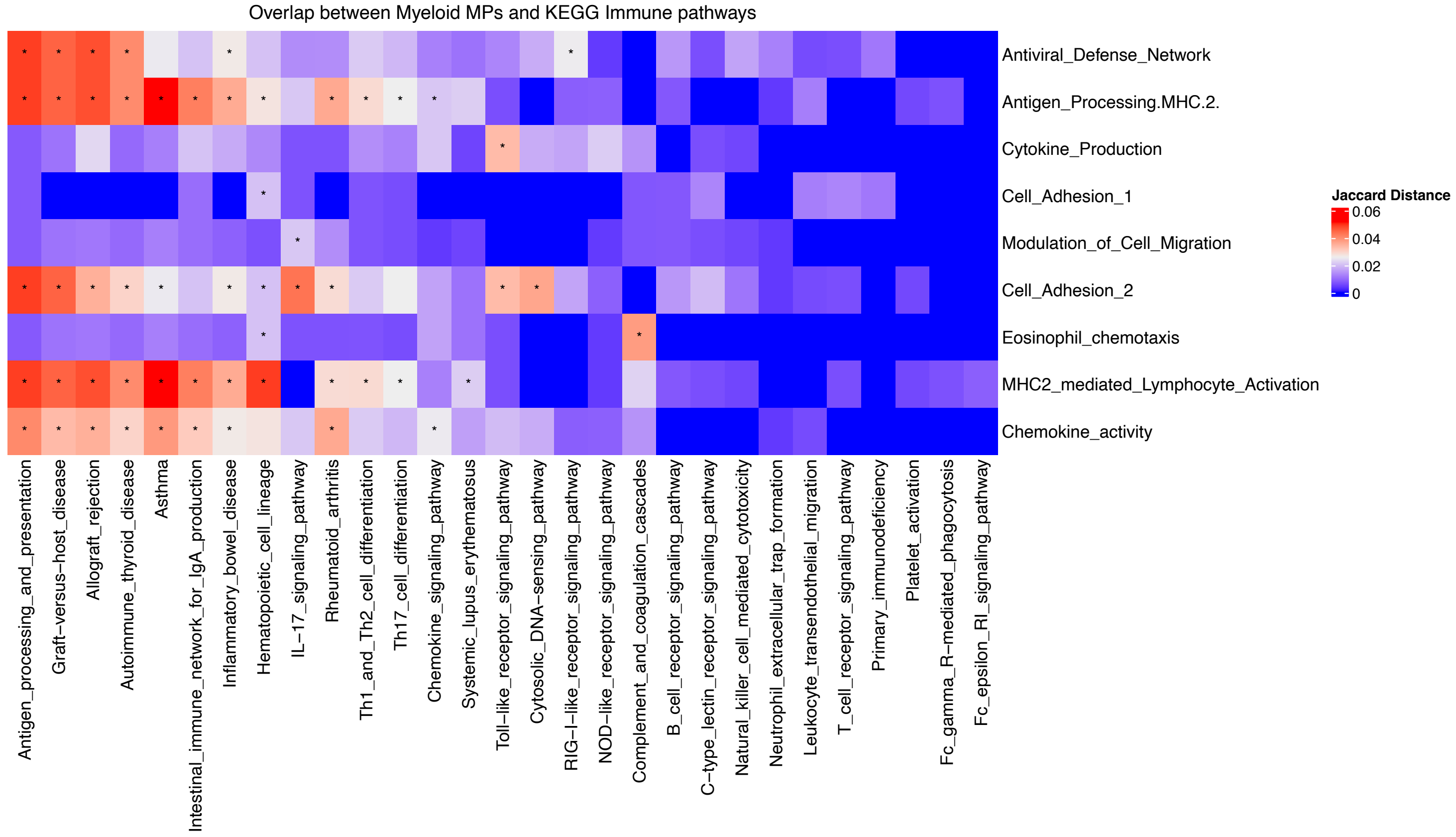

Overlap between Myeloid MPs and KEGG Immune pathways
Jaccard Distance
0.06
0.04
0.02
0
Antiviral_Defense_Network
Antigen_Processing.MHC.2.
Cytokine_Production
Cell_Adhesion_1
Modulation_of_Cell_Migration
Cell_Adhesion_2
Eosinophil_chemotaxis
MHC2_mediated_Lymphocyte_Activation
Chemokine_activity
Antigen_processing_and_presentation
Graft−versus−host_disease
Allograft_rejection
Autoimmune_thyroid_disease
Asthma
Intestinal_immune_network_for_IgA_production
Inflammatory_bowel_disease
Hematopoietic_cell_lineage
IL−17_signaling_pathway
Rheumatoid_arthritis
Th1_and_Th2_cell_differentiation
Th17_cell_differentiation
Chemokine_signaling_pathway
Systemic_lupus_erythematosus
Toll−like_receptor_signaling_pathway
Cytosolic_DNA−sensing_pathway
RIG−I−like_receptor_signaling_pathway
NOD−like_receptor_signaling_pathway
Complement_and_coagulation_cascades
B_cell_receptor_signaling_pathway
C−type_lectin_receptor_signaling_pathway
Natural_killer_cell_mediated_cytotoxicity
Neutrophil_extracellular_trap_formation
Leukocyte_transendothelial_migration
T_cell_receptor_signaling_pathway
Primary_immunodeficiency
Platelet_activation
Fc_gamma_R−mediated_phagocytosis
Fc_epsilon_RI_signaling_pathway

**Figure S6. [Gene sets comparison with KEGG], Related to Figure 1.**

Pairwise jaccard distance between Lymphoid irMPs (top)/Myeloid irMPs (bottom) and 29 irKEGG pathways. Pairwise comparisons with adjusted p-value<0.05 are overlaid with asterisks.

S7.1

### Cell Cycle 1

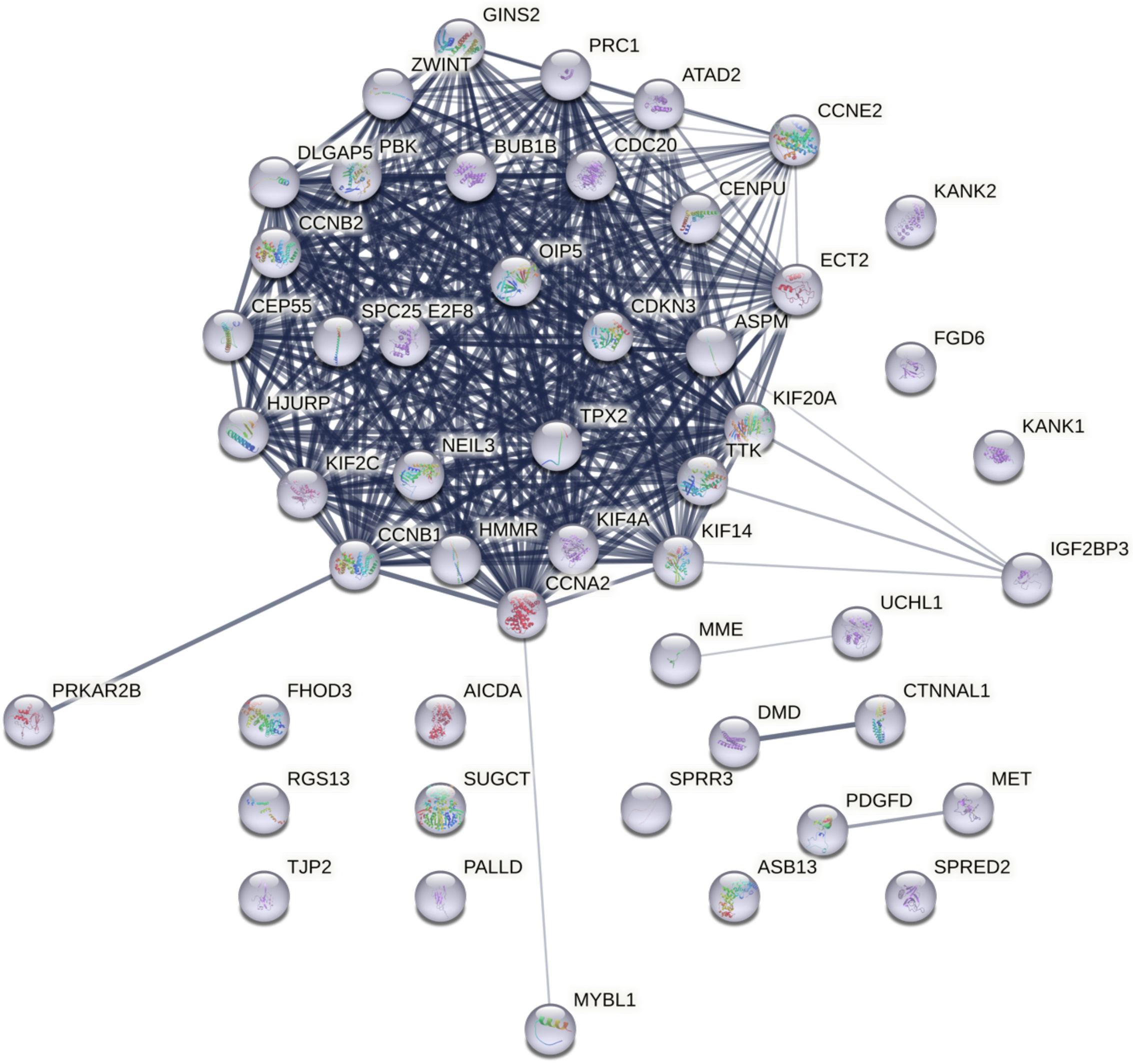

S7.2

### TCR Anchoring

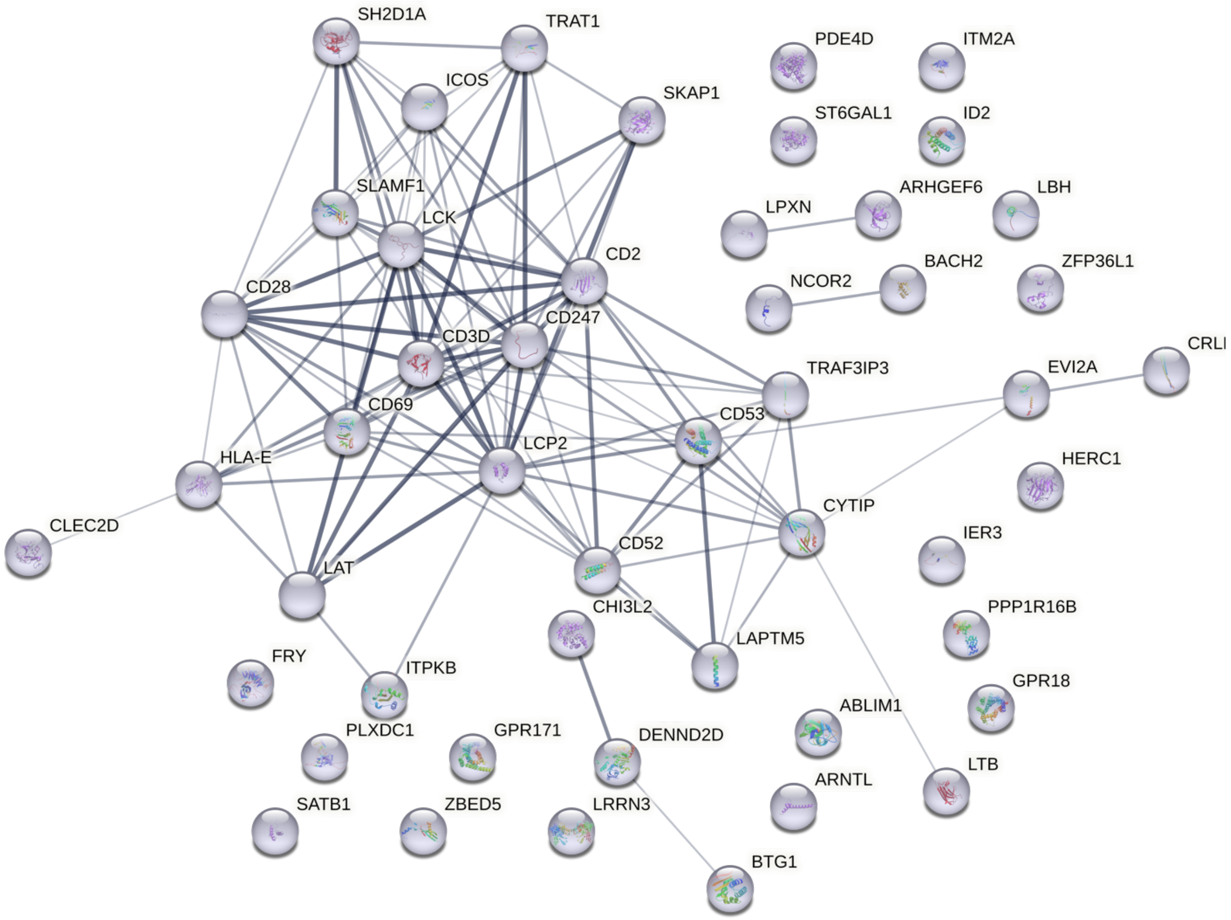

S7.3

### Histone associated lipid antigen presentation

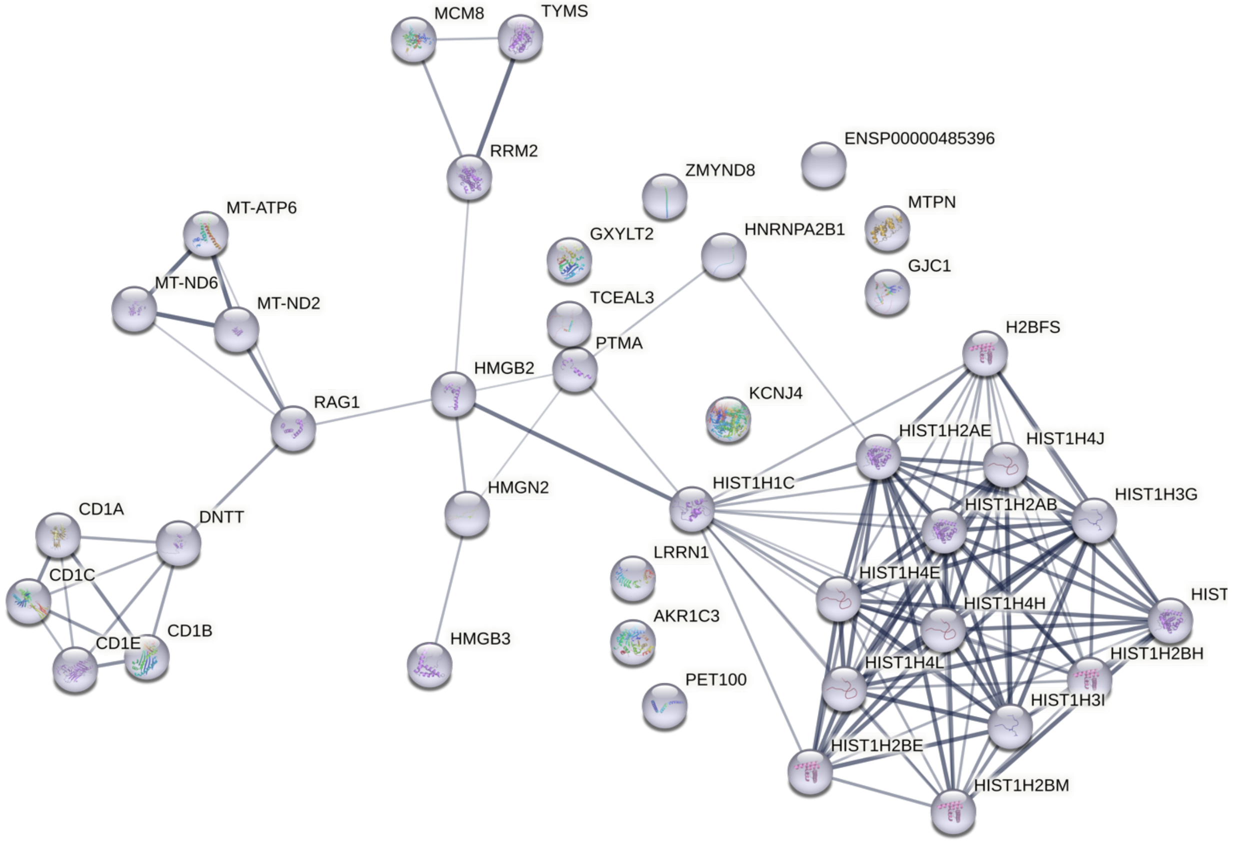

S7.4

### Interleukin induced Treg Activation

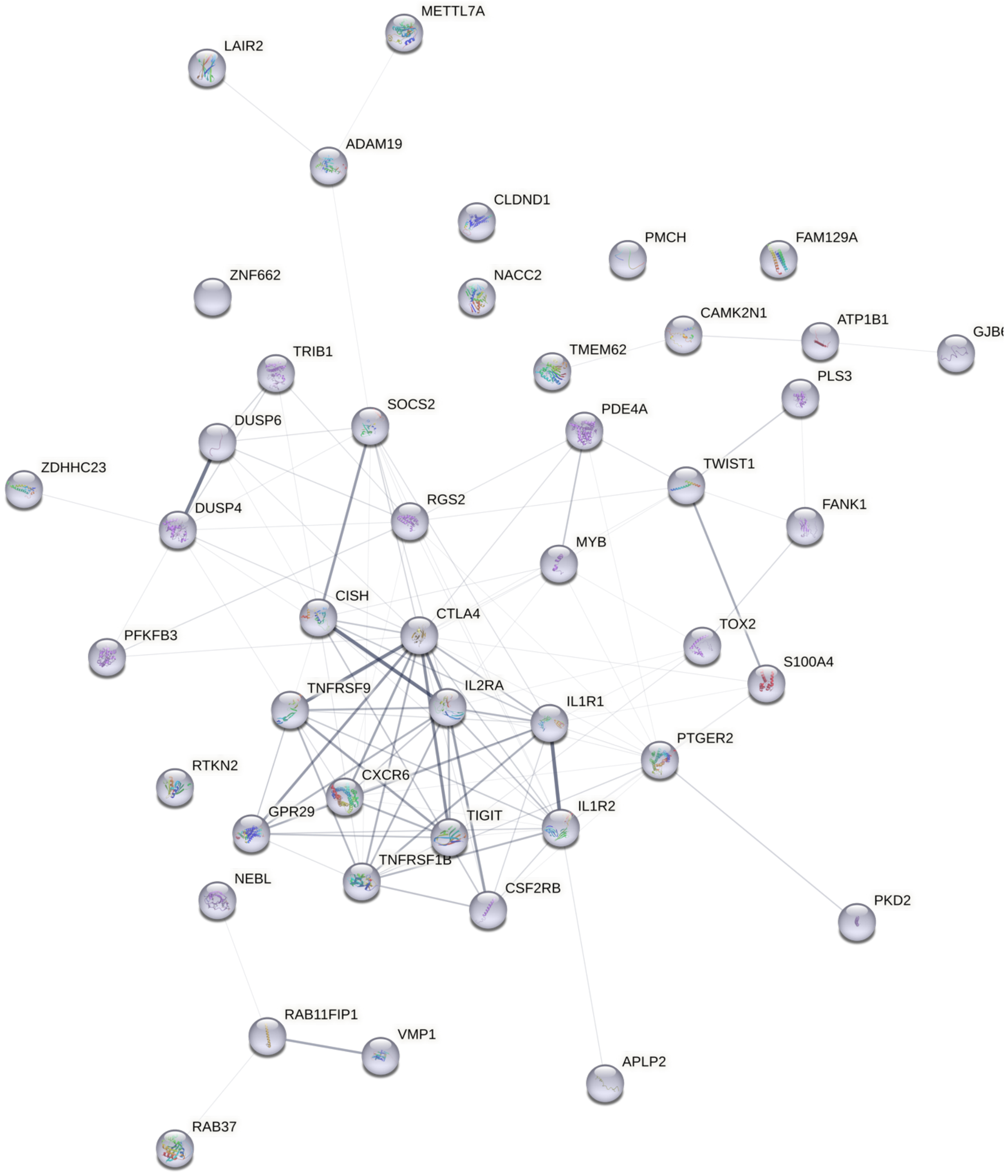

S7.5

### Antigen Processing

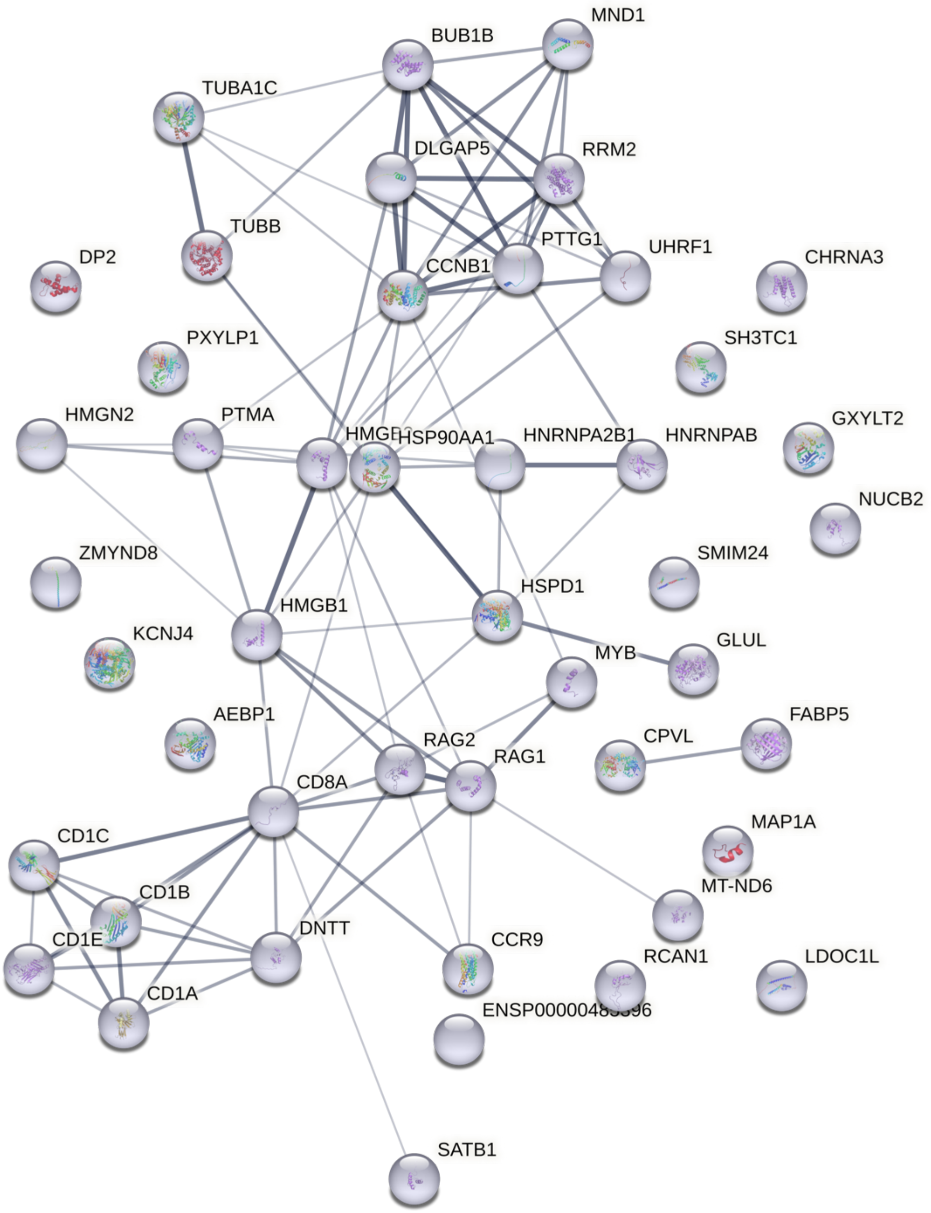

S7.6

### Cell Cycle 2

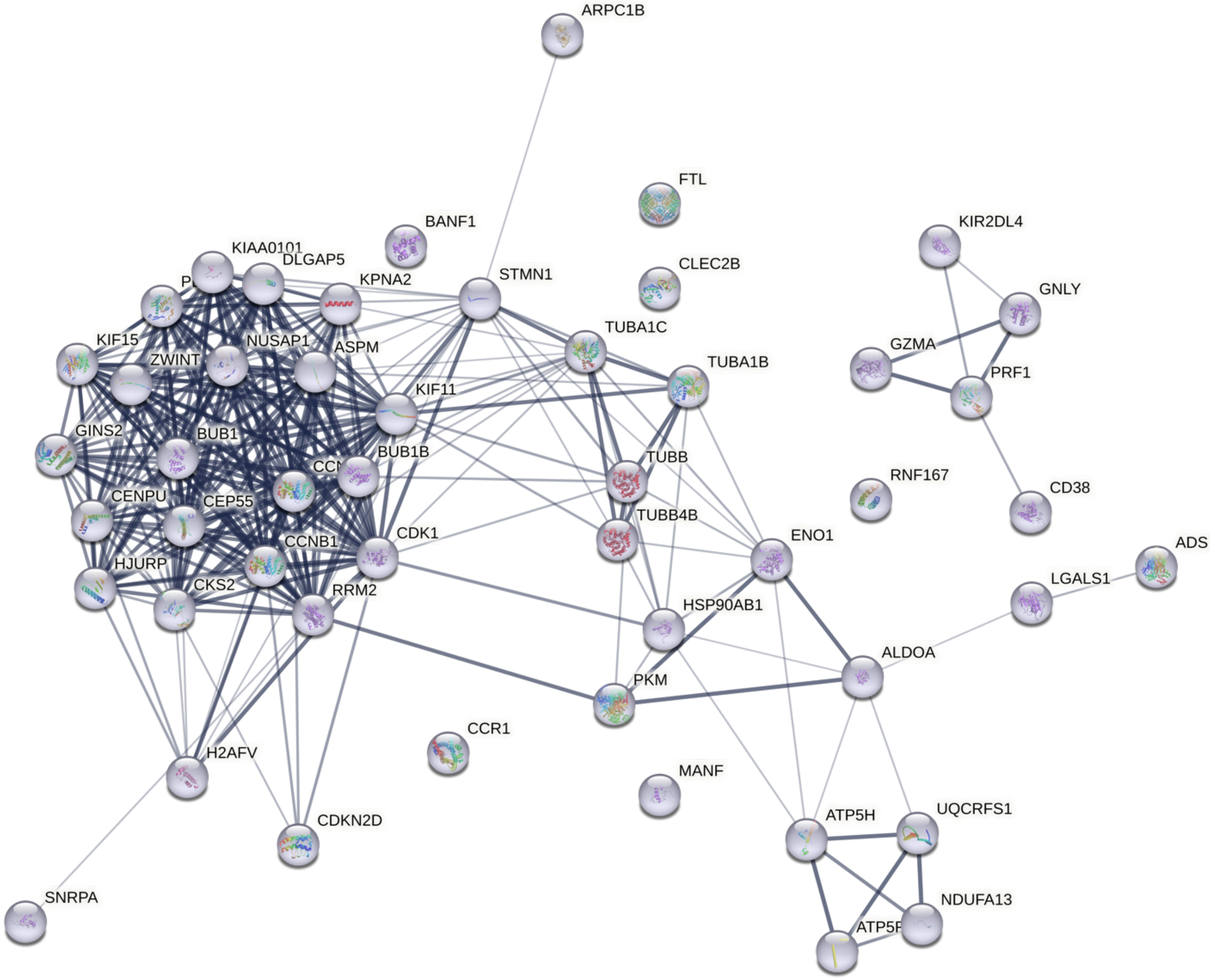

S7.7

### Chemokine Mediated T cell Activation

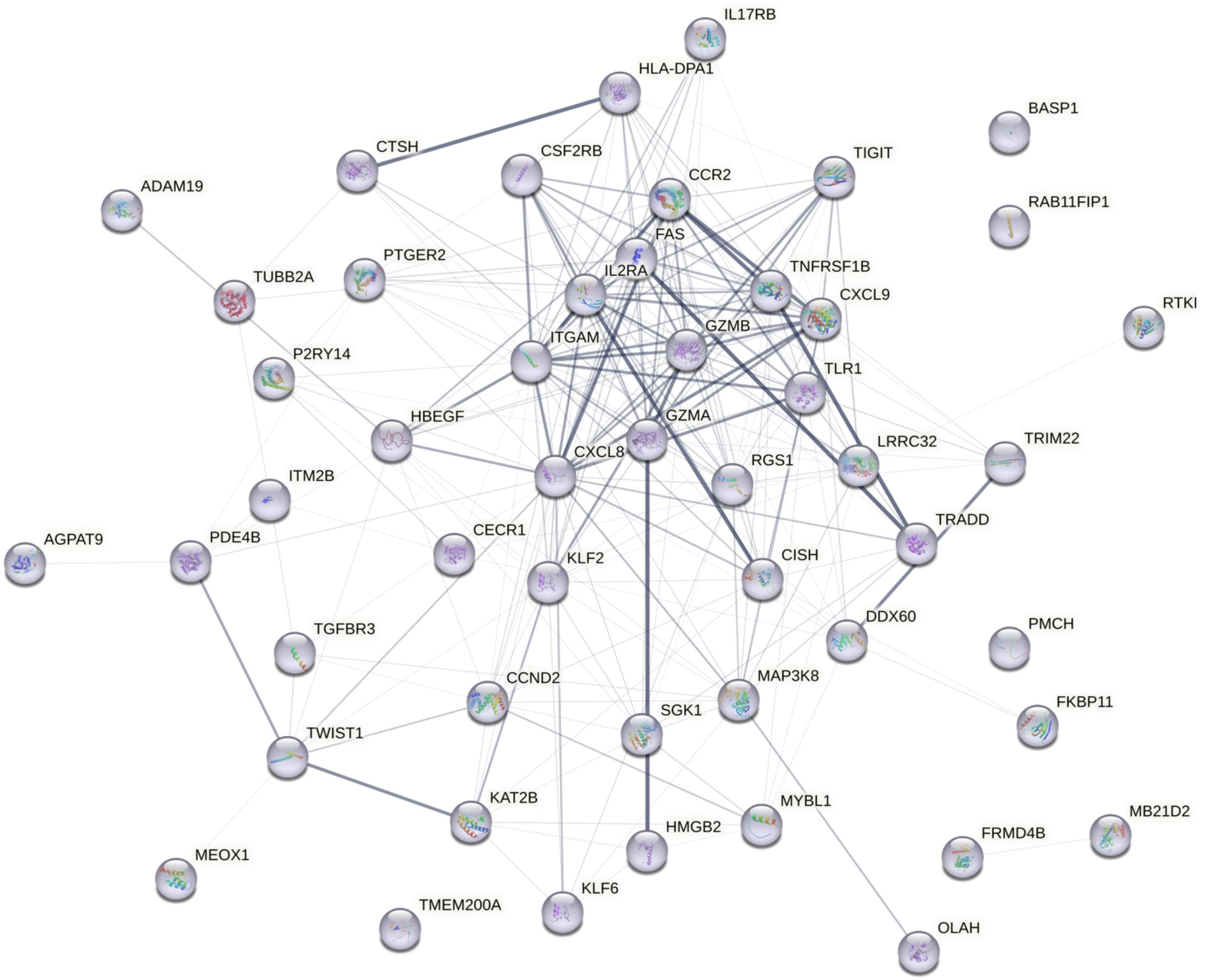

S7.8

### MHC-II mediated Immunity

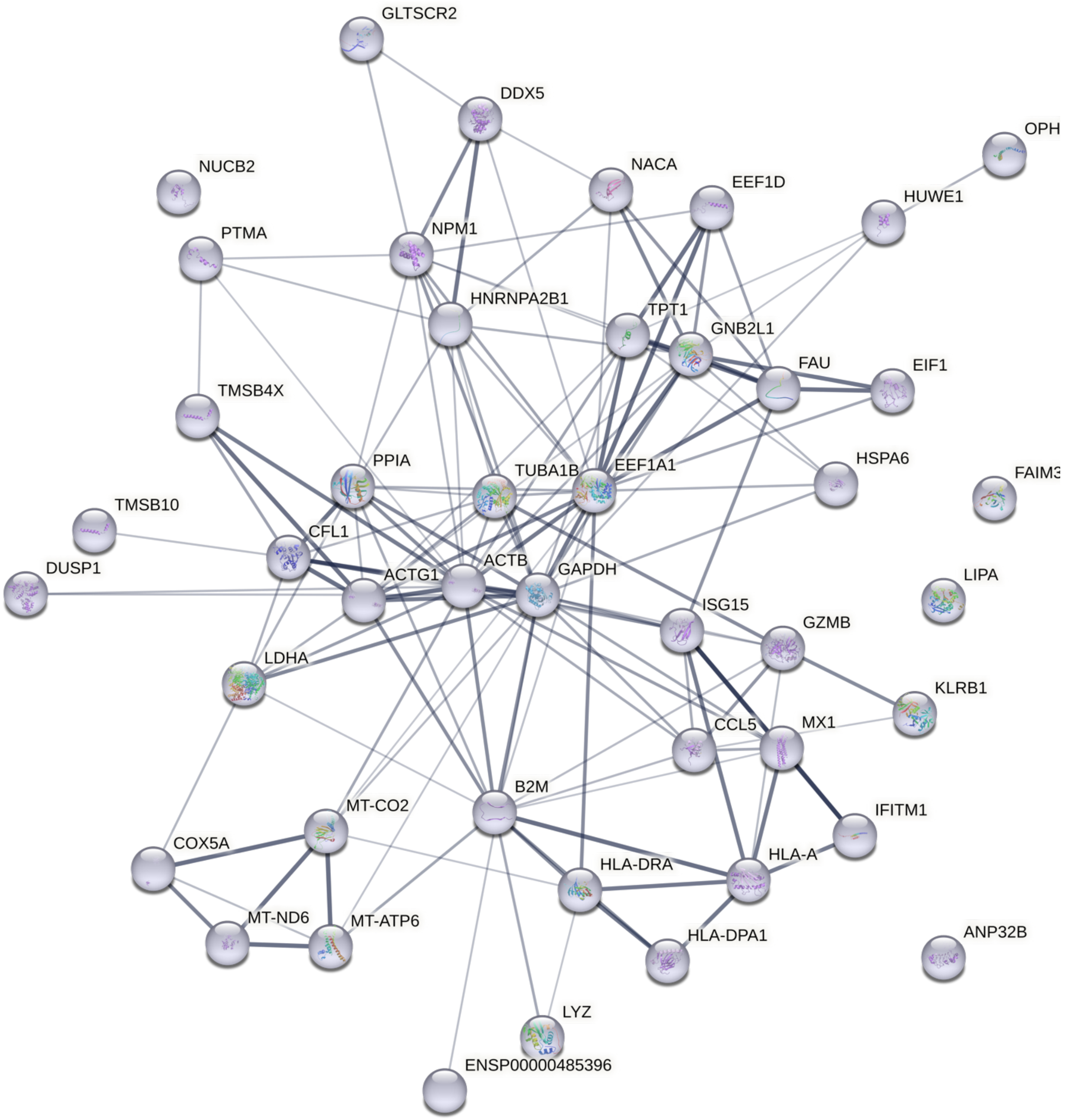

S7.9

#### Metabolic Process

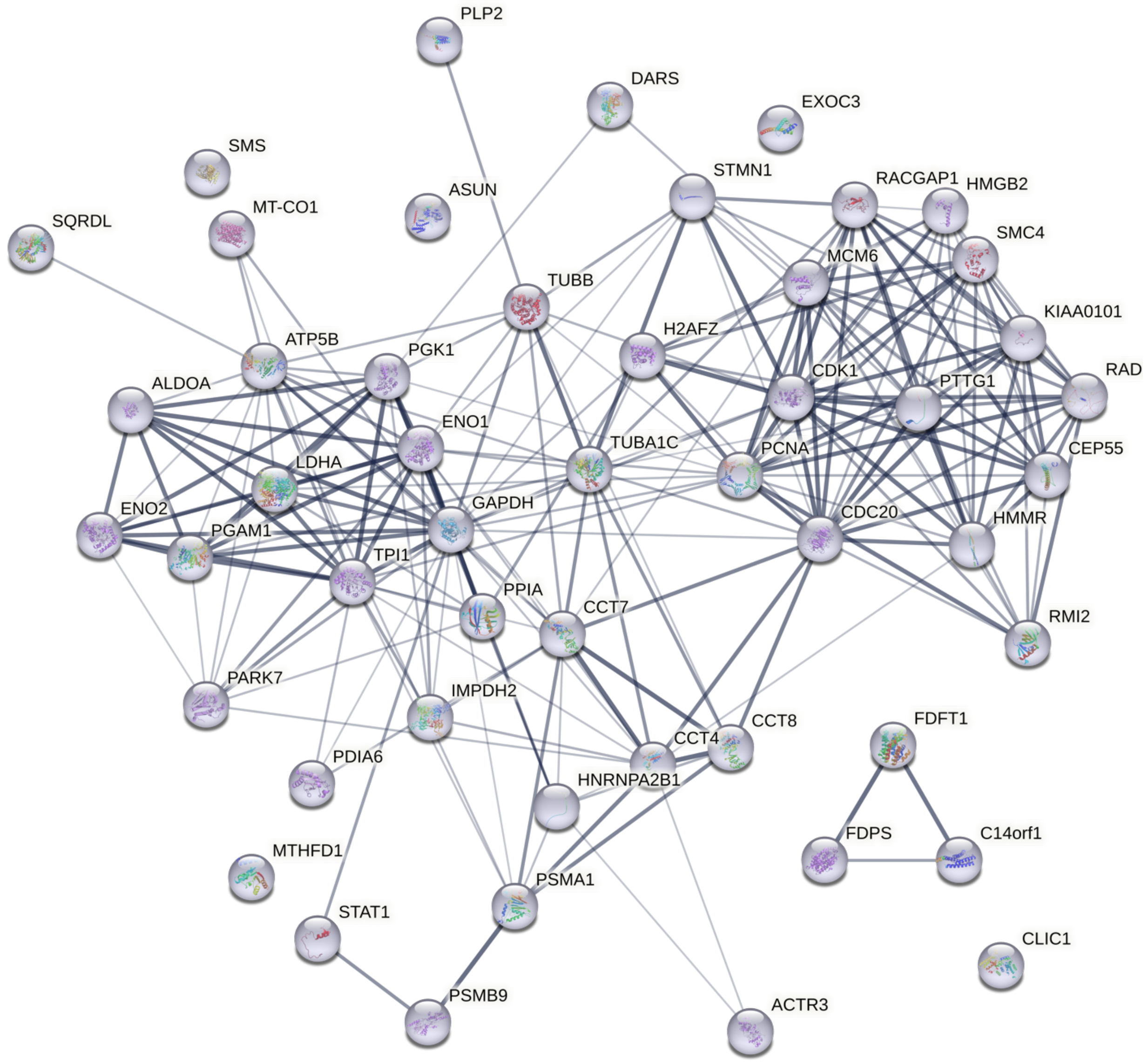

S7.10

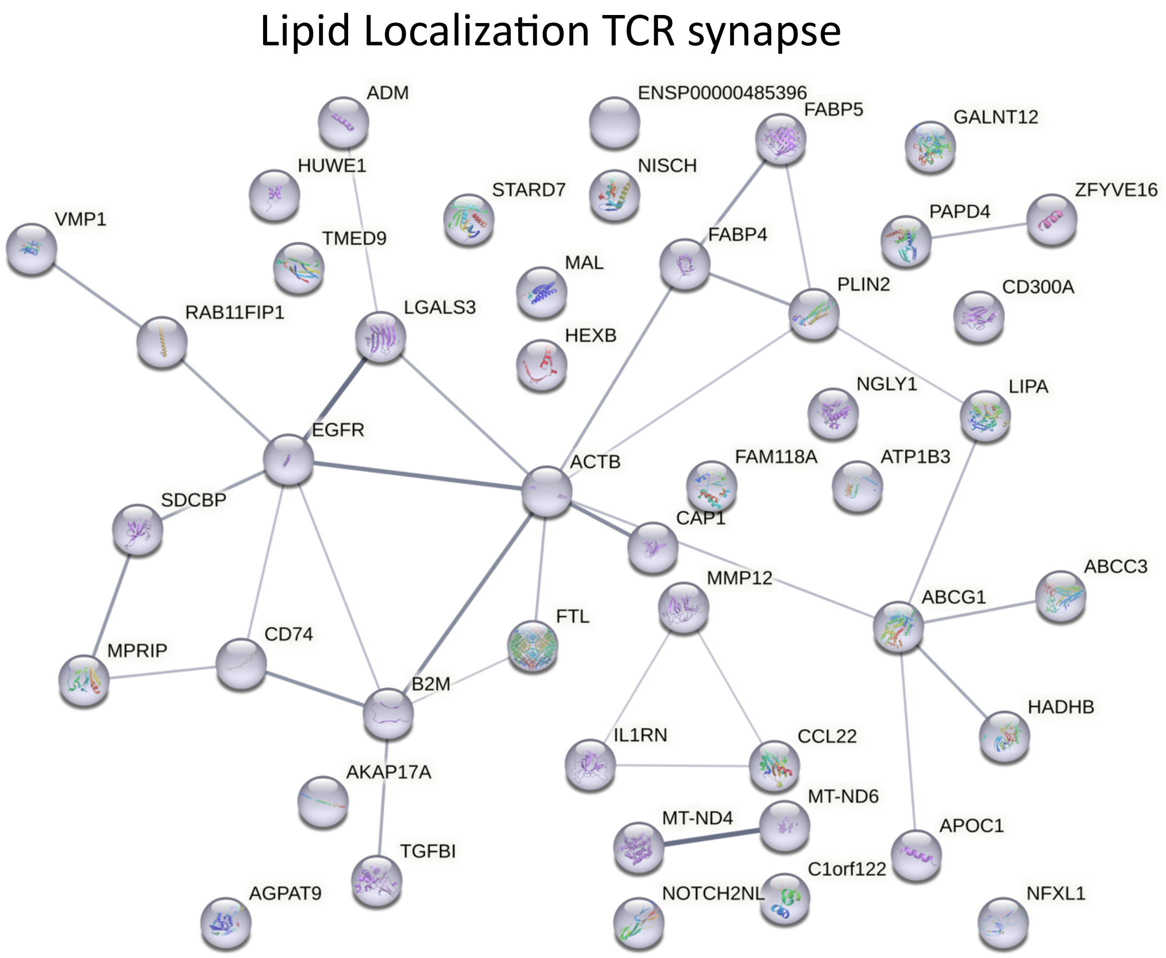
Lipid Localization TCR synapse
ADM
ENSP00000485396
FABP5
GALNT12
HUWE1
NISCH
STARD7
ZFYVE16
VMP1
PAPD4
TMED9
FABP4
MAL
PLIN2
CD300A
RAB11FIP1
LGALS3
HEXB
NGLY1
LIPA
EGFR
FAM118A
ATP1B3
ACTB
SDCBP
CAP1
ABCC3
MMP12
ABCG1
FTL
CD74
MPRIP
B2M
HADHB
IL1RN
CCL22
AKAP17A
MT-ND6
MT-ND4
APOC1
TGFBI
C1orf122
AGPAT9
NOTCH2NL
NFXL1

S7.11

### Cell Cycle Immune Response

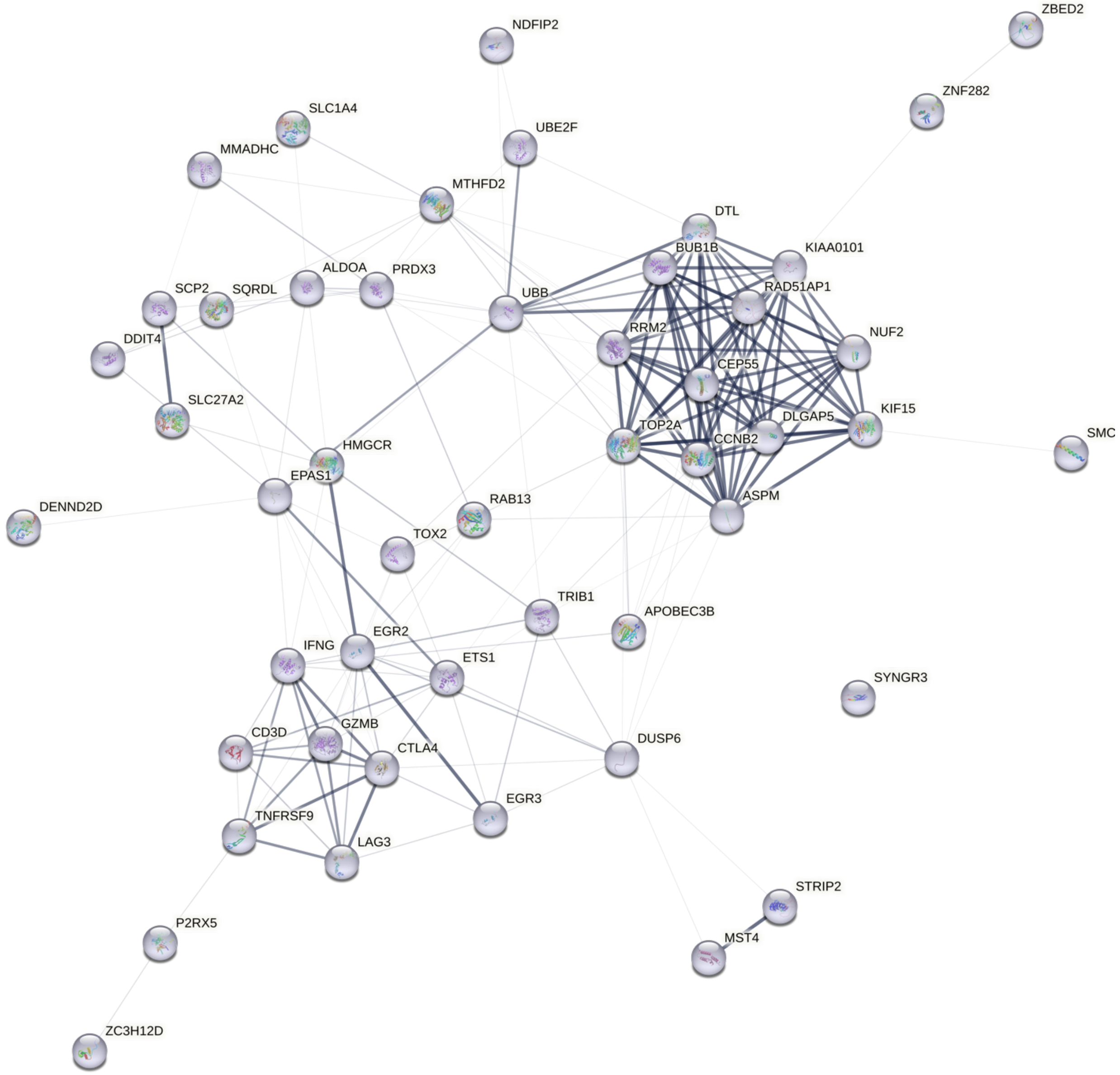

S7.12

### Lymphocyte Activation

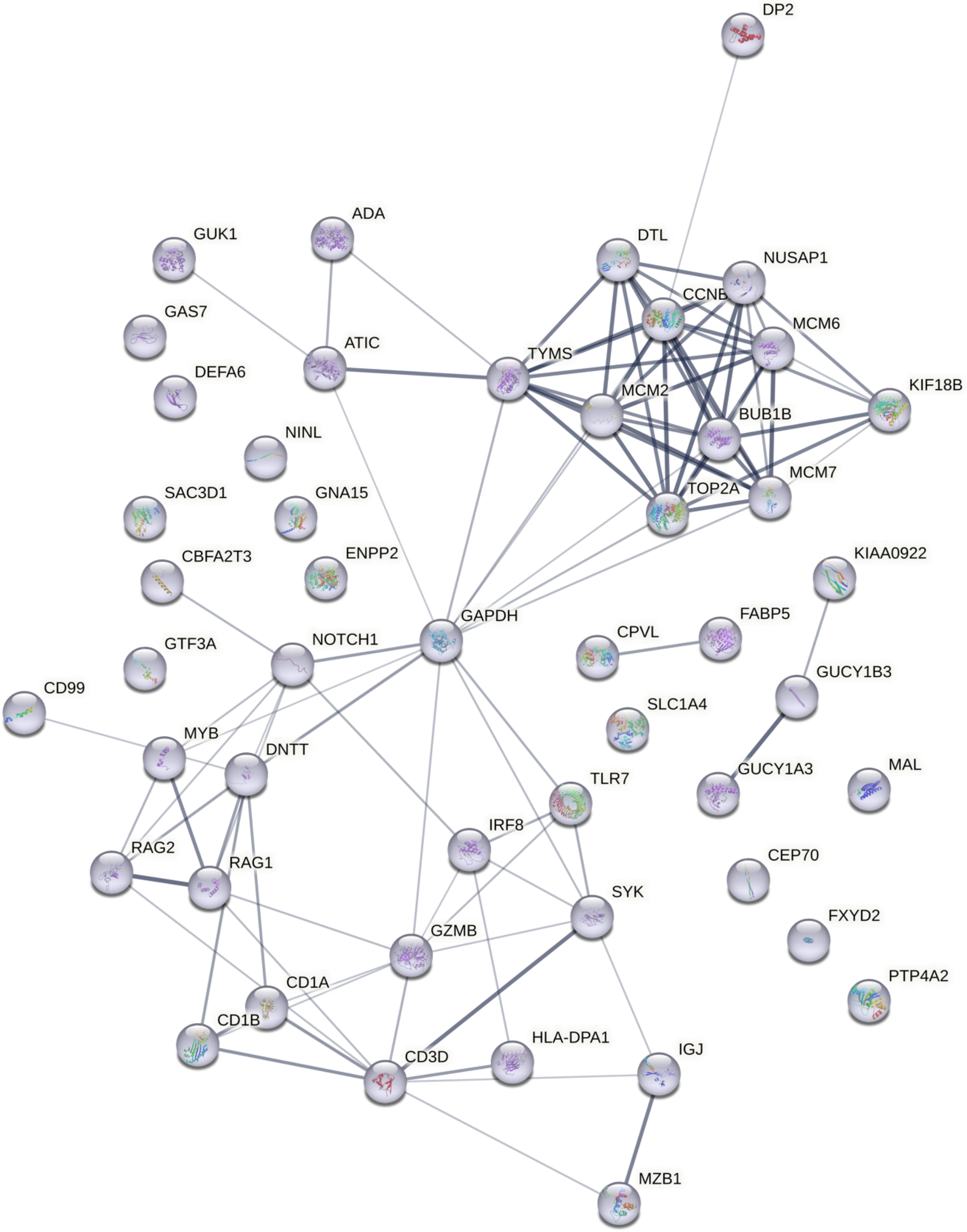

S7.13

### Interferon induced Antiviral Defense

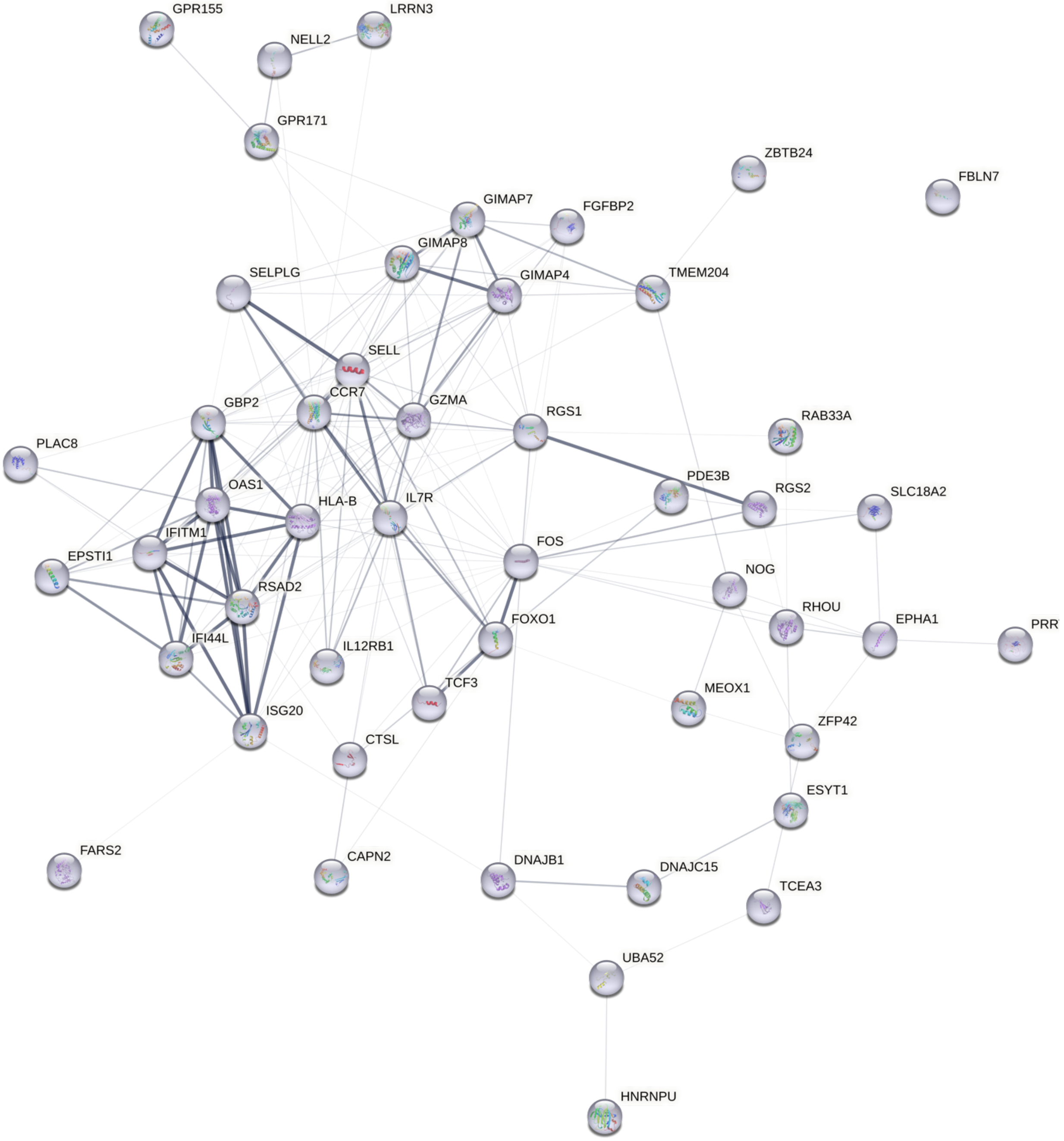

S7.14

### Type-I Interferon

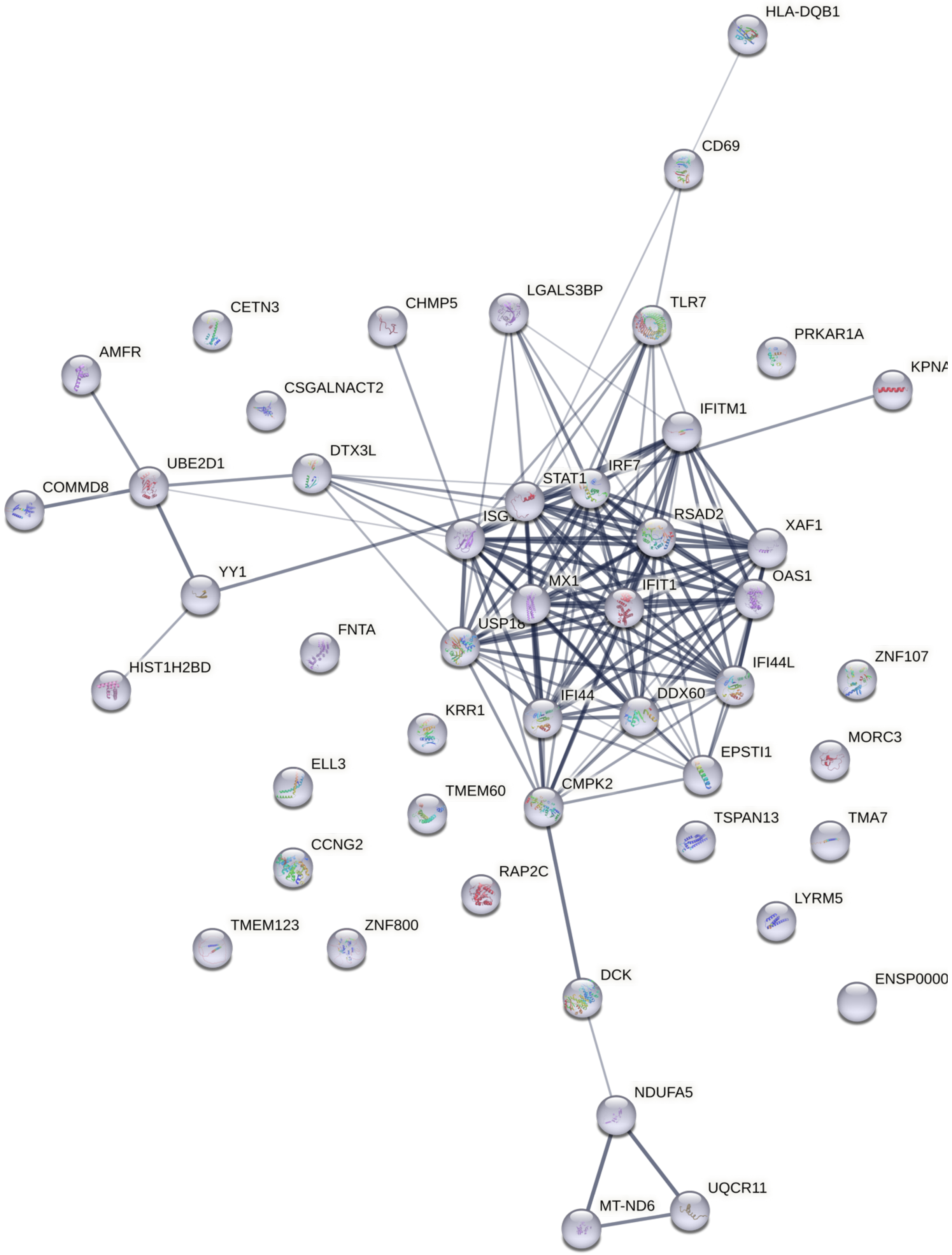

S7.15

### Cell Adhesion_1

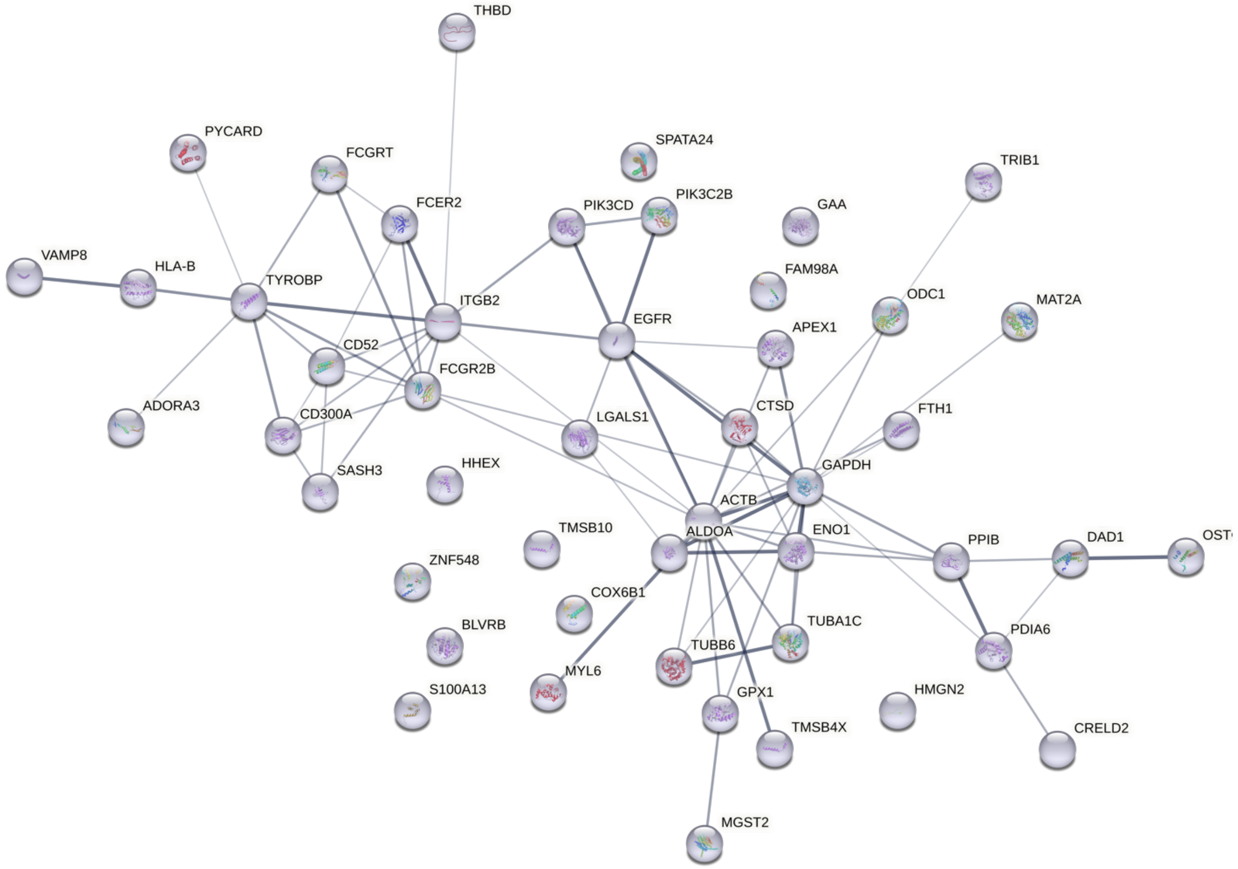

S7.16

### Cell Adhesion_2 (CD52)

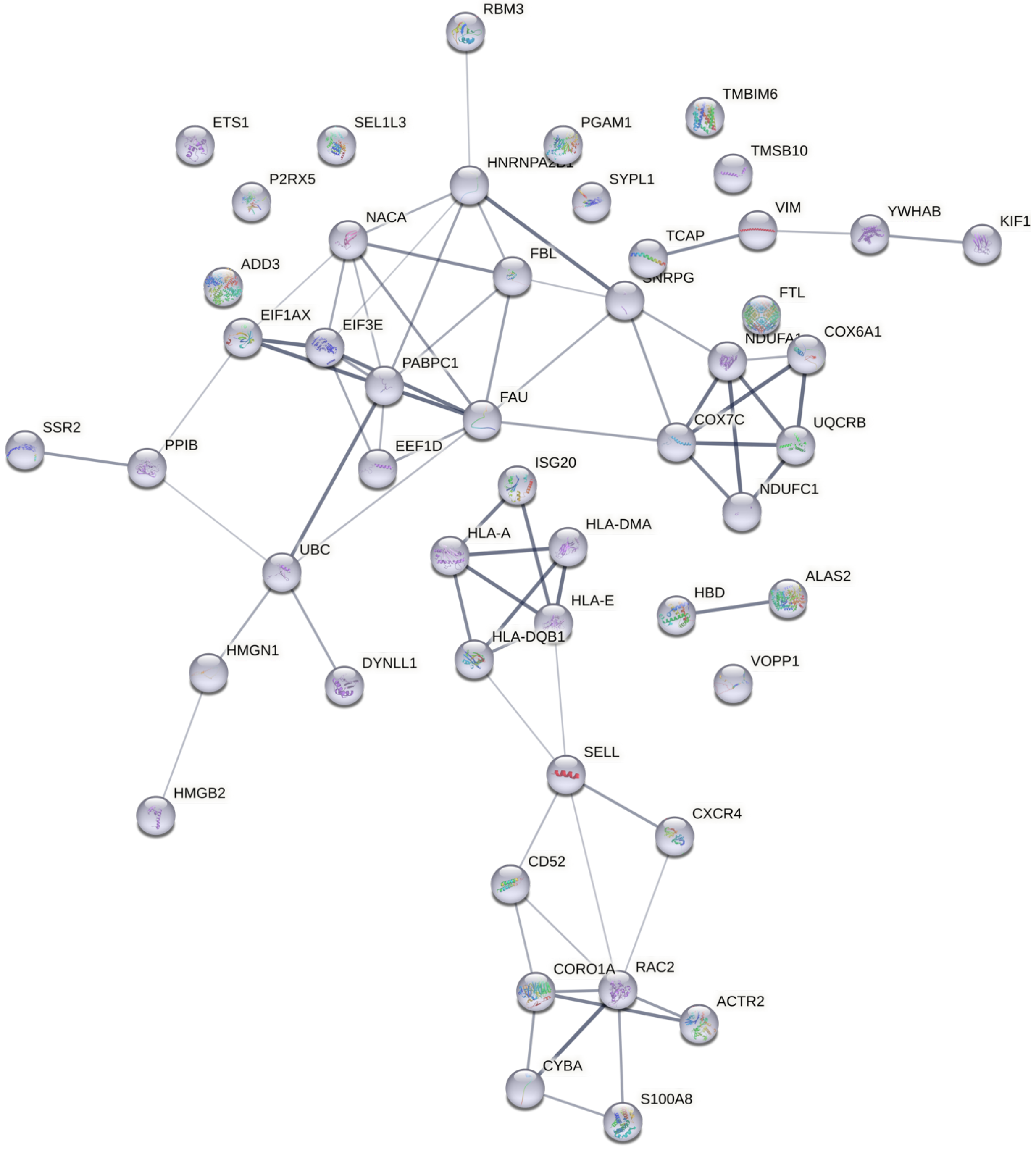

S7.17

### IFN-gamma induced cytotoxicity

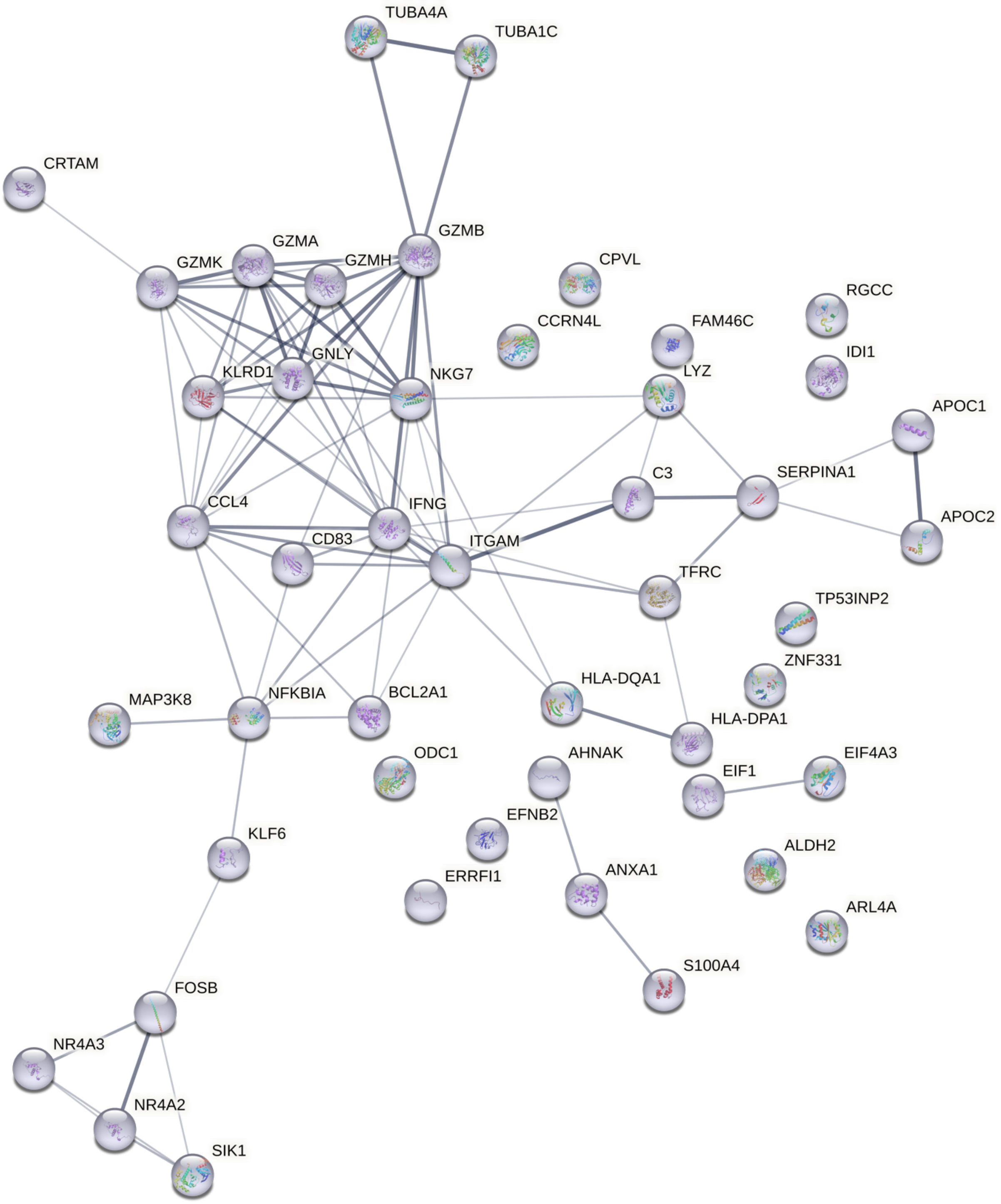

S7.18

### Cytokine Receptor Signaling

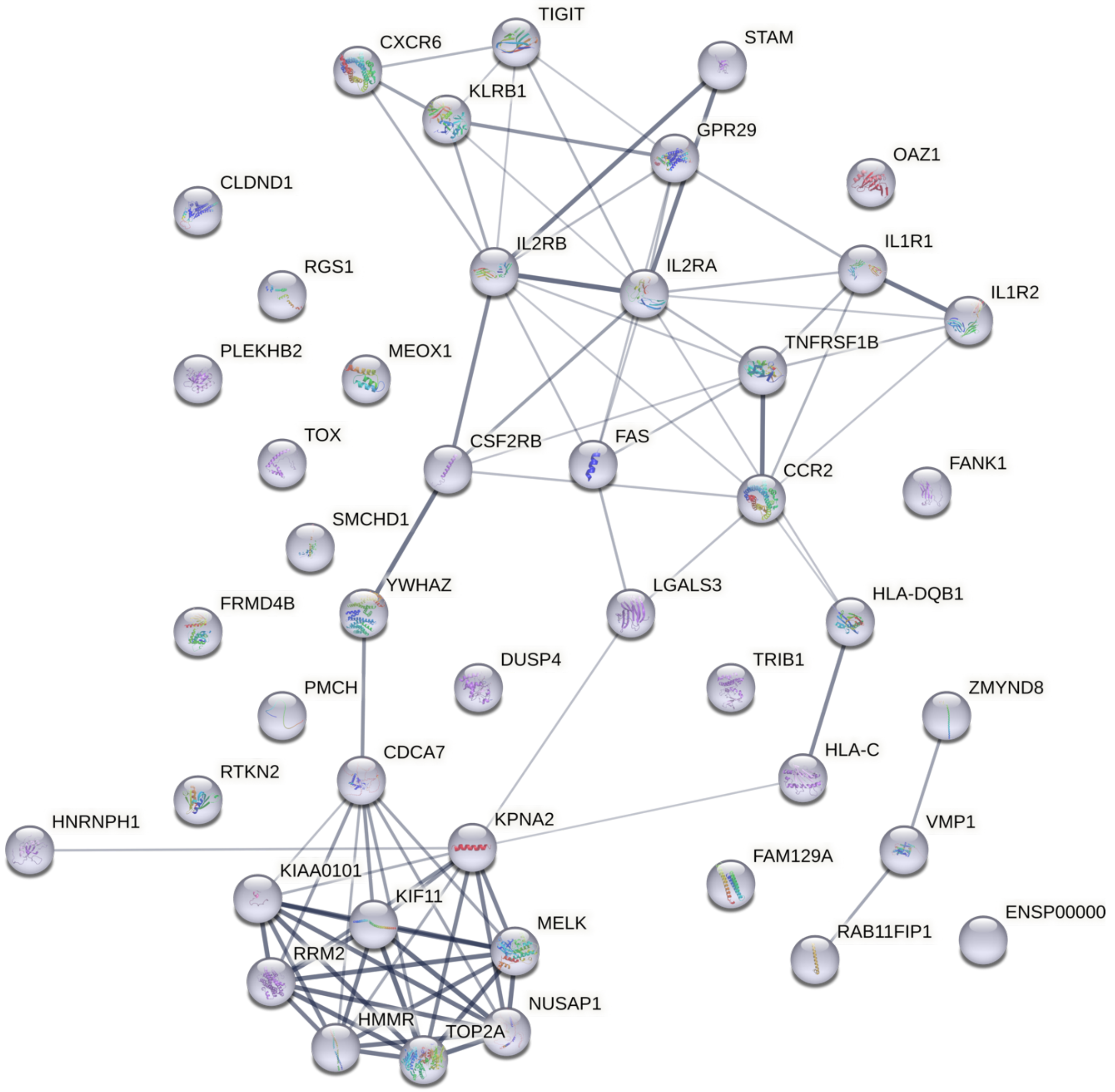

S7.19

### T cell Activation

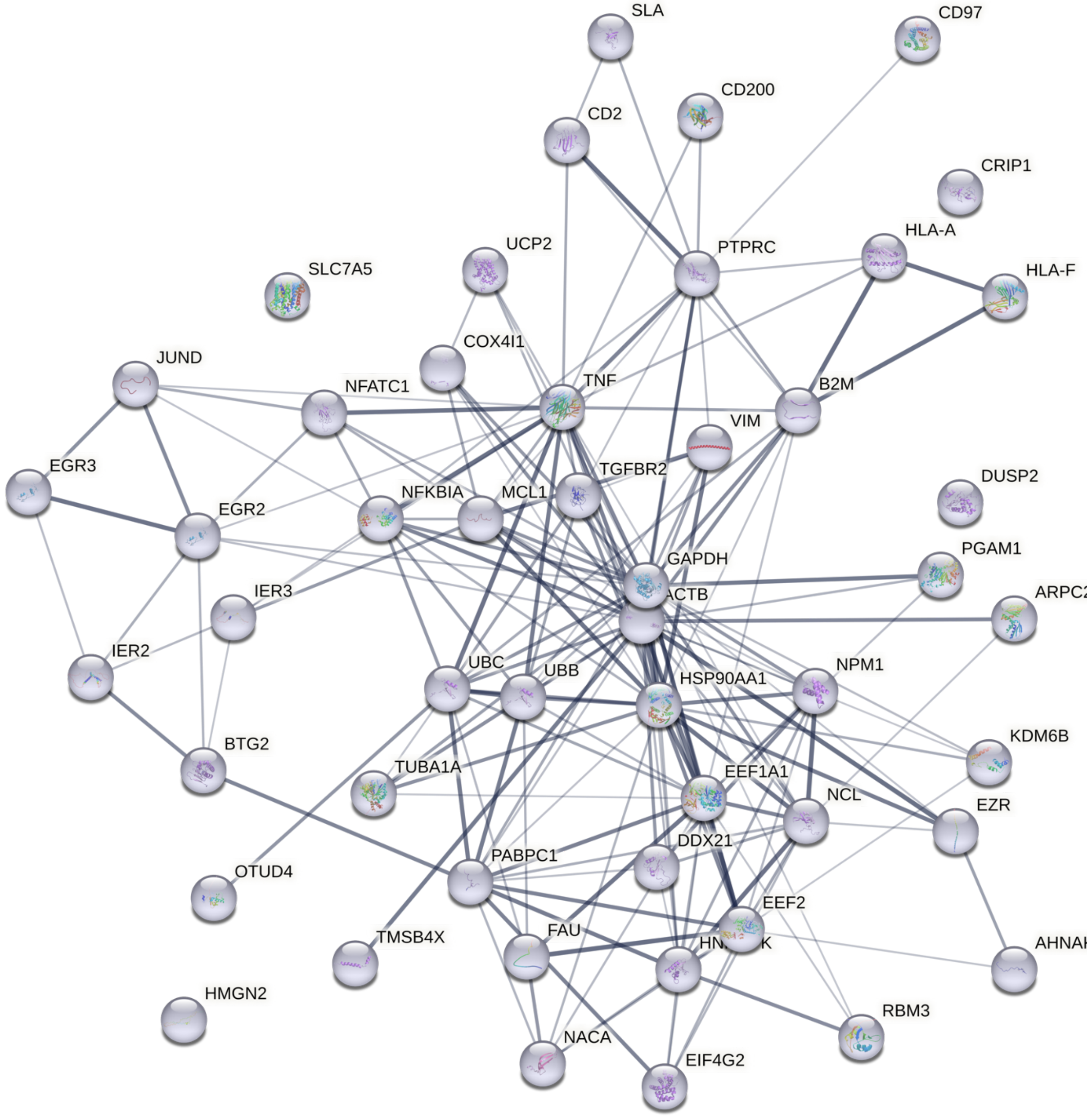

**Figure S7. [Protein-protein interaction network], Related to Figure 1.**

Protein-protein interaction network for all 19 lymphoid-derived irMPs. Each node represents a protein produced by a single, protein-coding gene locus. The protein 3D structure is known if the node is filled with a miniature 3D structure. Edge thickness refers to the level of confidence, with thicker edges indicating more confident protein interaction, and such interactions could come from any of the following: text mining, gene co-expression, neighborhood, gene fusion or co-occurrence.

S8.1

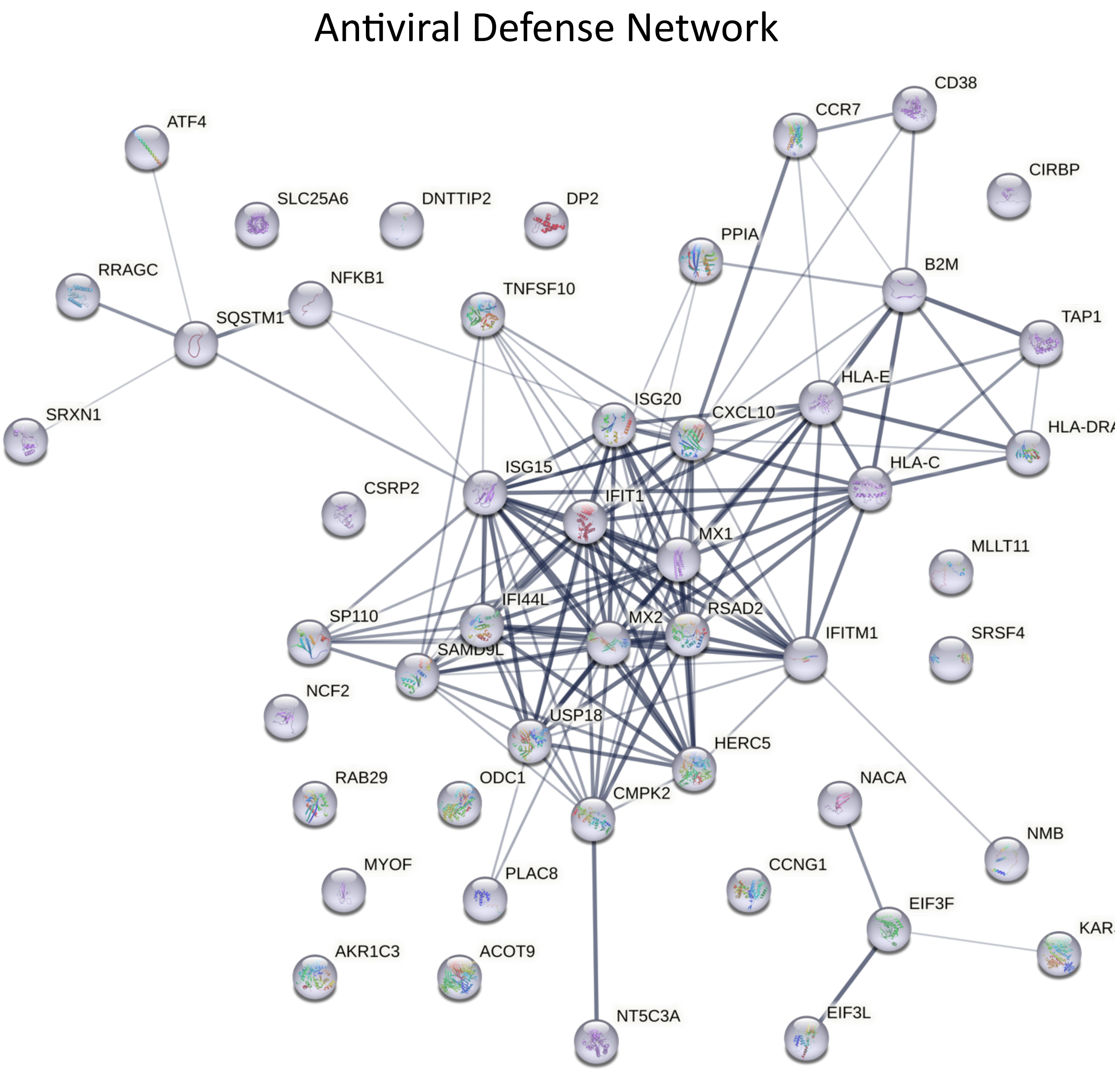
Antiviral Defense Network
ATF4
SLC25A6
DNTTIP2
DP2
CCR7
CD38
CIRBP
PPIA
B2M
RRAGC
NFKB1
TNFSF10
SQSTM1
TAP1
SRXN1
ISG20
CXCL10
HLA-E
HLA-DRA
HLA-C
CSRP2
ISG15
IFIT1
MX1
MLLT11
IFI44L
SP110
MX2
RSAD2
SAMD9L
IFITM1
SRSF4
NCF2
USP18
HERC5
RAB29
ODC1
CMPK2
NACA
NMB
MYOF
PLAC8
CCNG1
EIF3F
AKR1C3
ACOT9
NT5C3A
EIF3L

S8.2

#### Antigen Processing

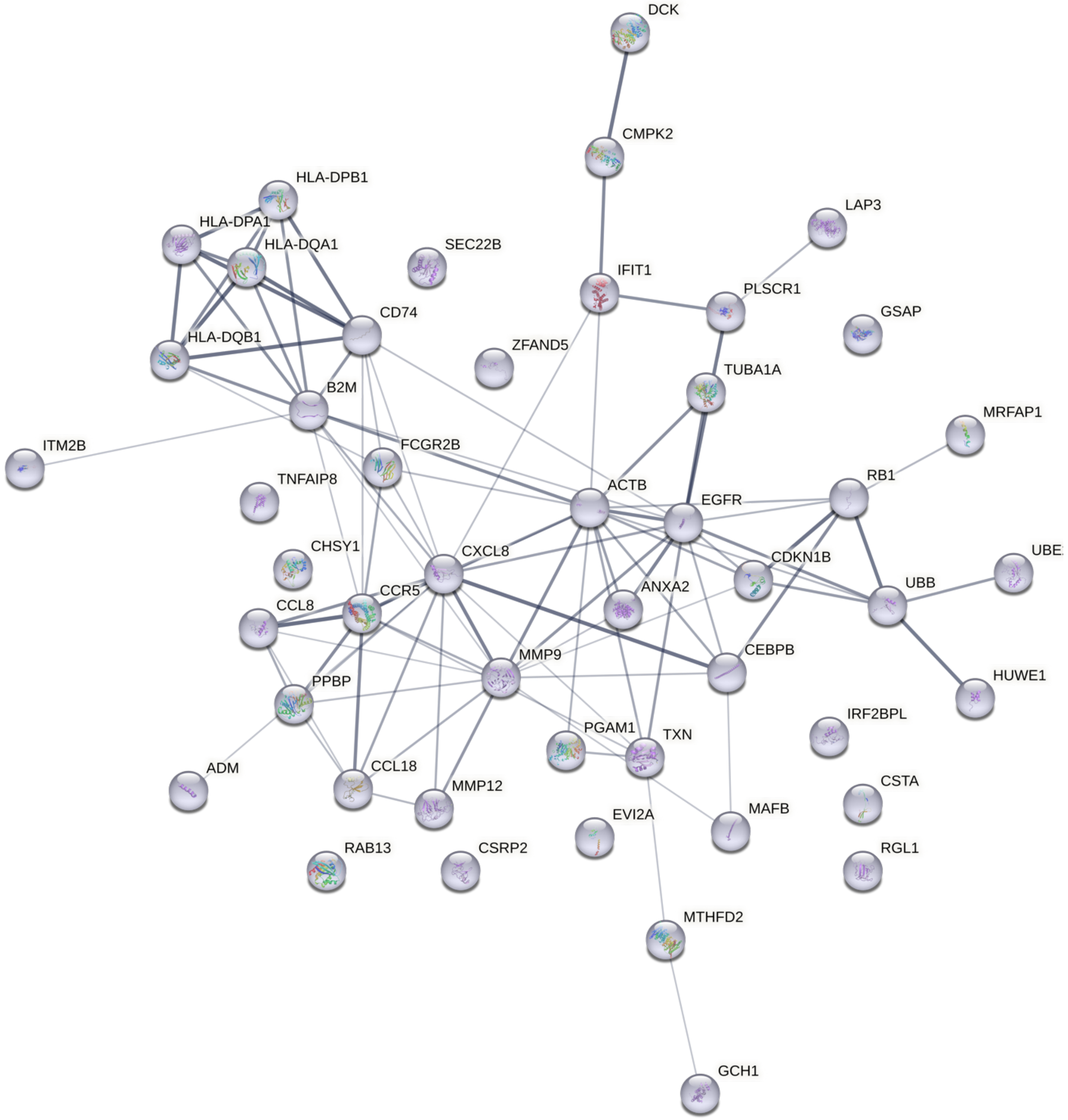

S8.3

#### Cytokine Production

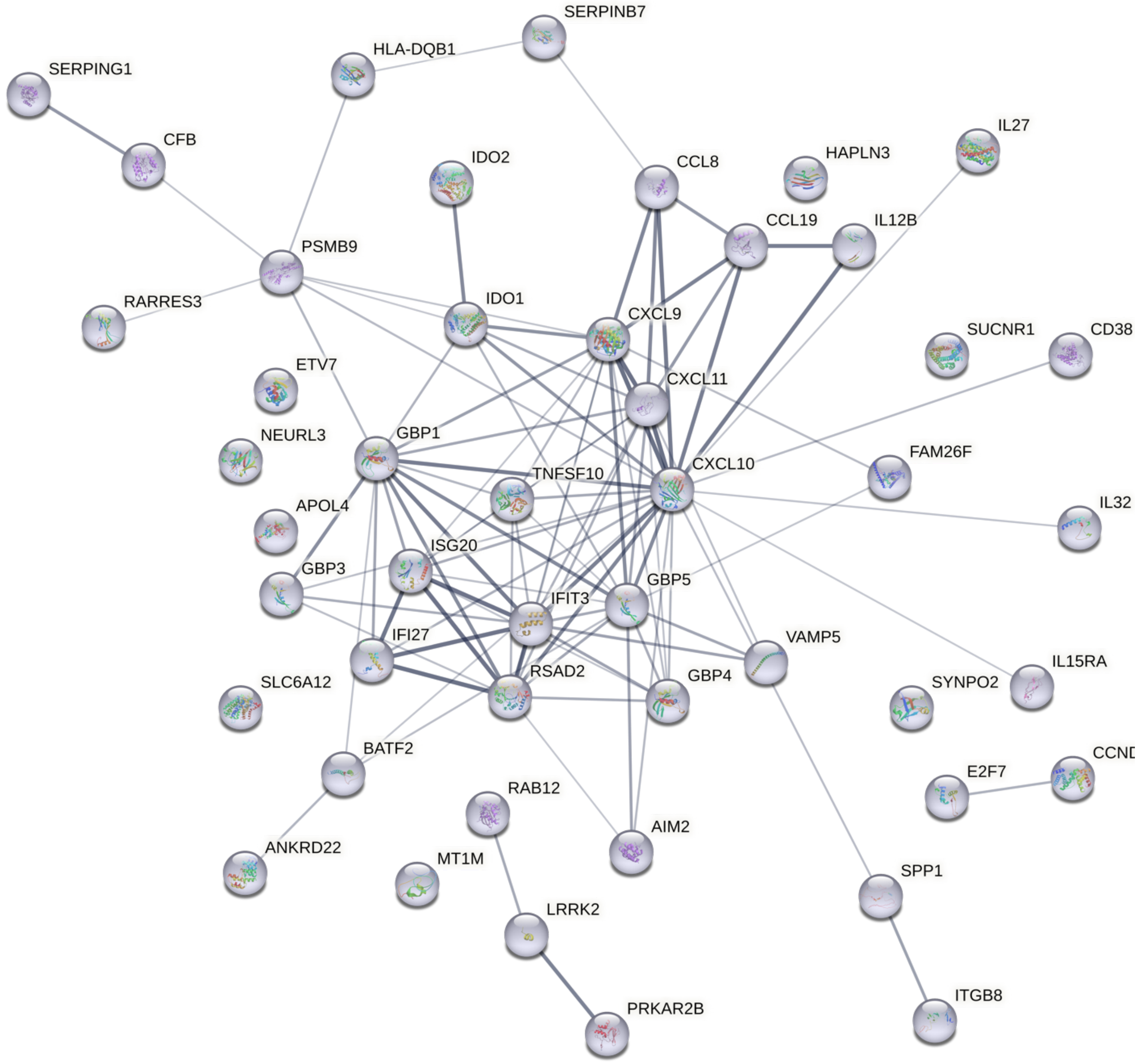

S8.4

### Cell Adhesion_1

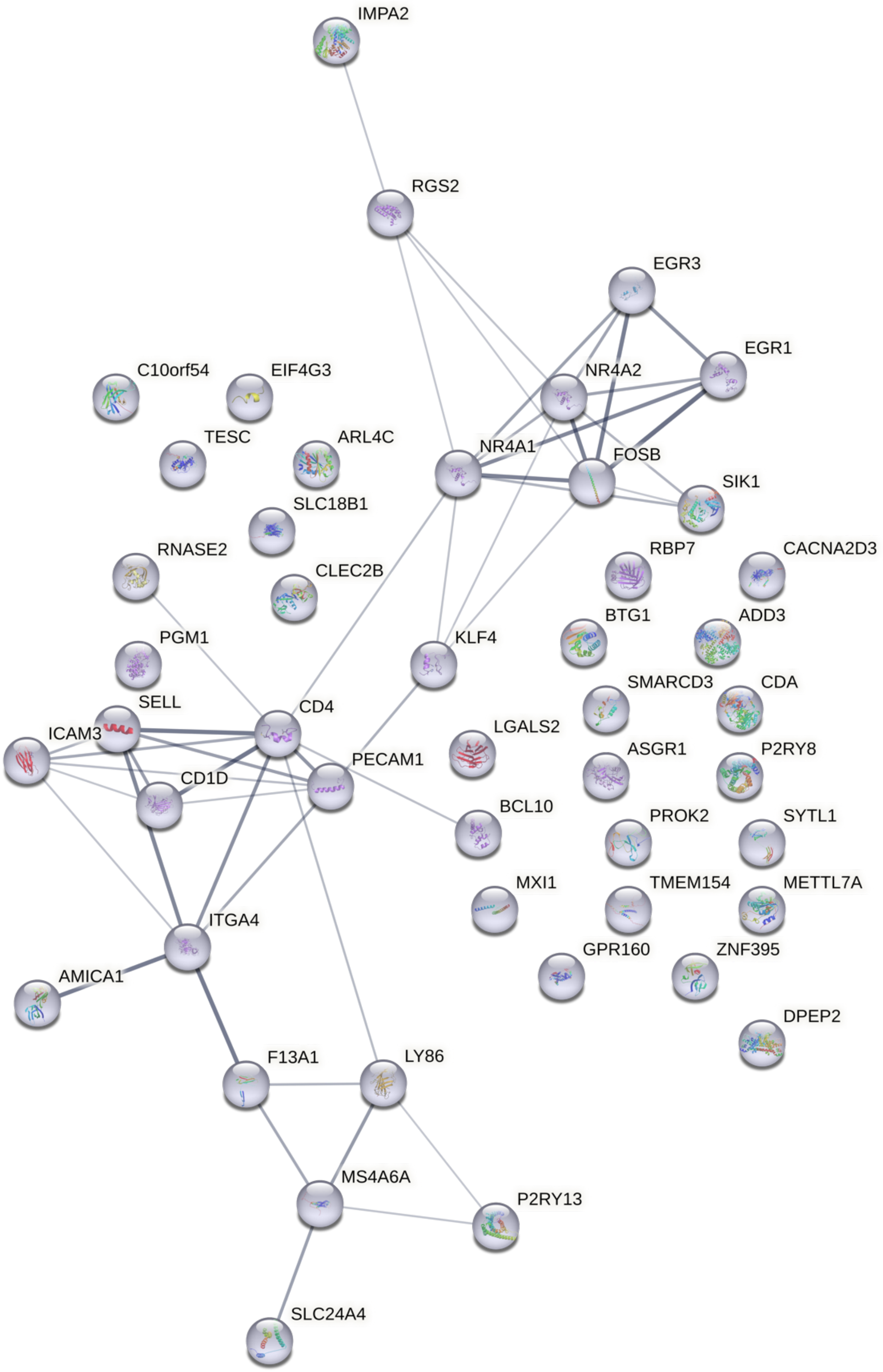

S8.5

### Modulation of Cell Migration

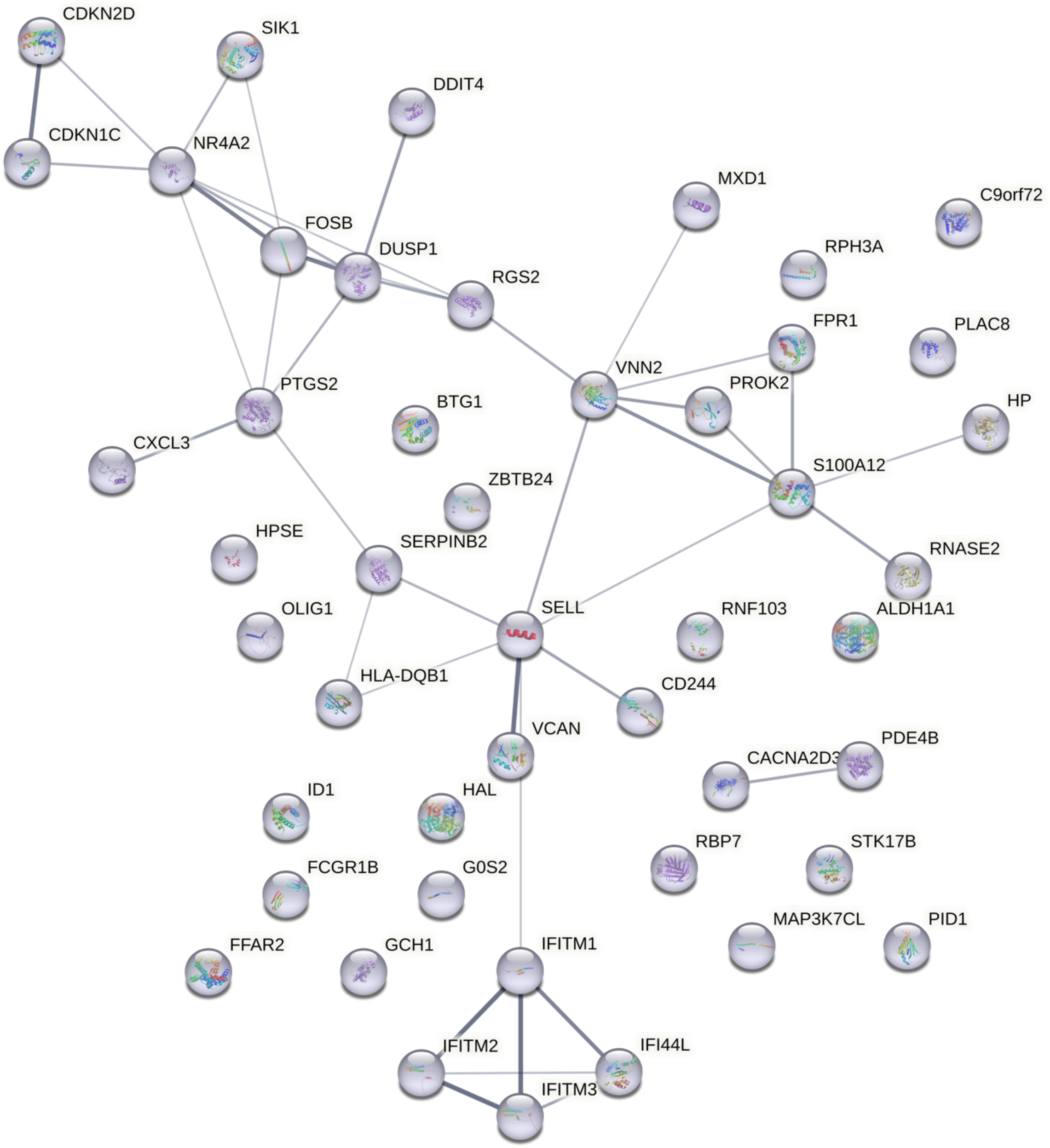

S8.6

#### Cell Adhesion_2

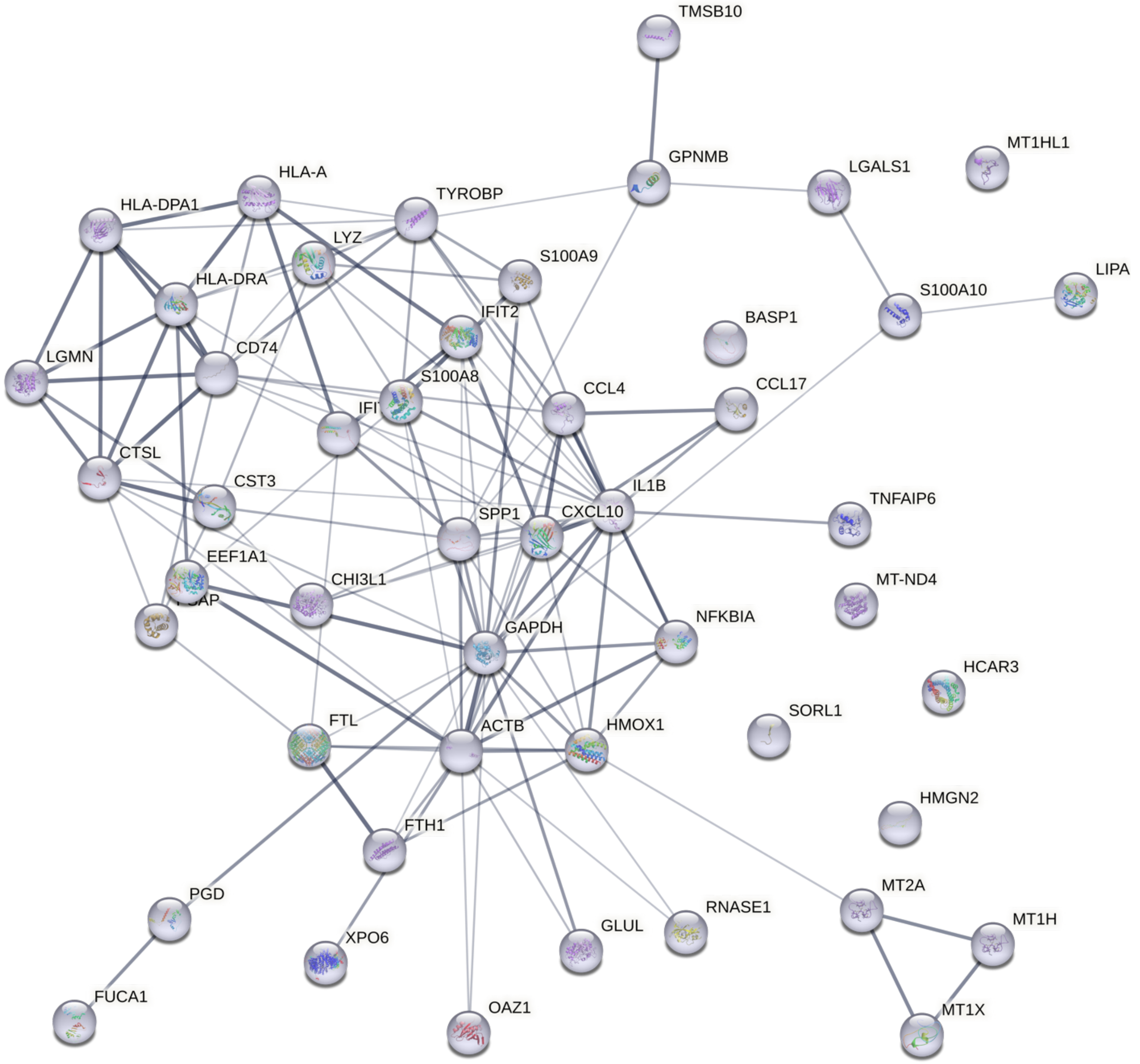

S8.7

### Eosinophil Chemotaxis

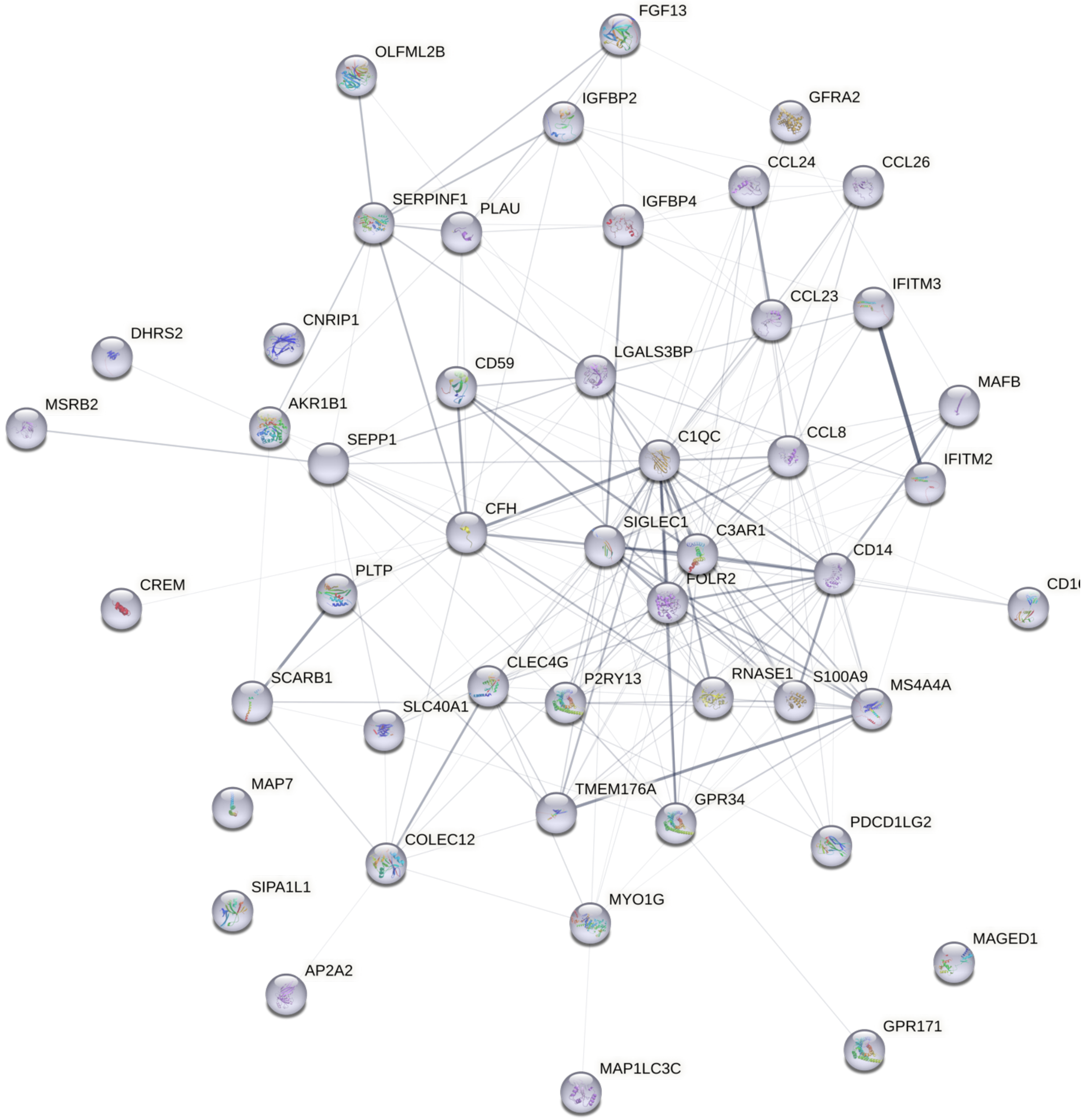

S8.8

### MHC2-mediated Lymphocyte Activation

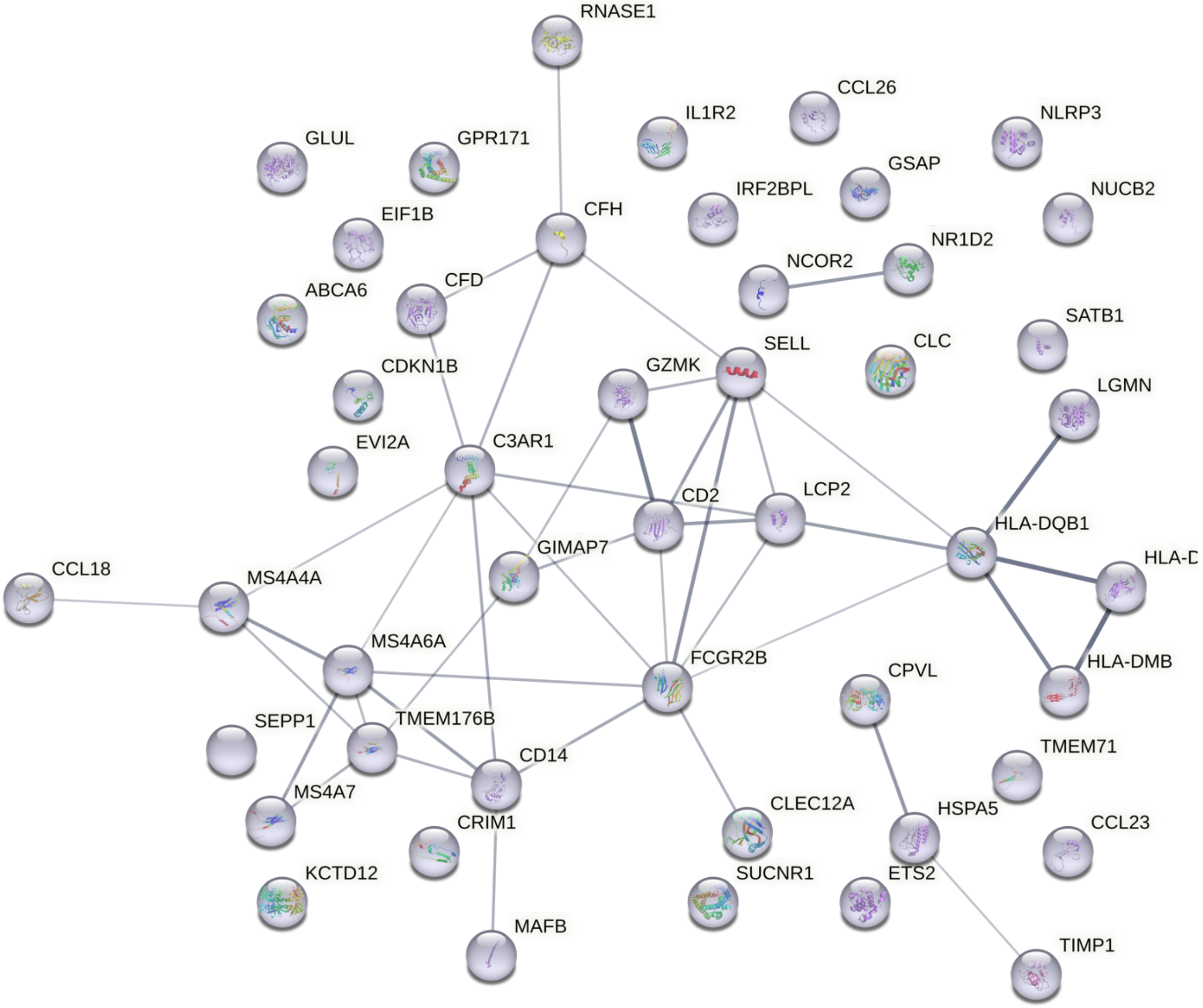

S8.9

### Chemokine Activity

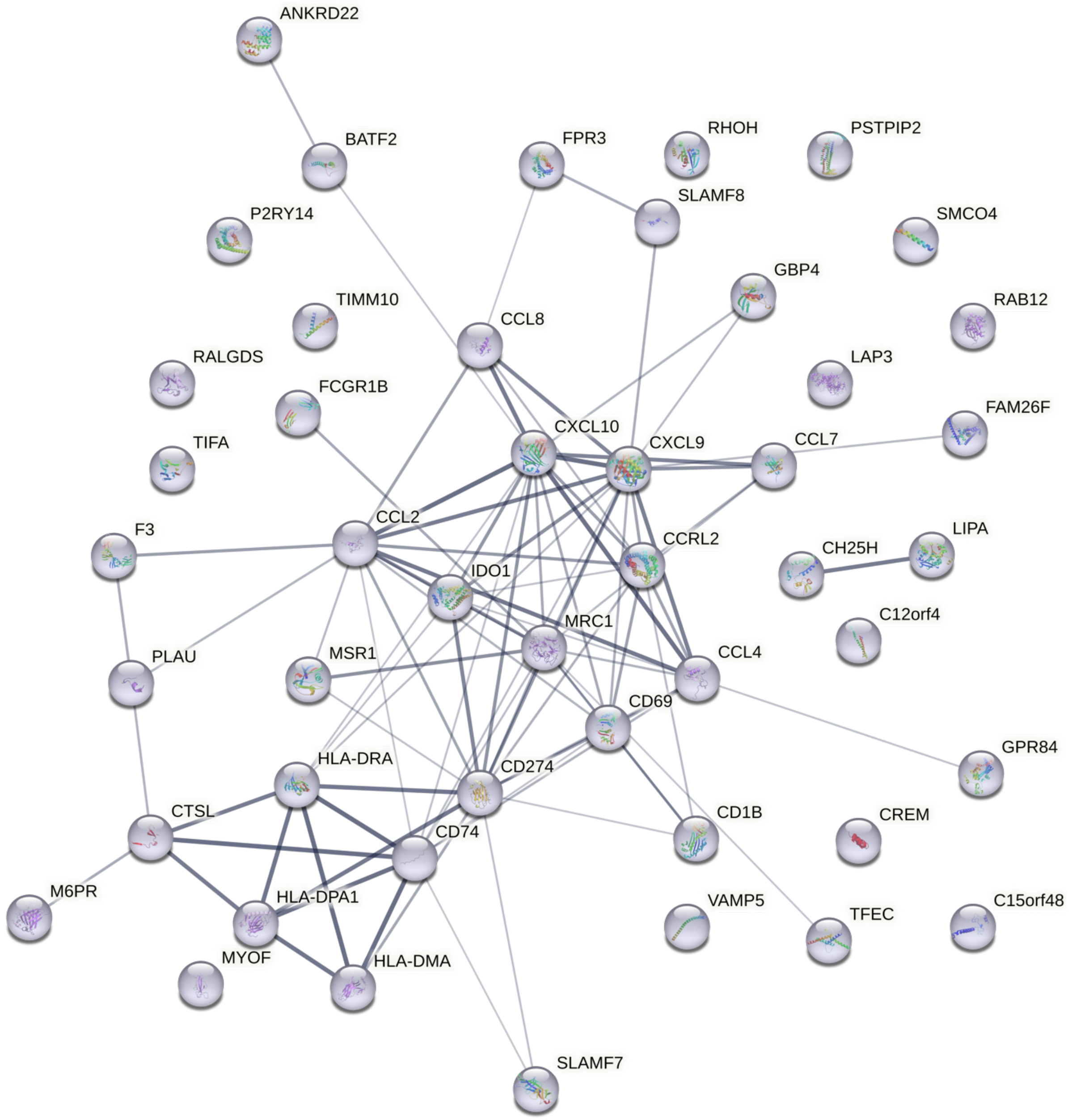

**Figure S8. [Protein-protein interaction network], Related to Figure 1.**
Protein-protein interaction network for all 9 myeloid -derived irMPs. Each node represents a protein produced by a single, protein-coding gene locus. The protein 3D structure is known if the node is filled with a miniature 3D structure. Edge thickness refers to the level of confidence, with thicker edges indicating more confident protein interaction, and such interactions could come from any of the following: text mining, gene co-expression, neighborhood, gene fusion or co-occurrence.

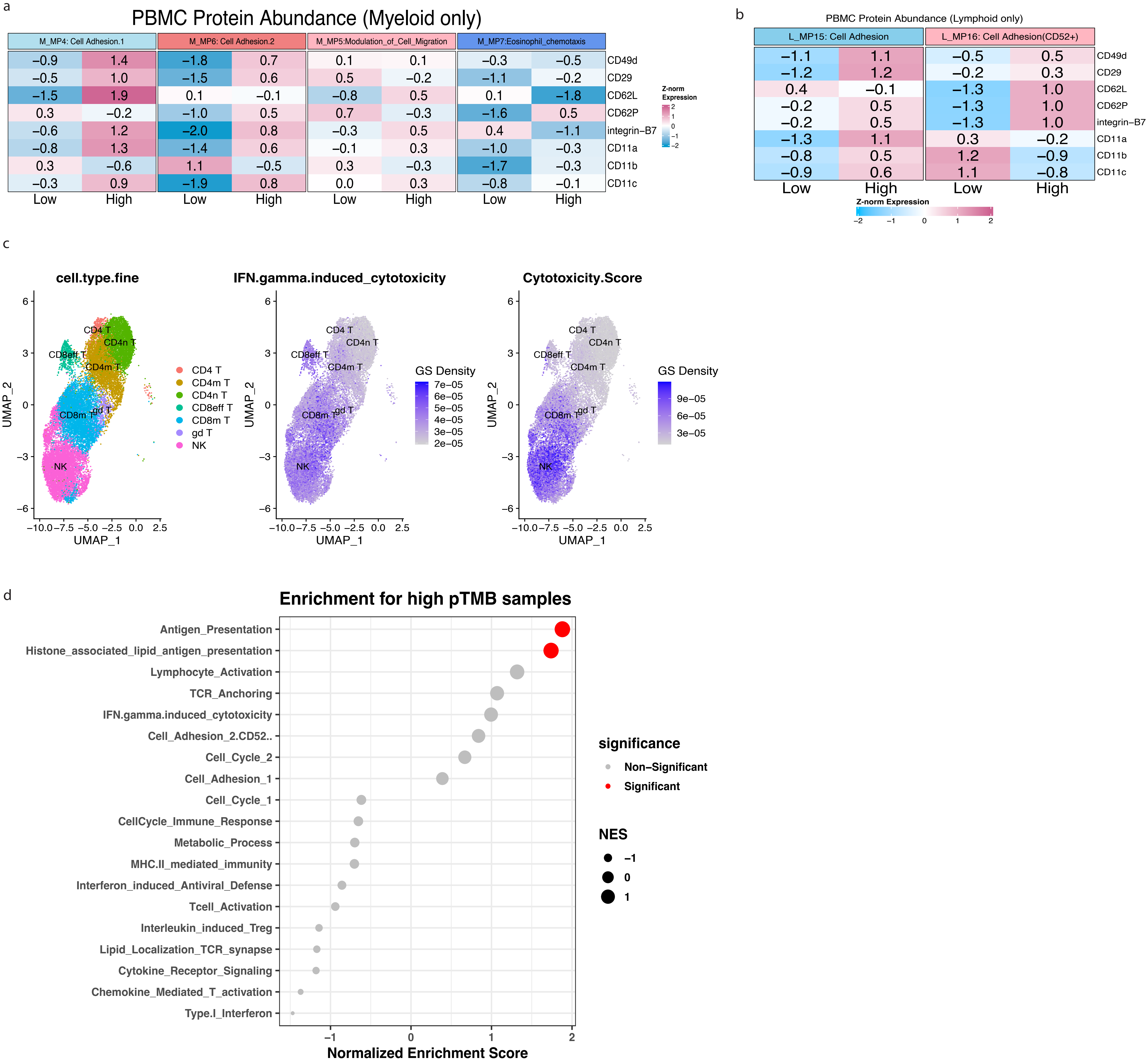
a
PBMC Protein Abundance (Myeloid only)
M_MP4: Cell Adhesion.1
M_MP6: Cell Adhesion.2
M_MP5:Modulation_of_Cell_Migration
M_MP7:Eosinophil_chemotaxis
−0.9 1.4 −1.8 0.7 0.1 0.1 −0.3 −0.5 CD49d
−0.5 1.0 −1.5 0.6 0.5 −0.2 −1.1 −0.2 CD29
−1.5 1.9 0.1 −0.1 −0.8 0.5 0.1 −1.8 CD62L
0.3 −0.2 −1.0 0.5 0.7 −0.3 −1.6 0.5 CD62P
−0.6 1.2 −2.0 0.8 −0.3 0.5 0.4 −1.1 integrin−B7
−0.8 1.3 −1.4 0.6 −0.1 0.3 −1.0 −0.3 CD11a
0.3 −0.6 1.1 −0.5 0.3 −0.3 −1.7 −0.3 CD11b
−0.3 0.9 −1.9 0.8 0.0 0.3 −0.8 −0.1 CD11c
Low High Low High Low High Low High
Z-norm Expression
2 1 0 −1 −2
b
PBMC Protein Abundance (Lymphoid only)
L_MP15: Cell Adhesion
L_MP16: Cell Adhesion(CD52+)
−1.1 1.1 −0.5 0.5 CD49d
−1.2 1.2 −0.2 0.3 CD29
0.4 −0.1 −1.3 1.0 CD62L
−0.2 0.5 −1.3 1.0 CD62P
−0.2 0.5 −1.3 1.0 integrin−B7
−1.3 1.1 0.3 −0.2 CD11a
−0.8 0.5 1.2 −0.9 CD11b
−0.9 0.6 1.1 −0.8 CD11c
Low High Low High
Z-norm Expression
−2 −1 0 1 2
c
cell.type.fine
IFN.gamma.induced_cytotoxicity
Cytotoxicity.Score
CD4 T
CD4n T
CD8eff T
CD4m T
CD8m T
gd T
NK
CD4 T
CD4m T
CD4n T
CD8eff T
CD8m T
gd T
NK
GS Density
7e−05
6e−05
5e−05
4e−05
3e−05
2e−05
GS Density
9e−05
6e−05
3e−05
UMAP_1
UMAP_2
−10.0 −7.5 −5.0 −2.5 0.0 2.5
6 3 0 −3 −6
d
Enrichment for high pTMB samples
Antigen_Presentation
Histone_associated_lipid_antigen_presentation
Lymphocyte_Activation
TCR_Anchoring
IFN.gamma.induced_cytotoxicity
Cell_Adhesion_2.CD52..
Cell_Cycle_2
Cell_Adhesion_1
Cell_Cycle_1
CellCycle_Immune_Response
Metabolic_Process
MHC.II_mediated_immunity
Interferon_induced_Antiviral_Defense
Tcell_Activation
Interleukin_induced_Treg
Lipid_Localization_TCR_synapse
Cytokine_Receptor_Signaling
Chemokine_Mediated_T_activation
Type.I_Interferon
−1 0 1 2
Normalized Enrichment Score
significance
Non−Significant
Significant
NES
−1
0
1

**Figure S9. [Gene set annotation validation], Related to Figure 2 and 3.**

**(a)** PBMC cells were dichotomized into two groups (High vs Low) based on the activity levels of **(a)** M_MP4, 5, 6, 7 and **(b)** L_MP15, L_MP16, estimated respectively from the RNA expression data. Shown in the heatmap table (and colored accordingly) are the average Z-transformed protein abundance of the surface proteins in each of the respective cell groups. **(c)** UMAP of all lymphoid cells from COVID-19 atlas colored by activity level of L_MP17 and a classical cytotoxicity score calculated by GSDensity. **(d)** Gene set enrichment results comparing TCGA samples with high and low persistent tumor mutational burden. Dot sizes represent normalized enrichment score (NES), and colors represent FDR adjusted p-value (red: adjusted p-value<0.05). NES>1 means upregulation.

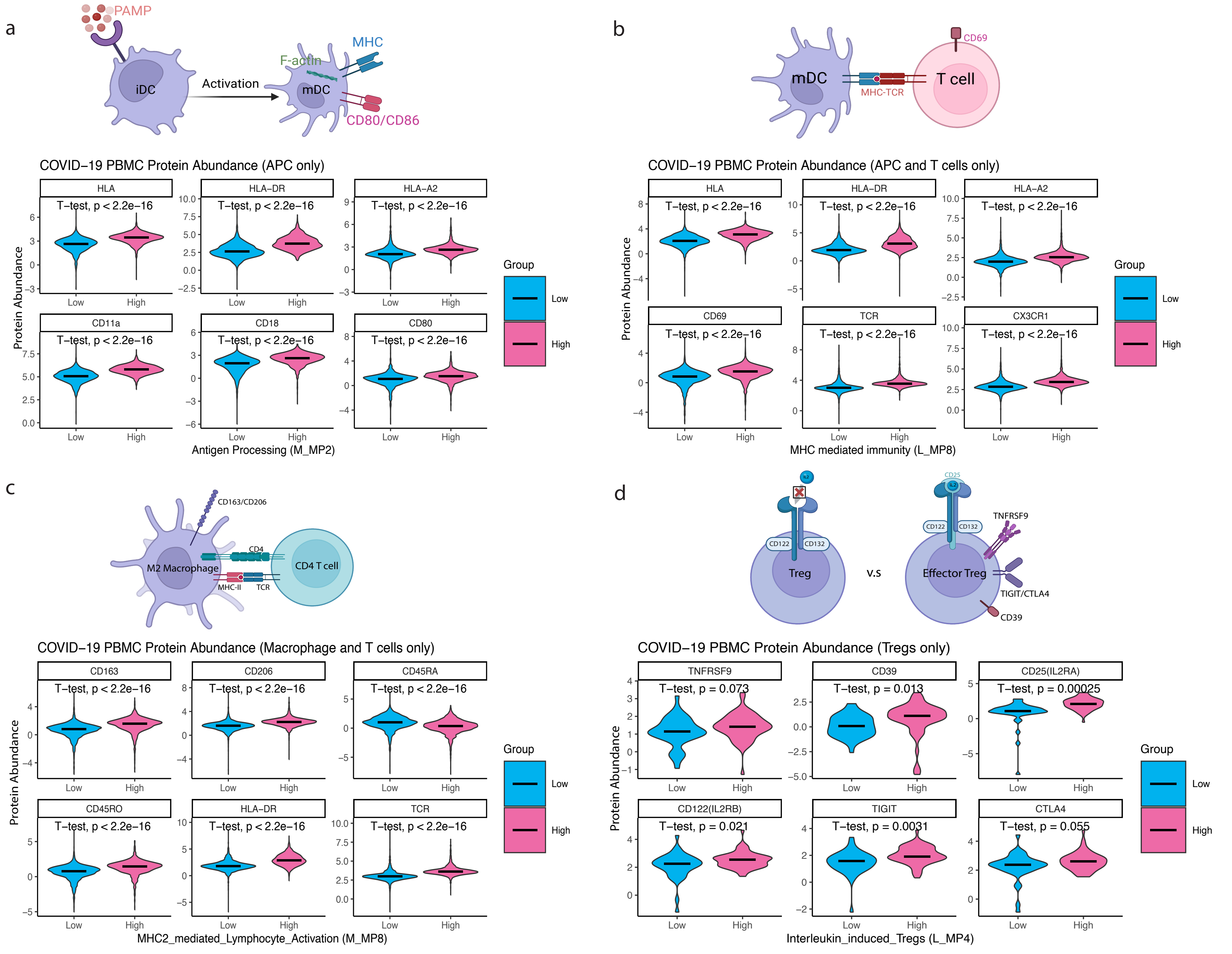
a
PAMP
iDC
Activation
F-actin
MHC
mDC
CD80/CD86
COVID−19 PBMC Protein Abundance (APC only)
HLA
HLA−DR
HLA−A2
CD11a
CD18
CD80
T−test, p < 2.2e−16
Protein Abundance
Low
High
Antigen Processing (M_MP2)
Group
b
mDC
MHC-TCR
T cell
CD69
COVID−19 PBMC Protein Abundance (APC and T cells only)
HLA
HLA−DR
HLA−A2
CD69
TCR
CX3CR1
MHC mediated immunity (L_MP8)
c
CD163/CD206
M2 Macrophage
CD4
CD4 T cell
MHC-II
TCR
COVID−19 PBMC Protein Abundance (Macrophage and T cells only)
CD163
CD206
CD45RA
CD45RO
HLA−DR
TCR
MHC2_mediated_Lymphocyte_Activation (M_MP8)
d
IL2
CD25
CD122
CD132
Treg
v.s
Effector Treg
TNFRSF9
TIGIT/CTLA4
CD39
COVID−19 PBMC Protein Abundance (Tregs only)
TNFRSF9
T−test, p = 0.073
CD39
T−test, p = 0.013
CD25(IL2RA)
T−test, p = 0.00025
CD122(IL2RB)
T−test, p = 0.021
TIGIT
T−test, p = 0.0031
CTLA4
T−test, p = 0.055
Interleukin_induced_Tregs (L_MP4)

**Figure S10. [Gene set annotation validation], Related to Figure 2 and 3.**

**(a)-(d)** Diagrams for p-MHC interface/pMHC-TCR interface/Treg activation. Violin plots are the abundance for selected surface protein markers comparing two dichotomized COVID-19 PBMC cell groups (Dichotomization of cells are based on ssGSEA of respective irMPs in the RNA level) (Different subsets of cells were kept, detailed on figure titles). Protein abundance between two groups was compared using T test with significance level 0.05.

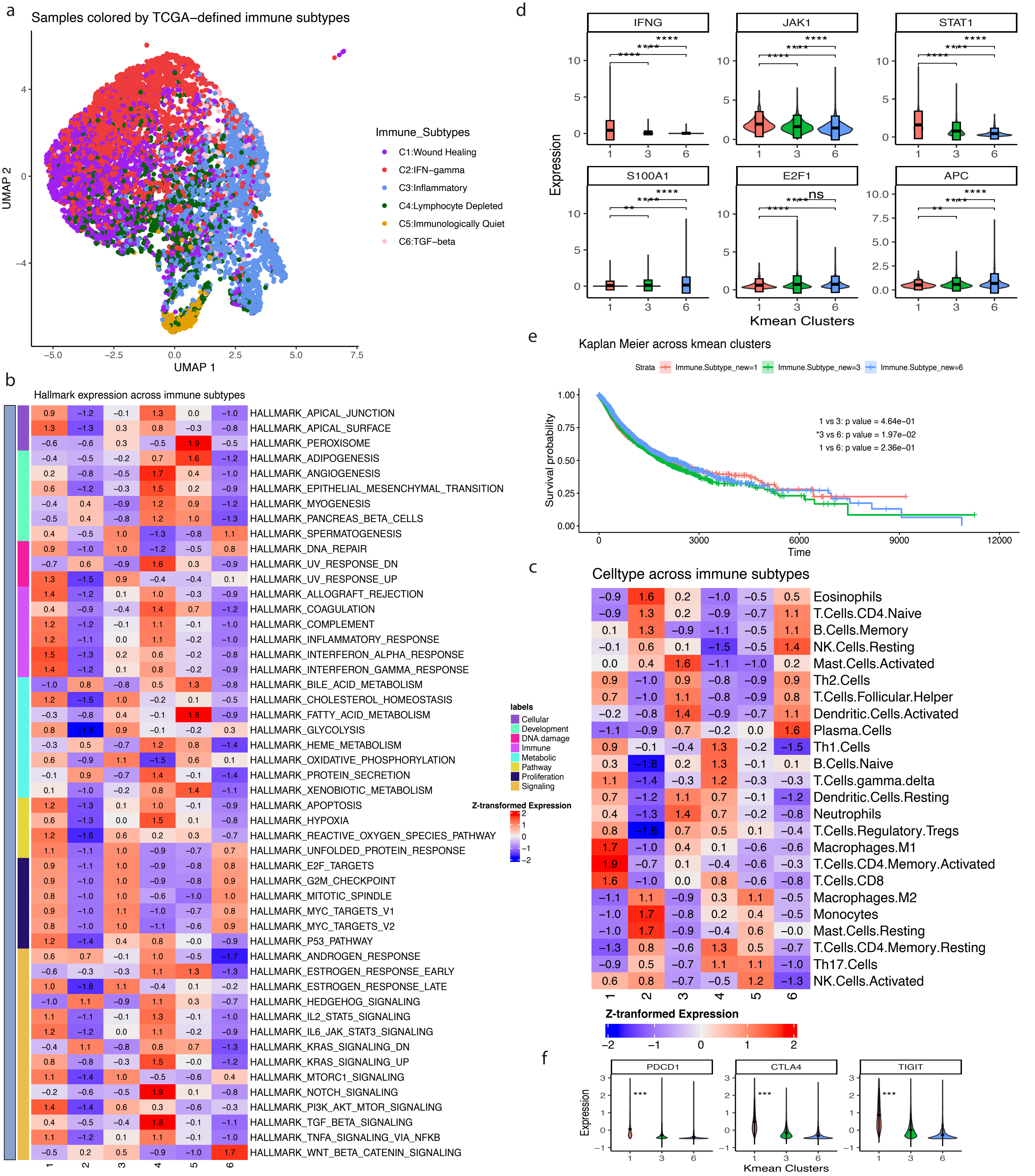

a
Samples colored by TCGA−defined immune subtypes
UMAP 2
UMAP 1
Immune_Subtypes
C1:Wound Healing
C2:IFN−gamma
C3:Inflammatory
C4:Lymphocyte Depleted
C5:Immunologically Quiet
C6:TGF−beta
b
Hallmark expression across immune subtypes
HALLMARK_APICAL_JUNCTION
HALLMARK_APICAL_SURFACE
HALLMARK_PEROXISOME
HALLMARK_ADIPOGENESIS
HALLMARK_ANGIOGENESIS
HALLMARK_EPITHELIAL_MESENCHYMAL_TRANSITION
HALLMARK_MYOGENESIS
HALLMARK_PANCREAS_BETA_CELLS
HALLMARK_SPERMATOGENESIS
HALLMARK_DNA_REPAIR
HALLMARK_UV_RESPONSE_DN
HALLMARK_UV_RESPONSE_UP
HALLMARK_ALLOGRAFT_REJECTION
HALLMARK_COAGULATION
HALLMARK_COMPLEMENT
HALLMARK_INFLAMMATORY_RESPONSE
HALLMARK_INTERFERON_ALPHA_RESPONSE
HALLMARK_INTERFERON_GAMMA_RESPONSE
HALLMARK_BILE_ACID_METABOLISM
HALLMARK_CHOLESTEROL_HOMEOSTASIS
HALLMARK_FATTY_ACID_METABOLISM
HALLMARK_GLYCOLYSIS
HALLMARK_HEME_METABOLISM
HALLMARK_OXIDATIVE_PHOSPHORYLATION
HALLMARK_PROTEIN_SECRETION
HALLMARK_XENOBIOTIC_METABOLISM
HALLMARK_APOPTOSIS
HALLMARK_HYPOXIA
HALLMARK_REACTIVE_OXYGEN_SPECIES_PATHWAY
HALLMARK_UNFOLDED_PROTEIN_RESPONSE
HALLMARK_E2F_TARGETS
HALLMARK_G2M_CHECKPOINT
HALLMARK_MITOTIC_SPINDLE
HALLMARK_MYC_TARGETS_V1
HALLMARK_MYC_TARGETS_V2
HALLMARK_P53_PATHWAY
HALLMARK_ANDROGEN_RESPONSE
HALLMARK_ESTROGEN_RESPONSE_EARLY
HALLMARK_ESTROGEN_RESPONSE_LATE
HALLMARK_HEDGEHOG_SIGNALING
HALLMARK_IL2_STAT5_SIGNALING
HALLMARK_IL6_JAK_STAT3_SIGNALING
HALLMARK_KRAS_SIGNALING_DN
HALLMARK_KRAS_SIGNALING_UP
HALLMARK_MTORC1_SIGNALING
HALLMARK_NOTCH_SIGNALING
HALLMARK_PI3K_AKT_MTOR_SIGNALING
HALLMARK_TGF_BETA_SIGNALING
HALLMARK_TNFA_SIGNALING_VIA_NFKB
HALLMARK_WNT_BETA_CATENIN_SIGNALING
labels
Cellular
Development
DNA.damage
Immune
Metabolic
Pathway
Proliferation
Signaling
Z-transformed Expression
d
IFNG
JAK1
STAT1
S100A1
E2F1
APC
Expression
Kmean Clusters
e
Kaplan Meier across kmean clusters
Strata
Immune.Subtype_new=1
Immune.Subtype_new=3
Immune.Subtype_new=6
1 vs 3: p value = 4.64e−01
*3 vs 6: p value = 1.97e−02
1 vs 6: p value = 2.36e−01
Survival probability
Time
c
Celltype across immune subtypes
Eosinophils
T.Cells.CD4.Naive
B.Cells.Memory
NK.Cells.Resting
Mast.Cells.Activated
Th2.Cells
T.Cells.Follicular.Helper
Dendritic.Cells.Activated
Plasma.Cells
Th1.Cells
B.Cells.Naive
T.Cells.gamma.delta
Dendritic.Cells.Resting
Neutrophils
T.Cells.Regulatory.Tregs
Macrophages.M1
T.Cells.CD4.Memory.Activated
T.Cells.CD8
Macrophages.M2
Monocytes
Mast.Cells.Resting
T.Cells.CD4.Memory.Resting
Th17.Cells
NK.Cells.Activated
Z-tranformed Expression
f
PDCD1
CTLA4
TIGIT
Expression
Kmean Clusters

**Figure S11. [Gene sets refine pan-cancer immune subtypes], Related to Figure 4.**

**(a)** UMAP of all TCGA samples (n=10128) using single sample gene-set enrichment score calculated from the 28 irMPs. Each dot is a TCGA sample with colors that represent different TCGA immune subtypes. (**b**) Heatmap comparing the Z-transformed expression of 50 Hallmark pathways across the six k-mean clusters. Rows are clustered based on Hallmark supergroups. (**c**) Heatmap comparing the Z-transformed expression of TCGA immune cell type scores across the six k-mean clusters. Rows are clustered based on hierarchical clustering. **(d)** Violin plot comparing gene expression of IFN-gamma associated genes (top) and wound-healing associated genes (bottom) across k-mean cluster 1, 3 and 6. **(e)** Kaplan Meier Overall Survival curves across the k-mean clusters 1, 3, and 6. Log-rank test is performed between pairs of interests. **(f)** Violin plot comparing gene expression of classical exhaustion markers across k-mean cluster 1, 3 and 6. Gene expression between any two groups was compared using T test with significance level 0.05.

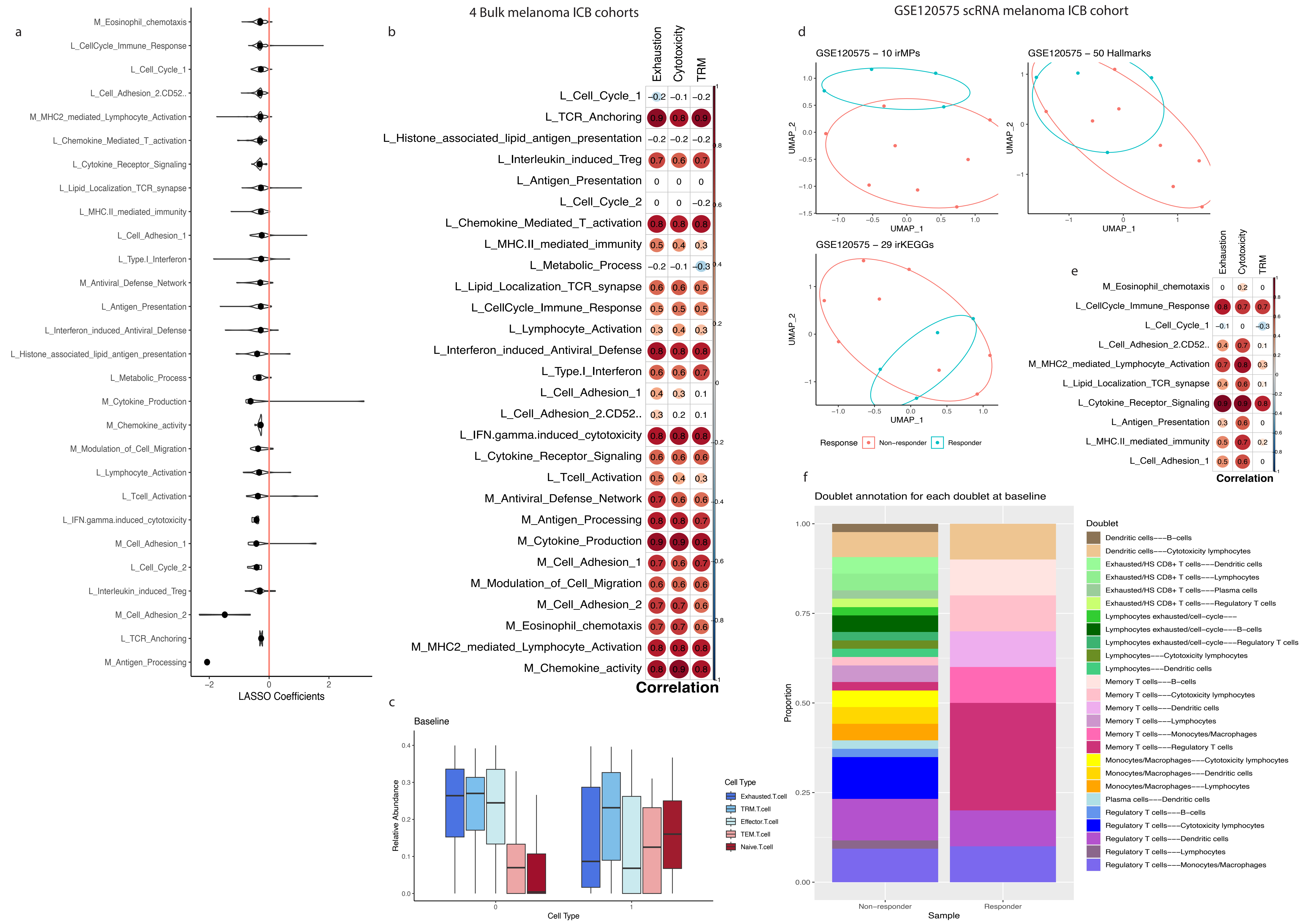
a
M_Eosinophil_chemotaxis
L_CellCycle_Immune_Response
L_Cell_Cycle_1
L_Cell_Adhesion_2.CD52..
M_MHC2_mediated_Lymphocyte_Activation
L_Chemokine_Mediated_T_activation
L_Cytokine_Receptor_Signaling
L_Lipid_Localization_TCR_synapse
L_MHC.II_mediated_immunity
L_Cell_Adhesion_1
L_Type.I_Interferon
M_Antiviral_Defense_Network
L_Antigen_Presentation
L_Interferon_induced_Antiviral_Defense
L_Histone_associated_lipid_antigen_presentation
L_Metabolic_Process
M_Cytokine_Production
M_Chemokine_activity
M_Modulation_of_Cell_Migration
L_Lymphocyte_Activation
L_Tcell_Activation
L_IFN.gamma.induced_cytotoxicity
M_Cell_Adhesion_1
L_Cell_Cycle_2
L_Interleukin_induced_Treg
M_Cell_Adhesion_2
L_TCR_Anchoring
M_Antigen_Processing
−2
0
2
LASSO Coefficients
4 Bulk melanoma ICB cohorts
b
Exhaustion
Cytotoxicity
TRM
L_Cell_Cycle_1 −0.2 −0.1 −0.2
L_TCR_Anchoring 0.9 0.8 0.9
L_Histone_associated_lipid_antigen_presentation −0.2 −0.2 −0.2
L_Interleukin_induced_Treg 0.7 0.6 0.7
L_Antigen_Presentation 0 0 0
L_Cell_Cycle_2 0 0 −0.2
L_Chemokine_Mediated_T_activation 0.8 0.8 0.8
L_MHC.II_mediated_immunity 0.5 0.4 0.3
L_Metabolic_Process −0.2 −0.1 −0.3
L_Lipid_Localization_TCR_synapse 0.6 0.6 0.5
L_CellCycle_Immune_Response 0.5 0.5 0.5
L_Lymphocyte_Activation 0.3 0.4 0.3
L_Interferon_induced_Antiviral_Defense 0.8 0.8 0.8
L_Type.I_Interferon 0.6 0.6 0.7
L_Cell_Adhesion_1 0.4 0.3 0.1
L_Cell_Adhesion_2.CD52.. 0.3 0.2 0.1
L_IFN.gamma.induced_cytotoxicity 0.8 0.8 0.8
L_Cytokine_Receptor_Signaling 0.6 0.6 0.6
L_Tcell_Activation 0.5 0.4 0.3
M_Antiviral_Defense_Network 0.7 0.6 0.6
M_Antigen_Processing 0.8 0.8 0.7
M_Cytokine_Production 0.9 0.9 0.8
M_Cell_Adhesion_1 0.7 0.6 0.7
M_Modulation_of_Cell_Migration 0.6 0.6 0.6
M_Cell_Adhesion_2 0.7 0.7 0.6
M_Eosinophil_chemotaxis 0.7 0.7 0.6
M_MHC2_mediated_Lymphocyte_Activation 0.8 0.8 0.8
M_Chemokine_activity 0.8 0.9 0.8
Correlation
c
Baseline
Relative Abundance
Cell Type
0
1
Cell Type
Exhausted.T.cell
TRM.T.cell
Effector.T.cell
TEM.T.cell
Naive.T.cell
GSE120575 scRNA melanoma ICB cohort
d
GSE120575 – 10 irMPs
GSE120575 – 50 Hallmarks
GSE120575 – 29 irKEGGs
UMAP_1
UMAP_2
Response
Non−responder
Responder
e
Exhaustion
Cytotoxicity
TRM
M_Eosinophil_chemotaxis 0 0.2 0
L_CellCycle_Immune_Response 0.8 0.7 0.7
L_Cell_Cycle_1 −0.1 0 −0.3
L_Cell_Adhesion_2.CD52.. 0.4 0.7 0.1
M_MHC2_mediated_Lymphocyte_Activation 0.7 0.8 0.3
L_Lipid_Localization_TCR_synapse 0.4 0.6 0.1
L_Cytokine_Receptor_Signaling 0.9 0.9 0.8
L_Antigen_Presentation 0.3 0.6 0
L_MHC.II_mediated_immunity 0.5 0.7 0.2
L_Cell_Adhesion_1 0.5 0.6 0
Correlation
f
Doublet annotation for each doublet at baseline
Proportion
Non−responder
Responder
Sample
Doublet
Dendritic cells−−−B−cells
Dendritic cells−−−Cytotoxicity lymphocytes
Exhausted/HS CD8+ T cells−−−Dendritic cells
Exhausted/HS CD8+ T cells−−−Lymphocytes
Exhausted/HS CD8+ T cells−−−Plasma cells
Exhausted/HS CD8+ T cells−−−Regulatory T cells
Lymphocytes exhausted/cell−cycle−−−
Lymphocytes exhausted/cell−cycle−−−B−cells
Lymphocytes exhausted/cell−cycle−−−Regulatory T cells
Lymphocytes−−−Cytotoxicity lymphocytes
Lymphocytes−−−Dendritic cells
Memory T cells−−−B−cells
Memory T cells−−−Cytotoxicity lymphocytes
Memory T cells−−−Dendritic cells
Memory T cells−−−Lymphocytes
Memory T cells−−−Monocytes/Macrophages
Memory T cells−−−Regulatory T cells
Monocytes/Macrophages−−−Cytotoxicity lymphocytes
Monocytes/Macrophages−−−Dendritic cells
Monocytes/Macrophages−−−Lymphocytes
Plasma cells−−−Dendritic cells
Regulatory T cells−−−B−cells
Regulatory T cells−−−Cytotoxicity lymphocytes
Regulatory T cells−−−Dendritic cells
Regulatory T cells−−−Lymphocytes
Regulatory T cells−−−Monocytes/Macrophages

**Figure S12. [Gene sets improve ICB response prediction], Related to Figure 5.**
**(a)** Distribution of fitted coefficients across 1000 LASSO regression for each of the irMP. **(b)** Correlation heatmap showing the correlation between activity of irMPs and immune-relative signatures at baseline, calculated from ssGSEA. Significant correlations are shown with colored dots. **(c)** Grouped boxplot comparing relative abundance of selected T cell subsets between responders and non-responders at baseline. **(d)** UMAP derived from 10 most selected irMPs activities (top left), 50 Hallmarks (top right), and 29 irKEGGs (bottom left). Color represents different responses. **(e)** Correlation heatmap showing the correlation between activity of top 10 selected irMPs and immune-relative signatures at baseline, calculated from ssGSEA. Significant correlations are shown with colored dots. **(f)** Stacked barplot comparing the relevant proportion of different types of doublets between responders and non-responders at baseline.

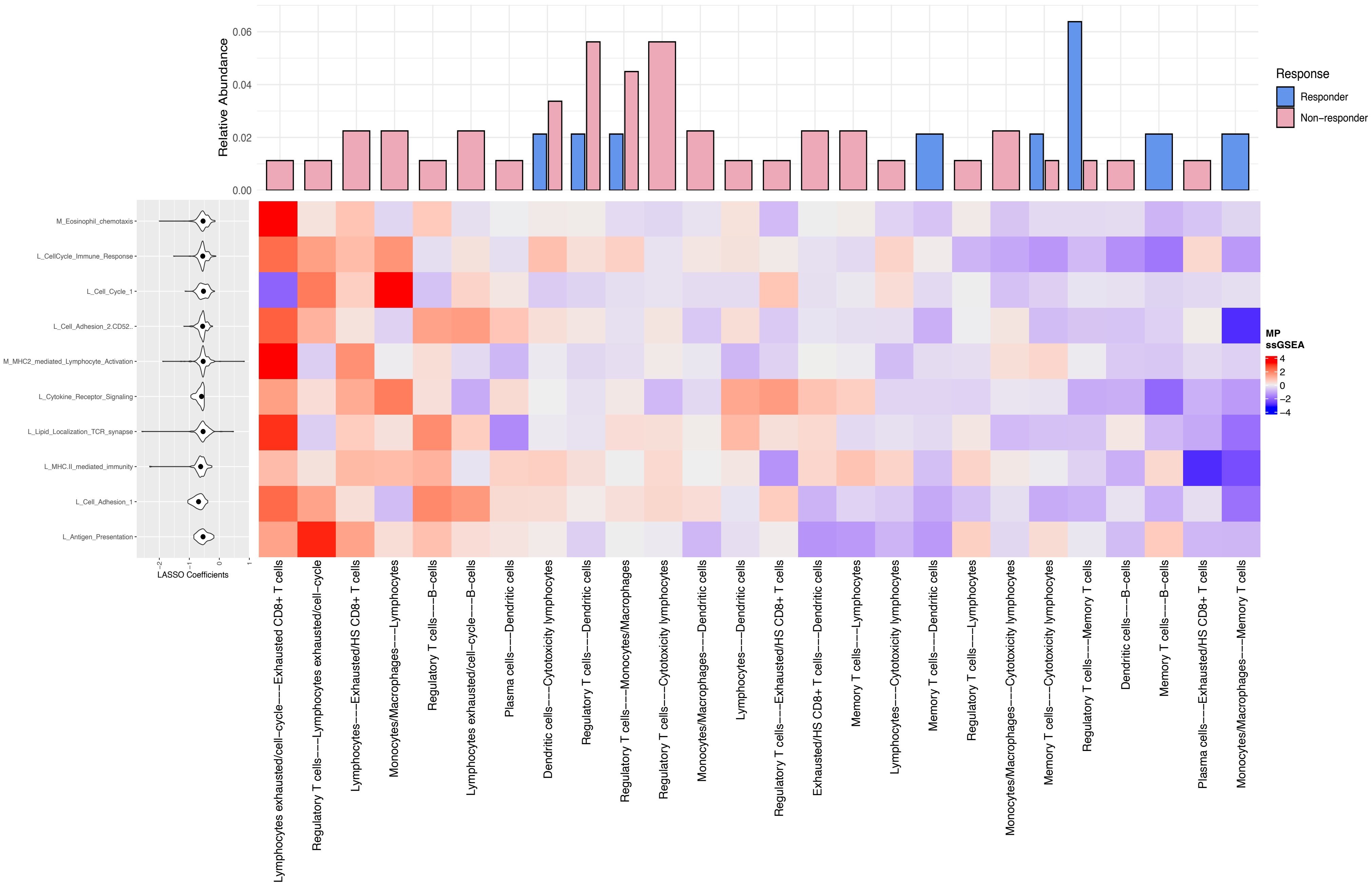

Relative Abundance
0.00
0.02
0.04
0.06
Response
Responder
Non−responder
MP
ssGSEA
4
2
0
−2
−4
M_Eosinophil_chemotaxis
L_CellCycle_Immune_Response
L_Cell_Cycle_1
L_Cell_Adhesion_2.CD52..
M_MHC2_mediated_Lymphocyte_Activation
L_Cytokine_Receptor_Signaling
L_Lipid_Localization_TCR_synapse
L_MHC.II_mediated_immunity
L_Cell_Adhesion_1
L_Antigen_Presentation
-2
-1
0
1
LASSO Coefficients
Lymphocytes exhausted/cell−cycle−−−Exhausted CD8+ T cells
Regulatory T cells−−−Lymphocytes exhausted/cell−cycle
Lymphocytes−−−Exhausted/HS CD8+ T cells
Monocytes/Macrophages−−−Lymphocytes
Regulatory T cells−−−B−cells
Lymphocytes exhausted/cell−cycle−−−B−cells
Plasma cells−−−Dendritic cells
Dendritic cells−−−Cytotoxicity lymphocytes
Regulatory T cells−−−Dendritic cells
Regulatory T cells−−−Monocytes/Macrophages
Regulatory T cells−−−Cytotoxicity lymphocytes
Monocytes/Macrophages−−−Dendritic cells
Lymphocytes−−−Dendritic cells
Regulatory T cells−−−Exhausted/HS CD8+ T cells
Exhausted/HS CD8+ T cells−−−Dendritic cells
Memory T cells−−−Lymphocytes
Lymphocytes−−−Cytotoxicity lymphocytes
Memory T cells−−−Dendritic cells
Regulatory T cells−−−Lymphocytes
Monocytes/Macrophages−−−Cytotoxicity lymphocytes
Memory T cells−−−Cytotoxicity lymphocytes
Regulatory T cells−−−Memory T cells
Dendritic cells−−−B−cells
Memory T cells−−−B−cells
Plasma cells−−−Exhausted/HS CD8+ T cells
Monocytes/Macrophages−−−Memory T cells

**Figure S13. [Gene sets improve ICB response prediction], Related to Figure 5.**

The average fitted coefficients for top 10 selected irMPs across 1000 iterations. Gene set activities for each type of doublets, calculated from ssGSEA. The histogram on top represents the relative abundance of each doublet type between responders and non-responders.

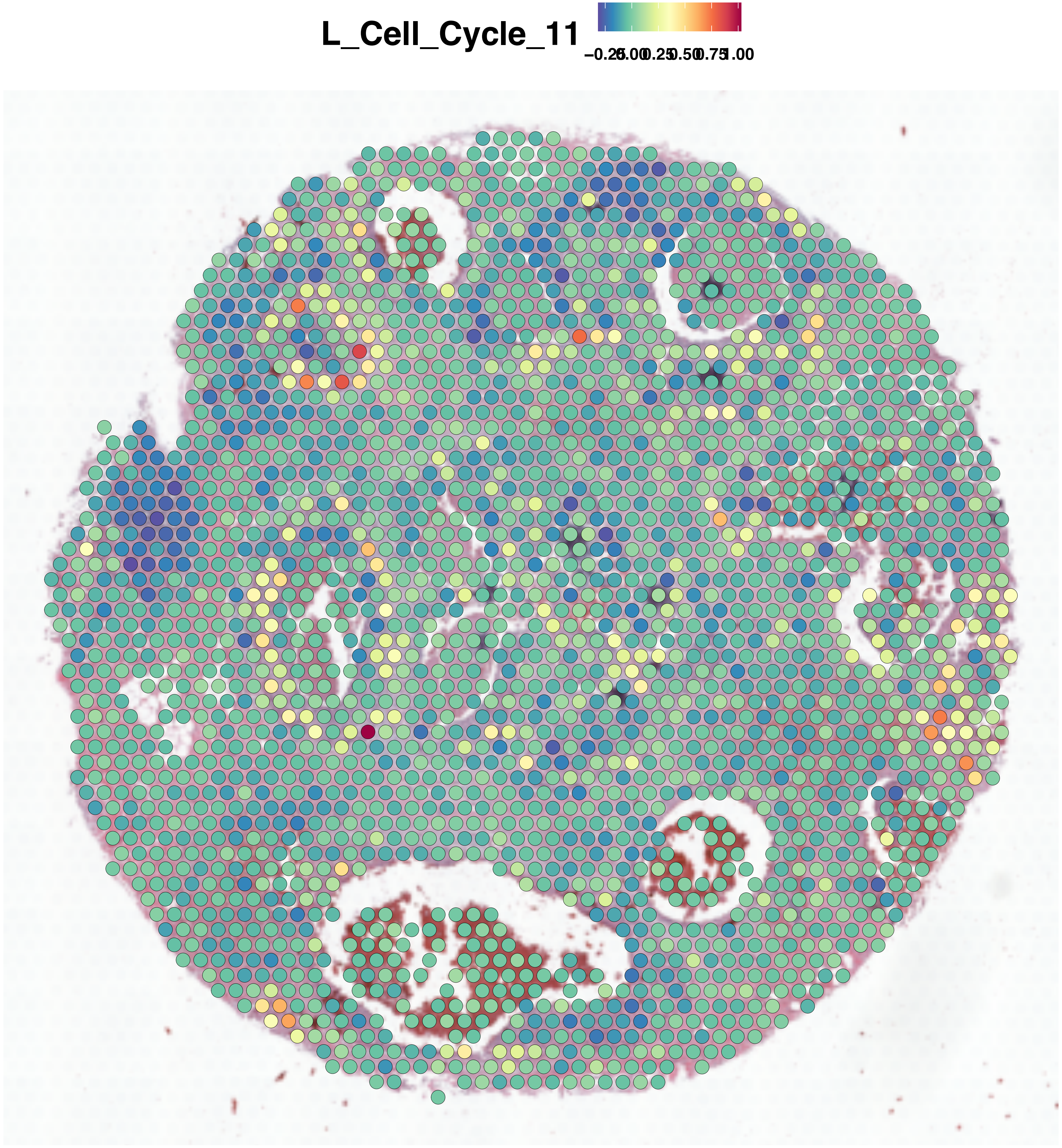
L_Cell_Cycle_11
−0.25
0.00
0.25
0.50
0.75
1.00

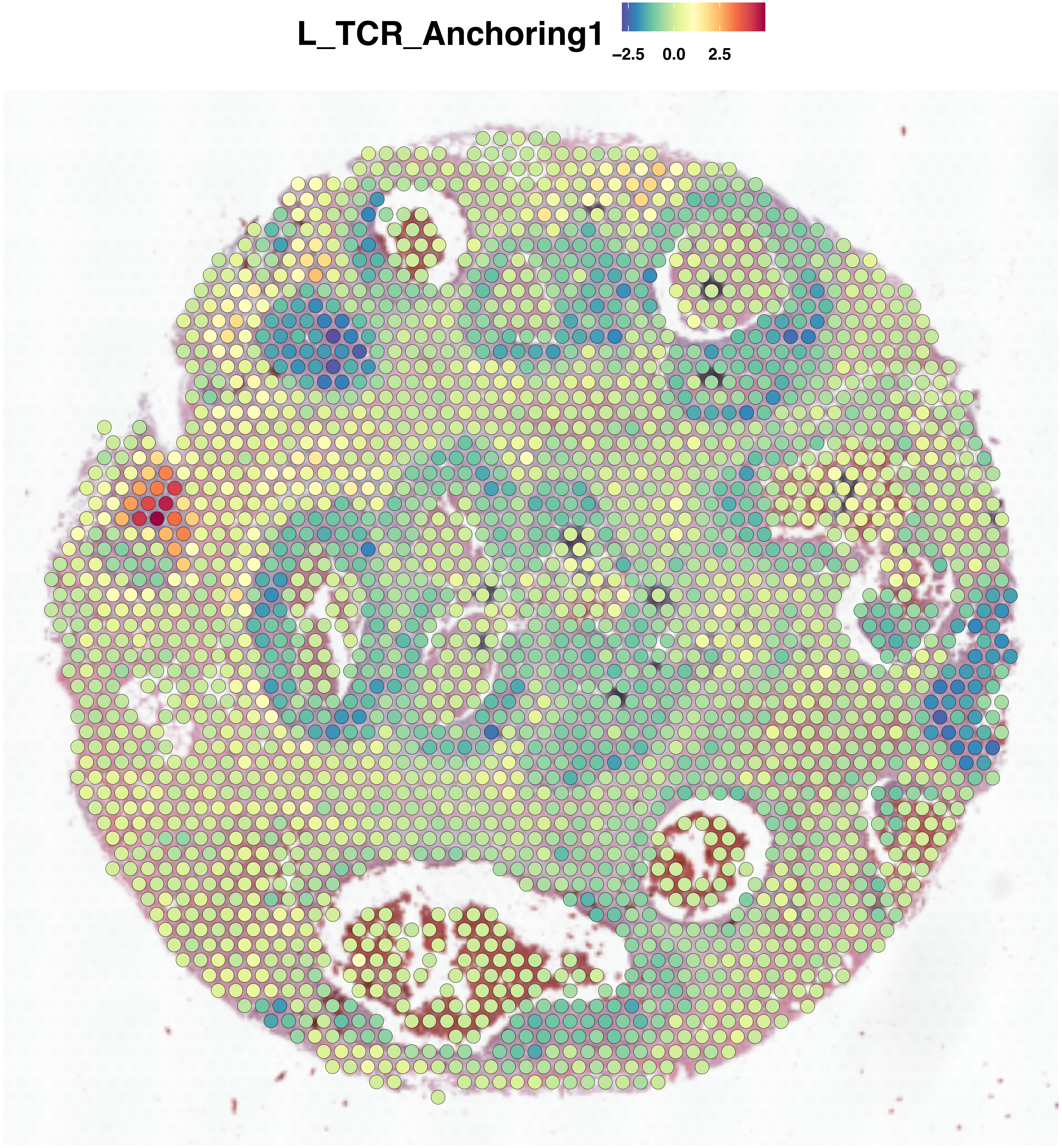
L_TCR_Anchoring1
−2.5
0.0
2.5

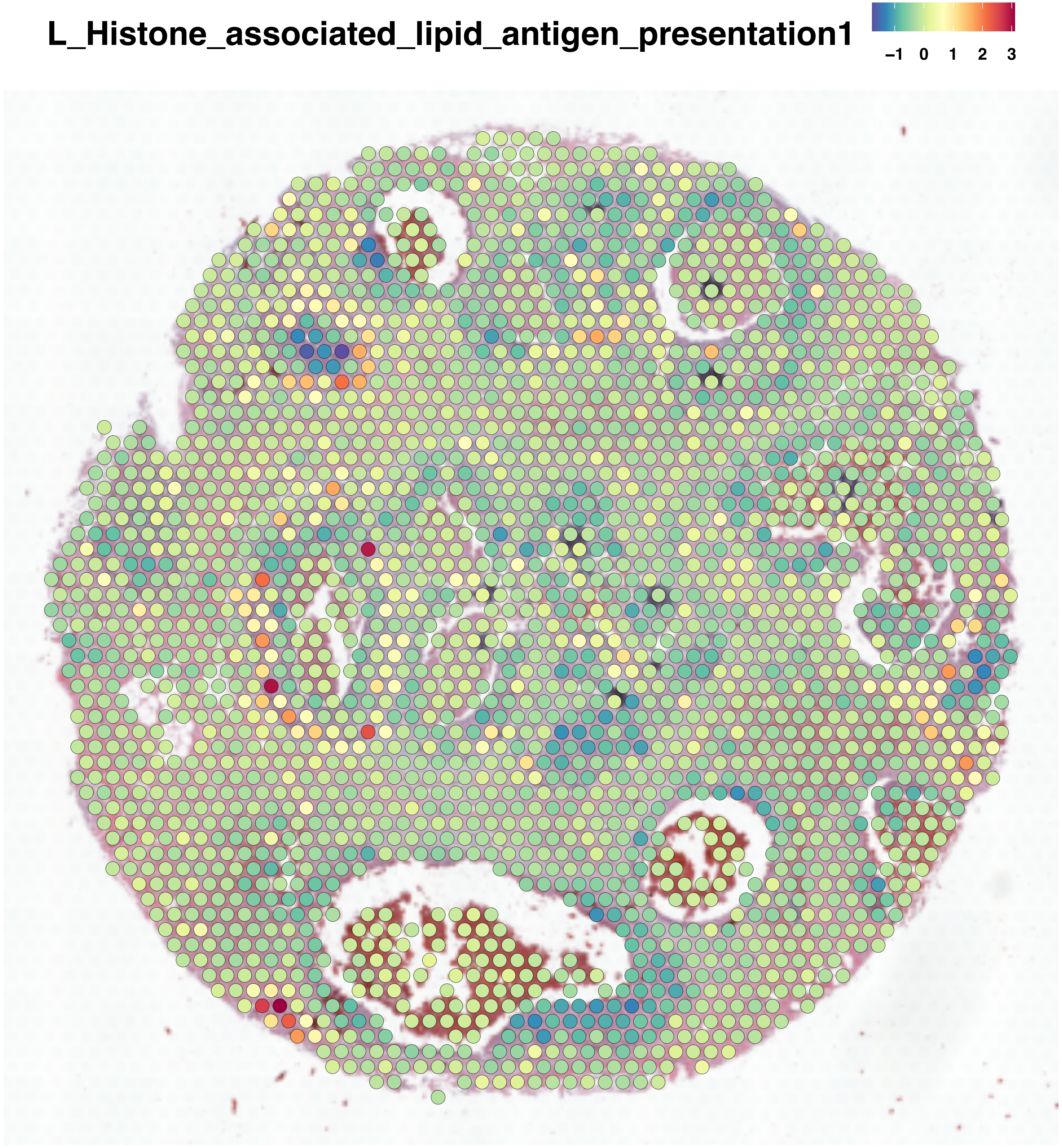
L_Histone_associated_lipid_antigen_presentation1
−1
0
1
2
3

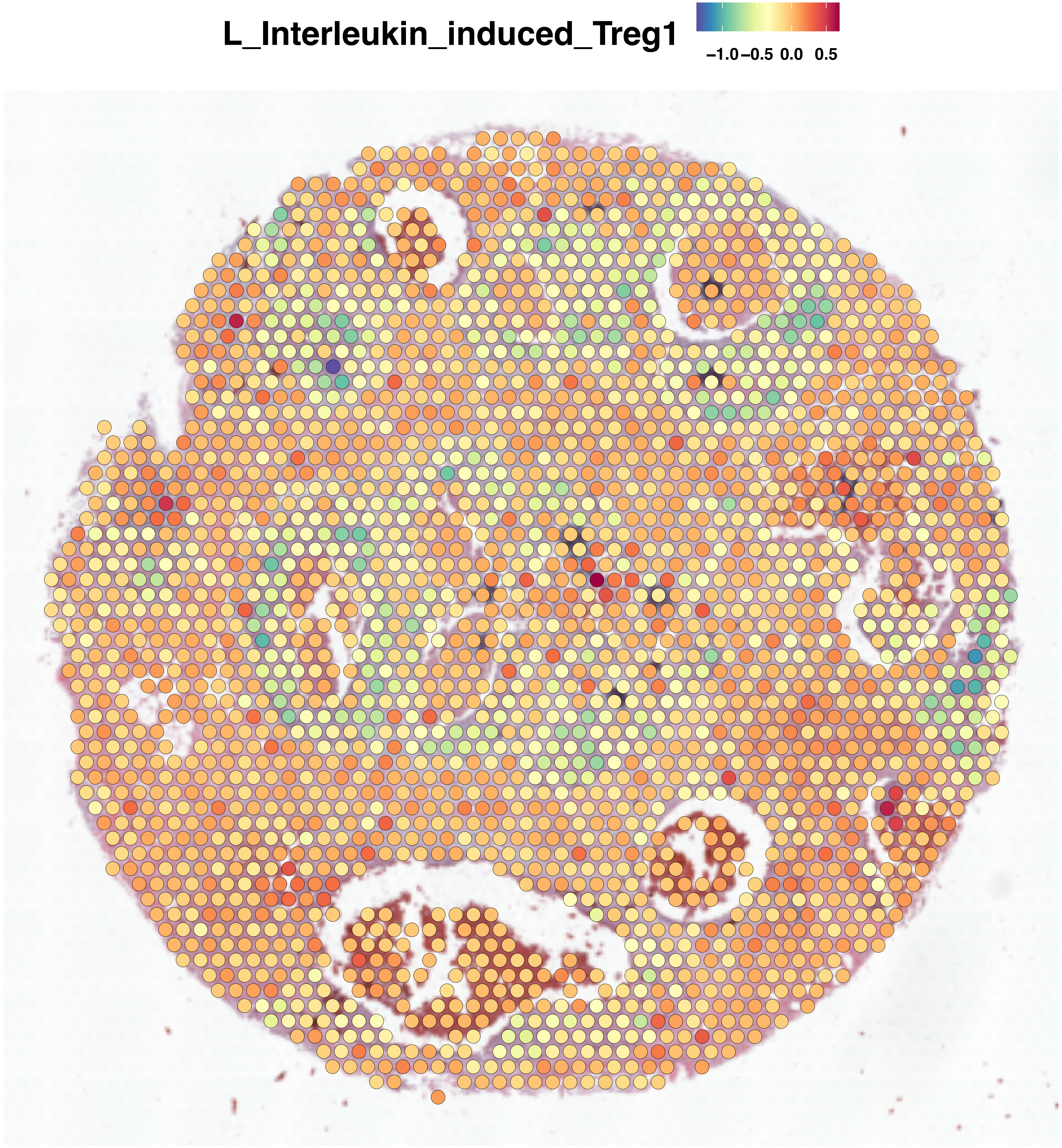
L_Interleukin_induced_Treg1
−1.0 −0.5 0.0 0.5

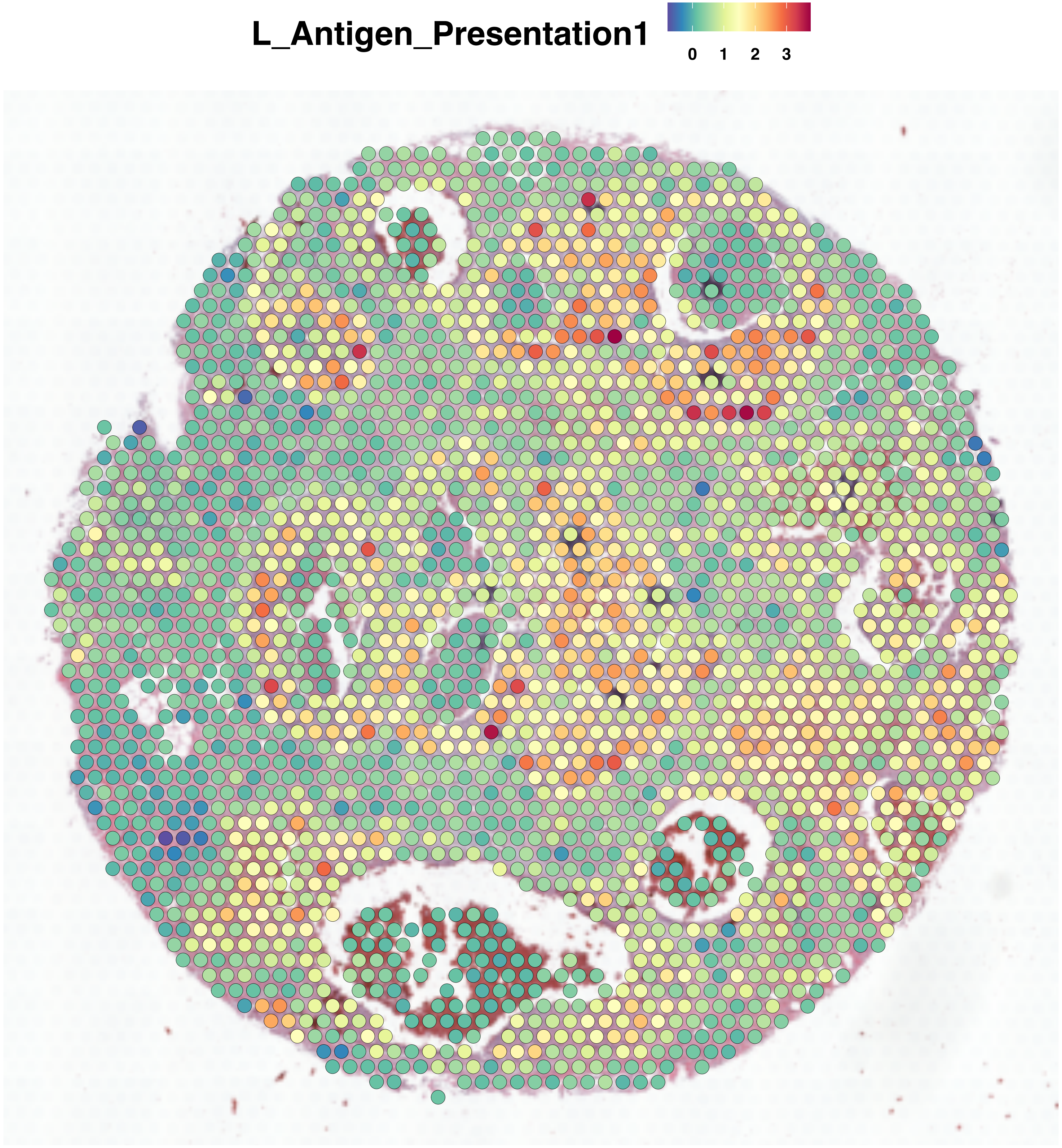
L_Antigen_Presentation1
0 1 2 3

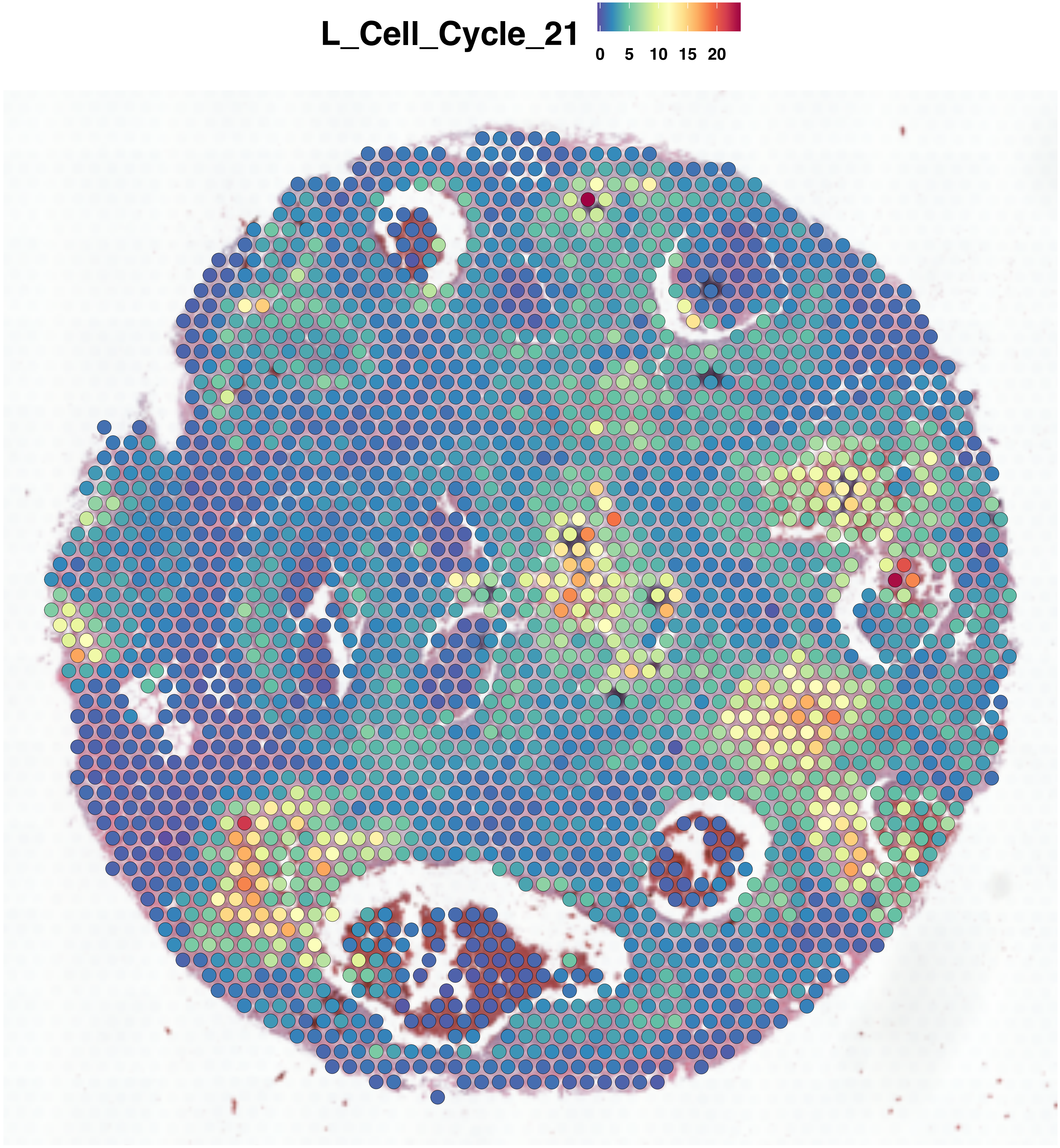
L_Cell_Cycle_21
0
5
10
15
20

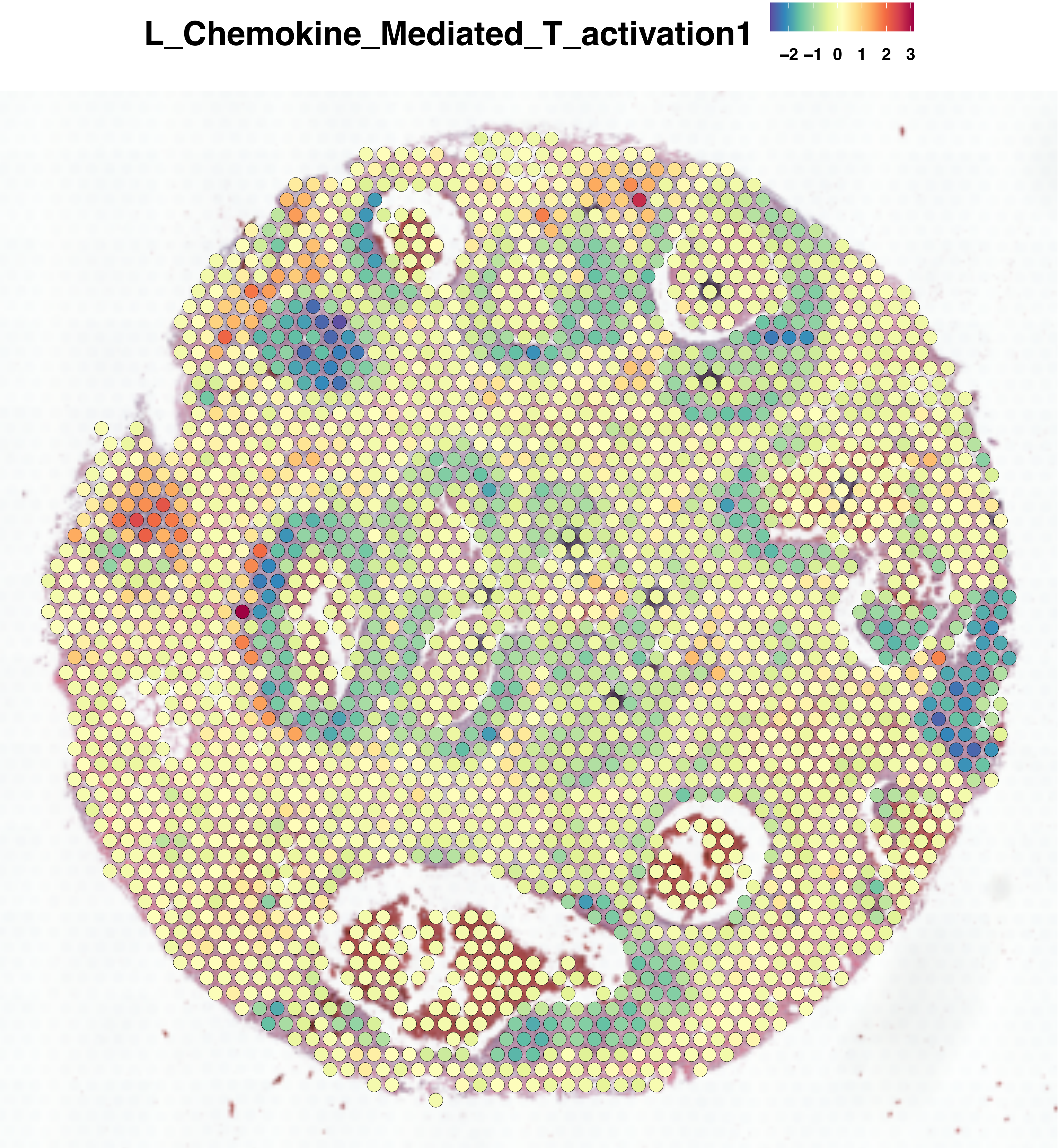
L_Chemokine_Mediated_T_activation1
−2 −1 0 1 2 3

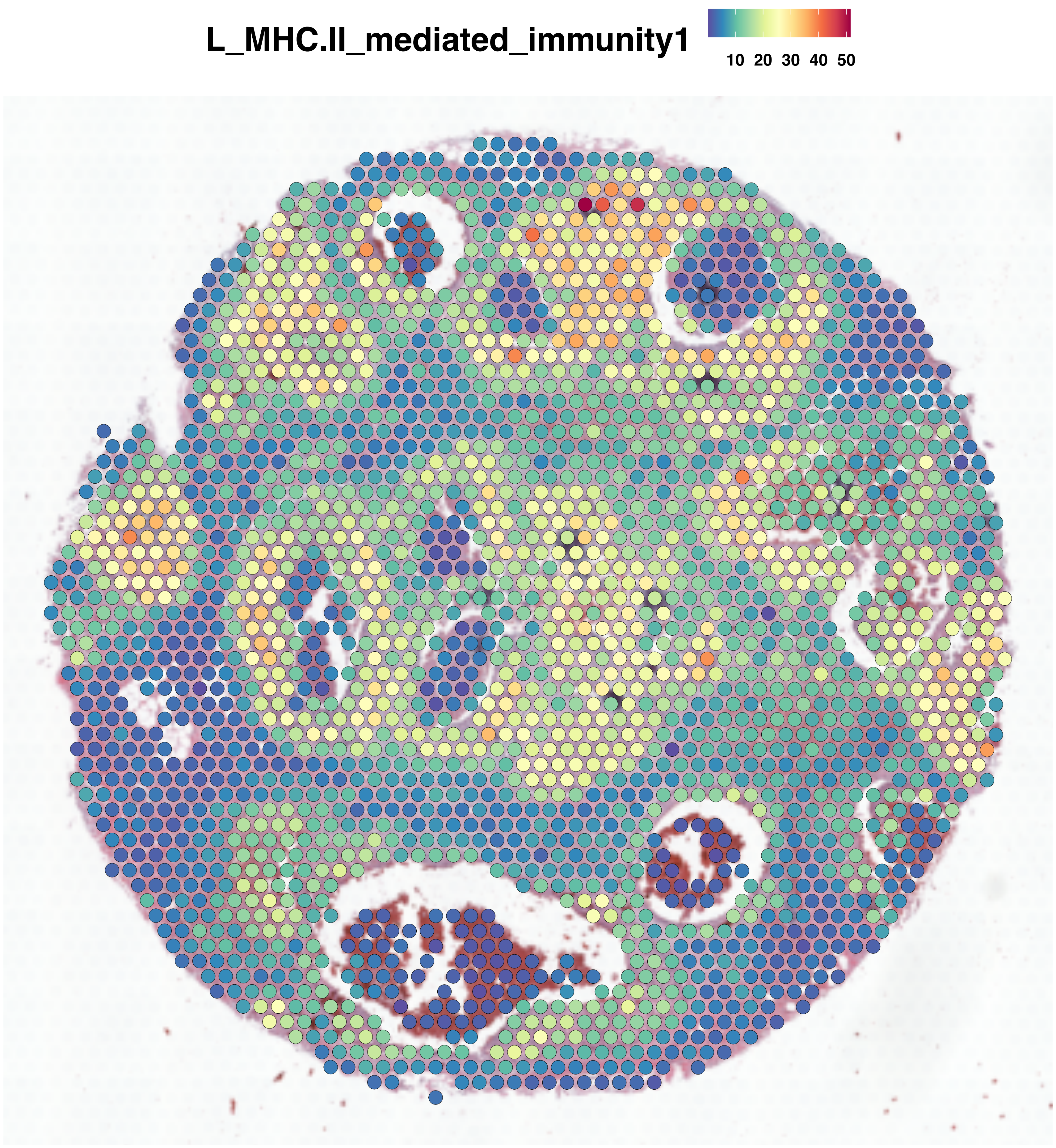
L_MHC.II_mediated_immunity1
10 20 30 40 50

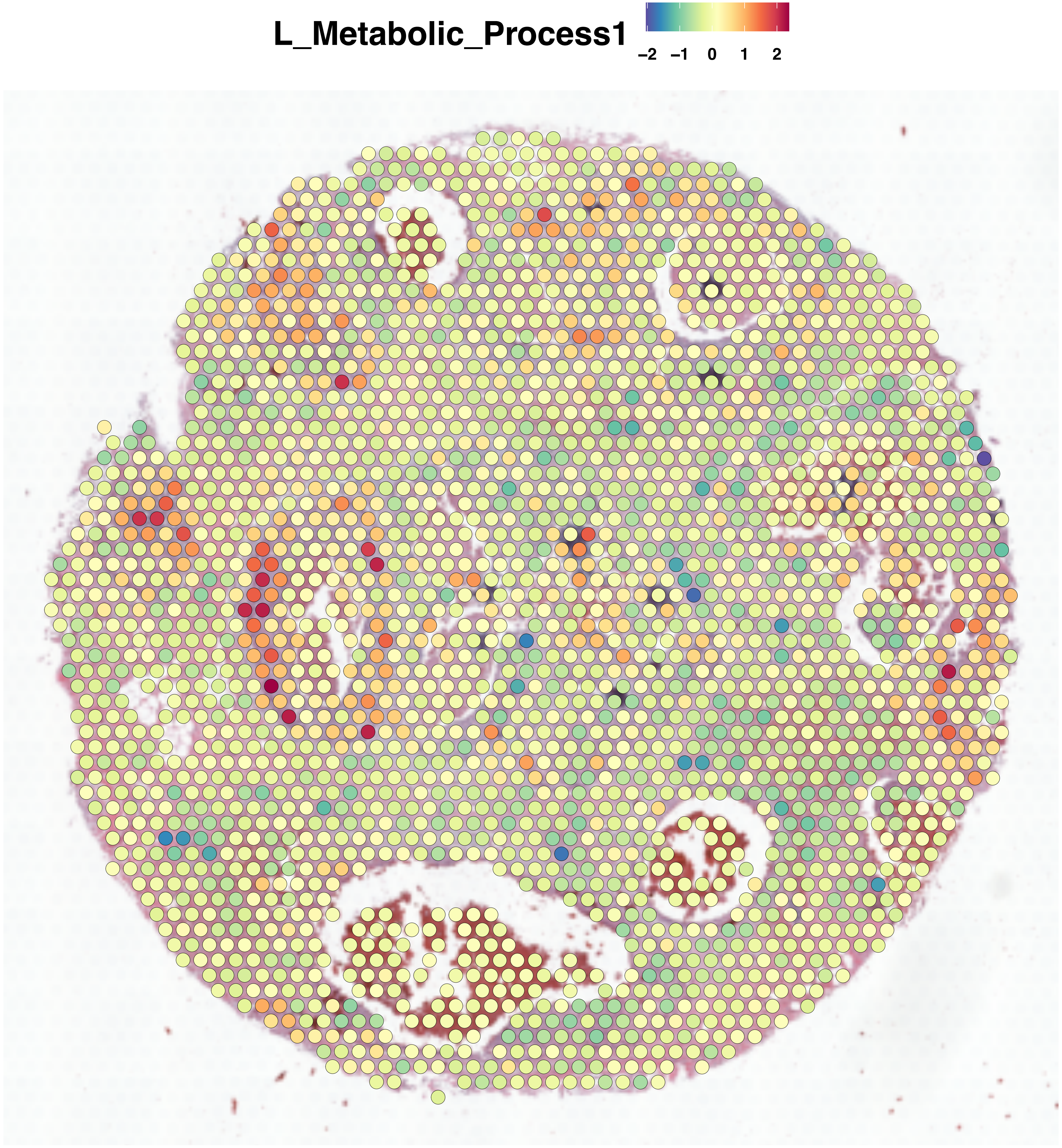
L_Metabolic_Process1
−2 −1 0 1 2

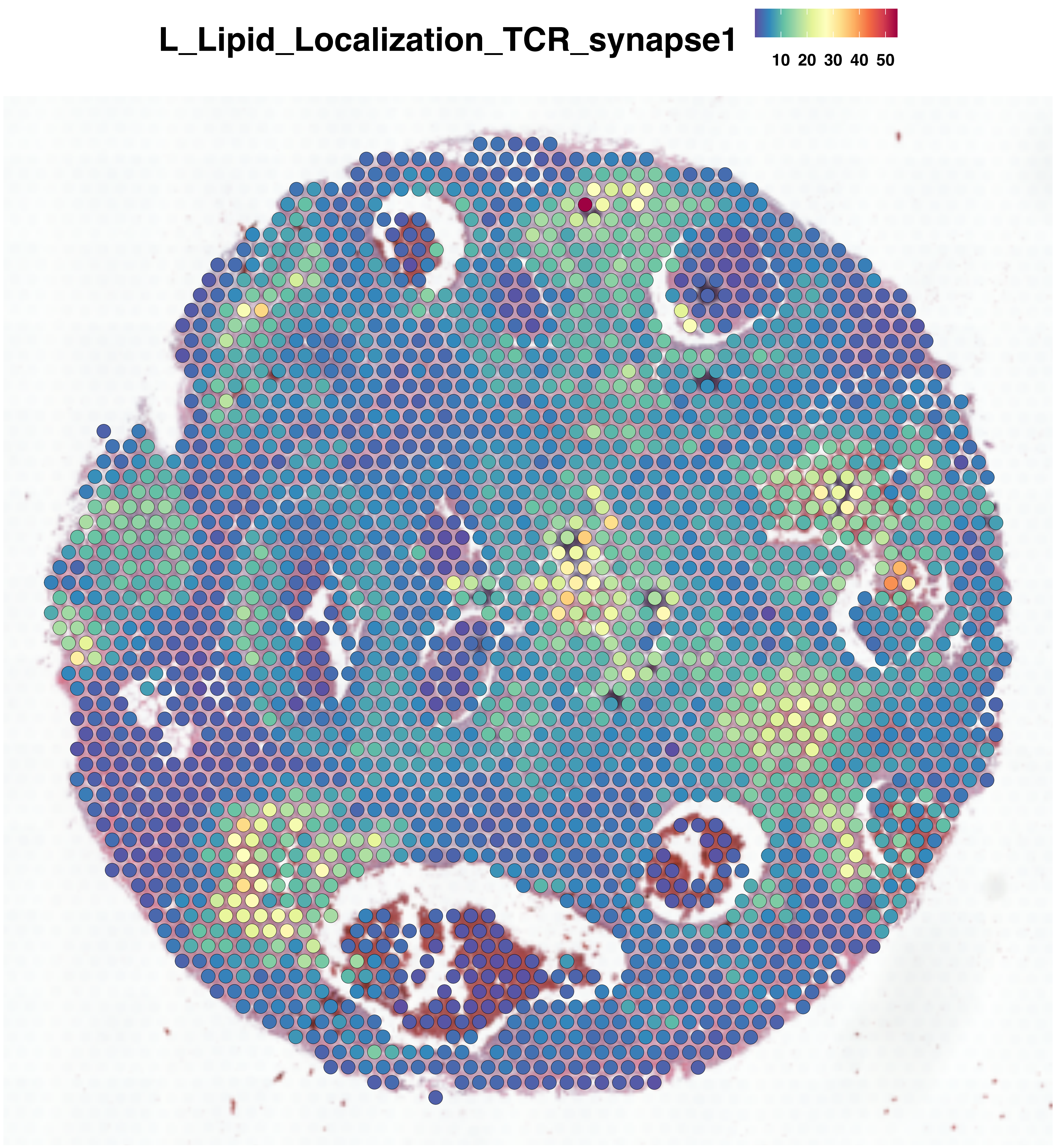
L_Lipid_Localization_TCR_synapse1
10 20 30 40 50

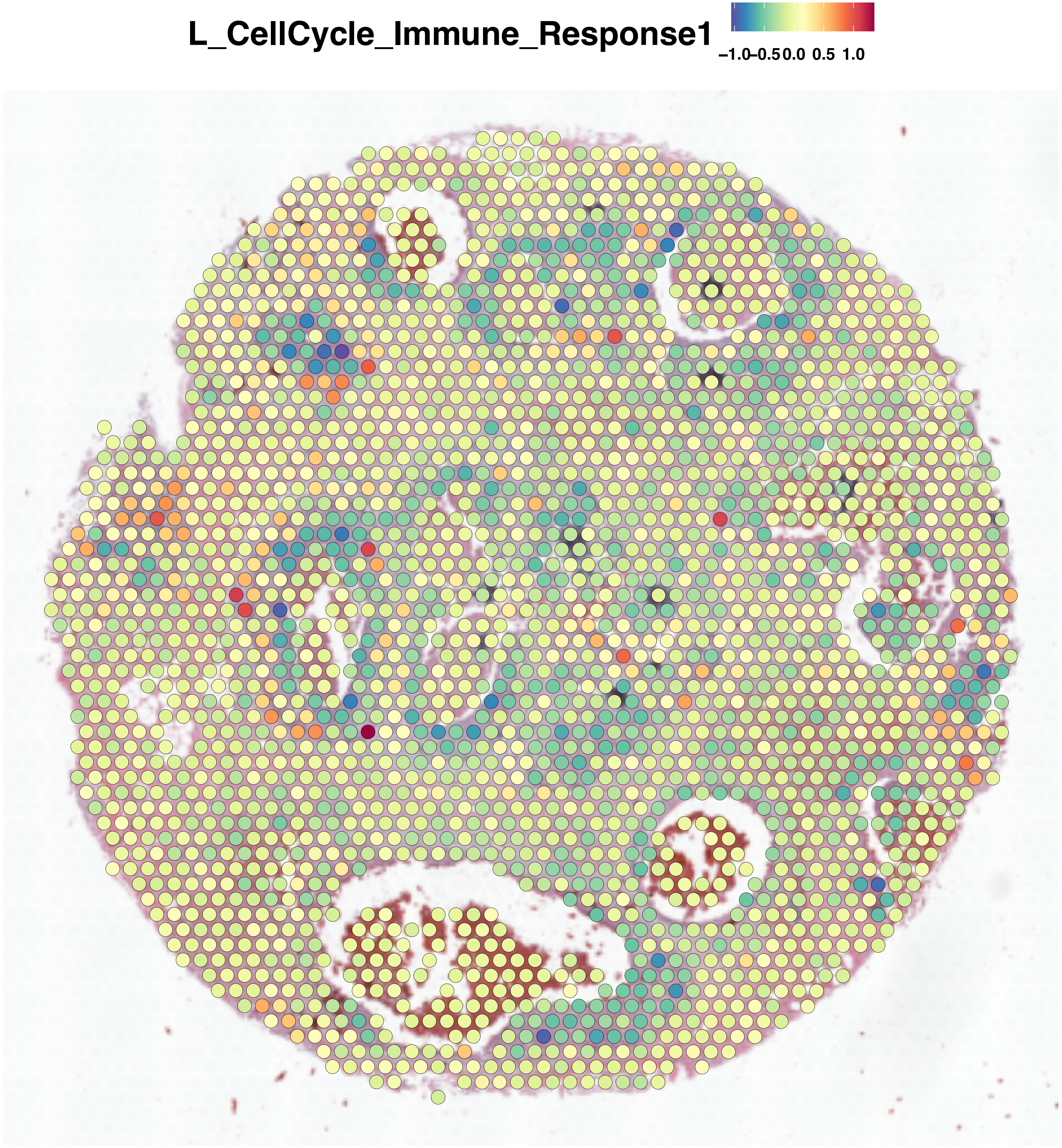
L_CellCycle_Immune_Response1
−1.0
−0.5
0.0
0.5
1.0

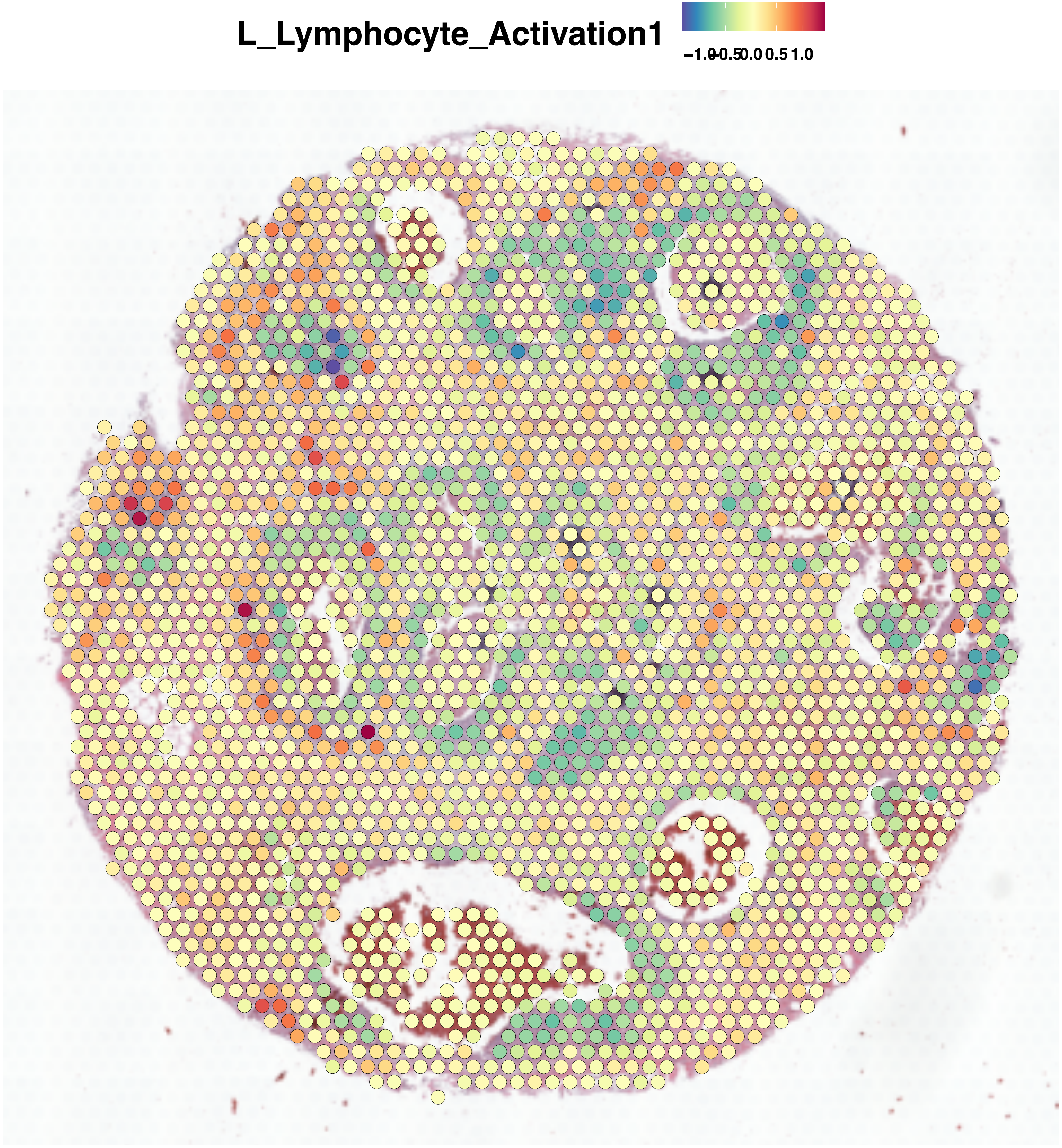
L_Lymphocyte_Activation1
−1.0
−0.5
0.0
0.5
1.0

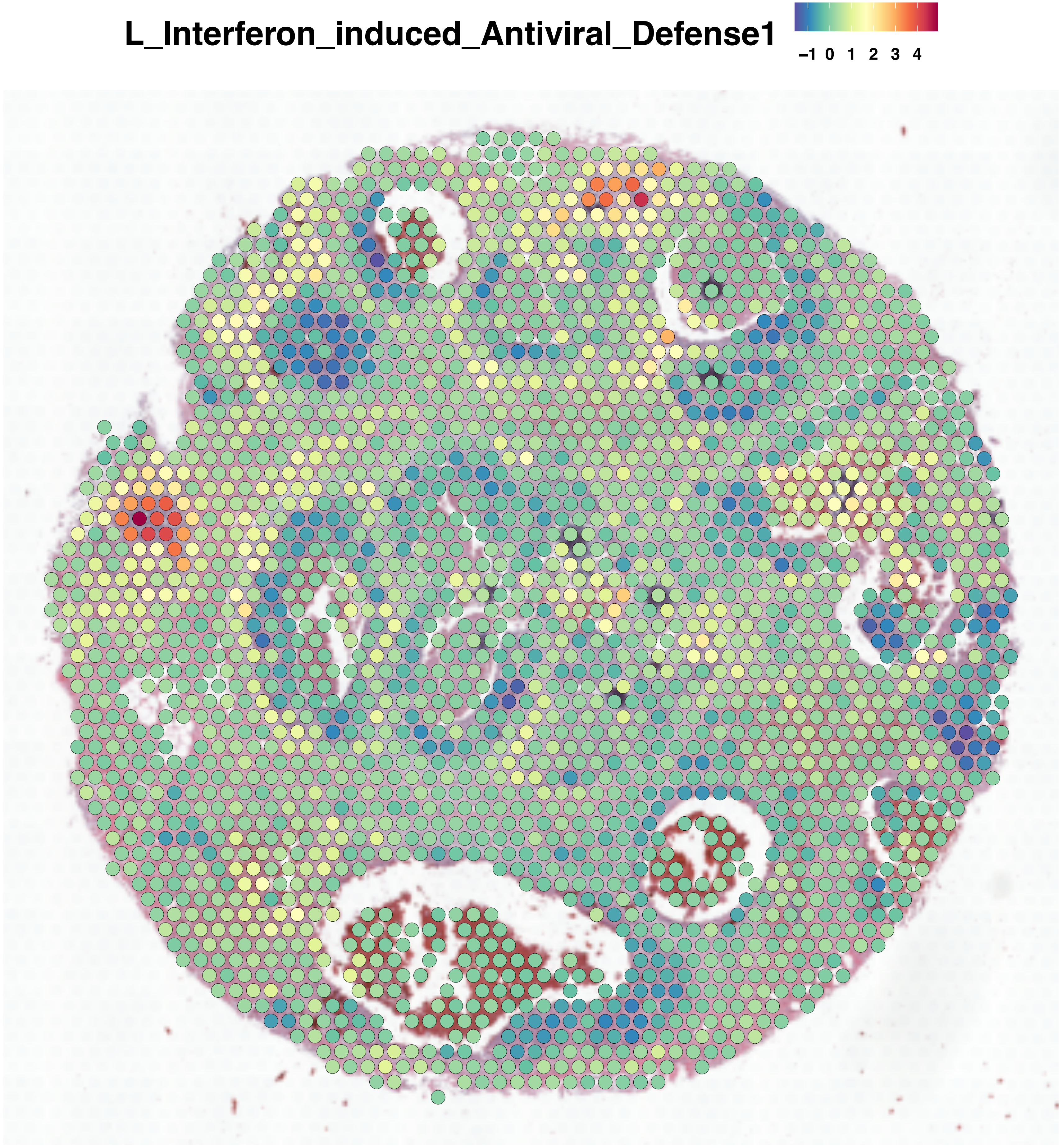

L_Interferon_induced_Antiviral_Defense1
−1 0 1 2 3 4

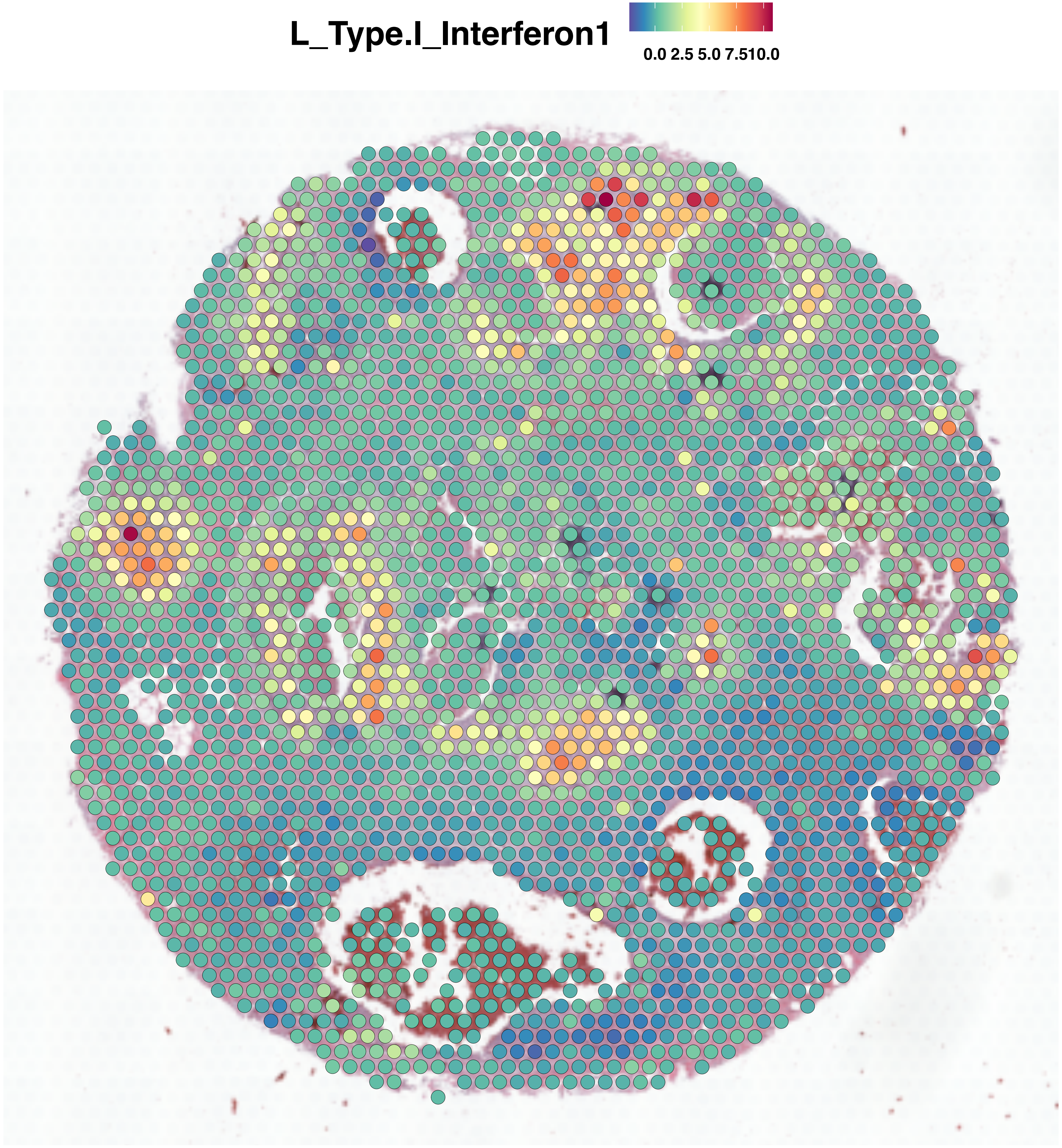
L_Type.I_Interferon1
0.0 2.5 5.0 7.510.0

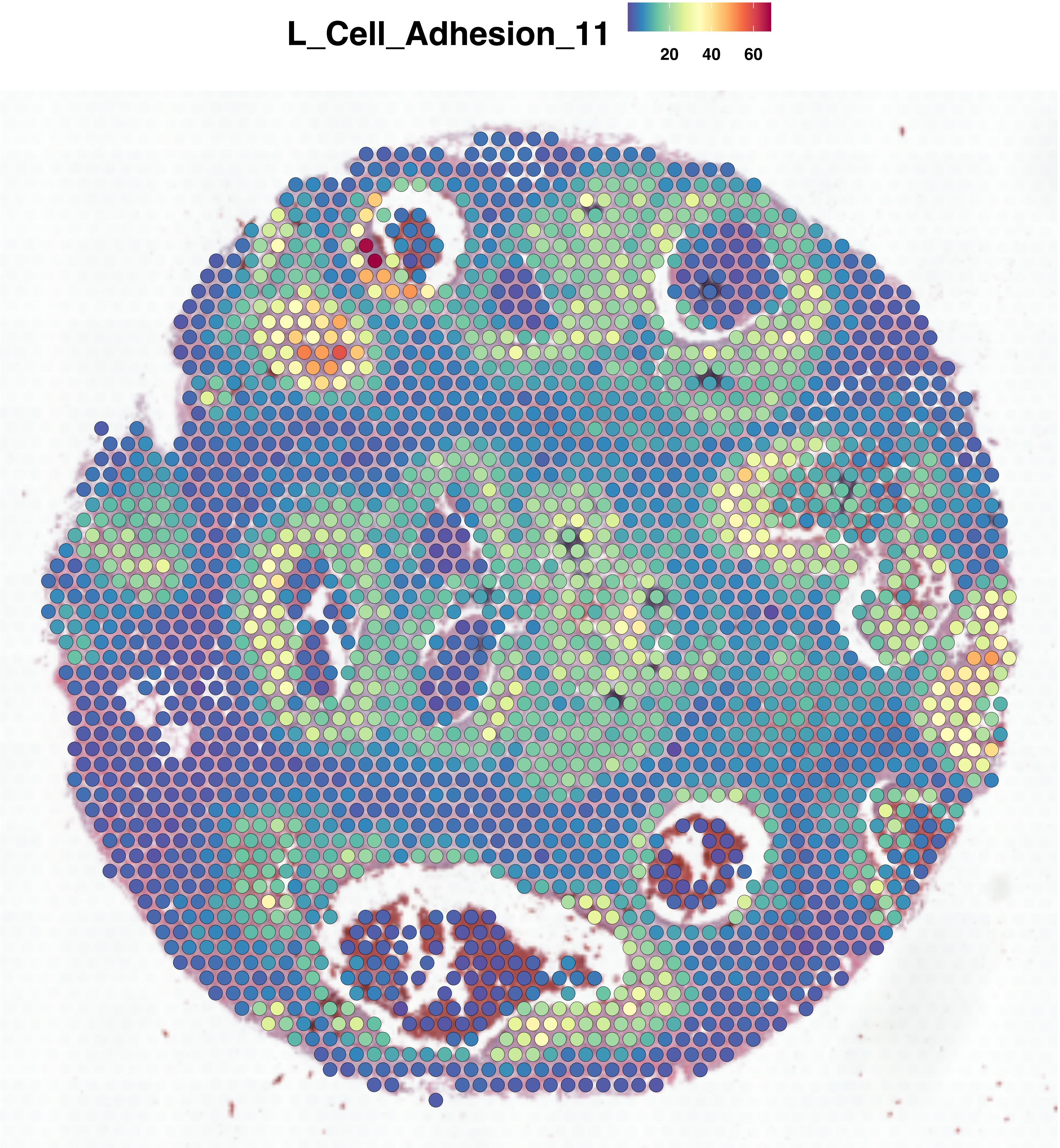
L_Cell_Adhesion_11
20
40
60

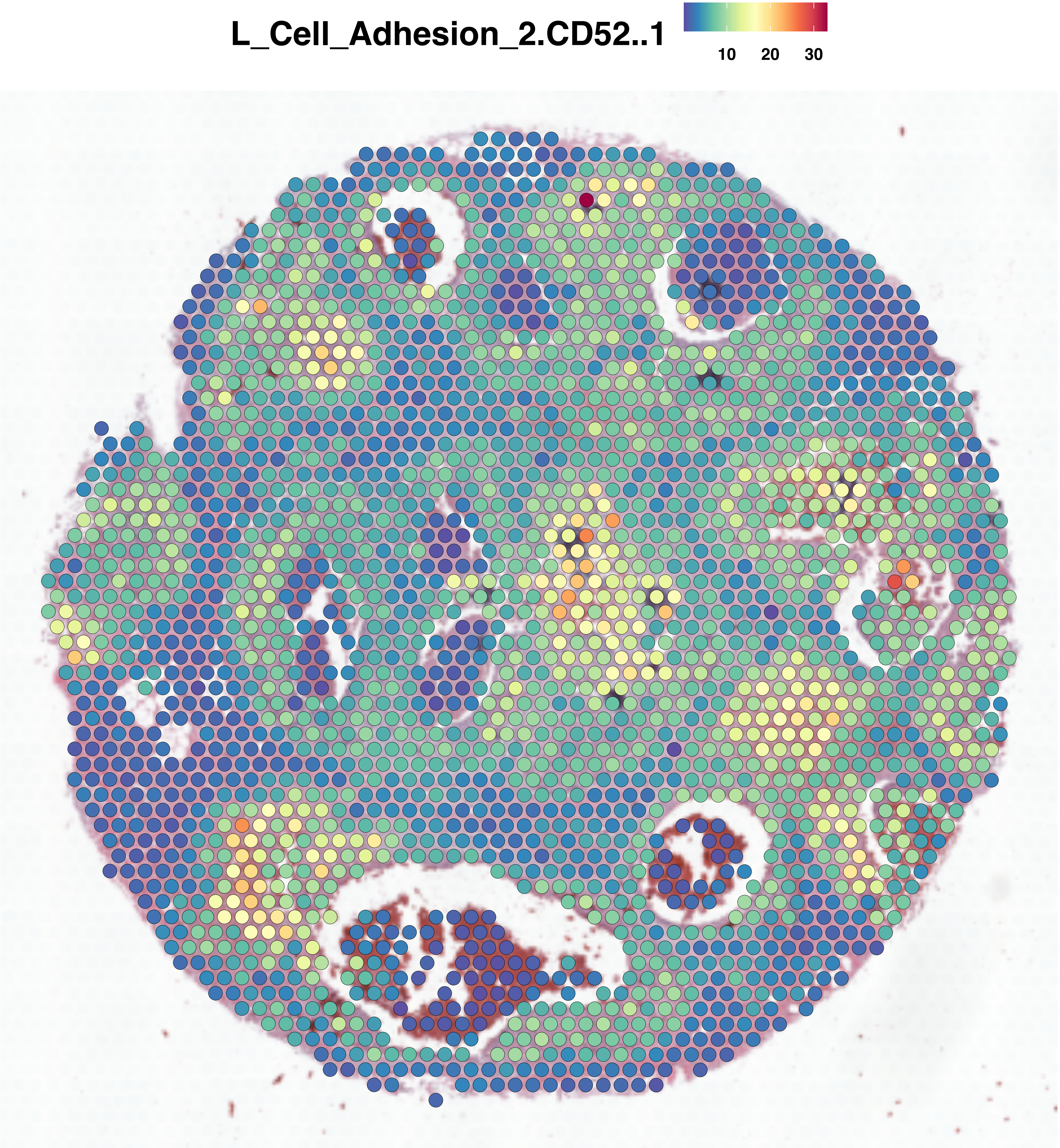
L_Cell_Adhesion_2.CD52..1
10
20
30

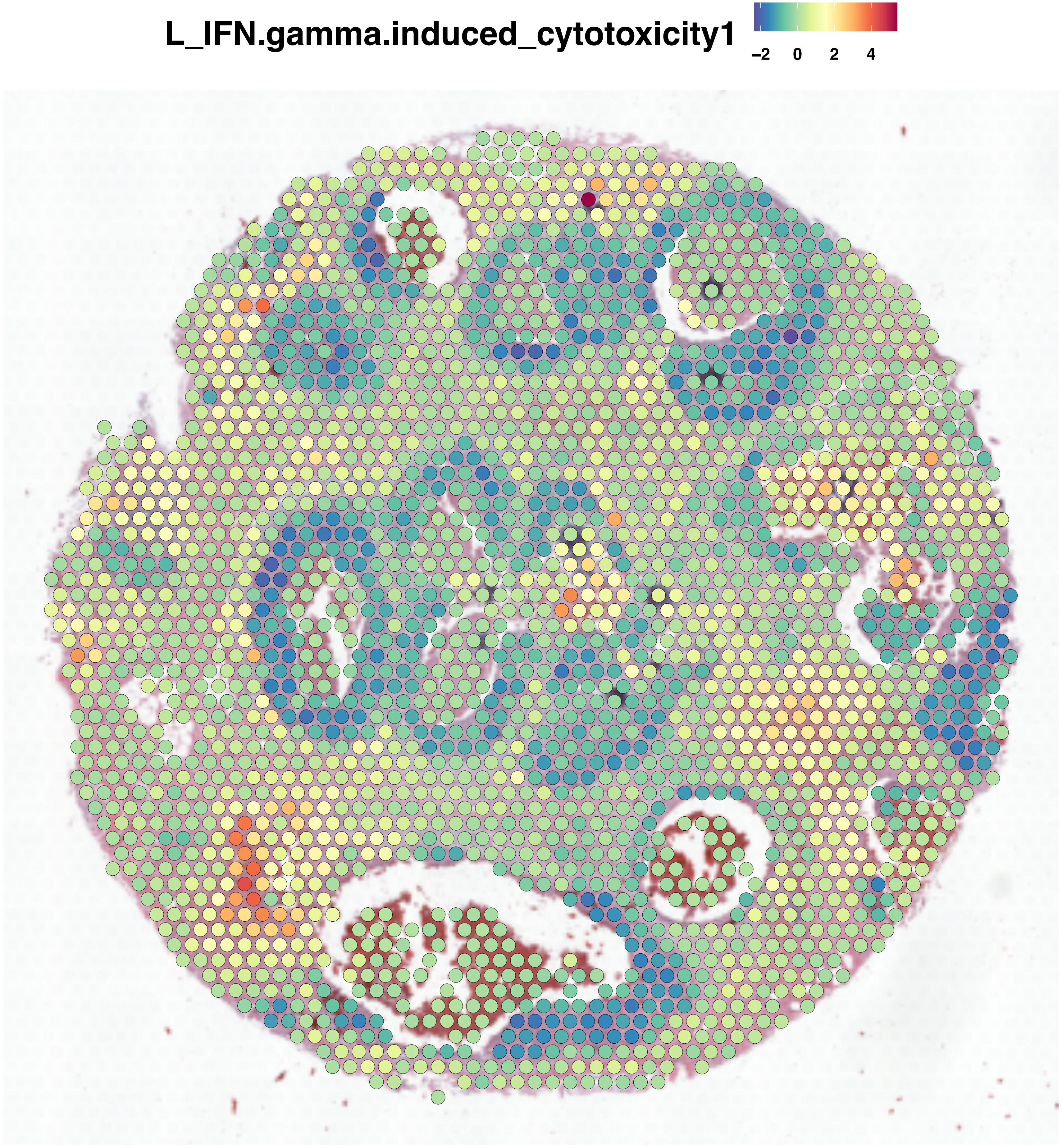
L_IFN.gamma.induced_cytotoxicity1
−2
0
2
4

L_Cytokine_Receptor_Signaling1

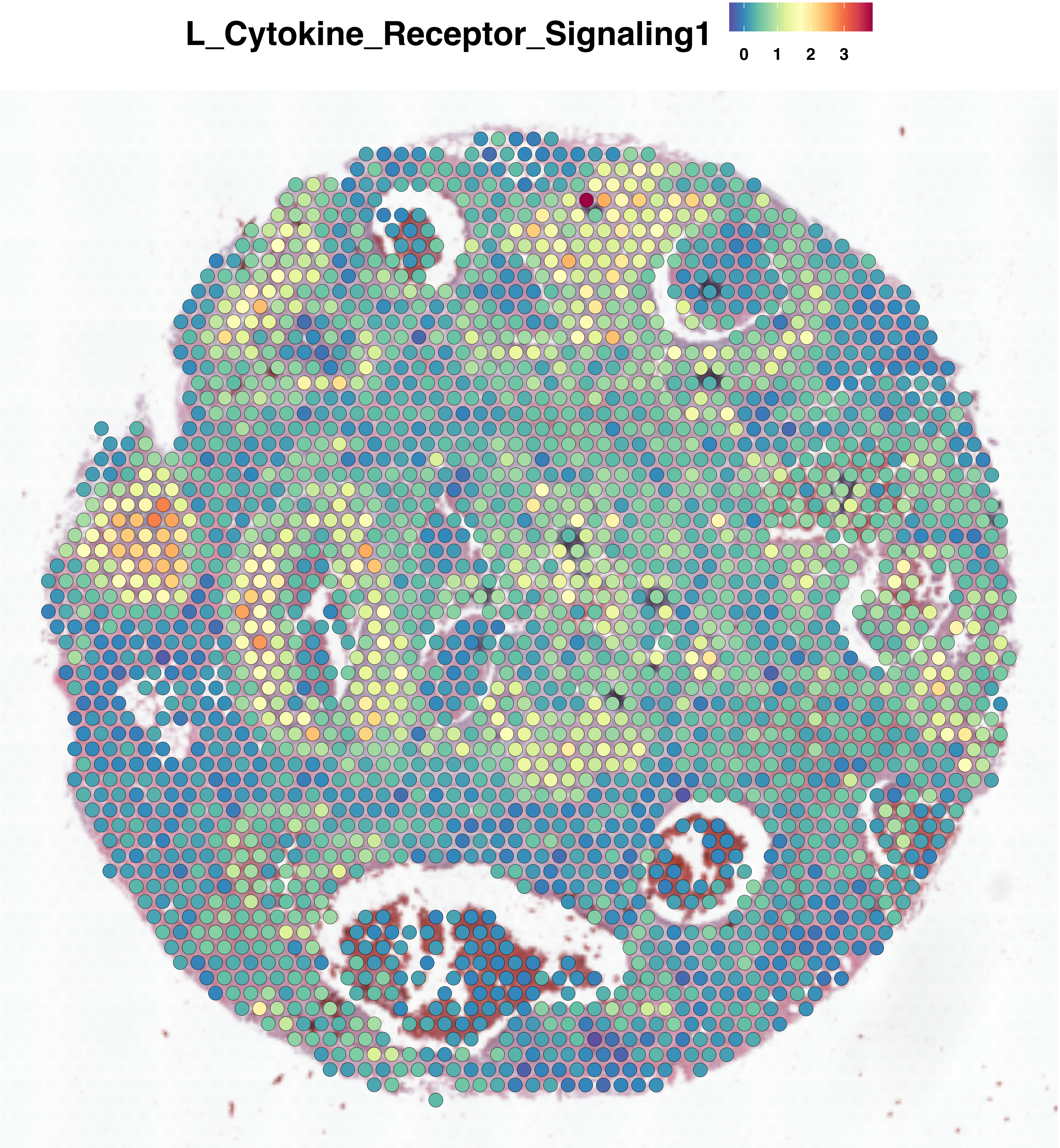

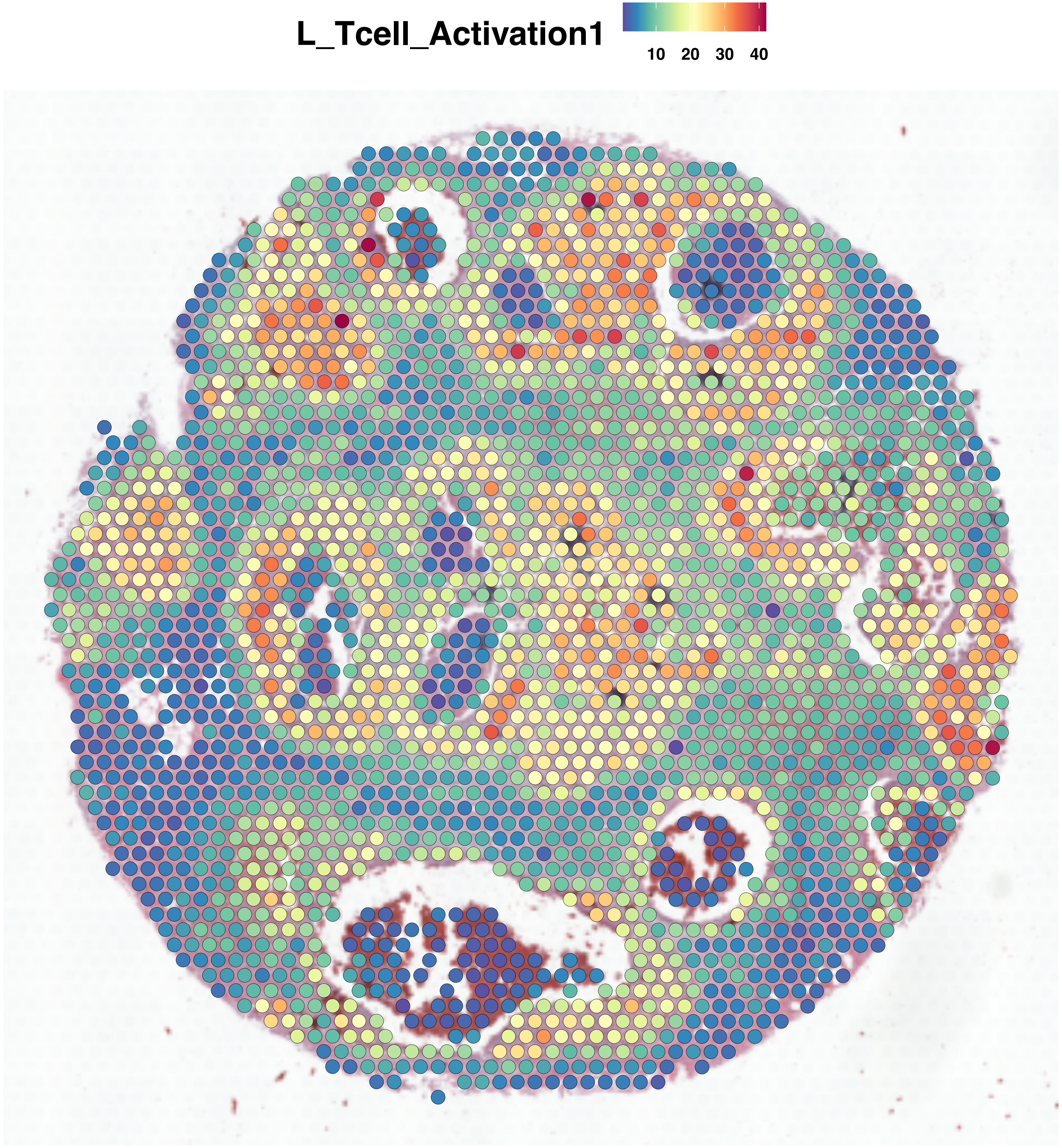
L_Tcell_Activation1
10 20 30 40

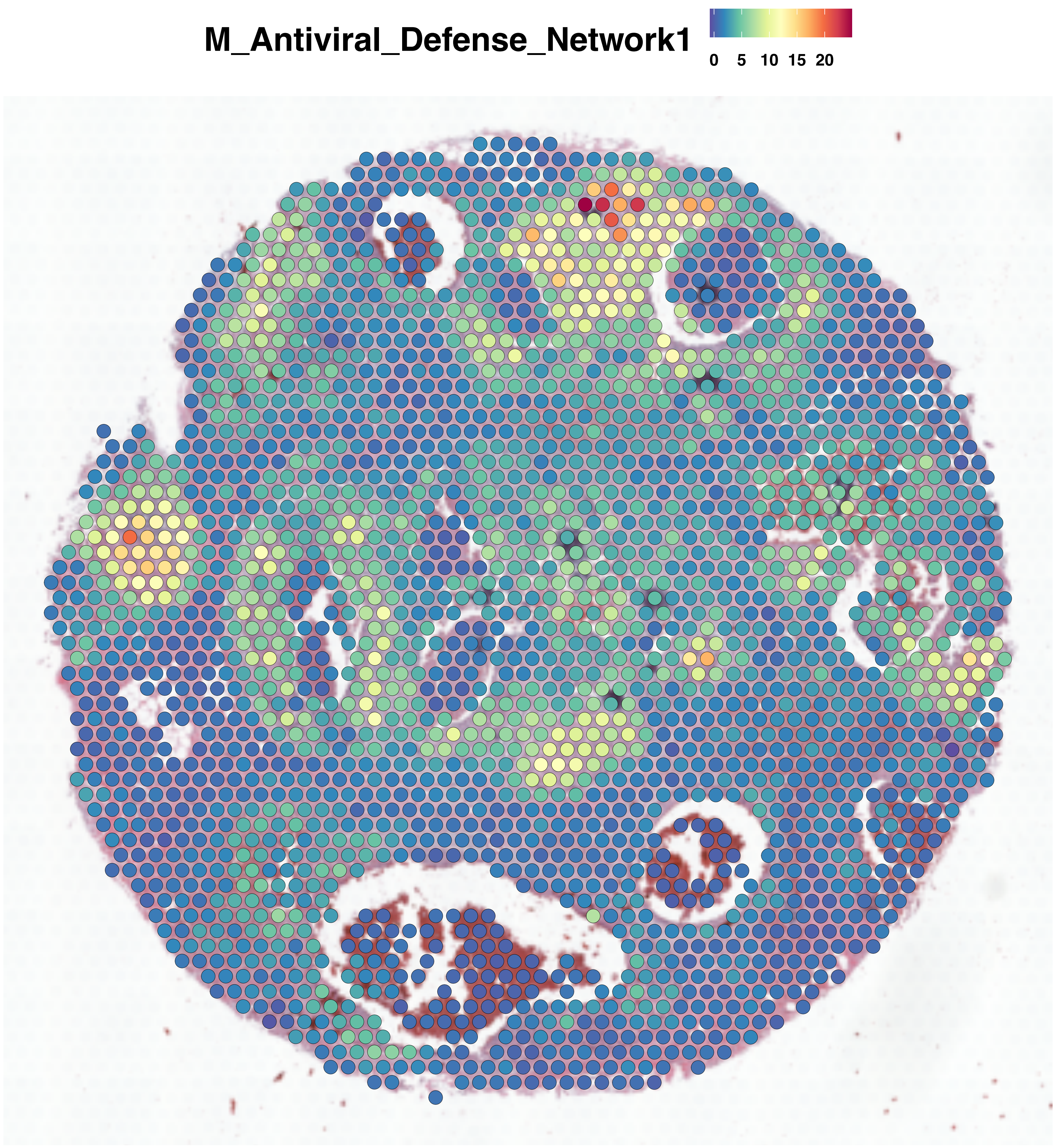
M_Antiviral_Defense_Network1
0
5
10
15
20

**M_Antigen_Processing1**

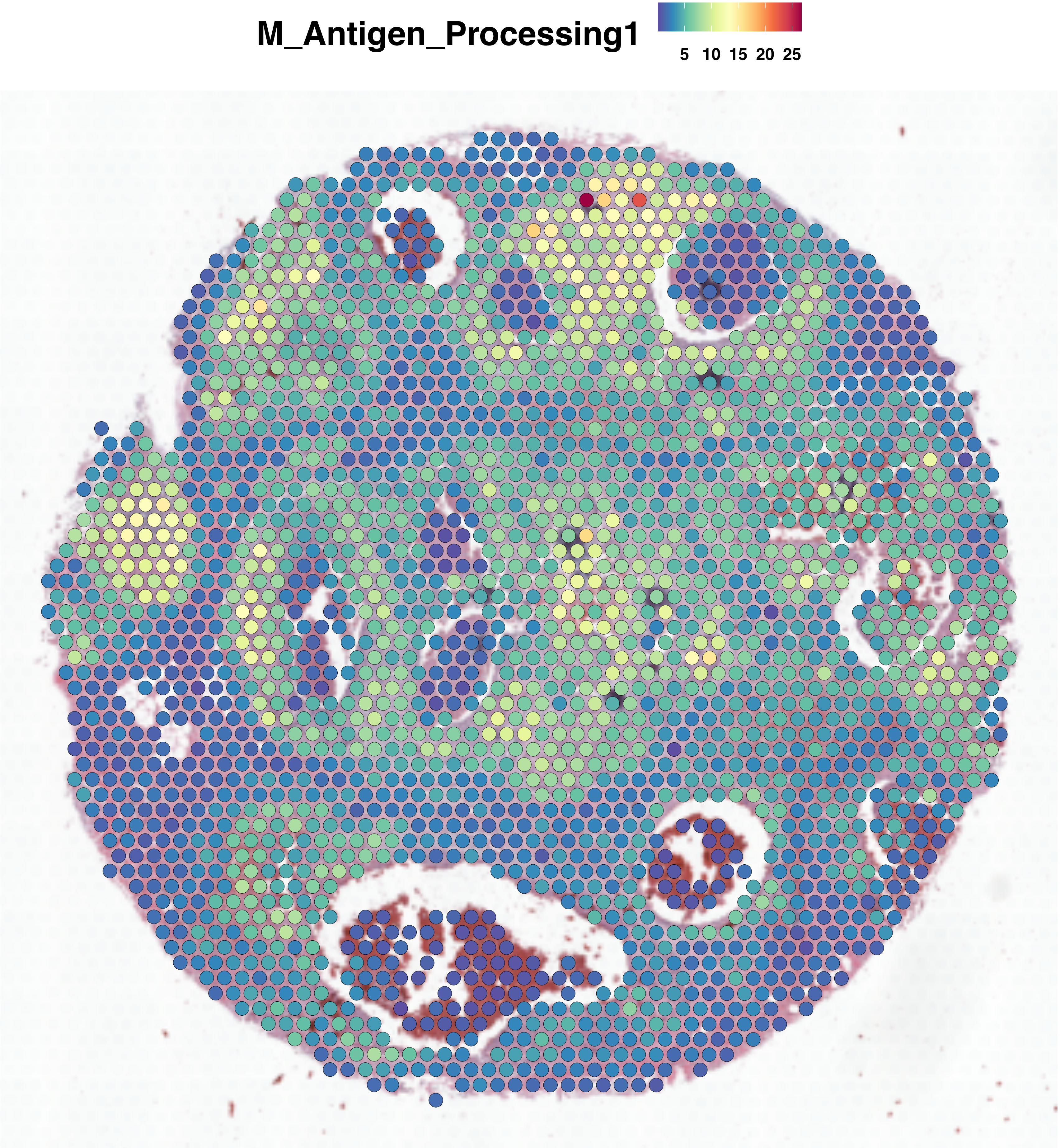

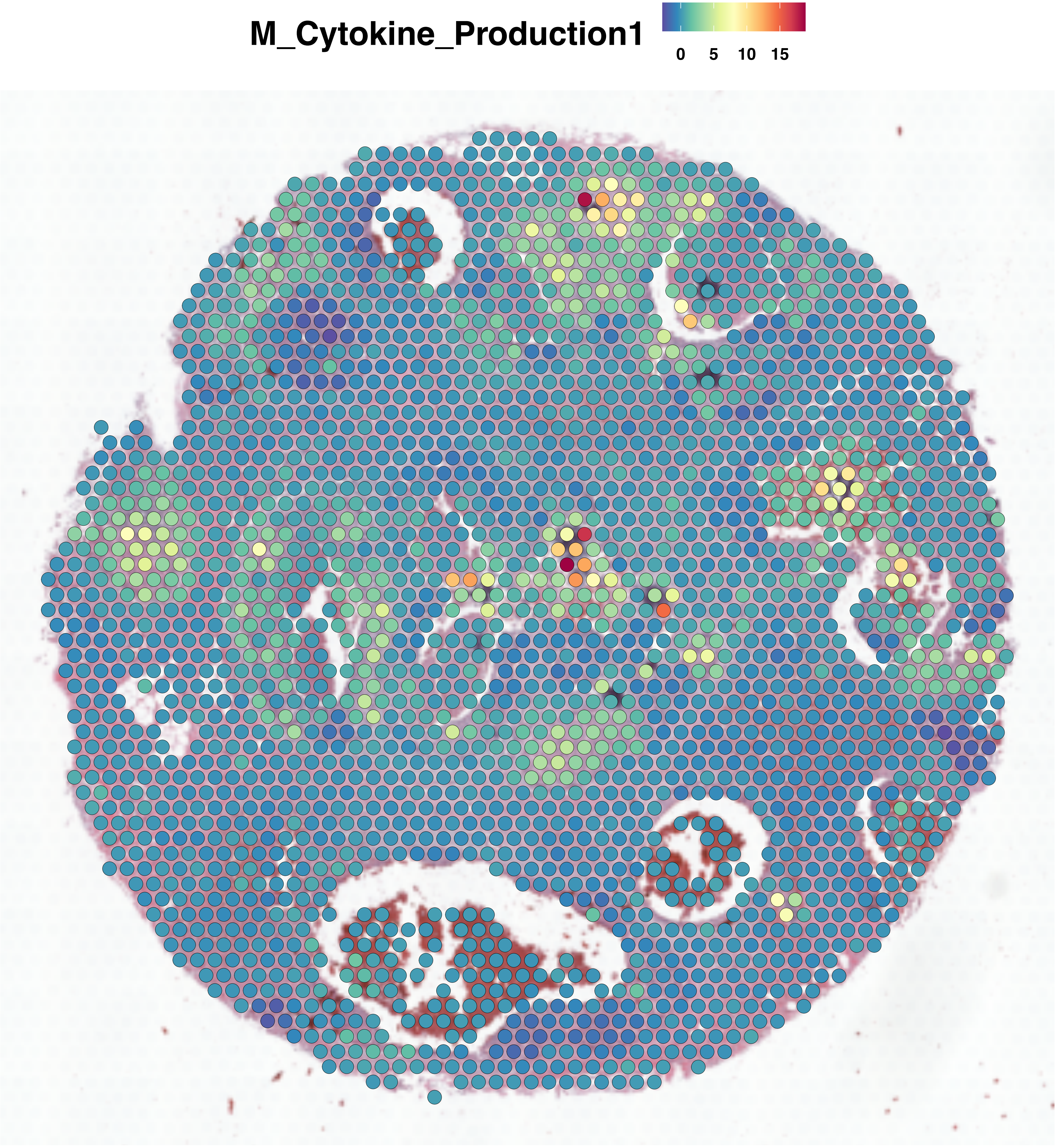

M_Cytokine_Production1
0
5
10
15

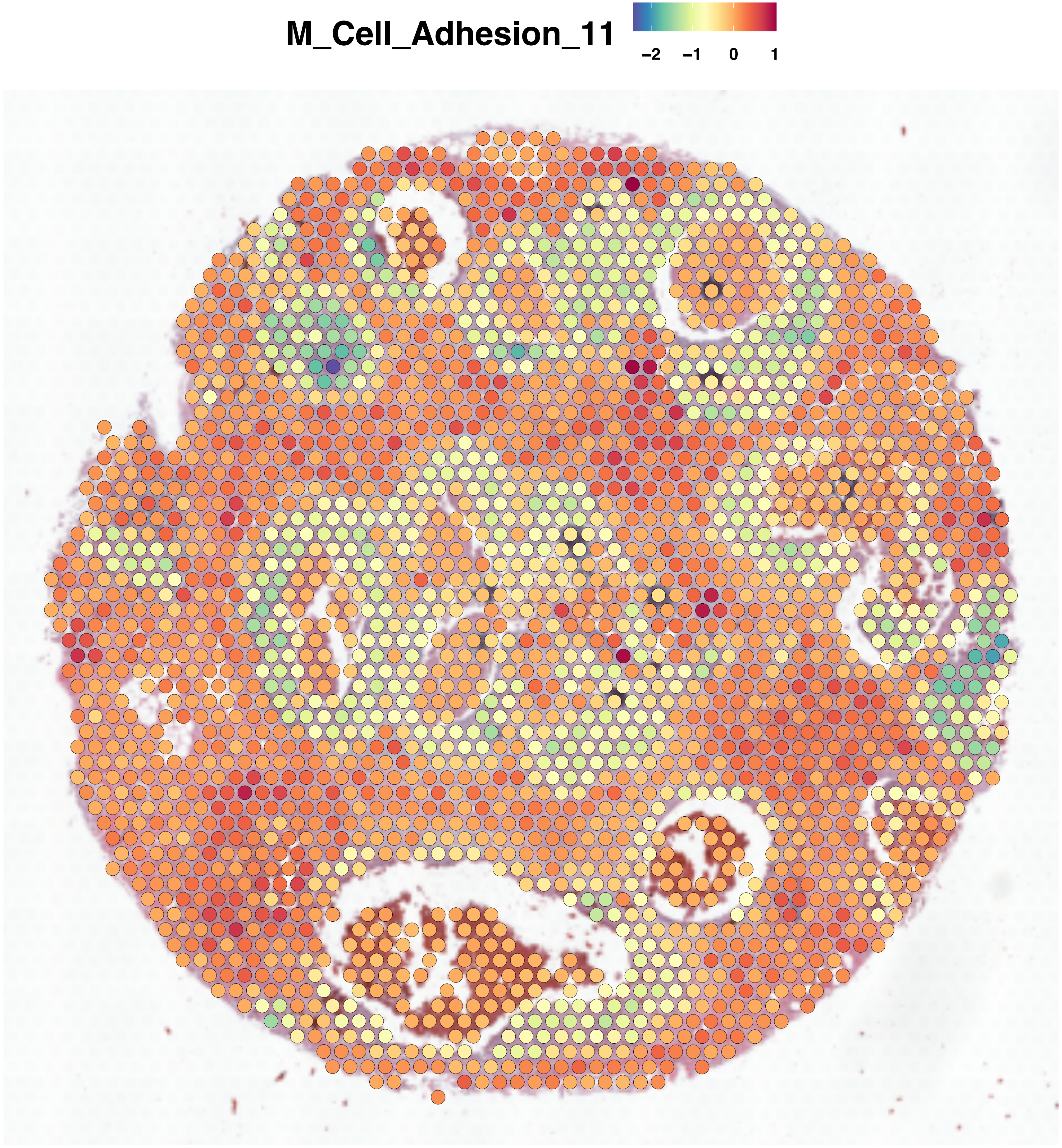
M_Cell_Adhesion_11
−2
−1
0
1

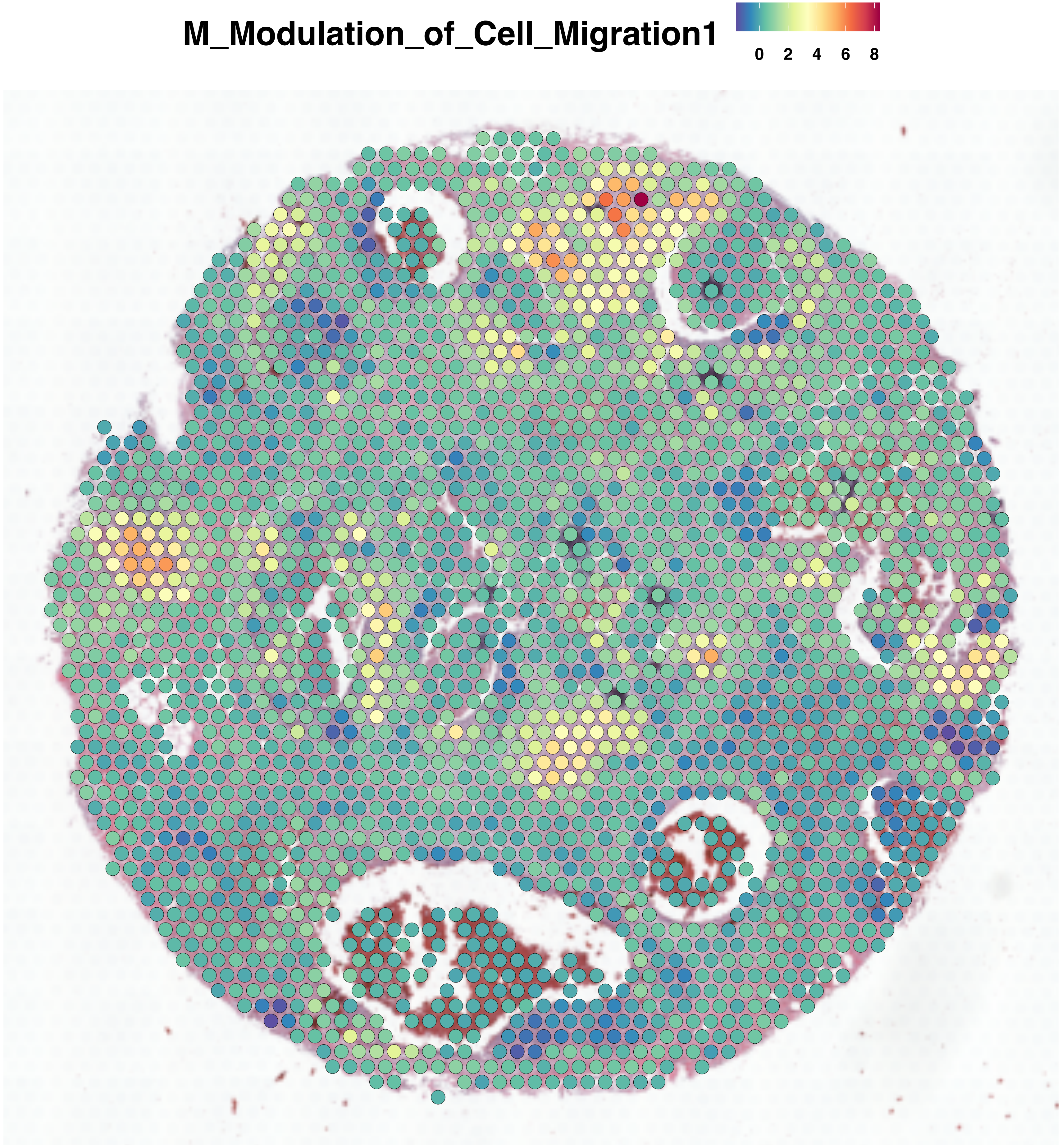
M_Modulation_of_Cell_Migration1
0 2 4 6 8

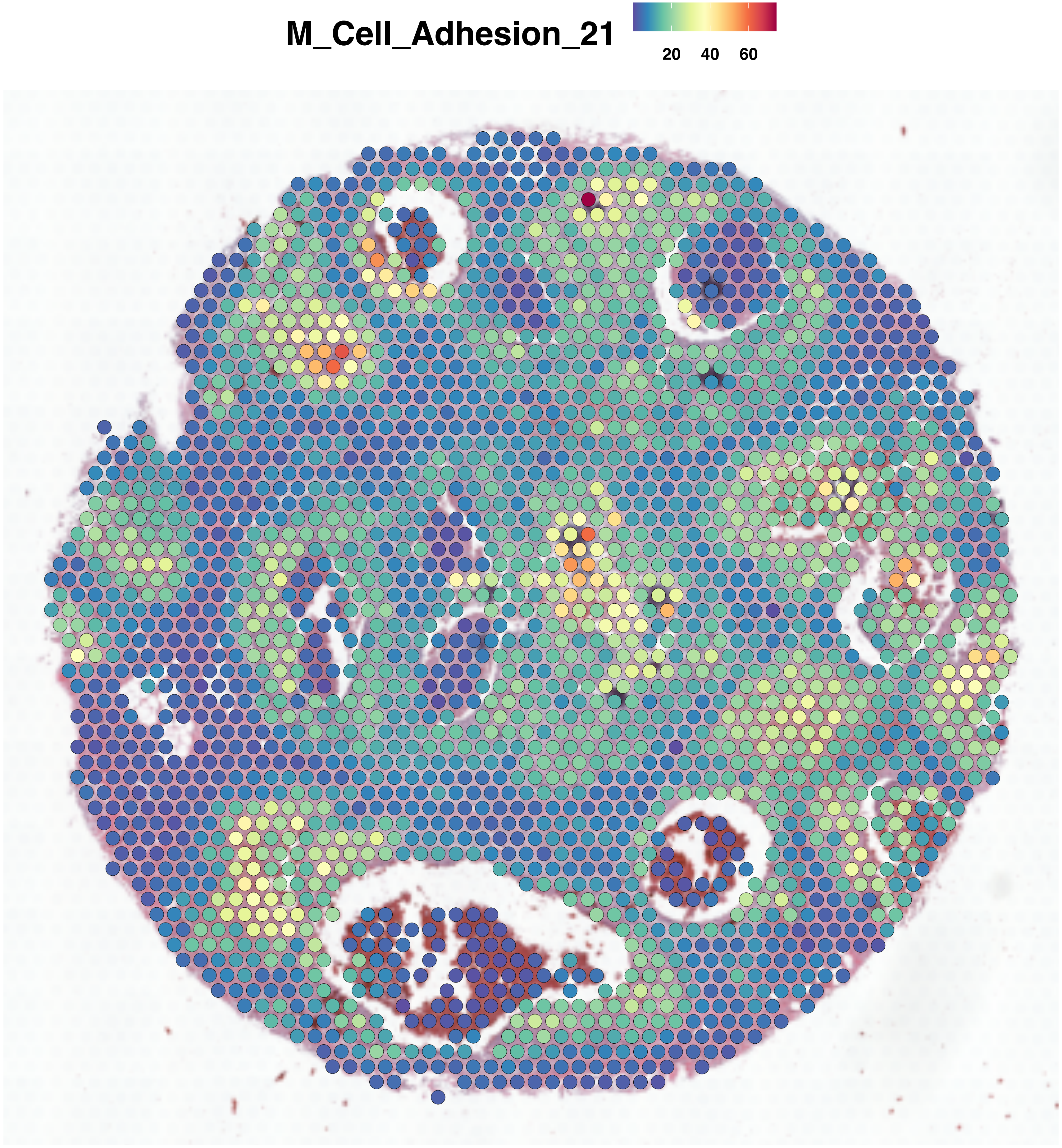
M_Cell_Adhesion_21
20
40
60

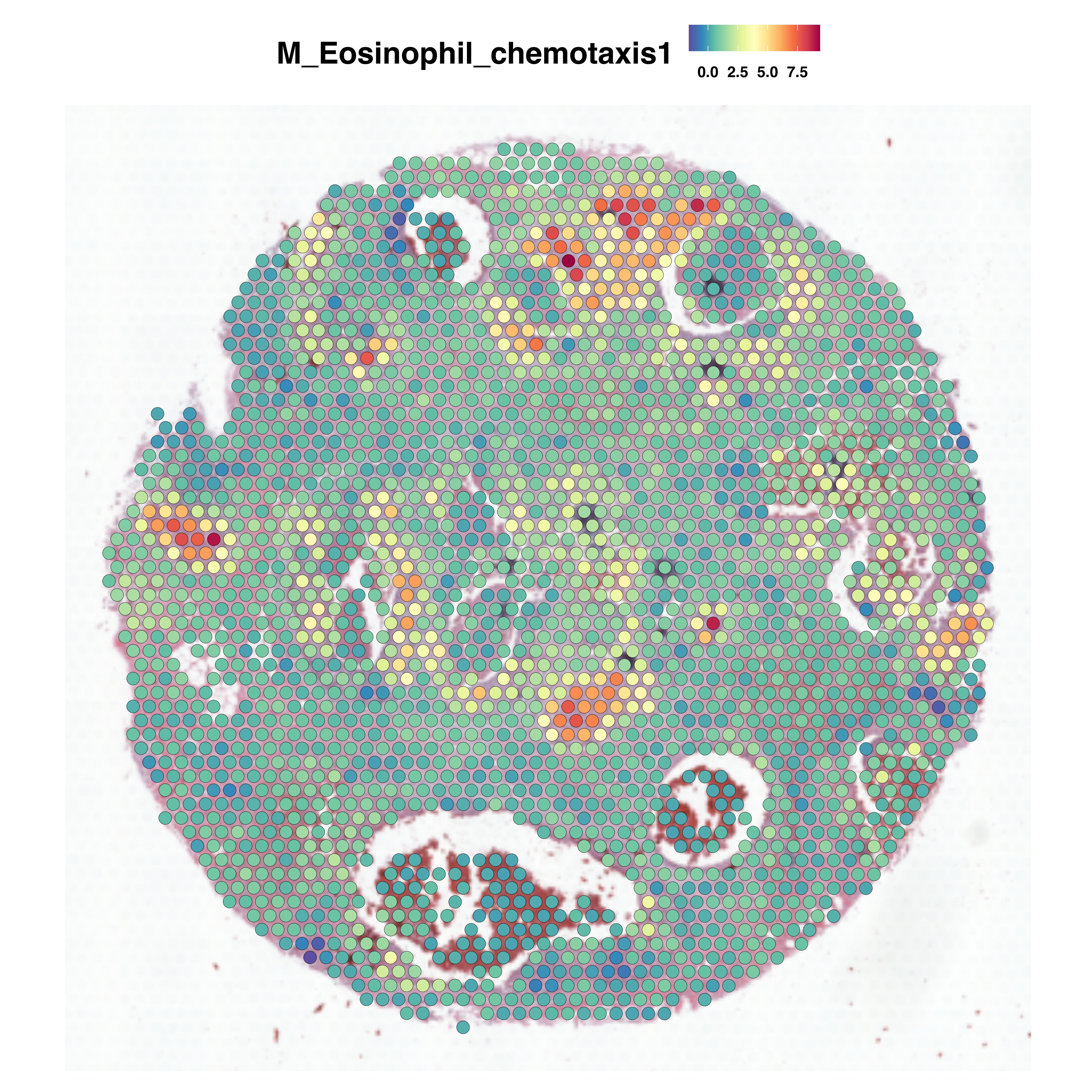
M_Eosinophil_chemotaxis1
0.0 2.5 5.0 7.5

### M_MHC2_mediated_Lymphocyte_Activation1

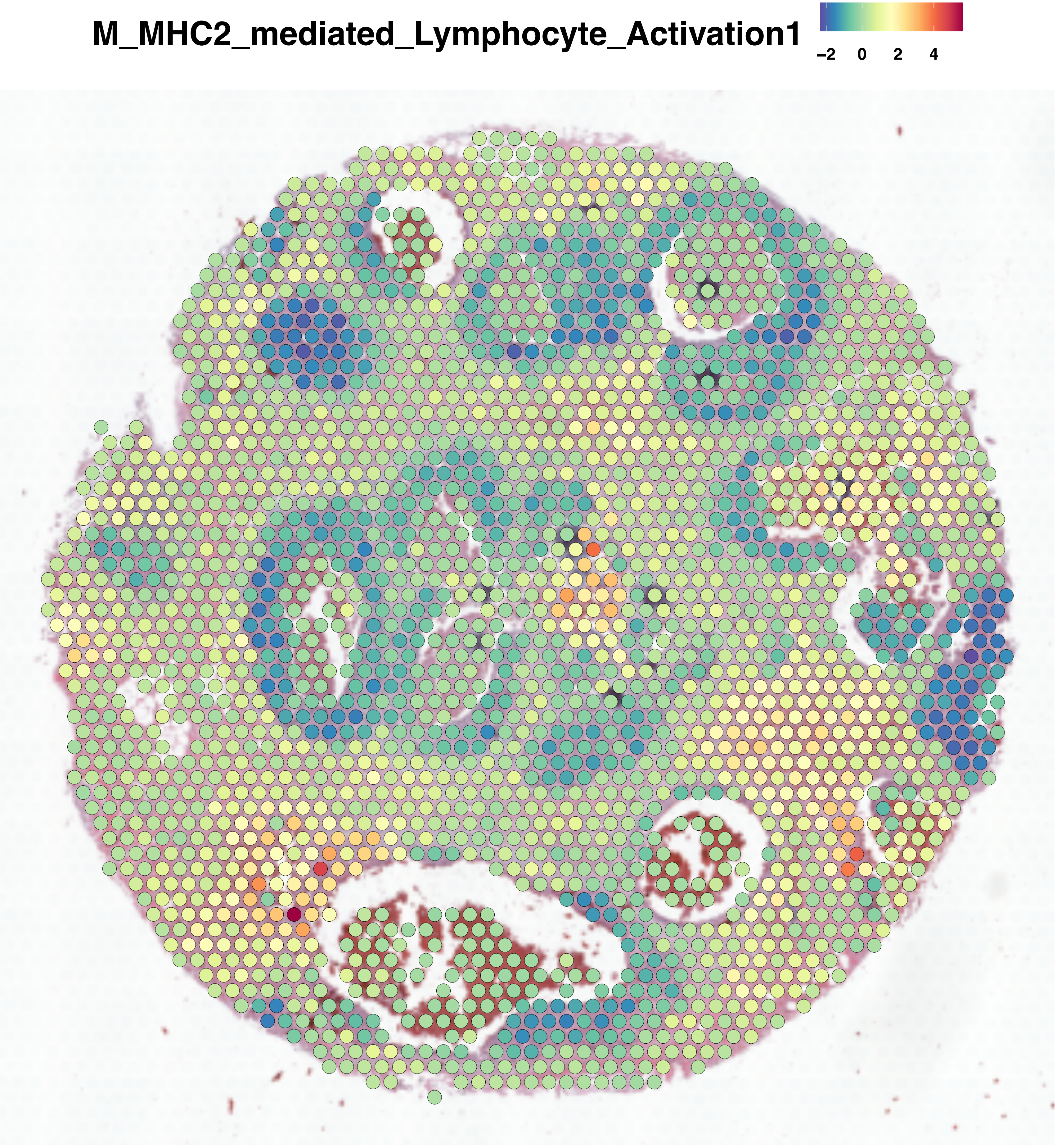

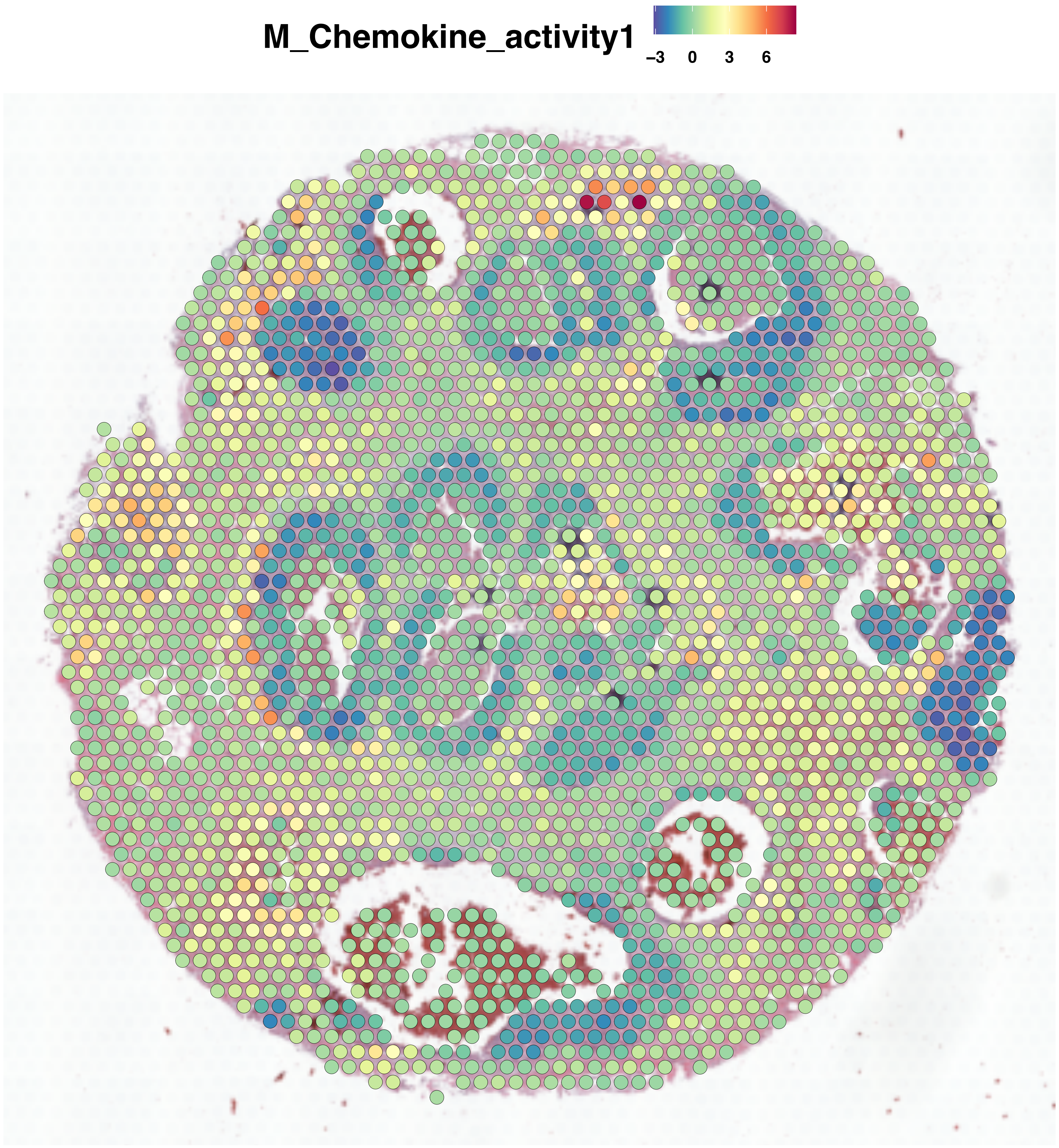
M_Chemokine_activity1
−3
0
3
6

**Figure S14. [Gene sets annotate spatial transcriptomics], Related to Figure 6.**
irMPs module score overlaid on breast cancer spatial spots. Color represents activity level. Warmer colors indicate higher level of corresponding irMP activity and colder colors indicate lower level of corresponding irMP activity.

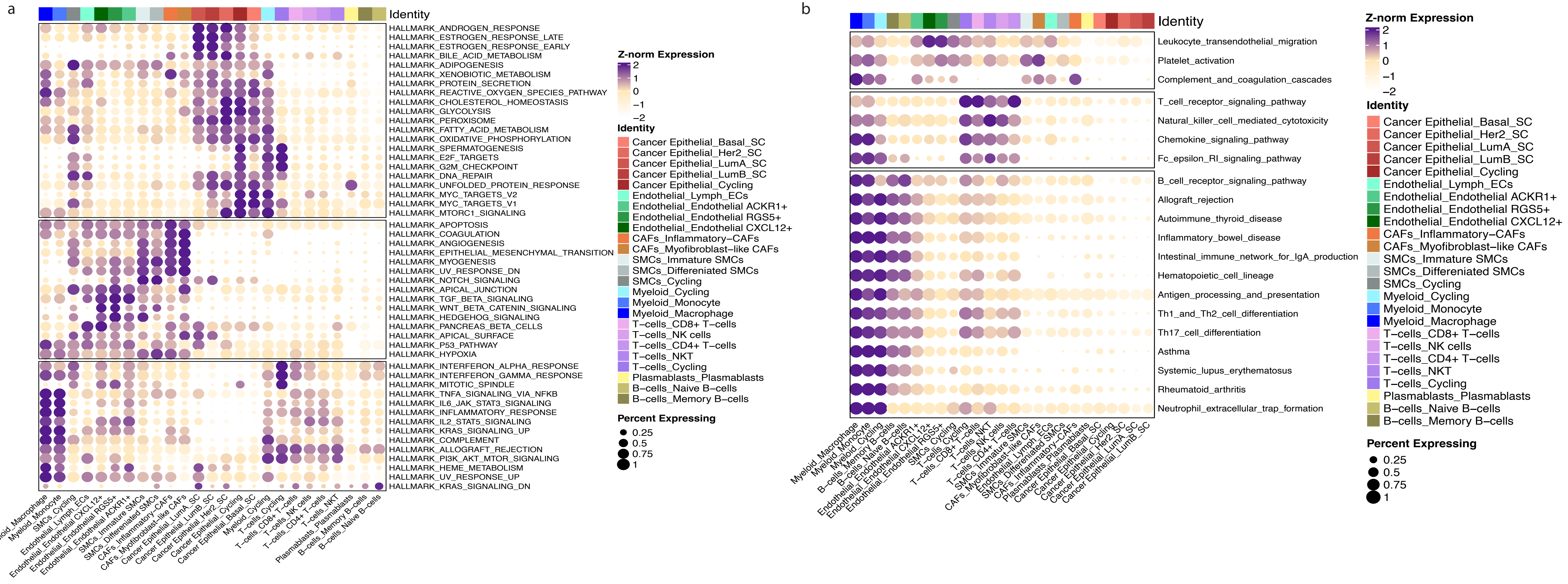
a
Identity
HALLMARK_ANDROGEN_RESPONSE
HALLMARK_ESTROGEN_RESPONSE_LATE
HALLMARK_ESTROGEN_RESPONSE_EARLY
HALLMARK_BILE_ACID_METABOLISM
HALLMARK_ADIPOGENESIS
HALLMARK_XENOBIOTIC_METABOLISM
HALLMARK_PROTEIN_SECRETION
HALLMARK_REACTIVE_OXYGEN_SPECIES_PATHWAY
HALLMARK_CHOLESTEROL_HOMEOSTASIS
HALLMARK_GLYCOLYSIS
HALLMARK_PEROXISOME
HALLMARK_FATTY_ACID_METABOLISM
HALLMARK_OXIDATIVE_PHOSPHORYLATION
HALLMARK_SPERMATOGENESIS
HALLMARK_E2F_TARGETS
HALLMARK_G2M_CHECKPOINT
HALLMARK_DNA_REPAIR
HALLMARK_UNFOLDED_PROTEIN_RESPONSE
HALLMARK_MYC_TARGETS_V2
HALLMARK_MYC_TARGETS_V1
HALLMARK_MTORC1_SIGNALING
HALLMARK_APOPTOSIS
HALLMARK_COAGULATION
HALLMARK_ANGIOGENESIS
HALLMARK_EPITHELIAL_MESENCHYMAL_TRANSITION
HALLMARK_MYOGENESIS
HALLMARK_UV_RESPONSE_DN
HALLMARK_NOTCH_SIGNALING
HALLMARK_APICAL_JUNCTION
HALLMARK_TGF_BETA_SIGNALING
HALLMARK_WNT_BETA_CATENIN_SIGNALING
HALLMARK_HEDGEHOG_SIGNALING
HALLMARK_PANCREAS_BETA_CELLS
HALLMARK_APICAL_SURFACE
HALLMARK_P53_PATHWAY
HALLMARK_HYPOXIA
HALLMARK_INTERFERON_ALPHA_RESPONSE
HALLMARK_INTERFERON_GAMMA_RESPONSE
HALLMARK_MITOTIC_SPINDLE
HALLMARK_TNFA_SIGNALING_VIA_NFKB
HALLMARK_IL6_JAK_STAT3_SIGNALING
HALLMARK_INFLAMMATORY_RESPONSE
HALLMARK_IL2_STAT5_SIGNALING
HALLMARK_KRAS_SIGNALING_UP
HALLMARK_COMPLEMENT
HALLMARK_ALLOGRAFT_REJECTION
HALLMARK_PI3K_AKT_MTOR_SIGNALING
HALLMARK_HEME_METABOLISM
HALLMARK_UV_RESPONSE_UP
HALLMARK_KRAS_SIGNALING_DN
Myeloid_Macrophage
Myeloid_Monocyte
SMCs_Cycling
Endothelial_Lymph_ECs
Endothelial_Endothelial CXCL12+
Endothelial_Endothelial RGS5+
Endothelial_Endothelial ACKR1+
SMCs_Immature SMCs
SMCs_Differeniated SMCs
CAFs_Inflammatory-CAFs
CAFs_Myofibroblast-like CAFs
Cancer Epithelial_LumA_SC
Cancer Epithelial_LumB_SC
Cancer Epithelial_Her2_SC
Cancer Epithelial_Cycling
Cancer Epithelial_Basal_SC
Myeloid_Cycling
T-cells_Cycling
T-cells_CD8+ T-cells
T-cells_NK cells
T-cells_CD4+ T-cells
T-cells_NKT
Plasmablasts_Plasmablasts
B-cells_Memory B-cells
B-cells_Naive B-cells
Z-norm Expression
2
1
0
−1
−2
Identity
Cancer Epithelial_Basal_SC
Cancer Epithelial_Her2_SC
Cancer Epithelial_LumA_SC
Cancer Epithelial_LumB_SC
Cancer Epithelial_Cycling
Endothelial_Lymph_ECs
Endothelial_Endothelial ACKR1+
Endothelial_Endothelial RGS5+
Endothelial_Endothelial CXCL12+
CAFs_Inflammatory-CAFs
CAFs_Myofibroblast-like CAFs
SMCs_Immature SMCs
SMCs_Differeniated SMCs
SMCs_Cycling
Myeloid_Cycling
Myeloid_Monocyte
Myeloid_Macrophage
T-cells_CD8+ T-cells
T-cells_NK cells
T-cells_CD4+ T-cells
T-cells_NKT
T-cells_Cycling
Plasmablasts_Plasmablasts
B-cells_Naive B-cells
B-cells_Memory B-cells
Percent Expressing
0.25
0.5
0.75
1
b
Identity
Leukocyte_transendothelial_migration
Platelet_activation
Complement_and_coagulation_cascades
T_cell_receptor_signaling_pathway
Natural_killer_cell_mediated_cytotoxicity
Chemokine_signaling_pathway
Fc_epsilon_RI_signaling_pathway
B_cell_receptor_signaling_pathway
Allograft_rejection
Autoimmune_thyroid_disease
Inflammatory_bowel_disease
Intestinal_immune_network_for_IgA_production
Hematopoietic_cell_lineage
Antigen_processing_and_presentation
Th1_and_Th2_cell_differentiation
Th17_cell_differentiation
Asthma
Systemic_lupus_erythematosus
Rheumatoid_arthritis
Neutrophil_extracellular_trap_formation
Myeloid_Macrophage
Myeloid_Monocyte
Myeloid_Cycling
B-cells_Memory B-cells
B-cells_Naive B-cells
Endothelial_Endothelial ACKR1+
Endothelial_Endothelial CXCL12+
Endothelial_Endothelial RGS5+
SMCs_Cycling
T-cells_Cycling
T-cells_CD8+ T-cells
T-cells_NKT
T-cells_NK cells
T-cells_CD4+ T-cells
SMCs_Immature SMCs
CAFs_Myofibroblast-like CAFs
Endothelial_Lymph_ECs
SMCs_Differeniated SMCs
CAFs_Inflammatory-CAFs
Plasmablasts_Plasmablasts
Cancer Epithelial_Basal_SC
Cancer Epithelial_Cycling
Cancer Epithelial_Her2_SC
Cancer Epithelial_LumA_SC
Cancer Epithelial_LumB_SC
Z-norm Expression
2
1
0
−1
−2
Identity
Cancer Epithelial_Basal_SC
Cancer Epithelial_Her2_SC
Cancer Epithelial_LumA_SC
Cancer Epithelial_LumB_SC
Cancer Epithelial_Cycling
Endothelial_Lymph_ECs
Endothelial_Endothelial ACKR1+
Endothelial_Endothelial RGS5+
Endothelial_Endothelial CXCL12+
CAFs_Inflammatory-CAFs
CAFs_Myofibroblast-like CAFs
SMCs_Immature SMCs
SMCs_Differeniated SMCs
SMCs_Cycling
Myeloid_Cycling
Myeloid_Monocyte
Myeloid_Macrophage
T-cells_CD8+ T-cells
T-cells_NK cells
T-cells_CD4+ T-cells
T-cells_NKT
T-cells_Cycling
Plasmablasts_Plasmablasts
B-cells_Naive B-cells
B-cells_Memory B-cells
Percent Expressing
0.25
0.5
0.75
1

**Figure S15. [Gene sets annotate spatial transcriptomics], Related to Figure 6.**
(**a**) Dot plot comparing the Z-transformed expression level of 50 Hallmark pathways across different celltrek-annotated cell types. (**b**) Dot plot comparing the Z-transformed expression level of 29 immune-related KEGG pathway across different celltrek-annotated cell types. Color represents the average expression level, and the dot size represents the percentage of cells expressing the irMP activity.

#### Ovarian Cancer

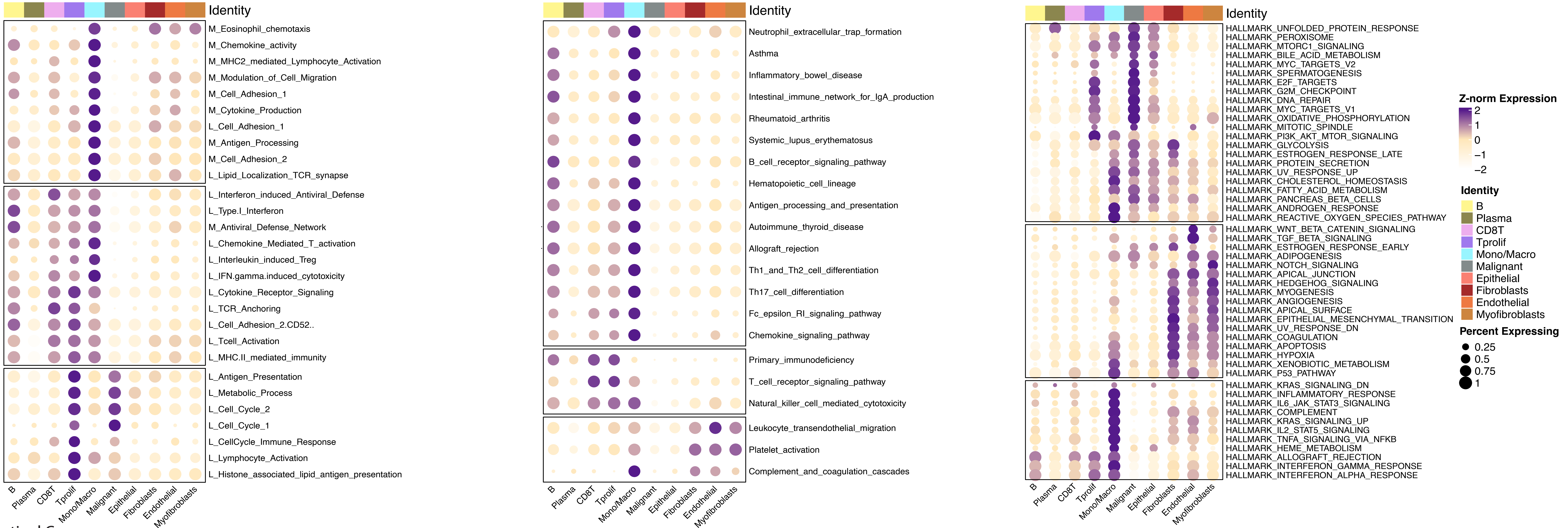


#### Intestinal Cancer

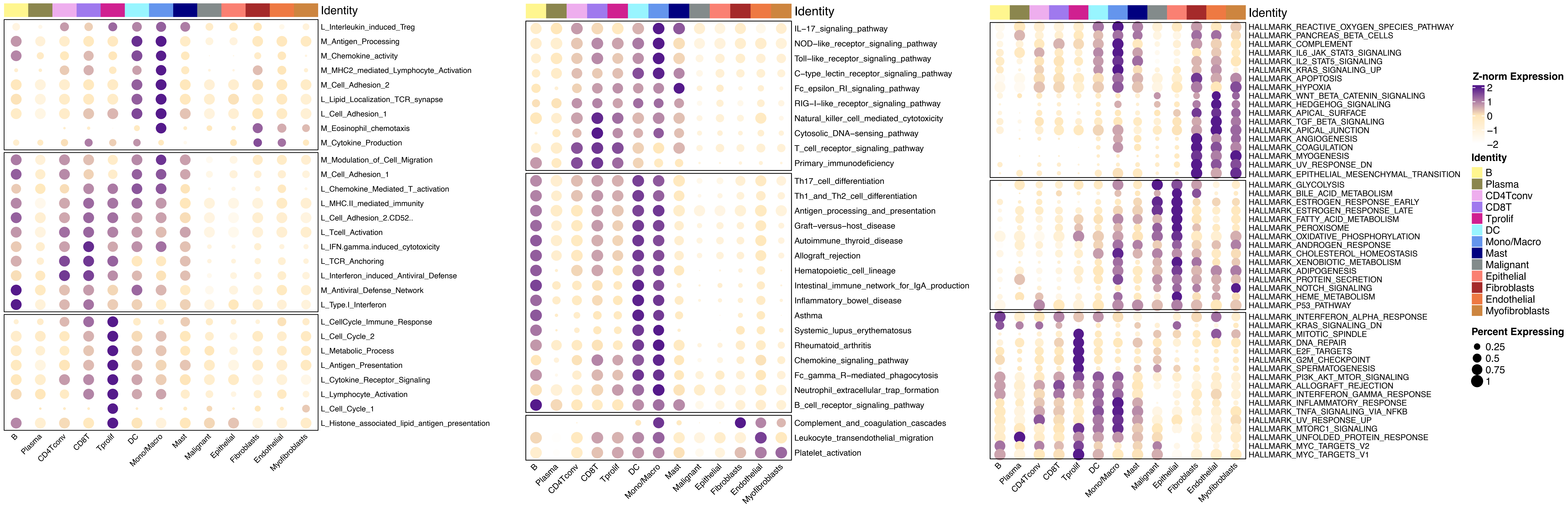

**Figure S16. [Gene sets annotate spatial transcriptomics], Related to Figure 6.**

Dot plot comparing the Z-transformed expression level of 28 irMPs, 29 irKEGGs and 50 Hallmark pathways across different celltrek-annotated cell types in ovarian cancer and intestine cancer.

| ID | Description | p.adjust | Count |
| --- | --- | --- | --- |
| GO:0000280 | nuclear division | 2.00456E-11 | 16 |
| GO:0007059 | chromosome segregation | 2.00456E-11 | 15 |
| GO:0098813 | nuclear chromosome segregation | 4.16211E-10 | 13 |
| GO:0044772 | mitotic cell cycle phase transition | 1.98568E-08 | 13 |
| HALLMARK_G2M_CHECKPOINT | HALLMARK_G2M_CHECKPOINT | 1.26215E-09 | 13 |
| GO:0140014 | mitotic nuclear division | 5.09241E-09 | 12 |
| GO:0000070 | mitotic sister chromatid segregation | 9.64624E-10 | 11 |
| GO:0000819 | sister chromatid segregation | 4.29875E-09 | 11 |
| HALLMARK_E2F_TARGETS | HALLMARK_E2F_TARGETS | 1.59963E-07 | 11 |
| GO:0007051 | spindle organization | 9.62668E-09 | 10 |
| GO:0010639 | negative regulation of organelle organization | 8.43886E-07 | 10 |
| GO:1901987 | regulation of cell cycle phase transition | 5.27568E-06 | 10 |
| HALLMARK_MITOTIC_SPINDLE | HALLMARK_MITOTIC_SPINDLE | 1.30571E-06 | 10 |
| GO:0010948 | negative regulation of cell cycle process | 3.01906E-06 | 9 |
| GO:1901990 | regulation of mitotic cell cycle phase transition | 6.78657E-06 | 9 |
| GO:0045786 | negative regulation of cell cycle | 1.79965E-05 | 9 |
| GO:0051983 | regulation of chromosome segregation | 1.58071E-07 | 8 |
| GO:0007052 | mitotic spindle organization | 1.58071E-07 | 8 |
| GO:1902850 | microtubule cytoskeleton organization involved in mitosis | 3.07588E-07 | 8 |
| GO:0045930 | negative regulation of mitotic cell cycle | 5.3426E-06 | 8 |
| GO:1901988 | negative regulation of cell cycle phase transition | 1.09526E-05 | 8 |
| GO:0140694 | non-membrane-bounded organelle assembly | 0.000116973 | 8 |
| GO:0071900 | regulation of protein serine/threonine kinase activity | 0.000136018 | 8 |
| GO:0019887 | protein kinase regulator activity | 0.000263145 | 7 |
| GO:0019207 | kinase regulator activity | 0.000292707 | 7 |
| GO:0008017 | microtubule binding | 0.000347193 | 7 |
| GO:0015631 | tubulin binding | 0.001475524 | 7 |
| GO:0030071 | regulation of mitotic metaphase/anaphase transition | 2.00349E-07 | 7 |
| GO:0007091 | metaphase/anaphase transition of mitotic cell cycle | 2.09364E-07 | 7 |
| GO:1902099 | regulation of metaphase/anaphase transition of cell cycle | 2.09364E-07 | 7 |
| GO:0010965 | regulation of mitotic sister chromatid separation | 2.40544E-07 | 7 |
| GO:0044784 | metaphase/anaphase transition of cell cycle | 2.40544E-07 | 7 |
| GO:0051306 | mitotic sister chromatid separation | 2.66555E-07 | 7 |
| GO:0033045 | regulation of sister chromatid segregation | 3.7365E-07 | 7 |
| GO:1905818 | regulation of chromosome separation | 4.53628E-07 | 7 |
| GO:0007088 | regulation of mitotic nuclear division | 6.74197E-07 | 7 |
| GO:0051304 | chromosome separation | 1.61736E-06 | 7 |
| GO:0007093 | mitotic cell cycle checkpoint signaling | 2.1232E-06 | 7 |
| GO:0051783 | regulation of nuclear division | 2.50555E-06 | 7 |
| GO:0000075 | cell cycle checkpoint signaling | 1.15342E-05 | 7 |
| GO:0000910 | cytokinesis | 1.16994E-05 | 7 |
| GO:0051302 | regulation of cell division | 1.27483E-05 | 7 |
| GO:0140013 | meiotic nuclear division | 1.40706E-05 | 7 |
| GO:1901991 | negative regulation of mitotic cell cycle phase transition | 1.40706E-05 | 7 |
| GO:1903046 | meiotic cell cycle process | 2.4047E-05 | 7 |
| GO:0033044 | regulation of chromosome organization | 7.43486E-05 | 7 |
| GO:0051321 | meiotic cell cycle | 0.000152225 | 7 |
| hsa04110 | Cell cycle | 3.55235E-06 | 7 |
| GO:0007094 | mitotic spindle assembly checkpoint signaling | 1.58071E-07 | 6 |
| GO:0071173 | spindle assembly checkpoint signaling | 1.58071E-07 | 6 |
| GO:0071174 | mitotic spindle checkpoint signaling | 1.58071E-07 | 6 |
| GO:0031577 | spindle checkpoint signaling | 1.67932E-07 | 6 |
| GO:0045841 | negative regulation of mitotic metaphase/anaphase transition | 1.78778E-07 | 6 |
| GO:1902100 | negative regulation of metaphase/anaphase transition of cell cycl | 2.00349E-07 | 6 |
| GO:0033046 | negative regulation of sister chromatid segregation | 2.00349E-07 | 6 |
| GO:0033048 | negative regulation of mitotic sister chromatid segregation | 2.00349E-07 | 6 |
| GO:2000816 | negative regulation of mitotic sister chromatid separation | 2.00349E-07 | 6 |
| GO:0051985 | negative regulation of chromosome segregation | 2.09364E-07 | 6 |
| GO:1905819 | negative regulation of chromosome separation | 2.09364E-07 | 6 |
| GO:0033047 | regulation of mitotic sister chromatid segregation | 2.49475E-07 | 6 |
| GO:0045839 | negative regulation of mitotic nuclear division | 2.49475E-07 | 6 |
| GO:0051784 | negative regulation of nuclear division | 4.50606E-07 | 6 |
| GO:2001251 | negative regulation of chromosome organization | 4.90592E-06 | 6 |
| GO:0051225 | spindle assembly | 1.98778E-05 | 6 |
| GO:0090068 | positive regulation of cell cycle process | 0.000943058 | 6 |
| GO:0007265 | Ras protein signal transduction | 0.004494329 | 6 |
| GO:0045787 | positive regulation of cell cycle | 0.005068123 | 6 |

| | | | |
|---|---|---|---|
| GO:0007018 | microtubule-based movement | 0.008795563 | 6 |
| GO:0033674 | positive regulation of kinase activity | 0.01809306 | 6 |
| GO:0032465 | regulation of cytokinesis | 9.01223E-05 | 5 |
| GO:0000079 | regulation of cyclin-dependent protein serine/threonine kinase ac | 0.000232793 | 5 |
| GO:1904029 | regulation of cyclin-dependent protein kinase activity | 0.000258865 | 5 |
| GO:0044843 | cell cycle G1/S phase transition | 0.009730801 | 5 |
| GO:0006260 | DNA replication | 0.010456276 | 5 |
| GO:0051056 | regulation of small GTPase mediated signal transduction | 0.012895459 | 5 |
| GO:0001933 | negative regulation of protein phosphorylation | 0.018269119 | 5 |
| GO:0042326 | negative regulation of phosphorylation | 0.028871643 | 5 |
| GO:0043254 | regulation of protein-containing complex assembly | 0.036510589 | 5 |
| GO:0045860 | positive regulation of protein kinase activity | 0.04008853 | 5 |
| GO:0022412 | cellular process involved in reproduction in multicellular organism | 0.042841017 | 5 |
| GO:0045936 | negative regulation of phosphate metabolic process | 0.04588861 | 5 |
| GO:0010563 | negative regulation of phosphorus metabolic process | 0.04588861 | 5 |
| hsa05166 | Human T-cell leukemia virus 1 infection | 0.005066617 | 5 |
| GO:0016538 | cyclin-dependent protein serine/threonine kinase regulator activi | 0.000347193 | 4 |
| GO:0003777 | microtubule motor activity | 0.000893595 | 4 |
| GO:0003774 | cytoskeletal motor activity | 0.004568766 | 4 |
| GO:0008608 | attachment of spindle microtubules to kinetochore | 9.01223E-05 | 4 |
| GO:0090307 | mitotic spindle assembly | 0.000706359 | 4 |
| GO:0000281 | mitotic cytokinesis | 0.001178627 | 4 |
| GO:0050000 | chromosome localization | 0.001367104 | 4 |
| GO:0048144 | fibroblast proliferation | 0.002374032 | 4 |
| GO:0061640 | cytoskeleton-dependent cytokinesis | 0.003292507 | 4 |
| GO:0006275 | regulation of DNA replication | 0.006054015 | 4 |
| GO:0007266 | Rho protein signal transduction | 0.006492597 | 4 |
| GO:0044839 | cell cycle G2/M phase transition | 0.009276193 | 4 |
| GO:0051494 | negative regulation of cytoskeleton organization | 0.010659486 | 4 |
| GO:0006261 | DNA-templated DNA replication | 0.010779515 | 4 |
| GO:1902904 | negative regulation of supramolecular fiber organization | 0.011326541 | 4 |
| GO:0046578 | regulation of Ras protein signal transduction | 0.016911168 | 4 |
| GO:0000082 | G1/S transition of mitotic cell cycle | 0.036123266 | 4 |
| GO:0051348 | negative regulation of transferase activity | 0.046128306 | 4 |
| GO:0051258 | protein polymerization | 0.04837857 | 4 |
| hsa04114 | Oocyte meiosis | 0.005335146 | 4 |
| hsa04218 | Cellular senescence | 0.007790408 | 4 |
| hsa04814 | Motor proteins | 0.011556335 | 4 |
| HALLMARK_SPERMATOGENESIS | HALLMARK_SPERMATOGENESIS | 0.039349182 | 4 |
| GO:0034508 | centromere complex assembly | 0.001292209 | 3 |
| GO:0032506 | cytokinetic process | 0.002423159 | 3 |
| GO:0048146 | positive regulation of fibroblast proliferation | 0.005247619 | 3 |
| GO:0007080 | mitotic metaphase plate congression | 0.006054015 | 3 |
| GO:0090329 | regulation of DNA-templated DNA replication | 0.006054015 | 3 |
| GO:0051445 | regulation of meiotic cell cycle | 0.008091096 | 3 |
| GO:0051310 | metaphase plate congression | 0.010321954 | 3 |
| GO:0051303 | establishment of chromosome localization | 0.014192022 | 3 |
| GO:0048145 | regulation of fibroblast proliferation | 0.016485534 | 3 |
| GO:0035023 | regulation of Rho protein signal transduction | 0.016862597 | 3 |
| GO:0051781 | positive regulation of cell division | 0.018269119 | 3 |
| GO:0070301 | cellular response to hydrogen peroxide | 0.018269119 | 3 |
| GO:0045132 | meiotic chromosome segregation | 0.022695802 | 3 |
| GO:0031109 | microtubule polymerization or depolymerization | 0.036510589 | 3 |
| GO:0042542 | response to hydrogen peroxide | 0.040926129 | 3 |
| GO:0007127 | meiosis I | 0.041469779 | 3 |
| GO:0061982 | meiosis I cell cycle process | 0.045489747 | 3 |
| GO:0000086 | G2/M transition of mitotic cell cycle | 0.048335229 | 3 |
| hsa04115 | p53 signaling pathway | 0.009234145 | 3 |
| hsa04914 | Progesterone-mediated oocyte maturation | 0.01671744 | 3 |
| GO:0044771 | meiotic cell cycle phase transition | 0.005247619 | 2 |
| GO:0051256 | mitotic spindle midzone assembly | 0.005247619 | 2 |
| GO:0000022 | mitotic spindle elongation | 0.006054015 | 2 |
| GO:0051231 | spindle elongation | 0.007883442 | 2 |
| GO:0051255 | spindle midzone assembly | 0.007883442 | 2 |
| GO:0000212 | meiotic spindle organization | 0.008795563 | 2 |
| GO:0051782 | negative regulation of cell division | 0.011326541 | 2 |
| GO:0051988 | regulation of attachment of spindle microtubules to kinetochore | 0.011326541 | 2 |
| GO:0061952 | midbody abscission | 0.011326541 | 2 |

| | | | |
|---|---|---|---|
| GO:0051444 | negative regulation of ubiquitin-protein transferase activity | 0.016457197 | 2 |
| GO:0035024 | negative regulation of Rho protein signal transduction | 0.017274979 | 2 |
| GO:0036120 | cellular response to platelet-derived growth factor stimulus | 0.018266312 | 2 |
| GO:1902410 | mitotic cytokinetic process | 0.018266312 | 2 |
| GO:0036119 | response to platelet-derived growth factor | 0.020636829 | 2 |
| GO:0010528 | regulation of transposition | 0.024953084 | 2 |
| GO:0010529 | negative regulation of transposition | 0.024953084 | 2 |
| GO:0051642 | centrosome localization | 0.029792238 | 2 |
| GO:0007143 | female meiotic nuclear division | 0.030836045 | 2 |
| GO:0032196 | transposition | 0.030836045 | 2 |
| GO:0061842 | microtubule organizing center localization | 0.030836045 | 2 |
| GO:0003382 | epithelial cell morphogenesis | 0.032427293 | 2 |
| GO:0035633 | maintenance of blood-brain barrier | 0.034042409 | 2 |
| GO:0030866 | cortical actin cytoskeleton organization | 0.041761948 | 2 |
| GO:0044786 | cell cycle DNA replication | 0.045217731 | 2 |
| GO:0071459 | protein localization to chromosome, centromeric region | 0.04588861 | 2 |

| ID | Description | p.adjust | Count |
|---|---|---|---|
| GO:0050851 | antigen receptor-mediated signaling pathway | 1.10111E-11 | 13 |
| GO:0002429 | immune response-activating cell surface receptor signaling pathw | 8.52078E-11 | 13 |
| GO:0002757 | immune response-activating signal transduction | 8.52078E-11 | 13 |
| GO:0002768 | immune response-regulating cell surface receptor signaling pathw | 2.10927E-10 | 13 |
| GO:0002253 | activation of immune response | 1.1959E-09 | 13 |
| GO:0050852 | T cell receptor signaling pathway | 5.02932E-13 | 12 |
| GO:0030098 | lymphocyte differentiation | 6.6707E-06 | 10 |
| GO:1903131 | mononuclear cell differentiation | 1.83502E-05 | 10 |
| GO:0050863 | regulation of T cell activation | 2.64498E-05 | 9 |
| HALLMARK_ALLOGRAFT_REJECTION | HALLMARK_ALLOGRAFT_REJECTION | 4.68465E-06 | 9 |
| GO:0046631 | alpha-beta T cell activation | 1.25965E-06 | 8 |
| GO:1903037 | regulation of leukocyte cell-cell adhesion | 0.000270511 | 8 |
| GO:0007159 | leukocyte cell-cell adhesion | 0.000430975 | 8 |
| GO:0002696 | positive regulation of leukocyte activation | 0.000680424 | 8 |
| GO:0050867 | positive regulation of cell activation | 0.000816758 | 8 |
| GO:0022407 | regulation of cell-cell adhesion | 0.001044308 | 8 |
| GO:1903039 | positive regulation of leukocyte cell-cell adhesion | 0.000312192 | 7 |
| GO:0030217 | T cell differentiation | 0.000442559 | 7 |
| GO:0022409 | positive regulation of cell-cell adhesion | 0.000680424 | 7 |
| GO:0051251 | positive regulation of lymphocyte activation | 0.002136804 | 7 |
| GO:0002683 | negative regulation of immune system process | 0.003904999 | 7 |
| GO:0002443 | leukocyte mediated immunity | 0.004551211 | 7 |
| GO:0045785 | positive regulation of cell adhesion | 0.005349 | 7 |
| GO:0001819 | positive regulation of cytokine production | 0.005349 | 7 |
| hsa04660 | T cell receptor signaling pathway | 1.08089E-06 | 7 |
| GO:0050870 | positive regulation of T cell activation | 0.001462294 | 6 |
| GO:0042113 | B cell activation | 0.005603031 | 6 |
| GO:0002449 | lymphocyte mediated immunity | 0.006652928 | 6 |
| GO:1903706 | regulation of hemopoiesis | 0.009840837 | 6 |
| hsa04650 | Natural killer cell mediated cytotoxicity | 2.74787E-05 | 6 |
| GO:0050854 | regulation of antigen receptor-mediated signaling pathway | 5.14968E-05 | 5 |
| GO:0050864 | regulation of B cell activation | 0.004943256 | 5 |
| GO:0045619 | regulation of lymphocyte differentiation | 0.005349 | 5 |
| GO:0002699 | positive regulation of immune effector process | 0.009840837 | 5 |
| GO:1902105 | regulation of leukocyte differentiation | 0.01596386 | 5 |
| GO:0006959 | humoral immune response | 0.015964122 | 5 |
| GO:0002460 | adaptive immune response based on somatic recombination of im | 0.025679864 | 5 |
| GO:0002697 | regulation of immune effector process | 0.026273957 | 5 |
| hsa05235 | PD-L1 expression and PD-1 checkpoint pathway in cancer | 5.92038E-05 | 5 |
| GO:0050856 | regulation of T cell receptor signaling pathway | 0.000309749 | 4 |
| GO:0033077 | T cell differentiation in thymus | 0.002435937 | 4 |
| GO:0002708 | positive regulation of lymphocyte mediated immunity | 0.005603031 | 4 |
| GO:0032609 | interferon-gamma production | 0.005603031 | 4 |
| GO:0032649 | regulation of interferon-gamma production | 0.005603031 | 4 |
| GO:0002821 | positive regulation of adaptive immune response | 0.005603031 | 4 |
| GO:0002702 | positive regulation of production of molecular mediator of immun | 0.007695919 | 4 |
| GO:0002705 | positive regulation of leukocyte mediated immunity | 0.009130849 | 4 |
| GO:0032946 | positive regulation of mononuclear cell proliferation | 0.009840837 | 4 |
| GO:0070665 | positive regulation of leukocyte proliferation | 0.013311286 | 4 |
| GO:0002706 | regulation of lymphocyte mediated immunity | 0.015197239 | 4 |
| GO:0032640 | tumor necrosis factor production | 0.01596386 | 4 |
| GO:0032680 | regulation of tumor necrosis factor production | 0.01596386 | 4 |
| GO:0002700 | regulation of production of molecular mediator of immune respon | 0.016458289 | 4 |
| GO:0071706 | tumor necrosis factor superfamily cytokine production | 0.016636791 | 4 |
| GO:1903555 | regulation of tumor necrosis factor superfamily cytokine producti | 0.016636791 | 4 |
| GO:0002819 | regulation of adaptive immune response | 0.019477488 | 4 |
| GO:0045637 | regulation of myeloid cell differentiation | 0.023247052 | 4 |
| GO:0002274 | myeloid leukocyte activation | 0.030007385 | 4 |
| GO:0002703 | regulation of leukocyte mediated immunity | 0.030516599 | 4 |
| GO:0032944 | regulation of mononuclear cell proliferation | 0.03087988 | 4 |
| GO:0070663 | regulation of leukocyte proliferation | 0.038627986 | 4 |
| GO:0002366 | leukocyte activation involved in immune response | 0.047468841 | 4 |
| GO:0048872 | homeostasis of number of cells | 0.048121036 | 4 |
| GO:0002263 | cell activation involved in immune response | 0.048121036 | 4 |
| hsa04658 | Th1 and Th2 cell differentiation | 0.001337066 | 4 |
| hsa04659 | Th17 cell differentiation | 0.001661689 | 4 |
| hsa04514 | Cell adhesion molecules | 0.006076985 | 4 |

| | | | |
|---|---|---|---|
| GO:0042169 | SH2 domain binding | 0.01299486 | 3 |
| GO:0045061 | thymic T cell selection | 0.00104353 | 3 |
| GO:0050858 | negative regulation of antigen receptor-mediated signaling pathv | 0.002902713 | 3 |
| GO:0031295 | T cell costimulation | 0.005603031 | 3 |
| GO:0031294 | lymphocyte costimulation | 0.005603031 | 3 |
| GO:0045058 | T cell selection | 0.006715553 | 3 |
| GO:0034113 | heterotypic cell-cell adhesion | 0.009840837 | 3 |
| GO:0032615 | interleukin-12 production | 0.009840837 | 3 |
| GO:0032623 | interleukin-2 production | 0.009840837 | 3 |
| GO:0032655 | regulation of interleukin-12 production | 0.009840837 | 3 |
| GO:0032663 | regulation of interleukin-2 production | 0.009840837 | 3 |
| GO:0046635 | positive regulation of alpha-beta T cell activation | 0.013558054 | 3 |
| GO:0042267 | natural killer cell mediated cytotoxicity | 0.013863719 | 3 |
| GO:0002228 | natural killer cell mediated immunity | 0.014560669 | 3 |
| GO:0032729 | positive regulation of interferon-gamma production | 0.014560669 | 3 |
| GO:0002720 | positive regulation of cytokine production involved in immune re: | 0.015197239 | 3 |
| GO:0045638 | negative regulation of myeloid cell differentiation | 0.022389354 | 3 |
| GO:0030101 | natural killer cell activation | 0.023247052 | 3 |
| GO:0042102 | positive regulation of T cell proliferation | 0.026273957 | 3 |
| GO:0002824 | positive regulation of adaptive immune response based on somat | 0.030516599 | 3 |
| GO:0046634 | regulation of alpha-beta T cell activation | 0.030516599 | 3 |
| GO:0002718 | regulation of cytokine production involved in immune response | 0.031038622 | 3 |
| GO:0002367 | cytokine production involved in immune response | 0.031902899 | 3 |
| GO:0045446 | endothelial cell differentiation | 0.032338478 | 3 |
| GO:0050853 | B cell receptor signaling pathway | 0.037369028 | 3 |
| GO:0001909 | leukocyte mediated cytotoxicity | 0.039578748 | 3 |
| GO:0003158 | endothelium development | 0.041954197 | 3 |
| GO:0014065 | phosphatidylinositol 3-kinase signaling | 0.045993879 | 3 |
| GO:0050671 | positive regulation of lymphocyte proliferation | 0.046488861 | 3 |
| GO:0030183 | B cell differentiation | 0.049108219 | 3 |
| hsa05340 | Primary immunodeficiency | 0.001522841 | 3 |
| hsa04064 | NF-kappa B signaling pathway | 0.018307514 | 3 |
| hsa05135 | Yersinia infection | 0.025464086 | 3 |
| hsa05162 | Measles | 0.025464086 | 3 |
| GO:0042608 | T cell receptor binding | 0.01299486 | 2 |
| GO:0002863 | positive regulation of inflammatory response to antigenic stimulu | 0.009840837 | 2 |
| GO:0045059 | positive thymic T cell selection | 0.010329513 | 2 |
| GO:0001787 | natural killer cell proliferation | 0.011459458 | 2 |
| GO:0034116 | positive regulation of heterotypic cell-cell adhesion | 0.011459458 | 2 |
| GO:0050862 | positive regulation of T cell receptor signaling pathway | 0.01283849 | 2 |
| GO:0033033 | negative regulation of myeloid cell apoptotic process | 0.014560669 | 2 |
| GO:0060644 | mammary gland epithelial cell differentiation | 0.014560669 | 2 |
| GO:0006925 | inflammatory cell apoptotic process | 0.021973543 | 2 |
| GO:0034114 | regulation of heterotypic cell-cell adhesion | 0.022608522 | 2 |
| GO:0046641 | positive regulation of alpha-beta T cell proliferation | 0.022608522 | 2 |
| GO:0050860 | negative regulation of T cell receptor signaling pathway | 0.022608522 | 2 |
| GO:0046629 | gamma-delta T cell activation | 0.023247052 | 2 |
| GO:0050857 | positive regulation of antigen receptor-mediated signaling pathw | 0.023247052 | 2 |
| GO:0032753 | positive regulation of interleukin-4 production | 0.024172712 | 2 |
| GO:0036037 | CD8-positive, alpha-beta T cell activation | 0.024172712 | 2 |
| GO:0045954 | positive regulation of natural killer cell mediated cytotoxicity | 0.024172712 | 2 |
| GO:0001773 | myeloid dendritic cell activation | 0.02660708 | 2 |
| GO:0002507 | tolerance induction | 0.030007385 | 2 |
| GO:0002717 | positive regulation of natural killer cell mediated immunity | 0.030516599 | 2 |
| GO:0033032 | regulation of myeloid cell apoptotic process | 0.03087988 | 2 |
| GO:0038094 | Fc-gamma receptor signaling pathway | 0.03087988 | 2 |
| GO:0045577 | regulation of B cell differentiation | 0.03087988 | 2 |
| GO:0032743 | positive regulation of interleukin-2 production | 0.031902899 | 2 |
| GO:0032633 | interleukin-4 production | 0.032689552 | 2 |
| GO:0032673 | regulation of interleukin-4 production | 0.032689552 | 2 |
| GO:0045777 | positive regulation of blood pressure | 0.03390697 | 2 |
| GO:0050869 | negative regulation of B cell activation | 0.03390697 | 2 |
| GO:0043368 | positive T cell selection | 0.037366642 | 2 |
| GO:0033028 | myeloid cell apoptotic process | 0.038604106 | 2 |
| GO:0006040 | amino sugar metabolic process | 0.041447323 | 2 |
| GO:0002714 | positive regulation of B cell mediated immunity | 0.041954197 | 2 |
| GO:0002891 | positive regulation of immunoglobulin mediated immune respons | 0.041954197 | 2 |
| GO:0032735 | positive regulation of interleukin-12 production | 0.041954197 | 2 |

| | | | |
|---|---|---|---|
| GO:0032689 | negative regulation of interferon-gamma production | 0.045175075 | 2 |
| GO:0046640 | regulation of alpha-beta T cell proliferation | 0.045175075 | 2 |
| GO:0002861 | regulation of inflammatory response to antigenic stimulus | 0.048121036 | 2 |
| GO:0042269 | regulation of natural killer cell mediated cytotoxicity | 0.048121036 | 2 |
| GO:0046633 | alpha-beta T cell proliferation | 0.048121036 | 2 |
| GO:0061028 | establishment of endothelial barrier | 0.049352503 | 2 |
| hsa05330 | Allograft rejection | 0.025464086 | 2 |
| hsa05332 | Graft-versus-host disease | 0.025464086 | 2 |
| hsa04940 | Type I diabetes mellitus | 0.025464086 | 2 |
| hsa04672 | Intestinal immune network for IgA production | 0.030494984 | 2 |
| hsa05320 | Autoimmune thyroid disease | 0.033134909 | 2 |
| hsa05416 | Viral myocarditis | 0.039450019 | 2 |
| hsa04664 | Fc epsilon RI signaling pathway | 0.047169939 | 2 |

| ID | Description | p.adjust | Count |
|---|---|---|---|
| GO:0033218 | amide binding | 0.009495596 | 5 |
| GO:0019882 | antigen processing and presentation | 0.000113011 | 5 |
| GO:0002456 | T cell mediated immunity | 0.000114814 | 5 |
| GO:0001906 | cell killing | 0.000631977 | 5 |
| GO:0009165 | nucleotide biosynthetic process | 0.003493727 | 5 |
| GO:1901293 | nucleoside phosphate biosynthetic process | 0.003493727 | 5 |
| GO:0002449 | lymphocyte mediated immunity | 0.006112681 | 5 |
| GO:0002460 | adaptive immune response based on somatic recombination of im | 0.006409837 | 5 |
| GO:0002443 | leukocyte mediated immunity | 0.014258444 | 5 |
| hsa04640 | Hematopoietic cell lineage | 1.62116E-05 | 5 |
| GO:0071723 | lipopeptide binding | 1.68833E-07 | 4 |
| GO:0003823 | antigen binding | 0.007684227 | 4 |
| GO:0042277 | peptide binding | 0.020045833 | 4 |
| GO:0002475 | antigen processing and presentation via MHC class Ib | 3.95208E-06 | 4 |
| GO:0019883 | antigen processing and presentation of endogenous antigen | 2.91101E-05 | 4 |
| GO:0001916 | positive regulation of T cell mediated cytotoxicity | 4.06121E-05 | 4 |
| GO:0001914 | regulation of T cell mediated cytotoxicity | 9.69179E-05 | 4 |
| GO:0019884 | antigen processing and presentation of exogenous antigen | 0.000114814 | 4 |
| GO:0001913 | T cell mediated cytotoxicity | 0.000118221 | 4 |
| GO:0001912 | positive regulation of leukocyte mediated cytotoxicity | 0.00018266 | 4 |
| GO:0002711 | positive regulation of T cell mediated immunity | 0.00018266 | 4 |
| GO:0031343 | positive regulation of cell killing | 0.000243749 | 4 |
| GO:0001910 | regulation of leukocyte mediated cytotoxicity | 0.000631977 | 4 |
| GO:0002709 | regulation of T cell mediated immunity | 0.000631977 | 4 |
| GO:0032392 | DNA geometric change | 0.000899057 | 4 |
| GO:0031341 | regulation of cell killing | 0.000948248 | 4 |
| GO:0071103 | DNA conformation change | 0.001038523 | 4 |
| GO:0002824 | positive regulation of adaptive immune response based on somat | 0.001305509 | 4 |
| GO:0002708 | positive regulation of lymphocyte mediated immunity | 0.001436534 | 4 |
| GO:0002821 | positive regulation of adaptive immune response | 0.001436534 | 4 |
| GO:0001909 | leukocyte mediated cytotoxicity | 0.002008999 | 4 |
| GO:0009142 | nucleoside triphosphate biosynthetic process | 0.002008999 | 4 |
| GO:0002705 | positive regulation of leukocyte mediated immunity | 0.002278439 | 4 |
| GO:0002706 | regulation of lymphocyte mediated immunity | 0.005007995 | 4 |
| GO:0002822 | regulation of adaptive immune response based on somatic recoml | 0.005146329 | 4 |
| GO:0002819 | regulation of adaptive immune response | 0.006112681 | 4 |
| GO:0002703 | regulation of leukocyte mediated immunity | 0.01098832 | 4 |
| GO:0002699 | positive regulation of immune effector process | 0.014062636 | 4 |
| GO:0009141 | nucleoside triphosphate metabolic process | 0.014369605 | 4 |
| GO:0006310 | DNA recombination | 0.026761061 | 4 |
| GO:0048545 | response to steroid hormone | 0.028314645 | 4 |
| GO:0002697 | regulation of immune effector process | 0.038627622 | 4 |
| hsa05146 | Amoebiasis | 0.000372008 | 4 |
| hsa04530 | Tight junction | 0.001826961 | 4 |
| hsa05208 | Chemical carcinogenesis - reactive oxygen species | 0.003872815 | 4 |
| hsa05014 | Amyotrophic lateral sclerosis | 0.015990351 | 4 |
| GO:0016655 | oxidoreductase activity, acting on NAD(P)H, quinone or similar co | 0.007617294 | 3 |
| GO:0016651 | oxidoreductase activity, acting on NAD(P)H | 0.009495596 | 3 |
| GO:0003697 | single-stranded DNA binding | 0.015376845 | 3 |
| GO:0016705 | oxidoreductase activity, acting on paired donors, with incorporati | 0.023787014 | 3 |
| GO:0140097 | catalytic activity, acting on DNA | 0.046705964 | 3 |
| GO:0042776 | proton motive force-driven mitochondrial ATP synthesis | 0.004625943 | 3 |
| GO:0015986 | proton motive force-driven ATP synthesis | 0.006025796 | 3 |
| GO:0033108 | mitochondrial respiratory chain complex assembly | 0.010272194 | 3 |
| GO:0006754 | ATP biosynthetic process | 0.011893982 | 3 |
| GO:0009206 | purine ribonucleoside triphosphate biosynthetic process | 0.014369605 | 3 |
| GO:0009145 | purine nucleoside triphosphate biosynthetic process | 0.014370917 | 3 |
| GO:0009201 | ribonucleoside triphosphate biosynthetic process | 0.015192824 | 3 |
| GO:0006119 | oxidative phosphorylation | 0.023835168 | 3 |
| GO:0009060 | aerobic respiration | 0.045063028 | 3 |
| GO:0071897 | DNA biosynthetic process | 0.046494088 | 3 |
| hsa00190 | Oxidative phosphorylation | 0.009456751 | 3 |
| hsa05415 | Diabetic cardiomyopathy | 0.019255907 | 3 |
| hsa04714 | Thermogenesis | 0.024826512 | 3 |
| hsa05012 | Parkinson disease | 0.028821996 | 3 |
| hsa05020 | Prion disease | 0.028821996 | 3 |
| hsa05016 | Huntington disease | 0.03659053 | 3 |

| | | | |
|---|---|---|---|
| HALLMARK_E2F_TARGETS | HALLMARK_E2F_TARGETS | 0.03536907 | 3 |
| GO:0000400 | four-way junction DNA binding | 0.009495596 | 2 |
| GO:0008301 | DNA binding, bending | 0.009495596 | 2 |
| GO:0032452 | histone demethylase activity | 0.015376845 | 2 |
| GO:0140457 | protein demethylase activity | 0.015376845 | 2 |
| GO:0000217 | DNA secondary structure binding | 0.020045833 | 2 |
| GO:0032451 | demethylase activity | 0.020045833 | 2 |
| GO:0008137 | NADH dehydrogenase (ubiquinone) activity | 0.020045833 | 2 |
| GO:0050136 | NADH dehydrogenase (quinone) activity | 0.020045833 | 2 |
| GO:0003954 | NADH dehydrogenase activity | 0.02027603 | 2 |
| GO:0003955 | NAD(P)H dehydrogenase (quinone) activity | 0.02027603 | 2 |
| GO:0016706 | 2-oxoglutarate-dependent dioxygenase activity | 0.031948002 | 2 |
| GO:0015453 | oxidoreduction-driven active transmembrane transporter activity | 0.042141123 | 2 |
| GO:0009263 | deoxyribonucleotide biosynthetic process | 0.006112681 | 2 |
| GO:0009265 | 2'-deoxyribonucleotide biosynthetic process | 0.006112681 | 2 |
| GO:0046385 | deoxyribose phosphate biosynthetic process | 0.006112681 | 2 |
| GO:0033151 | V(D)J recombination | 0.007345656 | 2 |
| GO:0070076 | histone lysine demethylation | 0.014370917 | 2 |
| GO:0016577 | histone demethylation | 0.015084467 | 2 |
| GO:0006482 | protein demethylation | 0.017247149 | 2 |
| GO:0008214 | protein dealkylation | 0.017247149 | 2 |
| GO:0009394 | 2'-deoxyribonucleotide metabolic process | 0.030042109 | 2 |
| GO:0009262 | deoxyribonucleotide metabolic process | 0.030265292 | 2 |
| GO:0019692 | deoxyribose phosphate metabolic process | 0.030265292 | 2 |
| GO:0006120 | mitochondrial electron transport, NADH to ubiquinone | 0.038048596 | 2 |
| GO:0010257 | NADH dehydrogenase complex assembly | 0.046494088 | 2 |
| GO:0032981 | mitochondrial respiratory chain complex I assembly | 0.046494088 | 2 |
| hsa00240 | Pyrimidine metabolism | 0.019255907 | 2 |
| hsa01232 | Nucleotide metabolism | 0.028821996 | 2 |

| ID | Description | p.adjust | Count |
|---|---|---|---|
| HALLMARK_IL2_STAT5_SIGNALING | HALLMARK_IL2_STAT5_SIGNALING | 2.25698E-06 | 9 |
| hsa04060 | Cytokine-cytokine receptor interaction | 8.14108E-05 | 8 |
| GO:0019955 | cytokine binding | 6.64348E-06 | 7 |
| GO:0004896 | cytokine receptor activity | 8.04388E-06 | 6 |
| GO:0140375 | immune receptor activity | 6.54644E-05 | 6 |
| HALLMARK_IL6_JAK_STAT3_SIGNALING | HALLMARK_IL6_JAK_STAT3_SIGNALING | 0.0003914 | 5 |
| HALLMARK_INFLAMMATORY_RESPONSE | HALLMARK_INFLAMMATORY_RESPONSE | 0.01262365 | 5 |
| GO:0019207 | kinase regulator activity | 0.032554655 | 4 |
| hsa04630 | JAK-STAT signaling pathway | 0.041706591 | 4 |
| GO:0019838 | growth factor binding | 0.040735445 | 3 |
| hsa04640 | Hematopoietic cell lineage | 0.049143601 | 3 |
| hsa04061 | Viral protein interaction with cytokine and cytokine receptor | 0.049143601 | 3 |
| GO:0008330 | protein tyrosine/threonine phosphatase activity | 0.010047172 | 2 |
| GO:0017017 | MAP kinase tyrosine/serine/threonine phosphatase activity | 0.013870264 | 2 |
| GO:0046935 | 1-phosphatidylinositol-3-kinase regulator activity | 0.017703526 | 2 |
| GO:0033549 | MAP kinase phosphatase activity | 0.018836769 | 2 |
| GO:0035014 | phosphatidylinositol 3-kinase regulator activity | 0.018836769 | 2 |
| GO:0016493 | C-C chemokine receptor activity | 0.023621286 | 2 |
| GO:0019957 | C-C chemokine binding | 0.023621286 | 2 |
| GO:0001637 | G protein-coupled chemoattractant receptor activity | 0.023621286 | 2 |
| GO:0004950 | chemokine receptor activity | 0.023621286 | 2 |
| GO:0019956 | chemokine binding | 0.032554655 | 2 |
| GO:0048487 | beta-tubulin binding | 0.041647821 | 2 |
| GO:0008138 | protein tyrosine/serine/threonine phosphatase activity | 0.047260306 | 2 |

| ID | Description | p.adjust | Count |
|---|---|---|---|
| GO:0002449 | lymphocyte mediated immunity | 5.3408E-05 | 9 |
| GO:0002443 | leukocyte mediated immunity | 0.000160762 | 9 |
| GO:0002456 | T cell mediated immunity | 3.92346E-07 | 8 |
| GO:0002460 | adaptive immune response based on somatic recombination of im | 0.000283477 | 8 |
| GO:0002697 | regulation of immune effector process | 0.001545632 | 7 |
| GO:0002709 | regulation of T cell mediated immunity | 2.98625E-05 | 6 |
| GO:0019882 | antigen processing and presentation | 5.3408E-05 | 6 |
| GO:0002706 | regulation of lymphocyte mediated immunity | 0.000384915 | 6 |
| GO:0002822 | regulation of adaptive immune response based on somatic recoml | 0.000398594 | 6 |
| GO:0001906 | cell killing | 0.00044285 | 6 |
| GO:0002819 | regulation of adaptive immune response | 0.000499024 | 6 |
| GO:0002703 | regulation of leukocyte mediated immunity | 0.001151931 | 6 |
| GO:0002699 | positive regulation of immune effector process | 0.001683129 | 6 |
| GO:0030217 | T cell differentiation | 0.003173452 | 6 |
| GO:0006310 | DNA recombination | 0.004935418 | 6 |
| GO:0030098 | lymphocyte differentiation | 0.012831049 | 6 |
| GO:1903131 | mononuclear cell differentiation | 0.022032873 | 6 |
| hsa04640 | Hematopoietic cell lineage | 5.90483E-05 | 6 |
| HALLMARK_E2F_TARGETS | HALLMARK_E2F_TARGETS | 0.002345809 | 6 |
| GO:0002711 | positive regulation of T cell mediated immunity | 6.4197E-05 | 5 |
| GO:0002562 | somatic diversification of immune receptors via germline recombi | 0.000104968 | 5 |
| GO:0016444 | somatic cell DNA recombination | 0.000104968 | 5 |
| GO:0002200 | somatic diversification of immune receptors | 0.000160762 | 5 |
| GO:0002824 | positive regulation of adaptive immune response based on somat | 0.000499024 | 5 |
| GO:0002708 | positive regulation of lymphocyte mediated immunity | 0.000584647 | 5 |
| GO:0002821 | positive regulation of adaptive immune response | 0.000584647 | 5 |
| GO:0001909 | leukocyte mediated cytotoxicity | 0.000921221 | 5 |
| GO:0002705 | positive regulation of leukocyte mediated immunity | 0.001056553 | 5 |
| GO:0002285 | lymphocyte activation involved in immune response | 0.004935418 | 5 |
| GO:0033044 | regulation of chromosome organization | 0.00920241 | 5 |
| GO:0002366 | leukocyte activation involved in immune response | 0.016902491 | 5 |
| GO:0002263 | cell activation involved in immune response | 0.017642578 | 5 |
| GO:0098813 | nuclear chromosome segregation | 0.023291388 | 5 |
| GO:0006338 | chromatin remodeling | 0.034194813 | 5 |
| GO:0050863 | regulation of T cell activation | 0.034502936 | 5 |
| GO:1903037 | regulation of leukocyte cell-cell adhesion | 0.034502936 | 5 |
| GO:0007059 | chromosome segregation | 0.036086812 | 5 |
| GO:1903706 | regulation of hemopoiesis | 0.046486926 | 5 |
| GO:0007159 | leukocyte cell-cell adhesion | 0.046486926 | 5 |
| hsa04530 | Tight junction | 0.006763629 | 5 |
| GO:0071723 | lipopeptide binding | 1.51314E-06 | 4 |
| GO:0003697 | single-stranded DNA binding | 0.020290723 | 4 |
| GO:0003823 | antigen binding | 0.037230014 | 4 |
| GO:0002475 | antigen processing and presentation via MHC class Ib | 1.96804E-05 | 4 |
| GO:0033151 | V(D)J recombination | 2.98625E-05 | 4 |
| GO:0019883 | antigen processing and presentation of endogenous antigen | 7.18936E-05 | 4 |
| GO:0001916 | positive regulation of T cell mediated cytotoxicity | 0.000109049 | 4 |
| GO:0001914 | regulation of T cell mediated cytotoxicity | 0.000270791 | 4 |
| GO:0019884 | antigen processing and presentation of exogenous antigen | 0.00043428 | 4 |
| GO:0001913 | T cell mediated cytotoxicity | 0.000459321 | 4 |
| GO:0001912 | positive regulation of leukocyte mediated cytotoxicity | 0.000659558 | 4 |
| GO:0031343 | positive regulation of cell killing | 0.000954852 | 4 |
| GO:0001910 | regulation of leukocyte mediated cytotoxicity | 0.002395038 | 4 |
| GO:0010965 | regulation of mitotic sister chromatid separation | 0.003585771 | 4 |
| GO:0051306 | mitotic sister chromatid separation | 0.003915773 | 4 |
| GO:0031341 | regulation of cell killing | 0.0039552 | 4 |
| GO:0033045 | regulation of sister chromatid segregation | 0.004631079 | 4 |
| GO:1905818 | regulation of chromosome separation | 0.004935418 | 4 |
| GO:0051983 | regulation of chromosome segregation | 0.008334697 | 4 |
| GO:0051304 | chromosome separation | 0.00888531 | 4 |
| GO:0045580 | regulation of T cell differentiation | 0.021142871 | 4 |
| GO:0002833 | positive regulation of response to biotic stimulus | 0.023689661 | 4 |
| GO:0000070 | mitotic sister chromatid segregation | 0.029563941 | 4 |
| GO:0045619 | regulation of lymphocyte differentiation | 0.030989995 | 4 |
| GO:0000819 | sister chromatid segregation | 0.041258798 | 4 |
| GO:0050870 | positive regulation of T cell activation | 0.046411527 | 4 |
| hsa05146 | Amoebiasis | 0.006763629 | 4 |

| | | | |
|---|---|---|---|
| hsa04110 | Cell cycle | 0.027280418 | 4 |
| GO:0033046 | negative regulation of sister chromatid segregation | 0.007661266 | 3 |
| GO:0033048 | negative regulation of mitotic sister chromatid segregation | 0.007661266 | 3 |
| GO:2000816 | negative regulation of mitotic sister chromatid separation | 0.007661266 | 3 |
| GO:0051985 | negative regulation of chromosome segregation | 0.008206487 | 3 |
| GO:1905819 | negative regulation of chromosome separation | 0.008206487 | 3 |
| GO:0033047 | regulation of mitotic sister chromatid segregation | 0.0090564 | 3 |
| GO:0032481 | positive regulation of type I interferon production | 0.01183099 | 3 |
| GO:0006303 | double-strand break repair via nonhomologous end joining | 0.013648318 | 3 |
| GO:0030071 | regulation of mitotic metaphase/anaphase transition | 0.028460084 | 3 |
| GO:0007091 | metaphase/anaphase transition of mitotic cell cycle | 0.029829247 | 3 |
| GO:1902099 | regulation of metaphase/anaphase transition of cell cycle | 0.029829247 | 3 |
| GO:2001251 | negative regulation of chromosome organization | 0.030989995 | 3 |
| GO:0044784 | metaphase/anaphase transition of cell cycle | 0.031229643 | 3 |
| GO:0032392 | DNA geometric change | 0.031703227 | 3 |
| GO:1901992 | positive regulation of mitotic cell cycle phase transition | 0.032070185 | 3 |
| GO:0045132 | meiotic chromosome segregation | 0.032070185 | 3 |
| GO:0032479 | regulation of type I interferon production | 0.034194813 | 3 |
| GO:0032606 | type I interferon production | 0.034194813 | 3 |
| GO:0071103 | DNA conformation change | 0.034502936 | 3 |
| GO:0002286 | T cell activation involved in immune response | 0.043704697 | 3 |
| GO:0007088 | regulation of mitotic nuclear division | 0.043704697 | 3 |
| GO:1901989 | positive regulation of cell cycle phase transition | 0.046411527 | 3 |
| hsa05340 | Primary immunodeficiency | 0.006763629 | 3 |
| GO:0050786 | RAGE receptor binding | 0.020290723 | 2 |
| GO:0000400 | four-way junction DNA binding | 0.037230014 | 2 |
| GO:0008301 | DNA binding, bending | 0.038017018 | 2 |
| GO:0042288 | MHC class I protein binding | 0.038338287 | 2 |
| GO:0070182 | DNA polymerase binding | 0.038338287 | 2 |
| GO:0002327 | immature B cell differentiation | 0.00920241 | 2 |
| GO:0002424 | T cell mediated immune response to tumor cell | 0.00920241 | 2 |
| GO:0002837 | regulation of immune response to tumor cell | 0.022611135 | 2 |
| GO:0002834 | regulation of response to tumor cell | 0.023744361 | 2 |
| GO:0032069 | regulation of nuclease activity | 0.025663539 | 2 |
| GO:0051131 | chaperone-mediated protein complex assembly | 0.027624728 | 2 |
| GO:0002418 | immune response to tumor cell | 0.032070185 | 2 |
| GO:0042026 | protein refolding | 0.032070185 | 2 |
| GO:0010390 | histone monoubiquitination | 0.046411527 | 2 |

| ID | Description | p.adjust | Count |
|---|---|---|---|
| GO:0000280 | nuclear division | 9.47239E-07 | 12 |
| HALLMARK_G2M_CHECKPOINT | HALLMARK_G2M_CHECKPOINT | 1.18656E-06 | 11 |
| GO:0044772 | mitotic cell cycle phase transition | 3.49547E-05 | 10 |
| GO:0140014 | mitotic nuclear division | 2.21208E-05 | 9 |
| GO:0007059 | chromosome segregation | 5.23532E-05 | 9 |
| GO:1901987 | regulation of cell cycle phase transition | 0.000121073 | 9 |
| HALLMARK_E2F_TARGETS | HALLMARK_E2F_TARGETS | 6.92847E-05 | 9 |
| GO:0007051 | spindle organization | 9.47999E-06 | 8 |
| GO:1901990 | regulation of mitotic cell cycle phase transition | 0.000150623 | 8 |
| GO:0045786 | negative regulation of cell cycle | 0.000203681 | 8 |
| GO:0000070 | mitotic sister chromatid segregation | 7.734E-05 | 7 |
| GO:0000819 | sister chromatid segregation | 0.000142328 | 7 |
| GO:1901988 | negative regulation of cell cycle phase transition | 0.000202417 | 7 |
| GO:0010948 | negative regulation of cell cycle process | 0.000259548 | 7 |
| GO:0098813 | nuclear chromosome segregation | 0.000266401 | 7 |
| GO:0045787 | positive regulation of cell cycle | 0.000505089 | 7 |
| HALLMARK_MITOTIC_SPINDLE | HALLMARK_MITOTIC_SPINDLE | 0.002844567 | 7 |
| GO:0015631 | tubulin binding | 0.008558911 | 6 |
| GO:0007088 | regulation of mitotic nuclear division | 6.36135E-05 | 6 |
| GO:0007093 | mitotic cell cycle checkpoint signaling | 0.000121073 | 6 |
| GO:0051783 | regulation of nuclear division | 0.000128355 | 6 |
| GO:1902850 | microtubule cytoskeleton organization involved in mitosis | 0.00019526 | 6 |
| GO:0000075 | cell cycle checkpoint signaling | 0.000205382 | 6 |
| GO:0001906 | cell killing | 0.000205778 | 6 |
| GO:0045930 | negative regulation of mitotic cell cycle | 0.000556702 | 6 |
| GO:0090068 | positive regulation of cell cycle process | 0.000743978 | 6 |
| GO:0140694 | non-membrane-bounded organelle assembly | 0.004820445 | 6 |
| hsa04110 | Cell cycle | 0.000959944 | 6 |
| hsa04814 | Motor proteins | 0.001534856 | 6 |
| hsa05132 | Salmonella infection | 0.004146227 | 6 |
| hsa05012 | Parkinson disease | 0.00499911 | 6 |
| hsa05020 | Prion disease | 0.00499911 | 6 |
| hsa05016 | Huntington disease | 0.007459335 | 6 |
| hsa05014 | Amyotrophic lateral sclerosis | 0.014060953 | 6 |
| hsa05010 | Alzheimer disease | 0.014358201 | 6 |
| hsa05022 | Pathways of neurodegeneration - multiple diseases | 0.038134092 | 6 |
| GO:0005200 | structural constituent of cytoskeleton | 0.000517532 | 5 |
| GO:0019887 | protein kinase regulator activity | 0.008558911 | 5 |
| GO:0019207 | kinase regulator activity | 0.008558911 | 5 |
| GO:0045296 | cadherin binding | 0.019755871 | 5 |
| GO:0005525 | GTP binding | 0.027200561 | 5 |
| GO:0019001 | guanyl nucleotide binding | 0.030056758 | 5 |
| GO:0032561 | guanyl ribonucleotide binding | 0.030056758 | 5 |
| GO:0030071 | regulation of mitotic metaphase/anaphase transition | 0.000201045 | 5 |
| GO:0007091 | metaphase/anaphase transition of mitotic cell cycle | 0.000202417 | 5 |
| GO:1902099 | regulation of metaphase/anaphase transition of cell cycle | 0.000202417 | 5 |
| GO:0010965 | regulation of mitotic sister chromatid separation | 0.000202417 | 5 |
| GO:0044784 | metaphase/anaphase transition of cell cycle | 0.000202417 | 5 |
| GO:0051306 | mitotic sister chromatid separation | 0.000202902 | 5 |
| GO:0033045 | regulation of sister chromatid segregation | 0.000205778 | 5 |
| GO:1905818 | regulation of chromosome separation | 0.000208636 | 5 |
| GO:1904029 | regulation of cyclin-dependent protein kinase activity | 0.000259548 | 5 |
| GO:0051225 | spindle assembly | 0.000338346 | 5 |
| GO:0051983 | regulation of chromosome segregation | 0.000375905 | 5 |
| GO:0007052 | mitotic spindle organization | 0.000380261 | 5 |
| GO:0051304 | chromosome separation | 0.000399862 | 5 |
| GO:0009060 | aerobic respiration | 0.001924873 | 5 |
| GO:0140013 | meiotic nuclear division | 0.001924873 | 5 |
| GO:1901991 | negative regulation of mitotic cell cycle phase transition | 0.001924873 | 5 |
| GO:1903046 | meiotic cell cycle process | 0.00286585 | 5 |
| GO:0045333 | cellular respiration | 0.004547534 | 5 |
| GO:0033044 | regulation of chromosome organization | 0.005242614 | 5 |
| GO:0051321 | meiotic cell cycle | 0.008800774 | 5 |
| GO:0009165 | nucleotide biosynthetic process | 0.010306364 | 5 |
| GO:1901293 | nucleoside phosphate biosynthetic process | 0.010483704 | 5 |
| GO:0019058 | viral life cycle | 0.012038501 | 5 |
| GO:0015980 | energy derivation by oxidation of organic compounds | 0.013923409 | 5 |

| | | | |
|---|---|---|---|
| GO:0010639 | negative regulation of organelle organization | 0.019765787 | 5 |
| GO:0071900 | regulation of protein serine/threonine kinase activity | 0.023408953 | 5 |
| GO:0016032 | viral process | 0.026746746 | 5 |
| GO:0009150 | purine ribonucleotide metabolic process | 0.032946245 | 5 |
| GO:0009259 | ribonucleotide metabolic process | 0.037924771 | 5 |
| GO:0019693 | ribose phosphate metabolic process | 0.040515004 | 5 |
| GO:0006163 | purine nucleotide metabolic process | 0.040587449 | 5 |
| GO:0016049 | cell growth | 0.045396526 | 5 |
| hsa04540 | Gap junction | 0.000959944 | 5 |
| hsa04914 | Progesterone-mediated oocyte maturation | 0.001030301 | 5 |
| hsa05130 | Pathogenic Escherichia coli infection | 0.007459335 | 5 |
| GO:0016538 | cyclin-dependent protein serine/threonine kinase regulator activi | 0.000517532 | 4 |
| GO:0016829 | lyase activity | 0.019755871 | 4 |
| GO:0008017 | microtubule binding | 0.03890715 | 4 |
| GO:0007094 | mitotic spindle assembly checkpoint signaling | 0.000202902 | 4 |
| GO:0071173 | spindle assembly checkpoint signaling | 0.000202902 | 4 |
| GO:0071174 | mitotic spindle checkpoint signaling | 0.000202902 | 4 |
| GO:0031577 | spindle checkpoint signaling | 0.000203681 | 4 |
| GO:0045841 | negative regulation of mitotic metaphase/anaphase transition | 0.000205382 | 4 |
| GO:1902100 | negative regulation of metaphase/anaphase transition of cell cycl | 0.000208636 | 4 |
| GO:0033046 | negative regulation of sister chromatid segregation | 0.000208636 | 4 |
| GO:0033048 | negative regulation of mitotic sister chromatid segregation | 0.000208636 | 4 |
| GO:2000816 | negative regulation of mitotic sister chromatid separation | 0.000208636 | 4 |
| GO:0051985 | negative regulation of chromosome segregation | 0.000229809 | 4 |
| GO:1905819 | negative regulation of chromosome separation | 0.000229809 | 4 |
| GO:0033047 | regulation of mitotic sister chromatid segregation | 0.000259548 | 4 |
| GO:0045839 | negative regulation of mitotic nuclear division | 0.000259548 | 4 |
| GO:0051784 | negative regulation of nuclear division | 0.000375905 | 4 |
| GO:2001251 | negative regulation of chromosome organization | 0.001814055 | 4 |
| GO:1901992 | positive regulation of mitotic cell cycle phase transition | 0.001924873 | 4 |
| GO:0009185 | ribonucleoside diphosphate metabolic process | 0.003028845 | 4 |
| GO:0000079 | regulation of cyclin-dependent protein serine/threonine kinase ac | 0.003288948 | 4 |
| GO:1901989 | positive regulation of cell cycle phase transition | 0.003796401 | 4 |
| GO:0045931 | positive regulation of mitotic cell cycle | 0.004544523 | 4 |
| GO:0009132 | nucleoside diphosphate metabolic process | 0.004725514 | 4 |
| GO:0001909 | leukocyte mediated cytotoxicity | 0.004820445 | 4 |
| GO:0006119 | oxidative phosphorylation | 0.006496726 | 4 |
| GO:0046034 | ATP metabolic process | 0.019765787 | 4 |
| GO:0009152 | purine ribonucleotide biosynthetic process | 0.019765787 | 4 |
| GO:0009260 | ribonucleotide biosynthetic process | 0.022659641 | 4 |
| GO:0046390 | ribose phosphate biosynthetic process | 0.024234362 | 4 |
| GO:0006164 | purine nucleotide biosynthetic process | 0.024234362 | 4 |
| GO:0009205 | purine ribonucleoside triphosphate metabolic process | 0.024234362 | 4 |
| GO:0009144 | purine nucleoside triphosphate metabolic process | 0.024720254 | 4 |
| GO:0009199 | ribonucleoside triphosphate metabolic process | 0.025179702 | 4 |
| GO:0072522 | purine-containing compound biosynthetic process | 0.025708416 | 4 |
| GO:0009141 | nucleoside triphosphate metabolic process | 0.030381214 | 4 |
| hsa04115 | p53 signaling pathway | 0.002811003 | 4 |
| hsa04114 | Oocyte meiosis | 0.011211539 | 4 |
| hsa04145 | Phagosome | 0.014358201 | 4 |
| hsa04218 | Cellular senescence | 0.014358201 | 4 |
| GO:0003725 | double-stranded RNA binding | 0.016204138 | 3 |
| GO:0030261 | chromosome condensation | 0.004003809 | 3 |
| GO:0006968 | cellular defense response | 0.004547534 | 3 |
| GO:1904031 | positive regulation of cyclin-dependent protein kinase activity | 0.004820445 | 3 |
| GO:0042267 | natural killer cell mediated cytotoxicity | 0.010239141 | 3 |
| GO:0002228 | natural killer cell mediated immunity | 0.010918011 | 3 |
| GO:0006096 | glycolytic process | 0.012946353 | 3 |
| GO:0006757 | ATP generation from ADP | 0.013249535 | 3 |
| GO:0000281 | mitotic cytokinesis | 0.013555703 | 3 |
| GO:0046031 | ADP metabolic process | 0.016657796 | 3 |
| GO:0006165 | nucleoside diphosphate phosphorylation | 0.019765787 | 3 |
| GO:0042773 | ATP synthesis coupled electron transport | 0.019765787 | 3 |
| GO:0042775 | mitochondrial ATP synthesis coupled electron transport | 0.019765787 | 3 |
| GO:2001243 | negative regulation of intrinsic apoptotic signaling pathway | 0.019765787 | 3 |
| GO:0031341 | regulation of cell killing | 0.019765787 | 3 |
| GO:0046939 | nucleotide phosphorylation | 0.019765787 | 3 |
| GO:0048144 | fibroblast proliferation | 0.02012774 | 3 |

| | | | |
|---|---|---|---|
| GO:0009135 | purine nucleoside diphosphate metabolic process | 0.02029718 | 3 |
| GO:0009179 | purine ribonucleoside diphosphate metabolic process | 0.02029718 | 3 |
| GO:0006754 | ATP biosynthetic process | 0.020836308 | 3 |
| GO:0006090 | pyruvate metabolic process | 0.02157738 | 3 |
| GO:0032526 | response to retinoic acid | 0.02157738 | 3 |
| GO:0061640 | cytoskeleton-dependent cytokinesis | 0.023752059 | 3 |
| GO:0009206 | purine ribonucleoside triphosphate biosynthetic process | 0.024720254 | 3 |
| GO:0009145 | purine nucleoside triphosphate biosynthetic process | 0.024720254 | 3 |
| GO:0022904 | respiratory electron transport chain | 0.026408315 | 3 |
| GO:0009201 | ribonucleoside triphosphate biosynthetic process | 0.026746746 | 3 |
| GO:0009142 | nucleoside triphosphate biosynthetic process | 0.032585866 | 3 |
| GO:1903900 | regulation of viral life cycle | 0.037149875 | 3 |
| GO:0016052 | carbohydrate catabolic process | 0.044542508 | 3 |
| GO:0044839 | cell cycle G2/M phase transition | 0.045528702 | 3 |
| GO:0050792 | regulation of viral process | 0.048339775 | 3 |
| hsa00010 | Glycolysis / Gluconeogenesis | 0.014060953 | 3 |
| hsa01230 | Biosynthesis of amino acids | 0.014358201 | 3 |
| hsa01200 | Carbon metabolism | 0.042470302 | 3 |
| GO:0008199 | ferric iron binding | 0.008558911 | 2 |
| GO:0061575 | cyclin-dependent protein serine/threonine kinase activator activi | 0.009607371 | 2 |
| GO:0042288 | MHC class I protein binding | 0.019755871 | 2 |
| GO:0023026 | MHC class II protein complex binding | 0.02466471 | 2 |
| GO:0023023 | MHC protein complex binding | 0.032732082 | 2 |
| GO:0042287 | MHC protein binding | 0.039387404 | 2 |
| GO:0070601 | centromeric sister chromatid cohesion | 0.0058167 | 2 |
| GO:0000212 | meiotic spindle organization | 0.008822636 | 2 |
| GO:0006735 | NADH regeneration | 0.009405398 | 2 |
| GO:0045144 | meiotic sister chromatid segregation | 0.009405398 | 2 |
| GO:0051177 | meiotic sister chromatid cohesion | 0.009405398 | 2 |
| GO:0061621 | canonical glycolysis | 0.009405398 | 2 |
| GO:0061718 | glucose catabolic process to pyruvate | 0.009405398 | 2 |
| GO:0007135 | meiosis II | 0.010239141 | 2 |
| GO:0061983 | meiosis II cell cycle process | 0.010239141 | 2 |
| GO:0061620 | glycolytic process through glucose-6-phosphate | 0.010947309 | 2 |
| GO:0061615 | glycolytic process through fructose-6-phosphate | 0.012038501 | 2 |
| GO:1903902 | positive regulation of viral life cycle | 0.017849156 | 2 |
| GO:0006007 | glucose catabolic process | 0.018955328 | 2 |
| GO:0019835 | cytolysis | 0.018955328 | 2 |
| GO:0010971 | positive regulation of G2/M transition of mitotic cell cycle | 0.020664774 | 2 |
| GO:0031342 | negative regulation of cell killing | 0.020664774 | 2 |
| GO:1902751 | positive regulation of cell cycle G2/M phase transition | 0.023752059 | 2 |
| GO:0034508 | centromere complex assembly | 0.024234362 | 2 |
| GO:0051642 | centrosome localization | 0.024234362 | 2 |
| GO:0061842 | microtubule organizing center localization | 0.024720254 | 2 |
| GO:0071353 | cellular response to interleukin-4 | 0.024720254 | 2 |
| GO:0070670 | response to interleukin-4 | 0.028007216 | 2 |
| GO:0006734 | NADH metabolic process | 0.029115032 | 2 |
| GO:0032435 | negative regulation of proteasomal ubiquitin-dependent protein ( | 0.029115032 | 2 |
| GO:0042307 | positive regulation of protein import into nucleus | 0.032946245 | 2 |
| GO:0019320 | hexose catabolic process | 0.035993342 | 2 |
| GO:0071459 | protein localization to chromosome, centromeric region | 0.037149875 | 2 |
| GO:0045840 | positive regulation of mitotic nuclear division | 0.038309048 | 2 |
| GO:0031640 | killing of cells of another organism | 0.040634213 | 2 |
| GO:0046365 | monosaccharide catabolic process | 0.040634213 | 2 |
| GO:1901976 | regulation of cell cycle checkpoint | 0.043849668 | 2 |
| GO:0045737 | positive regulation of cyclin-dependent protein serine/threonine l | 0.045905021 | 2 |
| GO:2000059 | negative regulation of ubiquitin-dependent protein catabolic proc | 0.045905021 | 2 |
| GO:1901799 | negative regulation of proteasomal protein catabolic process | 0.0473929 | 2 |

| ID | Description | p.adjust | Count |
|---|---|---|---|
| GO:0019221 | cytokine-mediated signaling pathway | 0.001808208 | 9 |
| HALLMARK_INFLAMMATORY_RESPONSE | HALLMARK_INFLAMMATORY_RESPONSE | 0.000108737 | 9 |
| GO:0002237 | response to molecule of bacterial origin | 0.001808208 | 8 |
| hsa04060 | Cytokine-cytokine receptor interaction | 0.001109207 | 8 |
| HALLMARK_IL2_STAT5_SIGNALING | HALLMARK_IL2_STAT5_SIGNALING | 0.000450182 | 8 |
| HALLMARK_ALLOGRAFT_REJECTION | HALLMARK_ALLOGRAFT_REJECTION | 0.000450182 | 8 |
| GO:0032496 | response to lipopolysaccharide | 0.007417435 | 7 |
| GO:0050863 | regulation of T cell activation | 0.009078017 | 7 |
| GO:0002443 | leukocyte mediated immunity | 0.015282311 | 7 |
| GO:0001819 | positive regulation of cytokine production | 0.016946213 | 7 |
| GO:0022407 | regulation of cell-cell adhesion | 0.016946213 | 7 |
| HALLMARK_IL6_JAK_STAT3_SIGNALING | HALLMARK_IL6_JAK_STAT3_SIGNALING | 5.78474E-05 | 7 |
| HALLMARK_INTERFERON_GAMMA_RESPONSE | HALLMARK_INTERFERON_GAMMA_RESPONSE | 0.002119744 | 7 |
| HALLMARK_TNFA_SIGNALING_VIA_NFKB | HALLMARK_TNFA_SIGNALING_VIA_NFKB | 0.002119744 | 7 |
| GO:0140375 | immune receptor activity | 0.000334111 | 6 |
| GO:0071219 | cellular response to molecule of bacterial origin | 0.007417435 | 6 |
| GO:0071216 | cellular response to biotic stimulus | 0.009126576 | 6 |
| GO:0070663 | regulation of leukocyte proliferation | 0.009256892 | 6 |
| GO:0048872 | homeostasis of number of cells | 0.01229906 | 6 |
| GO:0060326 | cell chemotaxis | 0.015282311 | 6 |
| GO:0070661 | leukocyte proliferation | 0.016946213 | 6 |
| GO:0002449 | lymphocyte mediated immunity | 0.018865789 | 6 |
| GO:1903037 | regulation of leukocyte cell-cell adhesion | 0.019390858 | 6 |
| GO:0007159 | leukocyte cell-cell adhesion | 0.026985675 | 6 |
| GO:0019955 | cytokine binding | 0.002760356 | 5 |
| GO:0005126 | cytokine receptor binding | 0.018346646 | 5 |
| GO:0042129 | regulation of T cell proliferation | 0.01229906 | 5 |
| GO:0042098 | T cell proliferation | 0.016946213 | 5 |
| GO:0071222 | cellular response to lipopolysaccharide | 0.016946213 | 5 |
| GO:0071356 | cellular response to tumor necrosis factor | 0.018865789 | 5 |
| GO:0050670 | regulation of lymphocyte proliferation | 0.019390858 | 5 |
| GO:0032944 | regulation of mononuclear cell proliferation | 0.019390858 | 5 |
| GO:0034612 | response to tumor necrosis factor | 0.022418352 | 5 |
| GO:0046651 | lymphocyte proliferation | 0.042585918 | 5 |
| GO:0032943 | mononuclear cell proliferation | 0.044214165 | 5 |
| GO:0022409 | positive regulation of cell-cell adhesion | 0.049252857 | 5 |
| hsa04061 | Viral protein interaction with cytokine and cytokine receptor | 0.001770656 | 5 |
| hsa04210 | Apoptosis | 0.005132874 | 5 |
| GO:0004896 | cytokine receptor activity | 0.006353395 | 4 |
| GO:0070665 | positive regulation of leukocyte proliferation | 0.039221635 | 4 |
| GO:0002262 | myeloid cell homeostasis | 0.03948811 | 4 |
| hsa04620 | Toll-like receptor signaling pathway | 0.00935444 | 4 |
| hsa04668 | TNF signaling pathway | 0.011572275 | 4 |
| hsa05162 | Measles | 0.018847486 | 4 |
| hsa04630 | JAK-STAT signaling pathway | 0.033596656 | 4 |
| hsa05164 | Influenza A | 0.034266054 | 4 |
| hsa05152 | Tuberculosis | 0.038000755 | 4 |
| hsa05202 | Transcriptional misregulation in cancer | 0.040721458 | 4 |
| hsa05130 | Pathogenic Escherichia coli infection | 0.040721458 | 4 |
| hsa05169 | Epstein-Barr virus infection | 0.041255686 | 4 |
| GO:0042379 | chemokine receptor binding | 0.020250635 | 3 |
| GO:0036230 | granulocyte activation | 0.016946213 | 3 |
| GO:0033619 | membrane protein proteolysis | 0.022418352 | 3 |
| GO:0032729 | positive regulation of interferon-gamma production | 0.042585918 | 3 |
| hsa05330 | Allograft rejection | 0.00935444 | 3 |
| hsa05332 | Graft-versus-host disease | 0.00935444 | 3 |
| hsa04940 | Type I diabetes mellitus | 0.00935444 | 3 |
| hsa05320 | Autoimmune thyroid disease | 0.012541612 | 3 |
| hsa04657 | IL-17 signaling pathway | 0.040721458 | 3 |
| hsa04640 | Hematopoietic cell lineage | 0.040721458 | 3 |
| GO:0005031 | tumor necrosis factor receptor activity | 0.01299486 | 2 |
| GO:0005035 | death receptor activity | 0.017933758 | 2 |
| GO:0045236 | CXCR chemokine receptor binding | 0.021812048 | 2 |
| GO:0050431 | transforming growth factor beta binding | 0.037596376 | 2 |

| ID | Description | p.adjust | Count |
|---|---|---|---|
| GO:0002449 | lymphocyte mediated immunity | 0.000186899 | 8 |
| GO:0051251 | positive regulation of lymphocyte activation | 0.000294545 | 8 |
| GO:0002696 | positive regulation of leukocyte activation | 0.000599498 | 8 |
| GO:0002443 | leukocyte mediated immunity | 0.000599498 | 8 |
| GO:0050867 | positive regulation of cell activation | 0.000608576 | 8 |
| hsa05014 | Amyotrophic lateral sclerosis | 0.000696318 | 8 |
| GO:0001906 | cell killing | 5.47164E-05 | 7 |
| GO:0050870 | positive regulation of T cell activation | 0.000183711 | 7 |
| GO:1903039 | positive regulation of leukocyte cell-cell adhesion | 0.000276829 | 7 |
| GO:0022409 | positive regulation of cell-cell adhesion | 0.000599498 | 7 |
| GO:0050863 | regulation of T cell activation | 0.001032492 | 7 |
| GO:1903037 | regulation of leukocyte cell-cell adhesion | 0.001032492 | 7 |
| GO:0007159 | leukocyte cell-cell adhesion | 0.001614009 | 7 |
| GO:0016032 | viral process | 0.001671819 | 7 |
| GO:0045785 | positive regulation of cell adhesion | 0.003379804 | 7 |
| GO:0022407 | regulation of cell-cell adhesion | 0.003578345 | 7 |
| hsa04145 | Phagosome | 2.60405E-05 | 7 |
| hsa05164 | Influenza A | 4.49961E-05 | 7 |
| hsa05020 | Prion disease | 0.000740387 | 7 |
| GO:0019079 | viral genome replication | 8.90944E-05 | 6 |
| GO:0046034 | ATP metabolic process | 0.000599498 | 6 |
| GO:0009205 | purine ribonucleoside triphosphate metabolic process | 0.000932029 | 6 |
| GO:0009144 | purine nucleoside triphosphate metabolic process | 0.001013054 | 6 |
| GO:0009199 | ribonucleoside triphosphate metabolic process | 0.001027916 | 6 |
| GO:0009141 | nucleoside triphosphate metabolic process | 0.001296477 | 6 |
| GO:0044403 | biological process involved in symbiotic interaction | 0.002364147 | 6 |
| GO:0019058 | viral life cycle | 0.002796834 | 6 |
| GO:0002440 | production of molecular mediator of immune response | 0.003329199 | 6 |
| GO:0002460 | adaptive immune response based on somatic recombination of im | 0.005047785 | 6 |
| GO:0009615 | response to virus | 0.007338205 | 6 |
| GO:0009150 | purine ribonucleotide metabolic process | 0.010375884 | 6 |
| GO:0009259 | ribonucleotide metabolic process | 0.012145559 | 6 |
| GO:0019693 | ribose phosphate metabolic process | 0.012709521 | 6 |
| GO:0006163 | purine nucleotide metabolic process | 0.012709521 | 6 |
| hsa05416 | Viral myocarditis | 1.02277E-05 | 6 |
| hsa04612 | Antigen processing and presentation | 1.25671E-05 | 6 |
| hsa05169 | Epstein-Barr virus infection | 0.001105344 | 6 |
| hsa04714 | Thermogenesis | 0.001854807 | 6 |
| hsa05012 | Parkinson disease | 0.003388474 | 6 |
| hsa05010 | Alzheimer disease | 0.014463157 | 6 |
| HALLMARK_ALLOGRAFT_REJECTION | HALLMARK_ALLOGRAFT_REJECTION | 0.00777901 | 6 |
| GO:0003823 | antigen binding | 0.003344023 | 5 |
| GO:0140297 | DNA-binding transcription factor binding | 0.044026151 | 5 |
| GO:0002478 | antigen processing and presentation of exogenous peptide antige | 2.16724E-05 | 5 |
| GO:0019884 | antigen processing and presentation of exogenous antigen | 2.86473E-05 | 5 |
| GO:0048002 | antigen processing and presentation of peptide antigen | 6.79591E-05 | 5 |
| GO:0045069 | regulation of viral genome replication | 0.000183711 | 5 |
| GO:0048525 | negative regulation of viral process | 0.00019761 | 5 |
| GO:0019882 | antigen processing and presentation | 0.000387087 | 5 |
| GO:0019730 | antimicrobial humoral response | 0.000599498 | 5 |
| GO:0001909 | leukocyte mediated cytotoxicity | 0.000751439 | 5 |
| GO:0034341 | response to interferon-gamma | 0.00089659 | 5 |
| GO:1903900 | regulation of viral life cycle | 0.000928083 | 5 |
| GO:0050792 | regulation of viral process | 0.001310133 | 5 |
| GO:0016064 | immunoglobulin mediated immune response | 0.003696048 | 5 |
| GO:0019724 | B cell mediated immunity | 0.003815903 | 5 |
| GO:0002377 | immunoglobulin production | 0.003815903 | 5 |
| GO:0001894 | tissue homeostasis | 0.008508093 | 5 |
| GO:0006959 | humoral immune response | 0.013521771 | 5 |
| GO:0060249 | anatomical structure homeostasis | 0.015176714 | 5 |
| GO:0010639 | negative regulation of organelle organization | 0.020188781 | 5 |
| GO:0042742 | defense response to bacterium | 0.02259141 | 5 |
| GO:1903829 | positive regulation of protein localization | 0.036563715 | 5 |
| GO:0001819 | positive regulation of cytokine production | 0.045731951 | 5 |
| GO:0006417 | regulation of translation | 0.045731951 | 5 |
| GO:0019221 | cytokine-mediated signaling pathway | 0.047779067 | 5 |
| hsa05330 | Allograft rejection | 1.25671E-05 | 5 |

| | | | |
|---|---|---|---|
| hsa05332 | Graft-versus-host disease | 1.25671E-05 | 5 |
| hsa04940 | Type I diabetes mellitus | 1.25671E-05 | 5 |
| hsa05320 | Autoimmune thyroid disease | 2.60405E-05 | 5 |
| hsa05415 | Diabetic cardiomyopathy | 0.006067382 | 5 |
| hsa05166 | Human T-cell leukemia virus 1 infection | 0.008596287 | 5 |
| hsa05016 | Huntington disease | 0.024587422 | 5 |
| HALLMARK_INTERFERON_GAMMA_RESPONSE | HALLMARK_INTERFERON_GAMMA_RESPONSE | 0.015083815 | 5 |
| HALLMARK_MYC_TARGETS_V1 | HALLMARK_MYC_TARGETS_V1 | 0.015083815 | 5 |
| GO:0023026 | MHC class II protein complex binding | 9.09413E-05 | 4 |
| GO:0023023 | MHC protein complex binding | 0.000150251 | 4 |
| GO:0051082 | unfolded protein binding | 0.005911204 | 4 |
| GO:0031625 | ubiquitin protein ligase binding | 0.044026151 | 4 |
| GO:0002399 | MHC class II protein complex assembly | 2.16724E-05 | 4 |
| GO:0002503 | peptide antigen assembly with MHC class II protein complex | 2.16724E-05 | 4 |
| GO:0002396 | MHC protein complex assembly | 2.86473E-05 | 4 |
| GO:0002501 | peptide antigen assembly with MHC protein complex | 2.86473E-05 | 4 |
| GO:0019886 | antigen processing and presentation of exogenous peptide antige | 9.55502E-05 | 4 |
| GO:0002495 | antigen processing and presentation of peptide antigen via MHC | 0.000145978 | 4 |
| GO:0002504 | antigen processing and presentation of peptide or polysaccharide | 0.000169375 | 4 |
| GO:0045071 | negative regulation of viral genome replication | 0.000599498 | 4 |
| GO:0002381 | immunoglobulin production involved in immunoglobulin-mediated | 0.001032492 | 4 |
| GO:0001895 | retina homeostasis | 0.001296477 | 4 |
| GO:0001910 | regulation of leukocyte mediated cytotoxicity | 0.001637637 | 4 |
| GO:0034109 | homotypic cell-cell adhesion | 0.002203683 | 4 |
| GO:0031341 | regulation of cell killing | 0.002782483 | 4 |
| GO:0006754 | ATP biosynthetic process | 0.003098381 | 4 |
| GO:0051702 | biological process involved in interaction with symbiont | 0.003815903 | 4 |
| GO:0009206 | purine ribonucleoside triphosphate biosynthetic process | 0.003828377 | 4 |
| GO:0009145 | purine nucleoside triphosphate biosynthetic process | 0.003828377 | 4 |
| GO:0071346 | cellular response to interferon-gamma | 0.003828377 | 4 |
| GO:0009201 | ribonucleoside triphosphate biosynthetic process | 0.004416573 | 4 |
| GO:0009142 | nucleoside triphosphate biosynthetic process | 0.005751822 | 4 |
| GO:0006119 | oxidative phosphorylation | 0.00748676 | 4 |
| GO:1902600 | proton transmembrane transport | 0.009383602 | 4 |
| GO:0099173 | postsynapse organization | 0.010375884 | 4 |
| GO:0002706 | regulation of lymphocyte mediated immunity | 0.012709521 | 4 |
| GO:0009060 | aerobic respiration | 0.015323207 | 4 |
| GO:0031396 | regulation of protein ubiquitination | 0.019146925 | 4 |
| GO:0009152 | purine ribonucleotide biosynthetic process | 0.020724412 | 4 |
| GO:0009260 | ribonucleotide biosynthetic process | 0.024446958 | 4 |
| GO:0045088 | regulation of innate immune response | 0.025720913 | 4 |
| GO:0045333 | cellular respiration | 0.025720913 | 4 |
| GO:0002703 | regulation of leukocyte mediated immunity | 0.025720913 | 4 |
| GO:0046390 | ribose phosphate biosynthetic process | 0.025720913 | 4 |
| GO:0006164 | purine nucleotide biosynthetic process | 0.025907332 | 4 |
| GO:0072522 | purine-containing compound biosynthetic process | 0.027833987 | 4 |
| GO:1903320 | regulation of protein modification by small protein conjugation or | 0.028031117 | 4 |
| GO:0051222 | positive regulation of protein transport | 0.043192848 | 4 |
| GO:0009165 | nucleotide biosynthetic process | 0.045731951 | 4 |
| GO:1901293 | nucleoside phosphate biosynthetic process | 0.046420115 | 4 |
| GO:1904951 | positive regulation of establishment of protein localization | 0.047715526 | 4 |
| GO:2001020 | regulation of response to DNA damage stimulus | 0.047950783 | 4 |
| hsa05140 | Leishmaniasis | 0.001854807 | 4 |
| hsa05323 | Rheumatoid arthritis | 0.003274981 | 4 |
| hsa05145 | Toxoplasmosis | 0.005552377 | 4 |
| hsa00190 | Oxidative phosphorylation | 0.009208775 | 4 |
| hsa04210 | Apoptosis | 0.009282817 | 4 |
| hsa04514 | Cell adhesion molecules | 0.014463157 | 4 |
| hsa05130 | Pathogenic Escherichia coli infection | 0.026237322 | 4 |
| hsa05208 | Chemical carcinogenesis - reactive oxygen species | 0.037127148 | 4 |
| hsa04810 | Regulation of actin cytoskeleton | 0.039422474 | 4 |
| HALLMARK_INTERFERON_ALPHA_RESPONSE | HALLMARK_INTERFERON_ALPHA_RESPONSE | 0.013963252 | 4 |
| GO:0042605 | peptide antigen binding | 0.004449257 | 3 |
| GO:0015453 | oxidoreduction-driven active transmembrane transporter activity | 0.011784104 | 3 |
| GO:0008135 | translation factor activity, RNA binding | 0.011872616 | 3 |
| GO:0005200 | structural constituent of cytoskeleton | 0.02077857 | 3 |
| GO:0090079 | translation regulator activity, nucleic acid binding | 0.02077857 | 3 |
| GO:0009055 | electron transfer activity | 0.02796191 | 3 |

| | | | |
|---|---|---|---|
| GO:0015078 | proton transmembrane transporter activity | 0.036220351 | 3 |
| GO:0043021 | ribonucleoprotein complex binding | 0.049193215 | 3 |
| GO:0002483 | antigen processing and presentation of endogenous peptide antig | 0.000599498 | 3 |
| GO:0019883 | antigen processing and presentation of endogenous antigen | 0.001110053 | 3 |
| GO:0001916 | positive regulation of T cell mediated cytotoxicity | 0.001637637 | 3 |
| GO:0001914 | regulation of T cell mediated cytotoxicity | 0.003284852 | 3 |
| GO:0001913 | T cell mediated cytotoxicity | 0.005047785 | 3 |
| GO:0001912 | positive regulation of leukocyte mediated cytotoxicity | 0.007148688 | 3 |
| GO:0002711 | positive regulation of T cell mediated immunity | 0.007338205 | 3 |
| GO:0019731 | antibacterial humoral response | 0.008192644 | 3 |
| GO:0031343 | positive regulation of cell killing | 0.008783693 | 3 |
| GO:0048488 | synaptic vesicle endocytosis | 0.008783693 | 3 |
| GO:0140238 | presynaptic endocytosis | 0.008783693 | 3 |
| GO:0070527 | platelet aggregation | 0.010375884 | 3 |
| GO:0042267 | natural killer cell mediated cytotoxicity | 0.011530172 | 3 |
| GO:0002228 | natural killer cell mediated immunity | 0.012344331 | 3 |
| GO:0032729 | positive regulation of interferon-gamma production | 0.012344331 | 3 |
| GO:0036465 | synaptic vesicle recycling | 0.012709521 | 3 |
| GO:0006446 | regulation of translational initiation | 0.013854758 | 3 |
| GO:0034340 | response to type I interferon | 0.014191011 | 3 |
| GO:0051851 | modulation by host of symbiont process | 0.014388149 | 3 |
| GO:0002709 | regulation of T cell mediated immunity | 0.016319747 | 3 |
| GO:0019646 | aerobic electron transport chain | 0.017979978 | 3 |
| GO:0001738 | morphogenesis of a polarized epithelium | 0.019146925 | 3 |
| GO:0042773 | ATP synthesis coupled electron transport | 0.020387896 | 3 |
| GO:0042775 | mitochondrial ATP synthesis coupled electron transport | 0.020387896 | 3 |
| GO:0042102 | positive regulation of T cell proliferation | 0.02259141 | 3 |
| GO:0048024 | regulation of mRNA splicing, via spliceosome | 0.023879013 | 3 |
| GO:0002456 | T cell mediated immunity | 0.025720913 | 3 |
| GO:0002824 | positive regulation of adaptive immune response based on somat | 0.025720913 | 3 |
| GO:0050830 | defense response to Gram-positive bacterium | 0.0272557 | 3 |
| GO:0002708 | positive regulation of lymphocyte mediated immunity | 0.0272557 | 3 |
| GO:0032609 | interferon-gamma production | 0.0272557 | 3 |
| GO:0032649 | regulation of interferon-gamma production | 0.0272557 | 3 |
| GO:0002821 | positive regulation of adaptive immune response | 0.027704655 | 3 |
| GO:0022904 | respiratory electron transport chain | 0.028406977 | 3 |
| GO:0006413 | translational initiation | 0.028758478 | 3 |
| GO:0070585 | protein localization to mitochondrion | 0.031400366 | 3 |
| GO:0030168 | platelet activation | 0.034028318 | 3 |
| GO:0050684 | regulation of mRNA processing | 0.037989444 | 3 |
| GO:0002705 | positive regulation of leukocyte mediated immunity | 0.03850229 | 3 |
| GO:0050671 | positive regulation of lymphocyte proliferation | 0.04243828 | 3 |
| GO:0032946 | positive regulation of mononuclear cell proliferation | 0.043941145 | 3 |
| GO:0090316 | positive regulation of intracellular protein transport | 0.047598523 | 3 |
| hsa05310 | Asthma | 0.001854807 | 3 |
| hsa04672 | Intestinal immune network for IgA production | 0.005552377 | 3 |
| hsa05321 | Inflammatory bowel disease | 0.009879898 | 3 |
| hsa04658 | Th1 and Th2 cell differentiation | 0.023417731 | 3 |
| hsa05150 | Staphylococcus aureus infection | 0.024587422 | 3 |
| hsa04640 | Hematopoietic cell lineage | 0.025749048 | 3 |
| hsa04659 | Th17 cell differentiation | 0.030626881 | 3 |
| GO:0030957 | Tat protein binding | 0.005911204 | 2 |
| GO:0032395 | MHC class II receptor activity | 0.005911204 | 2 |
| GO:0042608 | T cell receptor binding | 0.005911204 | 2 |
| GO:0099186 | structural constituent of postsynapse | 0.009066241 | 2 |
| GO:0043024 | ribosomal small subunit binding | 0.011784104 | 2 |
| GO:0140693 | molecular condensate scaffold activity | 0.011784104 | 2 |
| GO:0003746 | translation elongation factor activity | 0.011817517 | 2 |
| GO:0004129 | cytochrome-c oxidase activity | 0.011817517 | 2 |
| GO:0098918 | structural constituent of synapse | 0.011817517 | 2 |
| GO:0016675 | oxidoreductase activity, acting on a heme group of donors | 0.011872616 | 2 |
| GO:0003785 | actin monomer binding | 0.02077857 | 2 |
| GO:0098974 | postsynaptic actin cytoskeleton organization | 0.005047785 | 2 |
| GO:0042989 | sequestering of actin monomers | 0.005930992 | 2 |
| GO:0099188 | postsynaptic cytoskeleton organization | 0.007000345 | 2 |
| GO:2001171 | positive regulation of ATP biosynthetic process | 0.008712518 | 2 |
| GO:0019885 | antigen processing and presentation of endogenous peptide antig | 0.011556363 | 2 |
| GO:0044794 | positive regulation by host of viral process | 0.014191011 | 2 |

| | | | |
|---|---|---|---|
| GO:0150105 | protein localization to cell-cell junction | 0.015176714 | 2 |
| GO:0048025 | negative regulation of mRNA splicing, via spliceosome | 0.017507126 | 2 |
| GO:2001169 | regulation of ATP biosynthetic process | 0.017507126 | 2 |
| GO:0006123 | mitochondrial electron transport, cytochrome c to oxygen | 0.018699521 | 2 |
| GO:0019835 | cytolysis | 0.019536156 | 2 |
| GO:0050686 | negative regulation of mRNA processing | 0.019536156 | 2 |
| GO:0002726 | positive regulation of T cell cytokine production | 0.020188781 | 2 |
| GO:0030810 | positive regulation of nucleotide biosynthetic process | 0.020188781 | 2 |
| GO:1900373 | positive regulation of purine nucleotide biosynthetic process | 0.020188781 | 2 |
| GO:1903901 | negative regulation of viral life cycle | 0.020188781 | 2 |
| GO:0042026 | protein refolding | 0.021020212 | 2 |
| GO:0033119 | negative regulation of RNA splicing | 0.02240976 | 2 |
| GO:0090025 | regulation of monocyte chemotaxis | 0.023432546 | 2 |
| GO:0002474 | antigen processing and presentation of peptide antigen via MHC | 0.024456953 | 2 |
| GO:0045070 | positive regulation of viral genome replication | 0.026104532 | 2 |
| GO:0070633 | transepithelial transport | 0.0272557 | 2 |
| GO:0035633 | maintenance of blood-brain barrier | 0.028758478 | 2 |
| GO:1903580 | positive regulation of ATP metabolic process | 0.029981921 | 2 |
| GO:1903749 | positive regulation of establishment of protein localization to mit | 0.029981921 | 2 |
| GO:0002369 | T cell cytokine production | 0.03245365 | 2 |
| GO:0002724 | regulation of T cell cytokine production | 0.03245365 | 2 |
| GO:0043516 | regulation of DNA damage response, signal transduction by p53 c | 0.03245365 | 2 |
| GO:1900371 | regulation of purine nucleotide biosynthetic process | 0.040021133 | 2 |
| GO:0030808 | regulation of nucleotide biosynthetic process | 0.041591865 | 2 |
| GO:0031640 | killing of cells of another organism | 0.043941145 | 2 |
| GO:0044788 | modulation by host of viral process | 0.043941145 | 2 |
| GO:0042269 | regulation of natural killer cell mediated cytotoxicity | 0.045255691 | 2 |
| GO:0045646 | regulation of erythrocyte differentiation | 0.045255691 | 2 |
| GO:0045981 | positive regulation of nucleotide metabolic process | 0.045731951 | 2 |
| GO:1900544 | positive regulation of purine nucleotide metabolic process | 0.045731951 | 2 |
| GO:0035094 | response to nicotine | 0.047039445 | 2 |
| GO:1903747 | regulation of establishment of protein localization to mitochondri | 0.047787832 | 2 |
| GO:0002715 | regulation of natural killer cell mediated immunity | 0.048534755 | 2 |
| GO:0032873 | negative regulation of stress-activated MAPK cascade | 0.048534755 | 2 |
| GO:0070303 | negative regulation of stress-activated protein kinase signaling ca | 0.048534755 | 2 |

| ID | Description | p.adjust | Count |
|---|---|---|---|
| HALLMARK_E2F_TARGETS | HALLMARK_E2F_TARGETS | 1.57726E-06 | 11 |
| HALLMARK_MYC_TARGETS_V1 | HALLMARK_MYC_TARGETS_V1 | 1.57726E-06 | 11 |
| GO:0009150 | purine ribonucleotide metabolic process | 1.98489E-06 | 10 |
| GO:0009259 | ribonucleotide metabolic process | 2.37708E-06 | 10 |
| GO:0019693 | ribose phosphate metabolic process | 2.6488E-06 | 10 |
| GO:0006163 | purine nucleotide metabolic process | 2.6488E-06 | 10 |
| HALLMARK_GLYCOLYSIS | HALLMARK_GLYCOLYSIS | 1.1158E-05 | 10 |
| GO:0009205 | purine ribonucleoside triphosphate metabolic process | 1.90845E-07 | 9 |
| GO:0009144 | purine nucleoside triphosphate metabolic process | 2.10076E-07 | 9 |
| GO:0009199 | ribonucleoside triphosphate metabolic process | 2.10076E-07 | 9 |
| GO:0009141 | nucleoside triphosphate metabolic process | 3.61017E-07 | 9 |
| GO:0045296 | cadherin binding | 0.00011269 | 8 |
| GO:0006096 | glycolytic process | 5.37666E-09 | 8 |
| GO:0006757 | ATP generation from ADP | 5.37666E-09 | 8 |
| GO:0046031 | ADP metabolic process | 7.55847E-09 | 8 |
| GO:0006165 | nucleoside diphosphate phosphorylation | 8.798E-09 | 8 |
| GO:0046939 | nucleotide phosphorylation | 8.798E-09 | 8 |
| GO:0009135 | purine nucleoside diphosphate metabolic process | 8.798E-09 | 8 |
| GO:0009179 | purine ribonucleoside diphosphate metabolic process | 8.798E-09 | 8 |
| GO:0006090 | pyruvate metabolic process | 1.04558E-08 | 8 |
| GO:0009185 | ribonucleoside diphosphate metabolic process | 1.43946E-08 | 8 |
| GO:0009132 | nucleoside diphosphate metabolic process | 4.21482E-08 | 8 |
| GO:0016052 | carbohydrate catabolic process | 1.38051E-07 | 8 |
| GO:0046034 | ATP metabolic process | 1.34076E-06 | 8 |
| hsa00010 | Glycolysis / Gluconeogenesis | 4.18413E-09 | 8 |
| HALLMARK_MTORC1_SIGNALING | HALLMARK_MTORC1_SIGNALING | 0.000613568 | 8 |
| GO:0033044 | regulation of chromosome organization | 4.5682E-05 | 7 |
| hsa01230 | Biosynthesis of amino acids | 2.31386E-07 | 7 |
| hsa01200 | Carbon metabolism | 3.09789E-06 | 7 |
| HALLMARK_G2M_CHECKPOINT | HALLMARK_G2M_CHECKPOINT | 0.002779609 | 7 |
| HALLMARK_HYPOXIA | HALLMARK_HYPOXIA | 0.002779609 | 7 |
| GO:0019318 | hexose metabolic process | 0.000199166 | 6 |
| GO:0005996 | monosaccharide metabolic process | 0.00030232 | 6 |
| GO:0051054 | positive regulation of DNA metabolic process | 0.000763772 | 6 |
| GO:0000280 | nuclear division | 0.007489919 | 6 |
| hsa04066 | HIF-1 signaling pathway | 3.86153E-05 | 6 |
| GO:0003697 | single-stranded DNA binding | 0.000745178 | 5 |
| GO:0016829 | lyase activity | 0.003620724 | 5 |
| GO:0016887 | ATP hydrolysis activity | 0.015625698 | 5 |
| GO:2001252 | positive regulation of chromosome organization | 9.20627E-05 | 5 |
| GO:0061640 | cytoskeleton-dependent cytokinesis | 0.000112397 | 5 |
| GO:2000278 | regulation of DNA biosynthetic process | 0.000153829 | 5 |
| GO:0000723 | telomere maintenance | 0.000388831 | 5 |
| GO:0007338 | single fertilization | 0.000459747 | 5 |
| GO:0032200 | telomere organization | 0.000826205 | 5 |
| GO:0000910 | cytokinesis | 0.000856984 | 5 |
| GO:0006006 | glucose metabolic process | 0.000866444 | 5 |
| GO:0140013 | meiotic nuclear division | 0.000977942 | 5 |
| GO:0071897 | DNA biosynthetic process | 0.001058728 | 5 |
| GO:0009566 | fertilization | 0.001110504 | 5 |
| GO:1903046 | meiotic cell cycle process | 0.001391044 | 5 |
| GO:0050821 | protein stabilization | 0.001435296 | 5 |
| GO:0006457 | protein folding | 0.001719841 | 5 |
| GO:0051321 | meiotic cell cycle | 0.005108308 | 5 |
| GO:0098813 | nuclear chromosome segregation | 0.008097026 | 5 |
| GO:0031647 | regulation of protein stability | 0.008468952 | 5 |
| GO:0007059 | chromosome segregation | 0.014791914 | 5 |
| GO:0044282 | small molecule catabolic process | 0.015069239 | 5 |
| GO:0006979 | response to oxidative stress | 0.021180172 | 5 |
| GO:1903829 | positive regulation of protein localization | 0.024144605 | 5 |
| hsa05012 | Parkinson disease | 0.03157584 | 5 |
| GO:0016835 | carbon-oxygen lyase activity | 0.001867468 | 4 |
| GO:0051082 | unfolded protein binding | 0.00516113 | 4 |
| GO:0016853 | isomerase activity | 0.008559728 | 4 |
| GO:0006735 | NADH regeneration | 1.9947E-06 | 4 |
| GO:0061621 | canonical glycolysis | 1.9947E-06 | 4 |
| GO:0061718 | glucose catabolic process to pyruvate | 1.9947E-06 | 4 |

| | | | |
|---|---|---|---|
| GO:0061620 | glycolytic process through glucose-6-phosphate | 2.78603E-06 | 4 |
| GO:0061615 | glycolytic process through fructose-6-phosphate | 3.38251E-06 | 4 |
| GO:0006007 | glucose catabolic process | 1.05155E-05 | 4 |
| GO:0006734 | NADH metabolic process | 4.66947E-05 | 4 |
| GO:1904358 | positive regulation of telomere maintenance via telomere length | 4.66947E-05 | 4 |
| GO:0007339 | binding of sperm to zona pellucida | 6.18222E-05 | 4 |
| GO:0019320 | hexose catabolic process | 6.18222E-05 | 4 |
| GO:0046365 | monosaccharide catabolic process | 8.53193E-05 | 4 |
| GO:0035036 | sperm-egg recognition | 0.00010878 | 4 |
| GO:1904356 | regulation of telomere maintenance via telomere lengthening | 0.000197416 | 4 |
| GO:0032206 | positive regulation of telomere maintenance | 0.000276988 | 4 |
| GO:0009988 | cell-cell recognition | 0.000336902 | 4 |
| GO:2000573 | positive regulation of DNA biosynthetic process | 0.000388831 | 4 |
| GO:0010833 | telomere maintenance via telomere lengthening | 0.00052285 | 4 |
| GO:0000281 | mitotic cytokinesis | 0.000622463 | 4 |
| GO:1900182 | positive regulation of protein localization to nucleus | 0.000661479 | 4 |
| GO:0032204 | regulation of telomere maintenance | 0.001058728 | 4 |
| GO:1900180 | regulation of protein localization to nucleus | 0.002979715 | 4 |
| GO:0007051 | spindle organization | 0.0100806 | 4 |
| GO:0006403 | RNA localization | 0.011259646 | 4 |
| GO:0000070 | mitotic sister chromatid segregation | 0.011449213 | 4 |
| GO:0009152 | purine ribonucleotide biosynthetic process | 0.014411976 | 4 |
| GO:0008037 | cell recognition | 0.015948791 | 4 |
| GO:2001234 | negative regulation of apoptotic signaling pathway | 0.016248578 | 4 |
| GO:0009260 | ribonucleotide biosynthetic process | 0.016363132 | 4 |
| GO:0000819 | sister chromatid segregation | 0.017527431 | 4 |
| GO:0046390 | ribose phosphate biosynthetic process | 0.017617223 | 4 |
| GO:0006164 | purine nucleotide biosynthetic process | 0.01773761 | 4 |
| GO:0072522 | purine-containing compound biosynthetic process | 0.019455581 | 4 |
| GO:0009165 | nucleotide biosynthetic process | 0.03251811 | 4 |
| GO:0034504 | protein localization to nucleus | 0.032840741 | 4 |
| GO:1901293 | nucleoside phosphate biosynthetic process | 0.032840741 | 4 |
| GO:1904951 | positive regulation of establishment of protein localization | 0.033844139 | 4 |
| GO:0140014 | mitotic nuclear division | 0.037977067 | 4 |
| hsa04110 | Cell cycle | 0.03157584 | 4 |
| GO:0140662 | ATP-dependent protein folding chaperone | 0.003902021 | 3 |
| GO:0016765 | transferase activity, transferring alkyl or aryl (other than methyl | 0.008266921 | 3 |
| GO:0016836 | hydro-lyase activity | 0.008559728 | 3 |
| GO:0044183 | protein folding chaperone | 0.008798075 | 3 |
| GO:0005200 | structural constituent of cytoskeleton | 0.023438738 | 3 |
| GO:0016922 | nuclear receptor binding | 0.040551094 | 3 |
| GO:1904851 | positive regulation of establishment of protein localization to telo | 4.66947E-05 | 3 |
| GO:0070203 | regulation of establishment of protein localization to telomere | 5.48796E-05 | 3 |
| GO:1904869 | regulation of protein localization to Cajal body | 5.48796E-05 | 3 |
| GO:1904871 | positive regulation of protein localization to Cajal body | 5.48796E-05 | 3 |
| GO:0070202 | regulation of establishment of protein localization to chromosome | 6.18222E-05 | 3 |
| GO:1903405 | protein localization to nuclear body | 6.18222E-05 | 3 |
| GO:1904816 | positive regulation of protein localization to chromosome, telome | 6.18222E-05 | 3 |
| GO:1904867 | protein localization to Cajal body | 6.18222E-05 | 3 |
| GO:1904814 | regulation of protein localization to chromosome, telomeric regio | 9.2493E-05 | 3 |
| GO:1990173 | protein localization to nucleoplasm | 9.2493E-05 | 3 |
| GO:1904874 | positive regulation of telomerase RNA localization to Cajal body | 0.000110311 | 3 |
| GO:0070200 | establishment of protein localization to telomere | 0.000154092 | 3 |
| GO:1904872 | regulation of telomerase RNA localization to Cajal body | 0.000180863 | 3 |
| GO:0090670 | RNA localization to Cajal body | 0.000197416 | 3 |
| GO:0090671 | telomerase RNA localization to Cajal body | 0.000197416 | 3 |
| GO:0090672 | telomerase RNA localization | 0.000197416 | 3 |
| GO:0090685 | RNA localization to nucleus | 0.000197416 | 3 |
| GO:0070199 | establishment of protein localization to chromosome | 0.000505474 | 3 |
| GO:0070198 | protein localization to chromosome, telomeric region | 0.000661479 | 3 |
| GO:0032212 | positive regulation of telomere maintenance via telomerase | 0.000893547 | 3 |
| GO:1901998 | toxin transport | 0.00136585 | 3 |
| GO:0032210 | regulation of telomere maintenance via telomerase | 0.002979715 | 3 |
| GO:0032481 | positive regulation of type I interferon production | 0.004915029 | 3 |
| GO:0007004 | telomere maintenance via telomerase | 0.006273557 | 3 |
| GO:0061077 | chaperone-mediated protein folding | 0.006742817 | 3 |
| GO:0006278 | RNA-templated DNA biosynthetic process | 0.00764852 | 3 |
| GO:0002437 | inflammatory response to antigenic stimulus | 0.008516457 | 3 |

| | | | |
|---|---|---|---|
| GO:0006094 | gluconeogenesis | 0.009626628 | 3 |
| GO:0031397 | negative regulation of protein ubiquitination | 0.00986326 | 3 |
| GO:0019319 | hexose biosynthetic process | 0.010344119 | 3 |
| GO:0046364 | monosaccharide biosynthetic process | 0.011449213 | 3 |
| GO:1903321 | negative regulation of protein modification by small protein conju | 0.013397846 | 3 |
| GO:0032392 | DNA geometric change | 0.014411976 | 3 |
| GO:0045132 | meiotic chromosome segregation | 0.014791914 | 3 |
| GO:2001243 | negative regulation of intrinsic apoptotic signaling pathway | 0.014791914 | 3 |
| GO:0032479 | regulation of type I interferon production | 0.015948791 | 3 |
| GO:0032606 | type I interferon production | 0.015948791 | 3 |
| GO:0071103 | DNA conformation change | 0.016101201 | 3 |
| GO:1905818 | regulation of chromosome separation | 0.017617223 | 3 |
| GO:0034502 | protein localization to chromosome | 0.018997484 | 3 |
| GO:0009206 | purine ribonucleoside triphosphate biosynthetic process | 0.019455581 | 3 |
| GO:0009145 | purine nucleoside triphosphate biosynthetic process | 0.019486694 | 3 |
| GO:0071346 | cellular response to interferon-gamma | 0.019486694 | 3 |
| GO:0009201 | ribonucleoside triphosphate biosynthetic process | 0.021264016 | 3 |
| GO:0051225 | spindle assembly | 0.023452873 | 3 |
| GO:0042542 | response to hydrogen peroxide | 0.024144605 | 3 |
| GO:0051983 | regulation of chromosome segregation | 0.025025545 | 3 |
| GO:0007052 | mitotic spindle organization | 0.025379505 | 3 |
| GO:0009142 | nucleoside triphosphate biosynthetic process | 0.025731159 | 3 |
| GO:0051304 | chromosome separation | 0.025908142 | 3 |
| GO:0034341 | response to interferon-gamma | 0.027910612 | 3 |
| GO:0046165 | alcohol biosynthetic process | 0.029220811 | 3 |
| GO:1902850 | microtubule cytoskeleton organization involved in mitosis | 0.037977067 | 3 |
| GO:0002262 | myeloid cell homeostasis | 0.040933512 | 3 |
| GO:2001242 | regulation of intrinsic apoptotic signaling pathway | 0.042253945 | 3 |
| GO:0051099 | positive regulation of binding | 0.044676151 | 3 |
| GO:0004659 | prenyltransferase activity | 0.011102849 | 2 |
| GO:0035035 | histone acetyltransferase binding | 0.017329878 | 2 |
| GO:0000217 | DNA secondary structure binding | 0.036722031 | 2 |
| GO:0015035 | protein-disulfide reductase activity | 0.040322286 | 2 |
| GO:0001222 | transcription corepressor binding | 0.043046253 | 2 |
| GO:0015036 | disulfide oxidoreductase activity | 0.043046253 | 2 |
| GO:0006555 | methionine metabolic process | 0.00736926 | 2 |
| GO:0006089 | lactate metabolic process | 0.008059225 | 2 |
| GO:1902176 | negative regulation of oxidative stress-induced intrinsic apoptotic | 0.009626628 | 2 |
| GO:0032069 | regulation of nuclease activity | 0.011862435 | 2 |
| GO:0051444 | negative regulation of ubiquitin-protein transferase activity | 0.011862435 | 2 |
| GO:0031639 | plasminogen activation | 0.016101201 | 2 |
| GO:1902175 | regulation of oxidative stress-induced intrinsic apoptotic signaling | 0.017617223 | 2 |
| GO:0032148 | activation of protein kinase B activity | 0.01940127 | 2 |
| GO:1902042 | negative regulation of extrinsic apoptotic signaling pathway via d | 0.01940127 | 2 |
| GO:0040020 | regulation of meiotic nuclear division | 0.01986531 | 2 |
| GO:0000096 | sulfur amino acid metabolic process | 0.020955524 | 2 |
| GO:0045648 | positive regulation of erythrocyte differentiation | 0.021733208 | 2 |
| GO:0033260 | nuclear DNA replication | 0.025731159 | 2 |
| GO:0042401 | cellular biogenic amine biosynthetic process | 0.026703426 | 2 |
| GO:0009309 | amine biosynthetic process | 0.027680528 | 2 |
| GO:0032506 | cytokinetic process | 0.027680528 | 2 |
| GO:0046688 | response to copper ion | 0.028662131 | 2 |
| GO:0044786 | cell cycle DNA replication | 0.02964792 | 2 |
| GO:0044275 | cellular carbohydrate catabolic process | 0.032267737 | 2 |
| GO:0008631 | intrinsic apoptotic signaling pathway in response to oxidative stre | 0.032840741 | 2 |
| GO:0042398 | cellular modified amino acid biosynthetic process | 0.033844139 | 2 |
| GO:0002861 | regulation of inflammatory response to antigenic stimulus | 0.034629732 | 2 |
| GO:0045646 | regulation of erythrocyte differentiation | 0.034629732 | 2 |
| GO:0051972 | regulation of telomerase activity | 0.034629732 | 2 |
| GO:0009066 | aspartate family amino acid metabolic process | 0.03585901 | 2 |
| GO:0018198 | peptidyl-cysteine modification | 0.037977067 | 2 |
| GO:1902041 | regulation of extrinsic apoptotic signaling pathway via death dom | 0.037977067 | 2 |
| GO:0033046 | negative regulation of sister chromatid segregation | 0.038663955 | 2 |
| GO:0033048 | negative regulation of mitotic sister chromatid segregation | 0.038663955 | 2 |
| GO:2000816 | negative regulation of mitotic sister chromatid separation | 0.038663955 | 2 |
| GO:0051985 | negative regulation of chromosome segregation | 0.040933512 | 2 |
| GO:1905819 | negative regulation of chromosome separation | 0.040933512 | 2 |
| GO:1903202 | negative regulation of oxidative stress-induced cell death | 0.043476991 | 2 |

| | | | |
|---|---|---|---|
| GO:0033047 | regulation of mitotic sister chromatid segregation | 0.044505143 | 2 |
| GO:0051438 | regulation of ubiquitin-protein transferase activity | 0.044505143 | 2 |
| GO:0043388 | positive regulation of DNA binding | 0.045275403 | 2 |
| GO:0090329 | regulation of DNA-templated DNA replication | 0.045275403 | 2 |
| GO:0006695 | cholesterol biosynthetic process | 0.046302194 | 2 |
| GO:1902653 | secondary alcohol biosynthetic process | 0.046302194 | 2 |

| ID | Description | p.adjust | Count |
|---|---|---|---|
| GO:0010876 | lipid localization | 0.003149704 | 8 |
| GO:0006869 | lipid transport | 0.009016423 | 7 |
| GO:0051051 | negative regulation of transport | 0.01066551 | 7 |
| GO:0005543 | phospholipid binding | 0.021855082 | 6 |
| GO:0030100 | regulation of endocytosis | 0.003149704 | 6 |
| GO:0007162 | negative regulation of cell adhesion | 0.009016423 | 6 |
| GO:0022407 | regulation of cell-cell adhesion | 0.037426836 | 6 |
| GO:0005319 | lipid transporter activity | 0.005665478 | 5 |
| GO:0046890 | regulation of lipid biosynthetic process | 0.009016423 | 5 |
| GO:0006898 | receptor-mediated endocytosis | 0.024427391 | 5 |
| GO:1990778 | protein localization to cell periphery | 0.043420845 | 5 |
| GO:0062012 | regulation of small molecule metabolic process | 0.043420845 | 5 |
| GO:0019216 | regulation of lipid metabolic process | 0.043420845 | 5 |
| GO:0050863 | regulation of T cell activation | 0.047865444 | 5 |
| GO:1903037 | regulation of leukocyte cell-cell adhesion | 0.047865444 | 5 |
| GO:0015399 | primary active transmembrane transporter activity | 0.021855082 | 4 |
| GO:0045806 | negative regulation of endocytosis | 0.003149704 | 4 |
| GO:0010565 | regulation of cellular ketone metabolic process | 0.024427391 | 4 |
| GO:0045834 | positive regulation of lipid metabolic process | 0.031231058 | 4 |
| GO:0097530 | granulocyte migration | 0.031231058 | 4 |
| GO:0022408 | negative regulation of cell-cell adhesion | 0.043420845 | 4 |
| GO:0042180 | cellular ketone metabolic process | 0.047865444 | 4 |
| GO:0005504 | fatty acid binding | 0.017880077 | 3 |
| GO:0033293 | monocarboxylic acid binding | 0.022518584 | 3 |
| GO:0045940 | positive regulation of steroid metabolic process | 0.009016423 | 3 |
| GO:0035094 | response to nicotine | 0.02362114 | 3 |
| GO:0042304 | regulation of fatty acid biosynthetic process | 0.02362114 | 3 |
| GO:0015909 | long-chain fatty acid transport | 0.033125518 | 3 |
| GO:0097006 | regulation of plasma lipoprotein particle levels | 0.043420845 | 3 |
| GO:0019915 | lipid storage | 0.043420845 | 3 |
| GO:0048145 | regulation of fibroblast proliferation | 0.043420845 | 3 |
| GO:0046889 | positive regulation of lipid biosynthetic process | 0.043420845 | 3 |
| GO:0019217 | regulation of fatty acid metabolic process | 0.047865444 | 3 |
| GO:0015908 | fatty acid transport | 0.04946155 | 3 |
| GO:0019218 | regulation of steroid metabolic process | 0.04946155 | 3 |
| GO:0048144 | fibroblast proliferation | 0.04946155 | 3 |
| GO:1903076 | regulation of protein localization to plasma membrane | 0.04946155 | 3 |
| GO:0015914 | phospholipid transport | 0.04946155 | 3 |
| GO:0050998 | nitric-oxide synthase binding | 0.021855082 | 2 |
| GO:0005324 | long-chain fatty acid transporter activity | 0.021855082 | 2 |
| GO:0023026 | MHC class II protein complex binding | 0.041712589 | 2 |
| GO:0031392 | regulation of prostaglandin biosynthetic process | 0.031231058 | 2 |
| GO:0033700 | phospholipid efflux | 0.031231058 | 2 |
| GO:0045176 | apical protein localization | 0.034879322 | 2 |
| GO:2001279 | regulation of unsaturated fatty acid biosynthetic process | 0.034879322 | 2 |
| GO:0034375 | high-density lipoprotein particle remodeling | 0.037426836 | 2 |
| GO:0010893 | positive regulation of steroid biosynthetic process | 0.047865444 | 2 |
| GO:0072677 | eosinophil migration | 0.047865444 | 2 |
| GO:0019883 | antigen processing and presentation of endogenous antigen | 0.04946155 | 2 |

| ID | Description | p.adjust | Count |
|---|---|---|---|
| GO:0000280 | nuclear division | 0.000878363 | 9 |
| GO:0140013 | meiotic nuclear division | 0.00049675 | 7 |
| GO:1903046 | meiotic cell cycle process | 0.00049675 | 7 |
| GO:0051321 | meiotic cell cycle | 0.001771545 | 7 |
| HALLMARK_E2F_TARGETS | HALLMARK_E2F_TARGETS | 0.002650716 | 7 |
| GO:0030217 | T cell differentiation | 0.017142215 | 6 |
| GO:0042326 | negative regulation of phosphorylation | 0.026641335 | 6 |
| GO:1903706 | regulation of hemopoiesis | 0.031216409 | 6 |
| GO:0030098 | lymphocyte differentiation | 0.032082237 | 6 |
| GO:0045936 | negative regulation of phosphate metabolic process | 0.033904524 | 6 |
| GO:0010563 | negative regulation of phosphorus metabolic process | 0.033904524 | 6 |
| GO:0044772 | mitotic cell cycle phase transition | 0.041121804 | 6 |
| GO:1903131 | mononuclear cell differentiation | 0.041121804 | 6 |
| HALLMARK_MTORC1_SIGNALING | HALLMARK_MTORC1_SIGNALING | 0.01073556 | 6 |
| GO:0007051 | spindle organization | 0.017142215 | 5 |
| GO:0051348 | negative regulation of transferase activity | 0.032530755 | 5 |
| GO:1902105 | regulation of leukocyte differentiation | 0.041121804 | 5 |
| GO:0001933 | negative regulation of protein phosphorylation | 0.046700172 | 5 |
| GO:1901990 | regulation of mitotic cell cycle phase transition | 0.047350379 | 5 |
| GO:0042176 | regulation of protein catabolic process | 0.047350379 | 5 |
| GO:0050863 | regulation of T cell activation | 0.047740669 | 5 |
| GO:1903037 | regulation of leukocyte cell-cell adhesion | 0.047740669 | 5 |
| GO:0009185 | ribonucleoside diphosphate metabolic process | 0.022414828 | 4 |
| GO:0007127 | meiosis I | 0.026641335 | 4 |
| GO:0009132 | nucleoside diphosphate metabolic process | 0.026641335 | 4 |
| GO:0051304 | chromosome separation | 0.026641335 | 4 |
| GO:0061982 | meiosis I cell cycle process | 0.026641335 | 4 |
| GO:0051783 | regulation of nuclear division | 0.031119806 | 4 |
| GO:0007292 | female gamete generation | 0.033132428 | 4 |
| GO:0006694 | steroid biosynthetic process | 0.041121804 | 4 |
| GO:0045580 | regulation of T cell differentiation | 0.041121804 | 4 |
| GO:0045619 | regulation of lymphocyte differentiation | 0.047350379 | 4 |
| GO:0007143 | female meiotic nuclear division | 0.017142215 | 3 |
| GO:0045589 | regulation of regulatory T cell differentiation | 0.017142215 | 3 |
| GO:0045066 | regulatory T cell differentiation | 0.017142215 | 3 |
| GO:0006096 | glycolytic process | 0.041121804 | 3 |
| GO:0006757 | ATP generation from ADP | 0.041389929 | 3 |
| GO:0033077 | T cell differentiation in thymus | 0.04310687 | 3 |
| GO:0030071 | regulation of mitotic metaphase/anaphase transition | 0.047251717 | 3 |
| GO:0046031 | ADP metabolic process | 0.047251717 | 3 |
| GO:0007091 | metaphase/anaphase transition of mitotic cell cycle | 0.047350379 | 3 |
| GO:1902099 | regulation of metaphase/anaphase transition of cell cycle | 0.047350379 | 3 |
| GO:2001251 | negative regulation of chromosome organization | 0.047350379 | 3 |
| GO:0010965 | regulation of mitotic sister chromatid separation | 0.047350379 | 3 |
| GO:0044784 | metaphase/anaphase transition of cell cycle | 0.047350379 | 3 |
| GO:0006165 | nucleoside diphosphate phosphorylation | 0.047350379 | 3 |
| GO:1901992 | positive regulation of mitotic cell cycle phase transition | 0.047350379 | 3 |
| GO:0045132 | meiotic chromosome segregation | 0.047350379 | 3 |
| GO:0051306 | mitotic sister chromatid separation | 0.047350379 | 3 |
| GO:0046939 | nucleotide phosphorylation | 0.047350379 | 3 |
| GO:0045639 | positive regulation of myeloid cell differentiation | 0.047460796 | 3 |
| GO:0009135 | purine nucleoside diphosphate metabolic process | 0.047460796 | 3 |
| GO:0009179 | purine ribonucleoside diphosphate metabolic process | 0.047460796 | 3 |
| GO:0033045 | regulation of sister chromatid segregation | 0.048654292 | 3 |
| GO:0006090 | pyruvate metabolic process | 0.048654292 | 3 |
| GO:1902106 | negative regulation of leukocyte differentiation | 0.048654292 | 3 |
| GO:0006766 | vitamin metabolic process | 0.049682822 | 3 |
| GO:0030388 | fructose 1,6-bisphosphate metabolic process | 0.026641335 | 2 |
| GO:1900222 | negative regulation of amyloid-beta clearance | 0.026641335 | 2 |
| GO:0007144 | female meiosis I | 0.031119806 | 2 |
| GO:0000212 | meiotic spindle organization | 0.033132428 | 2 |
| GO:0045091 | regulation of single stranded viral RNA replication via double stra | 0.041121804 | 2 |
| GO:0039692 | single stranded viral RNA replication via double stranded DNA in | 0.041121804 | 2 |
| GO:1900221 | regulation of amyloid-beta clearance | 0.041121804 | 2 |
| GO:0051383 | kinetochore organization | 0.047251717 | 2 |
| GO:0045116 | protein neddylation | 0.047350379 | 2 |
| GO:0039694 | viral RNA genome replication | 0.049682822 | 2 |

| ID | Description | p.adjust | Count |
|---|---|---|---|
| GO:0030098 | lymphocyte differentiation | 0.009263796 | 7 |
| GO:1903131 | mononuclear cell differentiation | 0.009813734 | 7 |
| GO:0022407 | regulation of cell-cell adhesion | 0.009813734 | 7 |
| HALLMARK_E2F_TARGETS | HALLMARK_E2F_TARGETS | 0.007609589 | 7 |
| GO:0140097 | catalytic activity, acting on DNA | 0.002408251 | 6 |
| GO:0001906 | cell killing | 0.00345674 | 6 |
| GO:0002699 | positive regulation of immune effector process | 0.008064487 | 6 |
| GO:0030217 | T cell differentiation | 0.009263796 | 6 |
| GO:0006310 | DNA recombination | 0.009813734 | 6 |
| GO:0042113 | B cell activation | 0.009813734 | 6 |
| GO:0050863 | regulation of T cell activation | 0.012741974 | 6 |
| GO:1903037 | regulation of leukocyte cell-cell adhesion | 0.012741974 | 6 |
| GO:0002697 | regulation of immune effector process | 0.012741974 | 6 |
| GO:0007159 | leukocyte cell-cell adhesion | 0.016730926 | 6 |
| hsa04110 | Cell cycle | 0.000969286 | 6 |
| GO:0030183 | B cell differentiation | 0.008064487 | 5 |
| GO:0045580 | regulation of T cell differentiation | 0.009263796 | 5 |
| GO:0045619 | regulation of lymphocyte differentiation | 0.009813734 | 5 |
| GO:0050870 | positive regulation of T cell activation | 0.014581726 | 5 |
| GO:1903039 | positive regulation of leukocyte cell-cell adhesion | 0.016730926 | 5 |
| GO:0002366 | leukocyte activation involved in immune response | 0.019577642 | 5 |
| GO:0048872 | homeostasis of number of cells | 0.019577642 | 5 |
| GO:0002263 | cell activation involved in immune response | 0.019577642 | 5 |
| GO:0046651 | lymphocyte proliferation | 0.020786127 | 5 |
| GO:0032943 | mononuclear cell proliferation | 0.022010603 | 5 |
| GO:1902105 | regulation of leukocyte differentiation | 0.022982438 | 5 |
| GO:0022409 | positive regulation of cell-cell adhesion | 0.023361999 | 5 |
| GO:0002440 | production of molecular mediator of immune response | 0.026772159 | 5 |
| GO:0070661 | leukocyte proliferation | 0.031317785 | 5 |
| GO:0002460 | adaptive immune response based on somatic recombination of im | 0.036647048 | 5 |
| GO:0051251 | positive regulation of lymphocyte activation | 0.044303319 | 5 |
| GO:1903706 | regulation of hemopoiesis | 0.045610809 | 5 |
| hsa04640 | Hematopoietic cell lineage | 0.000969286 | 5 |
| GO:0008094 | ATP-dependent activity, acting on DNA | 0.012498222 | 4 |
| GO:0009123 | nucleoside monophosphate metabolic process | 0.008064487 | 4 |
| GO:0033077 | T cell differentiation in thymus | 0.009263796 | 4 |
| GO:0032508 | DNA duplex unwinding | 0.009263796 | 4 |
| GO:0032392 | DNA geometric change | 0.009813734 | 4 |
| GO:0032479 | regulation of type I interferon production | 0.009813734 | 4 |
| GO:0032606 | type I interferon production | 0.009813734 | 4 |
| GO:0071103 | DNA conformation change | 0.009813734 | 4 |
| GO:0045582 | positive regulation of T cell differentiation | 0.009813734 | 4 |
| GO:0045621 | positive regulation of lymphocyte differentiation | 0.012741974 | 4 |
| GO:0046631 | alpha-beta T cell activation | 0.020786127 | 4 |
| GO:1902107 | positive regulation of leukocyte differentiation | 0.022985292 | 4 |
| GO:1903708 | positive regulation of hemopoiesis | 0.022985292 | 4 |
| GO:0002285 | lymphocyte activation involved in immune response | 0.03206103 | 4 |
| GO:0000819 | sister chromatid segregation | 0.044040591 | 4 |
| GO:0050670 | regulation of lymphocyte proliferation | 0.044040591 | 4 |
| GO:0032944 | regulation of mononuclear cell proliferation | 0.044770518 | 4 |
| hsa05340 | Primary immunodeficiency | 0.000969286 | 4 |
| GO:0017116 | single-stranded DNA helicase activity | 0.002408251 | 3 |
| GO:0003678 | DNA helicase activity | 0.023189558 | 3 |
| GO:0000727 | double-strand break repair via break-induced replication | 0.00345674 | 3 |
| GO:0030174 | regulation of DNA-templated DNA replication initiation | 0.00345674 | 3 |
| GO:0006268 | DNA unwinding involved in DNA replication | 0.008064487 | 3 |
| GO:0046037 | GMP metabolic process | 0.008064487 | 3 |
| GO:0002313 | mature B cell differentiation involved in immune response | 0.009263796 | 3 |
| GO:0002335 | mature B cell differentiation | 0.009813734 | 3 |
| GO:0006270 | DNA replication initiation | 0.009813734 | 3 |
| GO:0009167 | purine ribonucleoside monophosphate metabolic process | 0.009813734 | 3 |
| GO:0009124 | nucleoside monophosphate biosynthetic process | 0.009813734 | 3 |
| GO:0009126 | purine nucleoside monophosphate metabolic process | 0.009813734 | 3 |
| GO:0009394 | 2'-deoxyribonucleotide metabolic process | 0.009813734 | 3 |
| GO:0009262 | deoxyribonucleotide metabolic process | 0.009813734 | 3 |
| GO:0019692 | deoxyribose phosphate metabolic process | 0.009813734 | 3 |
| GO:0031640 | killing of cells of another organism | 0.009813734 | 3 |

| | | | |
|---|---|---|---|
| GO:0019884 | antigen processing and presentation of exogenous antigen | 0.01130417 | 3 |
| GO:0046638 | positive regulation of alpha-beta T cell differentiation | 0.012741974 | 3 |
| GO:0090329 | regulation of DNA-templated DNA replication | 0.014686291 | 3 |
| GO:0009161 | ribonucleoside monophosphate metabolic process | 0.015505271 | 3 |
| GO:0032481 | positive regulation of type I interferon production | 0.016730926 | 3 |
| GO:0031100 | animal organ regeneration | 0.016730926 | 3 |
| GO:0031343 | positive regulation of cell killing | 0.01717383 | 3 |
| GO:0009620 | response to fungus | 0.017621984 | 3 |
| GO:0002260 | lymphocyte homeostasis | 0.018864872 | 3 |
| GO:0046635 | positive regulation of alpha-beta T cell activation | 0.019577642 | 3 |
| GO:0046637 | regulation of alpha-beta T cell differentiation | 0.019577642 | 3 |
| GO:0032729 | positive regulation of interferon-gamma production | 0.020786127 | 3 |
| GO:0002312 | B cell activation involved in immune response | 0.02711868 | 3 |
| GO:2001251 | negative regulation of chromosome organization | 0.032963434 | 3 |
| GO:0001776 | leukocyte homeostasis | 0.034755722 | 3 |
| GO:0031341 | regulation of cell killing | 0.035580841 | 3 |
| GO:0009135 | purine nucleoside diphosphate metabolic process | 0.036421455 | 3 |
| GO:0009179 | purine ribonucleoside diphosphate metabolic process | 0.036421455 | 3 |
| GO:0042100 | B cell proliferation | 0.036421455 | 3 |
| GO:0019882 | antigen processing and presentation | 0.039719874 | 3 |
| GO:0002824 | positive regulation of adaptive immune response based on somat | 0.044040591 | 3 |
| GO:0009185 | ribonucleoside diphosphate metabolic process | 0.044040591 | 3 |
| GO:0046634 | regulation of alpha-beta T cell activation | 0.044275583 | 3 |
| GO:0046632 | alpha-beta T cell differentiation | 0.044770518 | 3 |
| GO:0071346 | cellular response to interferon-gamma | 0.044770518 | 3 |
| GO:0032609 | interferon-gamma production | 0.044770518 | 3 |
| GO:0032649 | regulation of interferon-gamma production | 0.044770518 | 3 |
| GO:0002821 | positive regulation of adaptive immune response | 0.044770518 | 3 |
| hsa03030 | DNA replication | 0.007740427 | 3 |
| GO:0071723 | lipopeptide binding | 0.012498222 | 2 |
| GO:0043138 | 3'-5' DNA helicase activity | 0.020094024 | 2 |
| GO:0002327 | immature B cell differentiation | 0.014996223 | 2 |
| GO:0070243 | regulation of thymocyte apoptotic process | 0.016730926 | 2 |
| GO:0002475 | antigen processing and presentation via MHC class Ib | 0.016730926 | 2 |
| GO:0006167 | AMP biosynthetic process | 0.016730926 | 2 |
| GO:0006177 | GMP biosynthetic process | 0.016730926 | 2 |
| GO:1901533 | negative regulation of hematopoietic progenitor cell differentiati | 0.016730926 | 2 |
| GO:1902969 | mitotic DNA replication | 0.016730926 | 2 |
| GO:0009263 | deoxyribonucleotide biosynthetic process | 0.019289599 | 2 |
| GO:0009265 | 2'-deoxyribonucleotide biosynthetic process | 0.019289599 | 2 |
| GO:0046385 | deoxyribose phosphate biosynthetic process | 0.019289599 | 2 |
| GO:0009151 | purine deoxyribonucleotide metabolic process | 0.019812827 | 2 |
| GO:0033151 | V(D)J recombination | 0.020786127 | 2 |
| GO:0046653 | tetrahydrofolate metabolic process | 0.020786127 | 2 |
| GO:0070242 | thymocyte apoptotic process | 0.022420161 | 2 |
| GO:0009168 | purine ribonucleoside monophosphate biosynthetic process | 0.023361999 | 2 |
| GO:0009127 | purine nucleoside monophosphate biosynthetic process | 0.027131836 | 2 |
| GO:0070233 | negative regulation of T cell apoptotic process | 0.031528817 | 2 |
| GO:0009162 | deoxyribonucleoside monophosphate metabolic process | 0.032963434 | 2 |
| GO:0046033 | AMP metabolic process | 0.032963434 | 2 |
| GO:0019883 | antigen processing and presentation of endogenous antigen | 0.034488064 | 2 |
| GO:0035821 | modulation of process of another organism | 0.034488064 | 2 |
| GO:0006760 | folic acid-containing compound metabolic process | 0.035580841 | 2 |
| GO:0033081 | regulation of T cell differentiation in thymus | 0.035580841 | 2 |
| GO:0001916 | positive regulation of T cell mediated cytotoxicity | 0.043967934 | 2 |
| GO:0045070 | positive regulation of viral genome replication | 0.044275583 | 2 |
| GO:0007143 | female meiotic nuclear division | 0.044770518 | 2 |
| GO:0009112 | nucleobase metabolic process | 0.044770518 | 2 |
| GO:0009156 | ribonucleoside monophosphate biosynthetic process | 0.044770518 | 2 |
| GO:0042558 | pteridine-containing compound metabolic process | 0.044770518 | 2 |
| GO:0061081 | positive regulation of myeloid leukocyte cytokine production invo | 0.044770518 | 2 |
| hsa00670 | One carbon pool by folate | 0.049119593 | 2 |

| ID | Description | p.adjust | Count |
|---|---|---|---|
| GO:1903131 | mononuclear cell differentiation | 0.010859529 | 8 |
| HALLMARK_INTERFERON_ALPHA_RESPONSE | HALLMARK_INTERFERON_ALPHA_RESPONSE | 5.20318E-07 | 8 |
| GO:0005525 | GTP binding | 0.00233141 | 7 |
| GO:0019001 | guanyl nucleotide binding | 0.00233141 | 7 |
| GO:0032561 | guanyl ribonucleotide binding | 0.00233141 | 7 |
| GO:0003924 | GTPase activity | 0.005314044 | 6 |
| GO:0051607 | defense response to virus | 0.015140607 | 6 |
| GO:0140546 | defense response to symbiont | 0.015140607 | 6 |
| GO:0019058 | viral life cycle | 0.016465685 | 6 |
| GO:0009615 | response to virus | 0.029516519 | 6 |
| GO:0030098 | lymphocyte differentiation | 0.031260922 | 6 |
| GO:0016032 | viral process | 0.031260922 | 6 |
| HALLMARK_INTERFERON_GAMMA_RESPONSE | HALLMARK_INTERFERON_GAMMA_RESPONSE | 0.006216549 | 6 |
| GO:0030217 | T cell differentiation | 0.040724799 | 5 |
| HALLMARK_INFLAMMATORY_RESPONSE | HALLMARK_INFLAMMATORY_RESPONSE | 0.028825926 | 5 |
| GO:0045071 | negative regulation of viral genome replication | 0.010859529 | 4 |
| GO:0045069 | regulation of viral genome replication | 0.015140607 | 4 |
| GO:0002709 | regulation of T cell mediated immunity | 0.015140607 | 4 |
| GO:0048525 | negative regulation of viral process | 0.015140607 | 4 |
| GO:0019882 | antigen processing and presentation | 0.016465685 | 4 |
| GO:0002456 | T cell mediated immunity | 0.017974214 | 4 |
| GO:0071887 | leukocyte apoptotic process | 0.018968897 | 4 |
| GO:0019079 | viral genome replication | 0.022612385 | 4 |
| GO:1903900 | regulation of viral life cycle | 0.028699587 | 4 |
| GO:0050792 | regulation of viral process | 0.03691881 | 4 |
| GO:0002706 | regulation of lymphocyte mediated immunity | 0.049302866 | 4 |
| GO:0004896 | cytokine receptor activity | 0.044900107 | 3 |
| GO:0035456 | response to interferon-beta | 0.015140607 | 3 |
| GO:0001914 | regulation of T cell mediated cytotoxicity | 0.016465685 | 3 |
| GO:0001913 | T cell mediated cytotoxicity | 0.022612385 | 3 |
| GO:0045058 | T cell selection | 0.022612385 | 3 |
| GO:0009409 | response to cold | 0.022612385 | 3 |
| GO:0002711 | positive regulation of T cell mediated immunity | 0.029204644 | 3 |
| GO:0001965 | G-protein alpha-subunit binding | 0.044900107 | 2 |
| GO:0001671 | ATPase activator activity | 0.044900107 | 2 |

| ID | Description | p.adjust | Count |
|---|---|---|---|
| HALLMARK_INTERFERON_GAMMA_RESPONSE | HALLMARK_INTERFERON_GAMMA_RESPONSE | 5.44375E-13 | 15 |
| GO:0009615 | response to virus | 8.6326E-10 | 13 |
| HALLMARK_INTERFERON_ALPHA_RESPONSE | HALLMARK_INTERFERON_ALPHA_RESPONSE | 4.59825E-14 | 13 |
| GO:0051607 | defense response to virus | 8.5629E-10 | 12 |
| GO:0140546 | defense response to symbiont | 8.5629E-10 | 12 |
| GO:0016032 | viral process | 1.99823E-06 | 10 |
| GO:0034340 | response to type I interferon | 2.43115E-09 | 8 |
| GO:0050792 | regulation of viral process | 3.06238E-07 | 8 |
| GO:0019058 | viral life cycle | 2.16077E-05 | 8 |
| GO:0002831 | regulation of response to biotic stimulus | 4.96501E-05 | 8 |
| hsa05164 | Influenza A | 3.79118E-06 | 8 |
| GO:0071357 | cellular response to type I interferon | 6.44403E-08 | 7 |
| GO:0048525 | negative regulation of viral process | 1.76677E-07 | 7 |
| GO:1903900 | regulation of viral life cycle | 2.17835E-06 | 7 |
| GO:0019221 | cytokine-mediated signaling pathway | 0.003166822 | 7 |
| GO:0045071 | negative regulation of viral genome replication | 3.28815E-07 | 6 |
| GO:0060337 | type I interferon signaling pathway | 1.57479E-06 | 6 |
| GO:0045069 | regulation of viral genome replication | 2.90443E-06 | 6 |
| GO:0032479 | regulation of type I interferon production | 7.7037E-06 | 6 |
| GO:0032606 | type I interferon production | 7.7037E-06 | 6 |
| GO:0019079 | viral genome replication | 2.13125E-05 | 6 |
| GO:0001819 | positive regulation of cytokine production | 0.014473982 | 6 |
| hsa05160 | Hepatitis C | 0.000387166 | 6 |
| GO:0032608 | interferon-beta production | 9.67425E-06 | 5 |
| GO:0032648 | regulation of interferon-beta production | 9.67425E-06 | 5 |
| GO:0032481 | positive regulation of type I interferon production | 1.51358E-05 | 5 |
| GO:0002221 | pattern recognition receptor signaling pathway | 0.001861567 | 5 |
| hsa05162 | Measles | 0.001973205 | 5 |
| hsa05169 | Epstein-Barr virus infection | 0.008814927 | 5 |
| hsa05171 | Coronavirus disease - COVID-19 | 0.013257485 | 5 |
| GO:0035456 | response to interferon-beta | 3.44011E-05 | 4 |
| GO:0032728 | positive regulation of interferon-beta production | 7.07312E-05 | 4 |
| GO:0060338 | regulation of type I interferon-mediated signaling pathway | 8.30326E-05 | 4 |
| GO:0098586 | cellular response to virus | 0.003861186 | 4 |
| GO:0002224 | toll-like receptor signaling pathway | 0.004689699 | 4 |
| GO:0001959 | regulation of cytokine-mediated signaling pathway | 0.013012536 | 4 |
| GO:0060759 | regulation of response to cytokine stimulus | 0.014473982 | 4 |
| GO:0045088 | regulation of innate immune response | 0.035528359 | 4 |
| GO:0060339 | negative regulation of type I interferon-mediated signaling pathv | 0.000448761 | 3 |
| GO:0140374 | antiviral innate immune response | 0.000580174 | 3 |
| GO:0032727 | positive regulation of interferon-alpha production | 0.000640546 | 3 |
| GO:0032607 | interferon-alpha production | 0.001215866 | 3 |
| GO:0032647 | regulation of interferon-alpha production | 0.001215866 | 3 |
| GO:0043112 | receptor metabolic process | 0.009118427 | 3 |
| GO:0070936 | protein K48-linked ubiquitination | 0.012212077 | 3 |
| GO:0050688 | regulation of defense response to virus | 0.013741966 | 3 |
| GO:0045824 | negative regulation of innate immune response | 0.014473982 | 3 |
| GO:0001960 | negative regulation of cytokine-mediated signaling pathway | 0.021433106 | 3 |
| GO:0019646 | aerobic electron transport chain | 0.023811371 | 3 |
| GO:0060761 | negative regulation of response to cytokine stimulus | 0.024035991 | 3 |
| GO:0042773 | ATP synthesis coupled electron transport | 0.027788652 | 3 |
| GO:0042775 | mitochondrial ATP synthesis coupled electron transport | 0.027788652 | 3 |
| GO:0062207 | regulation of pattern recognition receptor signaling pathway | 0.032149904 | 3 |
| GO:0002832 | negative regulation of response to biotic stimulus | 0.038240705 | 3 |
| GO:0022904 | respiratory electron transport chain | 0.042377058 | 3 |
| GO:0039530 | MDA-5 signaling pathway | 0.007542382 | 2 |
| GO:0034154 | toll-like receptor 7 signaling pathway | 0.008772278 | 2 |
| GO:0009263 | deoxyribonucleotide biosynthetic process | 0.013480929 | 2 |
| GO:0009265 | 2'-deoxyribonucleotide biosynthetic process | 0.013480929 | 2 |
| GO:0046385 | deoxyribose phosphate biosynthetic process | 0.013480929 | 2 |
| GO:0035458 | cellular response to interferon-beta | 0.025548944 | 2 |
| GO:0002230 | positive regulation of defense response to virus by host | 0.038240705 | 2 |
| GO:0006221 | pyrimidine nucleotide biosynthetic process | 0.03962197 | 2 |
| GO:0039528 | cytoplasmic pattern recognition receptor signaling pathway in re | 0.042377058 | 2 |
| GO:0045648 | positive regulation of erythrocyte differentiation | 0.042377058 | 2 |
| GO:0072528 | pyrimidine-containing compound biosynthetic process | 0.049247826 | 2 |

| ID | Description | p.adjust | Count |
|---|---|---|---|
| GO:0002443 | leukocyte mediated immunity | 1.09972E-05 | 11 |
| GO:0002366 | leukocyte activation involved in immune response | 1.58151E-05 | 9 |
| GO:0002263 | cell activation involved in immune response | 1.58151E-05 | 9 |
| GO:0002697 | regulation of immune effector process | 6.78929E-05 | 9 |
| GO:0002274 | myeloid leukocyte activation | 3.45989E-05 | 8 |
| GO:0002703 | regulation of leukocyte mediated immunity | 3.45989E-05 | 8 |
| GO:0002699 | positive regulation of immune effector process | 5.64915E-05 | 8 |
| GO:1903037 | regulation of leukocyte cell-cell adhesion | 0.000441781 | 8 |
| GO:0007159 | leukocyte cell-cell adhesion | 0.000630158 | 8 |
| GO:0002696 | positive regulation of leukocyte activation | 0.000992221 | 8 |
| GO:0050867 | positive regulation of cell activation | 0.001172994 | 8 |
| GO:0022407 | regulation of cell-cell adhesion | 0.001429016 | 8 |
| GO:0002275 | myeloid cell activation involved in immune response | 7.80682E-06 | 7 |
| GO:0002444 | myeloid leukocyte mediated immunity | 8.35843E-06 | 7 |
| GO:0042113 | B cell activation | 0.00133149 | 7 |
| GO:0050863 | regulation of T cell activation | 0.001937095 | 7 |
| GO:0050727 | regulation of inflammatory response | 0.003216575 | 7 |
| GO:0032103 | positive regulation of response to external stimulus | 0.005285671 | 7 |
| GO:1903131 | mononuclear cell differentiation | 0.005835909 | 7 |
| hsa04145 | Phagosome | 0.000594655 | 7 |
| hsa05132 | Salmonella infection | 0.004542277 | 7 |
| GO:0043299 | leukocyte degranulation | 1.58151E-05 | 6 |
| GO:0001906 | cell killing | 0.000608317 | 6 |
| GO:0045055 | regulated exocytosis | 0.001489045 | 6 |
| GO:0070663 | regulation of leukocyte proliferation | 0.002485198 | 6 |
| GO:1903039 | positive regulation of leukocyte cell-cell adhesion | 0.002740455 | 6 |
| GO:0044403 | biological process involved in symbiotic interaction | 0.004677813 | 6 |
| GO:1902105 | regulation of leukocyte differentiation | 0.004913638 | 6 |
| GO:0022409 | positive regulation of cell-cell adhesion | 0.005246023 | 6 |
| GO:0070661 | leukocyte proliferation | 0.007273909 | 6 |
| GO:0006887 | exocytosis | 0.009450054 | 6 |
| GO:0071900 | regulation of protein serine/threonine kinase activity | 0.012135333 | 6 |
| GO:0051251 | positive regulation of lymphocyte activation | 0.012589874 | 6 |
| GO:1903706 | regulation of hemopoiesis | 0.013948925 | 6 |
| GO:0030098 | lymphocyte differentiation | 0.014201309 | 6 |
| GO:0002683 | negative regulation of immune system process | 0.018877313 | 6 |
| GO:0045785 | positive regulation of cell adhesion | 0.020982968 | 6 |
| GO:0001819 | positive regulation of cytokine production | 0.021105687 | 6 |
| GO:0042277 | peptide binding | 0.025319504 | 5 |
| GO:0033218 | amide binding | 0.04428418 | 5 |
| GO:0002886 | regulation of myeloid leukocyte mediated immunity | 6.78929E-05 | 5 |
| GO:0048002 | antigen processing and presentation of peptide antigen | 8.33098E-05 | 5 |
| GO:0032418 | lysosome localization | 0.000243363 | 5 |
| GO:1990849 | vacuolar localization | 0.000243363 | 5 |
| GO:0019882 | antigen processing and presentation | 0.000608317 | 5 |
| GO:0002705 | positive regulation of leukocyte mediated immunity | 0.00133149 | 5 |
| GO:0002819 | regulation of adaptive immune response | 0.004874052 | 5 |
| GO:0050670 | regulation of lymphocyte proliferation | 0.009539558 | 5 |
| GO:0032944 | regulation of mononuclear cell proliferation | 0.010126246 | 5 |
| GO:0050870 | positive regulation of T cell activation | 0.011197369 | 5 |
| GO:0046651 | lymphocyte proliferation | 0.019879681 | 5 |
| GO:0031349 | positive regulation of defense response | 0.020235278 | 5 |
| GO:0032943 | mononuclear cell proliferation | 0.020652837 | 5 |
| GO:0006909 | phagocytosis | 0.020652837 | 5 |
| GO:0016236 | macroautophagy | 0.020652837 | 5 |
| GO:0031647 | regulation of protein stability | 0.020982968 | 5 |
| GO:0051235 | maintenance of location | 0.023205349 | 5 |
| GO:0002831 | regulation of response to biotic stimulus | 0.026145855 | 5 |
| GO:0002449 | lymphocyte mediated immunity | 0.026547506 | 5 |
| GO:0070997 | neuron death | 0.026554488 | 5 |
| GO:0002460 | adaptive immune response based on somatic recombination of im | 0.027871974 | 5 |
| GO:0002253 | activation of immune response | 0.030635881 | 5 |
| GO:0050900 | leukocyte migration | 0.030794765 | 5 |
| GO:0032102 | negative regulation of response to external stimulus | 0.038067379 | 5 |
| hsa04066 | HIF-1 signaling pathway | 0.004542277 | 5 |
| hsa05418 | Fluid shear stress and atherosclerosis | 0.008057358 | 5 |
| hsa05225 | Hepatocellular carcinoma | 0.016823546 | 5 |

| | | | |
|---|---|---|---|
| hsa05130 | Pathogenic Escherichia coli infection | 0.031255072 | 5 |
| hsa05208 | Chemical carcinogenesis - reactive oxygen species | 0.039504547 | 5 |
| hsa04810 | Regulation of actin cytoskeleton | 0.039504547 | 5 |
| GO:0016829 | lyase activity | 0.025717459 | 4 |
| GO:0002495 | antigen processing and presentation of peptide antigen via MHC | 0.000208047 | 4 |
| GO:0002504 | antigen processing and presentation of peptide or polysaccharide | 0.000243363 | 4 |
| GO:0042119 | neutrophil activation | 0.00034281 | 4 |
| GO:0036230 | granulocyte activation | 0.000534092 | 4 |
| GO:0043300 | regulation of leukocyte degranulation | 0.000571833 | 4 |
| GO:0043303 | mast cell degranulation | 0.000630158 | 4 |
| GO:0002279 | mast cell activation involved in immune response | 0.00068286 | 4 |
| GO:0002448 | mast cell mediated immunity | 0.000707795 | 4 |
| GO:0032613 | interleukin-10 production | 0.001172994 | 4 |
| GO:0032653 | regulation of interleukin-10 production | 0.001172994 | 4 |
| GO:0030888 | regulation of B cell proliferation | 0.001234675 | 4 |
| GO:0045576 | mast cell activation | 0.001234675 | 4 |
| GO:0031341 | regulation of cell killing | 0.004733942 | 4 |
| GO:0048144 | fibroblast proliferation | 0.004790382 | 4 |
| GO:0042100 | B cell proliferation | 0.004790382 | 4 |
| GO:0002821 | positive regulation of adaptive immune response | 0.007264192 | 4 |
| GO:1903305 | regulation of regulated secretory pathway | 0.012269469 | 4 |
| GO:0030183 | B cell differentiation | 0.013835072 | 4 |
| GO:0050729 | positive regulation of inflammatory response | 0.013948925 | 4 |
| GO:0016052 | carbohydrate catabolic process | 0.014046903 | 4 |
| GO:0097530 | granulocyte migration | 0.014394605 | 4 |
| GO:0007254 | JNK cascade | 0.018877313 | 4 |
| GO:1905475 | regulation of protein localization to membrane | 0.018877313 | 4 |
| GO:0002706 | regulation of lymphocyte mediated immunity | 0.020347742 | 4 |
| GO:0002822 | regulation of adaptive immune response based on somatic recoml | 0.020652837 | 4 |
| GO:1902107 | positive regulation of leukocyte differentiation | 0.020652837 | 4 |
| GO:1903708 | positive regulation of hemopoiesis | 0.020652837 | 4 |
| GO:0043433 | negative regulation of DNA-binding transcription factor activity | 0.020982968 | 4 |
| GO:0050777 | negative regulation of immune response | 0.020982968 | 4 |
| GO:0050864 | regulation of B cell activation | 0.023205349 | 4 |
| GO:0017157 | regulation of exocytosis | 0.025453065 | 4 |
| GO:0006469 | negative regulation of protein kinase activity | 0.026145855 | 4 |
| GO:0046034 | ATP metabolic process | 0.026145855 | 4 |
| GO:0043491 | protein kinase B signaling | 0.026145855 | 4 |
| GO:0050866 | negative regulation of cell activation | 0.026547506 | 4 |
| GO:0051651 | maintenance of location in cell | 0.027842883 | 4 |
| GO:0019318 | hexose metabolic process | 0.028440664 | 4 |
| GO:0051403 | stress-activated MAPK cascade | 0.030635881 | 4 |
| GO:0033673 | negative regulation of kinase activity | 0.030635881 | 4 |
| GO:0045088 | regulation of innate immune response | 0.030635881 | 4 |
| GO:0009205 | purine ribonucleoside triphosphate metabolic process | 0.030635881 | 4 |
| GO:0097529 | myeloid leukocyte migration | 0.030635881 | 4 |
| GO:0031098 | stress-activated protein kinase signaling cascade | 0.031398788 | 4 |
| GO:0009144 | purine nucleoside triphosphate metabolic process | 0.031684187 | 4 |
| GO:0005996 | monosaccharide metabolic process | 0.031813193 | 4 |
| GO:0009199 | ribonucleoside triphosphate metabolic process | 0.031813193 | 4 |
| GO:0043122 | regulation of I-kappaB kinase/NF-kappaB signaling | 0.033521356 | 4 |
| GO:0009141 | nucleoside triphosphate metabolic process | 0.036825919 | 4 |
| GO:0051348 | negative regulation of transferase activity | 0.039821911 | 4 |
| GO:0007249 | I-kappaB kinase/NF-kappaB signaling | 0.044133083 | 4 |
| hsa00480 | Glutathione metabolism | 0.004542277 | 4 |
| hsa05416 | Viral myocarditis | 0.004542277 | 4 |
| hsa01230 | Biosynthesis of amino acids | 0.008057358 | 4 |
| hsa04650 | Natural killer cell mediated cytotoxicity | 0.039504547 | 4 |
| hsa04210 | Apoptosis | 0.039504547 | 4 |
| GO:0004602 | glutathione peroxidase activity | 0.003518706 | 3 |
| GO:0019865 | immunoglobulin binding | 0.003518706 | 3 |
| GO:0004601 | peroxidase activity | 0.013253744 | 3 |
| GO:0016684 | oxidoreductase activity, acting on peroxide as acceptor | 0.013253744 | 3 |
| GO:0016765 | transferase activity, transferring alkyl or aryl (other than methyl | 0.013253744 | 3 |
| GO:0016209 | antioxidant activity | 0.023633293 | 3 |
| GO:0005200 | structural constituent of cytoskeleton | 0.03295471 | 3 |
| GO:0031490 | chromatin DNA binding | 0.041153396 | 3 |
| GO:1902563 | regulation of neutrophil activation | 0.000564686 | 3 |

| | | | |
|---|---|---|---|
| GO:0030889 | negative regulation of B cell proliferation | 0.000852749 | 3 |
| GO:0019886 | antigen processing and presentation of exogenous peptide antige | 0.002451365 | 3 |
| GO:0050869 | negative regulation of B cell activation | 0.003876483 | 3 |
| GO:0002478 | antigen processing and presentation of exogenous peptide antige | 0.004790382 | 3 |
| GO:0031640 | killing of cells of another organism | 0.006474268 | 3 |
| GO:0019884 | antigen processing and presentation of exogenous antigen | 0.007410808 | 3 |
| GO:0006749 | glutathione metabolic process | 0.010511911 | 3 |
| GO:0031343 | positive regulation of cell killing | 0.013948925 | 3 |
| GO:0031638 | zymogen activation | 0.014046903 | 3 |
| GO:0002704 | negative regulation of leukocyte mediated immunity | 0.014297182 | 3 |
| GO:0032732 | positive regulation of interleukin-1 production | 0.017840927 | 3 |
| GO:0006096 | glycolytic process | 0.020652837 | 3 |
| GO:0006757 | ATP generation from ADP | 0.020652837 | 3 |
| GO:0001910 | regulation of leukocyte mediated cytotoxicity | 0.020982968 | 3 |
| GO:0048145 | regulation of fibroblast proliferation | 0.020982968 | 3 |
| GO:0050672 | negative regulation of lymphocyte proliferation | 0.020982968 | 3 |
| GO:0032945 | negative regulation of mononuclear cell proliferation | 0.021105687 | 3 |
| GO:0002709 | regulation of T cell mediated immunity | 0.021611093 | 3 |
| GO:1901216 | positive regulation of neuron death | 0.02262738 | 3 |
| GO:0046031 | ADP metabolic process | 0.02262738 | 3 |
| GO:0032088 | negative regulation of NF-kappaB transcription factor activity | 0.023205349 | 3 |
| GO:0070664 | negative regulation of leukocyte proliferation | 0.023725164 | 3 |
| GO:0006165 | nucleoside diphosphate phosphorylation | 0.025453065 | 3 |
| GO:0098869 | cellular oxidant detoxification | 0.025453065 | 3 |
| GO:0046939 | nucleotide phosphorylation | 0.026145855 | 3 |
| GO:0050764 | regulation of phagocytosis | 0.026145855 | 3 |
| GO:1903076 | regulation of protein localization to plasma membrane | 0.026145855 | 3 |
| GO:0009135 | purine nucleoside diphosphate metabolic process | 0.026332808 | 3 |
| GO:0009179 | purine ribonucleoside diphosphate metabolic process | 0.026332808 | 3 |
| GO:0032760 | positive regulation of tumor necrosis factor production | 0.026547506 | 3 |
| GO:0006754 | ATP biosynthetic process | 0.026698734 | 3 |
| GO:0030593 | neutrophil chemotaxis | 0.026698734 | 3 |
| GO:0006090 | pyruvate metabolic process | 0.027842883 | 3 |
| GO:0042116 | macrophage activation | 0.027975659 | 3 |
| GO:1903557 | positive regulation of tumor necrosis factor superfamily cytokine | 0.027975659 | 3 |
| GO:0045445 | myoblast differentiation | 0.02964152 | 3 |
| GO:0002456 | T cell mediated immunity | 0.029835434 | 3 |
| GO:0002824 | positive regulation of adaptive immune response based on somat | 0.030215633 | 3 |
| GO:0009185 | ribonucleoside diphosphate metabolic process | 0.030215633 | 3 |
| GO:1990748 | cellular detoxification | 0.030463139 | 3 |
| GO:0051702 | biological process involved in interaction with symbiont | 0.030635881 | 3 |
| GO:0000079 | regulation of cyclin-dependent protein serine/threonine kinase ac | 0.030635881 | 3 |
| GO:0009206 | purine ribonucleoside triphosphate biosynthetic process | 0.030635881 | 3 |
| GO:0009145 | purine nucleoside triphosphate biosynthetic process | 0.030635881 | 3 |
| GO:0002708 | positive regulation of lymphocyte mediated immunity | 0.030912056 | 3 |
| GO:0051897 | positive regulation of protein kinase B signaling | 0.031303369 | 3 |
| GO:1904029 | regulation of cyclin-dependent protein kinase activity | 0.031303369 | 3 |
| GO:0002698 | negative regulation of immune effector process | 0.031813193 | 3 |
| GO:0071901 | negative regulation of protein serine/threonine kinase activity | 0.031813193 | 3 |
| GO:0009201 | ribonucleoside triphosphate biosynthetic process | 0.03198415 | 3 |
| GO:0097237 | cellular response to toxic substance | 0.03198415 | 3 |
| GO:1904375 | regulation of protein localization to cell periphery | 0.033118687 | 3 |
| GO:0006690 | icosanoid metabolic process | 0.033521356 | 3 |
| GO:0006911 | phagocytosis, engulfment | 0.033960155 | 3 |
| GO:0032612 | interleukin-1 production | 0.035150706 | 3 |
| GO:0032652 | regulation of interleukin-1 production | 0.035150706 | 3 |
| GO:0050853 | B cell receptor signaling pathway | 0.035150706 | 3 |
| GO:0071621 | granulocyte chemotaxis | 0.035150706 | 3 |
| GO:1990266 | neutrophil migration | 0.035150706 | 3 |
| GO:0009132 | nucleoside diphosphate metabolic process | 0.036500291 | 3 |
| GO:0001909 | leukocyte mediated cytotoxicity | 0.036825919 | 3 |
| GO:0009142 | nucleoside triphosphate biosynthetic process | 0.036825919 | 3 |
| GO:0099024 | plasma membrane invagination | 0.038067379 | 3 |
| GO:0030833 | regulation of actin filament polymerization | 0.040243651 | 3 |
| GO:0046328 | regulation of JNK cascade | 0.040492159 | 3 |
| GO:0010324 | membrane invagination | 0.04181284 | 3 |
| GO:0014065 | phosphatidylinositol 3-kinase signaling | 0.043165375 | 3 |
| GO:0046718 | viral entry into host cell | 0.046218665 | 3 |

| | | | |
|---|---|---|---|
| GO:1903531 | negative regulation of secretion by cell | 0.046218665 | 3 |
| GO:0098754 | detoxification | 0.048511083 | 3 |
| hsa00010 | Glycolysis / Gluconeogenesis | 0.039504547 | 3 |
| hsa01524 | Platinum drug resistance | 0.047028814 | 3 |
| GO:0016303 | 1-phosphatidylinositol-3-kinase activity | 0.013253744 | 2 |
| GO:0019864 | IgG binding | 0.013253744 | 2 |
| GO:0035004 | phosphatidylinositol 3-kinase activity | 0.013253744 | 2 |
| GO:0016307 | phosphatidylinositol phosphate kinase activity | 0.018858128 | 2 |
| GO:0003756 | protein disulfide isomerase activity | 0.019607202 | 2 |
| GO:0016864 | intramolecular oxidoreductase activity, transposing S-S bonds | 0.019607202 | 2 |
| GO:0052742 | phosphatidylinositol kinase activity | 0.020054435 | 2 |
| GO:0004364 | glutathione transferase activity | 0.028256171 | 2 |
| GO:0003785 | actin monomer binding | 0.03295471 | 2 |
| GO:0042989 | sequestering of actin monomers | 0.009370453 | 2 |
| GO:0002887 | negative regulation of myeloid leukocyte mediated immunity | 0.010511911 | 2 |
| GO:2001198 | regulation of dendritic cell differentiation | 0.011840275 | 2 |
| GO:0043301 | negative regulation of leukocyte degranulation | 0.012819613 | 2 |
| GO:2001171 | positive regulation of ATP biosynthetic process | 0.012819613 | 2 |
| GO:0002283 | neutrophil activation involved in immune response | 0.019497301 | 2 |
| GO:0002923 | regulation of humoral immune response mediated by circulating i | 0.019497301 | 2 |
| GO:0032930 | positive regulation of superoxide anion generation | 0.019497301 | 2 |
| GO:0002577 | regulation of antigen processing and presentation | 0.02045243 | 2 |
| GO:0002888 | positive regulation of myeloid leukocyte mediated immunity | 0.02045243 | 2 |
| GO:0036092 | phosphatidylinositol-3-phosphate biosynthetic process | 0.020652837 | 2 |
| GO:0015874 | norepinephrine transport | 0.020982968 | 2 |
| GO:0032928 | regulation of superoxide anion generation | 0.020982968 | 2 |
| GO:0050765 | negative regulation of phagocytosis | 0.020982968 | 2 |
| GO:2001169 | regulation of ATP biosynthetic process | 0.021679392 | 2 |
| GO:0002922 | positive regulation of humoral immune response | 0.02262738 | 2 |
| GO:0018279 | protein N-linked glycosylation via asparagine | 0.02262738 | 2 |
| GO:0050855 | regulation of B cell receptor signaling pathway | 0.02262738 | 2 |
| GO:0001911 | negative regulation of leukocyte mediated cytotoxicity | 0.023205349 | 2 |
| GO:0018196 | peptidyl-asparagine modification | 0.023205349 | 2 |
| GO:0032693 | negative regulation of interleukin-10 production | 0.023205349 | 2 |
| GO:0043302 | positive regulation of leukocyte degranulation | 0.023205349 | 2 |
| GO:0046716 | muscle cell cellular homeostasis | 0.023205349 | 2 |
| GO:0030810 | positive regulation of nucleotide biosynthetic process | 0.023960867 | 2 |
| GO:0051238 | sequestering of metal ion | 0.023960867 | 2 |
| GO:1900373 | positive regulation of purine nucleotide biosynthetic process | 0.023960867 | 2 |
| GO:1903306 | negative regulation of regulated secretory pathway | 0.023960867 | 2 |
| GO:0031342 | negative regulation of cell killing | 0.026145855 | 2 |
| GO:0002313 | mature B cell differentiation involved in immune response | 0.028440664 | 2 |
| GO:0002862 | negative regulation of inflammatory response to antigenic stimul | 0.0297258 | 2 |
| GO:0043304 | regulation of mast cell degranulation | 0.0297258 | 2 |
| GO:0048147 | negative regulation of fibroblast proliferation | 0.030463139 | 2 |
| GO:0050858 | negative regulation of antigen receptor-mediated signaling pathv | 0.030463139 | 2 |
| GO:0033006 | regulation of mast cell activation involved in immune response | 0.030635881 | 2 |
| GO:0090322 | regulation of superoxide metabolic process | 0.030635881 | 2 |
| GO:1903580 | positive regulation of ATP metabolic process | 0.031813193 | 2 |
| GO:0002335 | mature B cell differentiation | 0.033056448 | 2 |
| GO:0045920 | negative regulation of exocytosis | 0.033960155 | 2 |
| GO:0045730 | respiratory burst | 0.035150706 | 2 |
| GO:0001914 | regulation of T cell mediated cytotoxicity | 0.036825919 | 2 |
| GO:0014912 | negative regulation of smooth muscle cell migration | 0.036825919 | 2 |
| GO:0032733 | positive regulation of interleukin-10 production | 0.036825919 | 2 |
| GO:0045454 | cell redox homeostasis | 0.036825919 | 2 |
| GO:0050999 | regulation of nitric-oxide synthase activity | 0.038067379 | 2 |
| GO:0042554 | superoxide anion generation | 0.039492914 | 2 |
| GO:1900371 | regulation of purine nucleotide biosynthetic process | 0.039492914 | 2 |
| GO:0021762 | substantia nigra development | 0.040492159 | 2 |
| GO:0030808 | regulation of nucleotide biosynthetic process | 0.040492159 | 2 |
| GO:0033003 | regulation of mast cell activation | 0.04181284 | 2 |
| GO:0002920 | regulation of humoral immune response | 0.043256981 | 2 |
| GO:0001774 | microglial cell activation | 0.044133083 | 2 |
| GO:0002861 | regulation of inflammatory response to antigenic stimulus | 0.044133083 | 2 |
| GO:0090199 | regulation of release of cytochrome c from mitochondria | 0.044133083 | 2 |
| GO:0097028 | dendritic cell differentiation | 0.044133083 | 2 |
| GO:0045981 | positive regulation of nucleotide metabolic process | 0.045583196 | 2 |

| | | | |
|---|---|---|---|
| GO:1900544 | positive regulation of purine nucleotide metabolic process | 0.045583196 | 2 |
| GO:0002269 | leukocyte activation involved in inflammatory response | 0.048511083 | 2 |
| GO:0001913 | T cell mediated cytotoxicity | 0.049988042 | 2 |
| GO:1903307 | positive regulation of regulated secretory pathway | 0.049988042 | 2 |

| ID | Description | p.adjust | Count |
|---|---|---|---|
| GO:0007159 | leukocyte cell-cell adhesion | 0.000971286 | 9 |
| GO:0032103 | positive regulation of response to external stimulus | 0.011490616 | 7 |
| hsa05415 | Diabetic cardiomyopathy | 0.000555401 | 7 |
| hsa05020 | Prion disease | 0.001309624 | 7 |
| GO:0022900 | electron transport chain | 0.000971286 | 6 |
| GO:1903039 | positive regulation of leukocyte cell-cell adhesion | 0.006645669 | 6 |
| GO:0060326 | cell chemotaxis | 0.011490616 | 6 |
| GO:0022409 | positive regulation of cell-cell adhesion | 0.011490616 | 6 |
| GO:0051235 | maintenance of location | 0.014450902 | 6 |
| GO:0050863 | regulation of T cell activation | 0.017768799 | 6 |
| GO:1903037 | regulation of leukocyte cell-cell adhesion | 0.017768799 | 6 |
| GO:0042742 | defense response to bacterium | 0.018244914 | 6 |
| GO:0050900 | leukocyte migration | 0.01964904 | 6 |
| GO:0006875 | cellular metal ion homeostasis | 0.023701474 | 6 |
| GO:0007015 | actin filament organization | 0.027089506 | 6 |
| GO:0002443 | leukocyte mediated immunity | 0.027089506 | 6 |
| GO:0045785 | positive regulation of cell adhesion | 0.030164588 | 6 |
| GO:0022407 | regulation of cell-cell adhesion | 0.030974296 | 6 |
| GO:0030003 | cellular cation homeostasis | 0.032251358 | 6 |
| hsa04145 | Phagosome | 0.000555401 | 6 |
| hsa05208 | Chemical carcinogenesis - reactive oxygen species | 0.00272177 | 6 |
| hsa05012 | Parkinson disease | 0.005114009 | 6 |
| hsa05014 | Amyotrophic lateral sclerosis | 0.018848007 | 6 |
| hsa05022 | Pathways of neurodegeneration - multiple diseases | 0.048186758 | 6 |
| GO:0019646 | aerobic electron transport chain | 0.000971286 | 5 |
| GO:0042773 | ATP synthesis coupled electron transport | 0.000971286 | 5 |
| GO:0042775 | mitochondrial ATP synthesis coupled electron transport | 0.000971286 | 5 |
| GO:0022904 | respiratory electron transport chain | 0.001899824 | 5 |
| GO:0006119 | oxidative phosphorylation | 0.003819151 | 5 |
| GO:0009060 | aerobic respiration | 0.011490616 | 5 |
| GO:0051651 | maintenance of location in cell | 0.015915153 | 5 |
| GO:0030595 | leukocyte chemotaxis | 0.017768799 | 5 |
| GO:0045333 | cellular respiration | 0.017768799 | 5 |
| GO:0050870 | positive regulation of T cell activation | 0.019085141 | 5 |
| GO:0051924 | regulation of calcium ion transport | 0.021694926 | 5 |
| GO:0031349 | positive regulation of defense response | 0.027089506 | 5 |
| GO:0002440 | production of molecular mediator of immune response | 0.03158345 | 5 |
| GO:0015980 | energy derivation by oxidation of organic compounds | 0.03158345 | 5 |
| GO:0002449 | lymphocyte mediated immunity | 0.039452143 | 5 |
| GO:0051251 | positive regulation of lymphocyte activation | 0.048406238 | 5 |
| hsa05416 | Viral myocarditis | 0.000437169 | 5 |
| hsa00190 | Oxidative phosphorylation | 0.002391683 | 5 |
| hsa04932 | Non-alcoholic fatty liver disease | 0.003529488 | 5 |
| hsa04514 | Cell adhesion molecules | 0.00355884 | 5 |
| hsa05169 | Epstein-Barr virus infection | 0.00880645 | 5 |
| hsa05166 | Human T-cell leukemia virus 1 infection | 0.012590356 | 5 |
| hsa04714 | Thermogenesis | 0.014422509 | 5 |
| hsa05016 | Huntington disease | 0.036672893 | 5 |
| GO:0090079 | translation regulator activity, nucleic acid binding | 0.012550845 | 4 |
| GO:0009055 | electron transfer activity | 0.012550845 | 4 |
| GO:0045730 | respiratory burst | 0.000971286 | 4 |
| GO:0002478 | antigen processing and presentation of exogenous peptide antige | 0.000971286 | 4 |
| GO:0019884 | antigen processing and presentation of exogenous antigen | 0.001077299 | 4 |
| GO:0048002 | antigen processing and presentation of peptide antigen | 0.002513347 | 4 |
| GO:0019882 | antigen processing and presentation | 0.011490616 | 4 |
| GO:0050830 | defense response to Gram-positive bacterium | 0.014450902 | 4 |
| GO:0045727 | positive regulation of translation | 0.01964904 | 4 |
| GO:0070665 | positive regulation of leukocyte proliferation | 0.027089506 | 4 |
| GO:0034250 | positive regulation of cellular amide metabolic process | 0.027089506 | 4 |
| GO:0042129 | regulation of T cell proliferation | 0.027089506 | 4 |
| GO:0097553 | calcium ion transmembrane import into cytosol | 0.027089506 | 4 |
| GO:1903169 | regulation of calcium ion transmembrane transport | 0.029886215 | 4 |
| GO:0007204 | positive regulation of cytosolic calcium ion concentration | 0.035020077 | 4 |
| GO:0042098 | T cell proliferation | 0.037257728 | 4 |
| GO:0002377 | immunoglobulin production | 0.039452143 | 4 |
| GO:0050670 | regulation of lymphocyte proliferation | 0.047985691 | 4 |
| GO:0032944 | regulation of mononuclear cell proliferation | 0.048406238 | 4 |

| | | | |
|---|---|---|---|
| hsa05330 | Allograft rejection | 0.000555401 | 4 |
| hsa05332 | Graft-versus-host disease | 0.000555401 | 4 |
| hsa04940 | Type I diabetes mellitus | 0.000555401 | 4 |
| hsa05320 | Autoimmune thyroid disease | 0.000943842 | 4 |
| hsa04612 | Antigen processing and presentation | 0.00272177 | 4 |
| hsa05170 | Human immunodeficiency virus 1 infection | 0.048186758 | 4 |
| GO:0042605 | peptide antigen binding | 0.012550845 | 3 |
| GO:0015453 | oxidoreduction-driven active transmembrane transporter activity | 0.029099554 | 3 |
| GO:0008135 | translation factor activity, RNA binding | 0.0393368 | 3 |
| GO:0016651 | oxidoreductase activity, acting on NAD(P)H | 0.0393368 | 3 |
| GO:0042743 | hydrogen peroxide metabolic process | 0.017768799 | 3 |
| GO:0019731 | antibacterial humoral response | 0.021383737 | 3 |
| GO:0042267 | natural killer cell mediated cytotoxicity | 0.027089506 | 3 |
| GO:0002228 | natural killer cell mediated immunity | 0.027089506 | 3 |
| GO:0030593 | neutrophil chemotaxis | 0.039782217 | 3 |
| GO:0042102 | positive regulation of T cell proliferation | 0.039782217 | 3 |
| GO:0046916 | cellular transition metal ion homeostasis | 0.048139061 | 3 |
| hsa04672 | Intestinal immune network for IgA production | 0.007525748 | 3 |
| hsa05140 | Leishmaniasis | 0.020728369 | 3 |
| hsa04260 | Cardiac muscle contraction | 0.027821011 | 3 |
| hsa04670 | Leukocyte transendothelial migration | 0.048186758 | 3 |
| GO:0042608 | T cell receptor binding | 0.012550845 | 2 |
| GO:0050786 | RAGE receptor binding | 0.012550845 | 2 |
| GO:0070486 | leukocyte aggregation | 0.01964904 | 2 |
| GO:0002399 | MHC class II protein complex assembly | 0.025584544 | 2 |
| GO:0002503 | peptide antigen assembly with MHC class II protein complex | 0.025584544 | 2 |
| GO:0019885 | antigen processing and presentation of endogenous peptide antig | 0.027089506 | 2 |
| GO:0010522 | regulation of calcium ion transport into cytosol | 0.027089506 | 2 |
| GO:0002483 | antigen processing and presentation of endogenous peptide antig | 0.027089506 | 2 |
| GO:0045953 | negative regulation of natural killer cell mediated cytotoxicity | 0.027089506 | 2 |
| GO:2001185 | regulation of CD8-positive, alpha-beta T cell activation | 0.027089506 | 2 |
| GO:0002396 | MHC protein complex assembly | 0.027089506 | 2 |
| GO:0002501 | peptide antigen assembly with MHC protein complex | 0.027089506 | 2 |
| GO:0002716 | negative regulation of natural killer cell mediated immunity | 0.027089506 | 2 |
| GO:0032069 | regulation of nuclease activity | 0.030974296 | 2 |
| GO:0006123 | mitochondrial electron transport, cytochrome c to oxygen | 0.033884172 | 2 |
| GO:0001911 | negative regulation of leukocyte mediated cytotoxicity | 0.035020077 | 2 |
| GO:0046641 | positive regulation of alpha-beta T cell proliferation | 0.035020077 | 2 |
| GO:0019883 | antigen processing and presentation of endogenous antigen | 0.036711646 | 2 |
| GO:0051238 | sequestering of metal ion | 0.036711646 | 2 |
| GO:0036037 | CD8-positive, alpha-beta T cell activation | 0.038402875 | 2 |
| GO:0031342 | negative regulation of cell killing | 0.039452143 | 2 |
| GO:0033119 | negative regulation of RNA splicing | 0.039452143 | 2 |
| GO:0010592 | positive regulation of lamellipodium assembly | 0.039782217 | 2 |
| GO:0060402 | calcium ion transport into cytosol | 0.039782217 | 2 |
| GO:0002474 | antigen processing and presentation of peptide antigen via MHC | 0.041211972 | 2 |
| GO:0019886 | antigen processing and presentation of exogenous peptide antige | 0.041211972 | 2 |
| GO:0001916 | positive regulation of T cell mediated cytotoxicity | 0.043419145 | 2 |
| GO:0002495 | antigen processing and presentation of peptide antigen via MHC | 0.048406238 | 2 |
| GO:0045648 | positive regulation of erythrocyte differentiation | 0.048406238 | 2 |
| hsa05310 | Asthma | 0.035541305 | 2 |
| hsa00260 | Glycine, serine and threonine metabolism | 0.048186758 | 2 |

| ID | Description | p.adjust | Count |
|---|---|---|---|
| GO:0022407 | regulation of cell-cell adhesion | 5.95923E-06 | 11 |
| HALLMARK_TNFA_SIGNALING_VIA_NFKB | HALLMARK_TNFA_SIGNALING_VIA_NFKB | 6.48969E-06 | 11 |
| GO:1903037 | regulation of leukocyte cell-cell adhesion | 5.95923E-06 | 10 |
| GO:0007159 | leukocyte cell-cell adhesion | 1.19975E-05 | 10 |
| GO:0002696 | positive regulation of leukocyte activation | 1.96163E-05 | 10 |
| GO:0002443 | leukocyte mediated immunity | 2.0781E-05 | 10 |
| GO:0050867 | positive regulation of cell activation | 2.0781E-05 | 10 |
| GO:0001906 | cell killing | 1.03535E-06 | 9 |
| GO:1903039 | positive regulation of leukocyte cell-cell adhesion | 5.95923E-06 | 9 |
| GO:0022409 | positive regulation of cell-cell adhesion | 1.53074E-05 | 9 |
| GO:0050863 | regulation of T cell activation | 3.07151E-05 | 9 |
| GO:0045785 | positive regulation of cell adhesion | 0.000233203 | 9 |
| GO:0050870 | positive regulation of T cell activation | 2.0781E-05 | 8 |
| GO:0002699 | positive regulation of immune effector process | 2.56855E-05 | 8 |
| GO:0002449 | lymphocyte mediated immunity | 0.000267726 | 8 |
| GO:0002697 | regulation of immune effector process | 0.000318497 | 8 |
| GO:0051251 | positive regulation of lymphocyte activation | 0.000434431 | 8 |
| GO:0001819 | positive regulation of cytokine production | 0.001300668 | 8 |
| GO:0019216 | regulation of lipid metabolic process | 0.001300668 | 7 |
| GO:0032103 | positive regulation of response to external stimulus | 0.004356186 | 7 |
| hsa04145 | Phagosome | 4.57937E-05 | 7 |
| HALLMARK_ALLOGRAFT_REJECTION | HALLMARK_ALLOGRAFT_REJECTION | 0.008376122 | 7 |
| HALLMARK_COMPLEMENT | HALLMARK_COMPLEMENT | 0.008376122 | 7 |
| GO:0046890 | regulation of lipid biosynthetic process | 0.000434431 | 6 |
| GO:0006898 | receptor-mediated endocytosis | 0.00197066 | 6 |
| GO:0002366 | leukocyte activation involved in immune response | 0.003704466 | 6 |
| GO:0002263 | cell activation involved in immune response | 0.003750158 | 6 |
| GO:0031349 | positive regulation of defense response | 0.00401307 | 6 |
| GO:0006959 | humoral immune response | 0.004356186 | 6 |
| GO:0062012 | regulation of small molecule metabolic process | 0.005102952 | 6 |
| GO:0002460 | adaptive immune response based on somatic recombination of im | 0.007452785 | 6 |
| hsa05140 | Leishmaniasis | 2.49128E-05 | 6 |
| HALLMARK_IL2_STAT5_SIGNALING | HALLMARK_IL2_STAT5_SIGNALING | 0.02623853 | 6 |
| GO:0008236 | serine-type peptidase activity | 0.005033471 | 5 |
| GO:0017171 | serine hydrolase activity | 0.005033471 | 5 |
| GO:0004857 | enzyme inhibitor activity | 0.047121206 | 5 |
| GO:0019835 | cytolysis | 2.73766E-06 | 5 |
| GO:0001909 | leukocyte mediated cytotoxicity | 0.001300668 | 5 |
| GO:0002705 | positive regulation of leukocyte mediated immunity | 0.001396061 | 5 |
| GO:0045834 | positive regulation of lipid metabolic process | 0.00197066 | 5 |
| GO:0046631 | alpha-beta T cell activation | 0.003290976 | 5 |
| GO:0002703 | regulation of leukocyte mediated immunity | 0.007452785 | 5 |
| GO:1901214 | regulation of neuron death | 0.018101546 | 5 |
| GO:0070997 | neuron death | 0.024972325 | 5 |
| GO:0006887 | exocytosis | 0.024972325 | 5 |
| GO:0001818 | negative regulation of cytokine production | 0.025361736 | 5 |
| GO:0051346 | negative regulation of hydrolase activity | 0.025755115 | 5 |
| GO:0006631 | fatty acid metabolic process | 0.030901496 | 5 |
| GO:1903706 | regulation of hemopoiesis | 0.032883884 | 5 |
| GO:0050727 | regulation of inflammatory response | 0.033008847 | 5 |
| GO:0030098 | lymphocyte differentiation | 0.034521217 | 5 |
| GO:1903131 | mononuclear cell differentiation | 0.049477236 | 5 |
| hsa05332 | Graft-versus-host disease | 2.49128E-05 | 5 |
| hsa04210 | Apoptosis | 0.002743433 | 5 |
| hsa05152 | Tuberculosis | 0.005512087 | 5 |
| HALLMARK_COAGULATION | HALLMARK_COAGULATION | 0.02623853 | 5 |
| GO:0004252 | serine-type endopeptidase activity | 0.01819729 | 4 |
| GO:0031343 | positive regulation of cell killing | 0.001300668 | 4 |
| GO:0042267 | natural killer cell mediated cytotoxicity | 0.001878875 | 4 |
| GO:0002228 | natural killer cell mediated immunity | 0.00197066 | 4 |
| GO:0019217 | regulation of fatty acid metabolic process | 0.003875298 | 4 |
| GO:0006576 | cellular biogenic amine metabolic process | 0.003922734 | 4 |
| GO:0031341 | regulation of cell killing | 0.004356186 | 4 |
| GO:0006641 | triglyceride metabolic process | 0.004356186 | 4 |
| GO:0044106 | cellular amine metabolic process | 0.004557511 | 4 |
| GO:0048259 | regulation of receptor-mediated endocytosis | 0.005030444 | 4 |
| GO:0035710 | CD4-positive, alpha-beta T cell activation | 0.005248411 | 4 |

| | | | |
|---|---|---|---|
| GO:0046634 | regulation of alpha-beta T cell activation | 0.005248411 | 4 |
| GO:0009308 | amine metabolic process | 0.005400634 | 4 |
| GO:0002708 | positive regulation of lymphocyte mediated immunity | 0.005558235 | 4 |
| GO:0006639 | acylglycerol metabolic process | 0.007465036 | 4 |
| GO:0006638 | neutral lipid metabolic process | 0.007560934 | 4 |
| GO:0010565 | regulation of cellular ketone metabolic process | 0.007739654 | 4 |
| GO:0062013 | positive regulation of small molecule metabolic process | 0.010975517 | 4 |
| GO:0050729 | positive regulation of inflammatory response | 0.011223729 | 4 |
| GO:0002706 | regulation of lymphocyte mediated immunity | 0.017073315 | 4 |
| GO:0045580 | regulation of T cell differentiation | 0.017073315 | 4 |
| GO:0002822 | regulation of adaptive immune response based on somatic recoml | 0.017349606 | 4 |
| GO:0042129 | regulation of T cell proliferation | 0.017349606 | 4 |
| GO:1905952 | regulation of lipid localization | 0.01747261 | 4 |
| GO:0002819 | regulation of adaptive immune response | 0.020554166 | 4 |
| GO:0002285 | lymphocyte activation involved in immune response | 0.022866628 | 4 |
| GO:0030100 | regulation of endocytosis | 0.02325929 | 4 |
| GO:0045619 | regulation of lymphocyte differentiation | 0.023457739 | 4 |
| GO:0042098 | T cell proliferation | 0.024069982 | 4 |
| GO:0016064 | immunoglobulin mediated immune response | 0.024972325 | 4 |
| GO:0019724 | B cell mediated immunity | 0.024972325 | 4 |
| GO:0042180 | cellular ketone metabolic process | 0.024972325 | 4 |
| GO:0002274 | myeloid leukocyte activation | 0.029978051 | 4 |
| GO:0050670 | regulation of lymphocyte proliferation | 0.030489288 | 4 |
| GO:0032944 | regulation of mononuclear cell proliferation | 0.031416759 | 4 |
| GO:1901617 | organic hydroxy compound biosynthetic process | 0.031686684 | 4 |
| GO:0070663 | regulation of leukocyte proliferation | 0.039841943 | 4 |
| GO:0030217 | T cell differentiation | 0.049477236 | 4 |
| hsa05330 | Allograft rejection | 0.000333795 | 4 |
| hsa04940 | Type I diabetes mellitus | 0.000440655 | 4 |
| hsa04612 | Antigen processing and presentation | 0.003320038 | 4 |
| hsa04658 | Th1 and Th2 cell differentiation | 0.005486174 | 4 |
| hsa05150 | Staphylococcus aureus infection | 0.005512087 | 4 |
| hsa04640 | Hematopoietic cell lineage | 0.005512087 | 4 |
| hsa04659 | Th17 cell differentiation | 0.006727796 | 4 |
| hsa05145 | Toxoplasmosis | 0.006886082 | 4 |
| hsa05322 | Systemic lupus erythematosus | 0.012259027 | 4 |
| hsa05164 | Influenza A | 0.024419487 | 4 |
| hsa05202 | Transcriptional misregulation in cancer | 0.030739224 | 4 |
| GO:0055102 | lipase inhibitor activity | 0.001504498 | 3 |
| GO:0023023 | MHC protein complex binding | 0.005033471 | 3 |
| GO:0042053 | regulation of dopamine metabolic process | 0.001170152 | 3 |
| GO:0042069 | regulation of catecholamine metabolic process | 0.001170152 | 3 |
| GO:0033238 | regulation of cellular amine metabolic process | 0.003748374 | 3 |
| GO:0043372 | positive regulation of CD4-positive, alpha-beta T cell differentiati | 0.003852051 | 3 |
| GO:0042401 | cellular biogenic amine biosynthetic process | 0.004356186 | 3 |
| GO:0097242 | amyloid-beta clearance | 0.004356186 | 3 |
| GO:0009309 | amine biosynthetic process | 0.004356186 | 3 |
| GO:0045923 | positive regulation of fatty acid metabolic process | 0.004356186 | 3 |
| GO:0042417 | dopamine metabolic process | 0.004557511 | 3 |
| GO:0090207 | regulation of triglyceride metabolic process | 0.005030444 | 3 |
| GO:2000516 | positive regulation of CD4-positive, alpha-beta T cell activation | 0.005267088 | 3 |
| GO:0031640 | killing of cells of another organism | 0.00542043 | 3 |
| GO:0042304 | regulation of fatty acid biosynthetic process | 0.006707316 | 3 |
| GO:0046638 | positive regulation of alpha-beta T cell differentiation | 0.007452785 | 3 |
| GO:0006584 | catecholamine metabolic process | 0.007687932 | 3 |
| GO:0009712 | catechol-containing compound metabolic process | 0.007687932 | 3 |
| GO:0043370 | regulation of CD4-positive, alpha-beta T cell differentiation | 0.007870781 | 3 |
| GO:0001912 | positive regulation of leukocyte mediated cytotoxicity | 0.008921191 | 3 |
| GO:1905953 | negative regulation of lipid localization | 0.011223729 | 3 |
| GO:0033344 | cholesterol efflux | 0.013791672 | 3 |
| GO:0046635 | positive regulation of alpha-beta T cell activation | 0.014583235 | 3 |
| GO:0046637 | regulation of alpha-beta T cell differentiation | 0.014583235 | 3 |
| GO:0002381 | immunoglobulin production involved in immunoglobulin-mediated | 0.014803538 | 3 |
| GO:0043299 | leukocyte degranulation | 0.017073315 | 3 |
| GO:2000514 | regulation of CD4-positive, alpha-beta T cell activation | 0.017073315 | 3 |
| GO:0032371 | regulation of sterol transport | 0.017349606 | 3 |
| GO:0032374 | regulation of cholesterol transport | 0.017349606 | 3 |
| GO:0001910 | regulation of leukocyte mediated cytotoxicity | 0.019811435 | 3 |

| | | | |
|---|---|---|---|
| GO:0043367 | CD4-positive, alpha-beta T cell differentiation | 0.019931595 | 3 |
| GO:0046889 | positive regulation of lipid biosynthetic process | 0.019931595 | 3 |
| GO:0060191 | regulation of lipase activity | 0.020585176 | 3 |
| GO:1901216 | positive regulation of neuron death | 0.020585176 | 3 |
| GO:0002275 | myeloid cell activation involved in immune response | 0.02325929 | 3 |
| GO:1905477 | positive regulation of protein localization to membrane | 0.024972325 | 3 |
| GO:0042102 | positive regulation of T cell proliferation | 0.027156205 | 3 |
| GO:0002444 | myeloid leukocyte mediated immunity | 0.027672626 | 3 |
| GO:0045833 | negative regulation of lipid metabolic process | 0.028373032 | 3 |
| GO:0018958 | phenol-containing compound metabolic process | 0.02889617 | 3 |
| GO:0002526 | acute inflammatory response | 0.030489288 | 3 |
| GO:0045582 | positive regulation of T cell differentiation | 0.031448355 | 3 |
| GO:0046632 | alpha-beta T cell differentiation | 0.031448355 | 3 |
| GO:0071346 | cellular response to interferon-gamma | 0.031772868 | 3 |
| GO:0030301 | cholesterol transport | 0.032883884 | 3 |
| GO:0051101 | regulation of DNA binding | 0.032883884 | 3 |
| GO:0032612 | interleukin-1 production | 0.038079147 | 3 |
| GO:0032652 | regulation of interleukin-1 production | 0.038079147 | 3 |
| GO:0045621 | positive regulation of lymphocyte differentiation | 0.038079147 | 3 |
| GO:0015918 | sterol transport | 0.041289379 | 3 |
| GO:0034341 | response to interferon-gamma | 0.044346211 | 3 |
| GO:0051384 | response to glucocorticoid | 0.045592439 | 3 |
| GO:0050671 | positive regulation of lymphocyte proliferation | 0.047580448 | 3 |
| GO:0032946 | positive regulation of mononuclear cell proliferation | 0.049477236 | 3 |
| hsa05320 | Autoimmune thyroid disease | 0.008618497 | 3 |
| hsa05134 | Legionellosis | 0.009446446 | 3 |
| hsa05321 | Inflammatory bowel disease | 0.012838681 | 3 |
| hsa04610 | Complement and coagulation cascades | 0.02552442 | 3 |
| hsa05323 | Rheumatoid arthritis | 0.029643741 | 3 |
| hsa04657 | IL-17 signaling pathway | 0.029643741 | 3 |
| hsa05142 | Chagas disease | 0.031718391 | 3 |
| hsa04064 | NF-kappa B signaling pathway | 0.031718391 | 3 |
| hsa04620 | Toll-like receptor signaling pathway | 0.031718391 | 3 |
| hsa04660 | T cell receptor signaling pathway | 0.046279566 | 3 |
| GO:0032395 | MHC class II receptor activity | 0.008254159 | 2 |
| GO:0004859 | phospholipase inhibitor activity | 0.010169684 | 2 |
| GO:0035259 | nuclear glucocorticoid receptor binding | 0.010169684 | 2 |
| GO:0023026 | MHC class II protein complex binding | 0.034793555 | 2 |
| GO:0002423 | natural killer cell mediated immune response to tumor cell | 0.006956521 | 2 |
| GO:0098883 | synapse pruning | 0.007687932 | 2 |
| GO:0042416 | dopamine biosynthetic process | 0.008549784 | 2 |
| GO:0033700 | phospholipid efflux | 0.009804446 | 2 |
| GO:0002399 | MHC class II protein complex assembly | 0.013791672 | 2 |
| GO:0002503 | peptide antigen assembly with MHC class II protein complex | 0.013791672 | 2 |
| GO:0061469 | regulation of type B pancreatic cell proliferation | 0.013791672 | 2 |
| GO:0060192 | negative regulation of lipase activity | 0.014803538 | 2 |
| GO:0010985 | negative regulation of lipoprotein particle clearance | 0.017349606 | 2 |
| GO:0002396 | MHC protein complex assembly | 0.01747261 | 2 |
| GO:0002501 | peptide antigen assembly with MHC protein complex | 0.01747261 | 2 |
| GO:0002888 | positive regulation of myeloid leukocyte mediated immunity | 0.01747261 | 2 |
| GO:0051004 | regulation of lipoprotein lipase activity | 0.01747261 | 2 |
| GO:0009713 | catechol-containing compound biosynthetic process | 0.019931595 | 2 |
| GO:0042423 | catecholamine biosynthetic process | 0.019931595 | 2 |
| GO:0010866 | regulation of triglyceride biosynthetic process | 0.020585176 | 2 |
| GO:0045723 | positive regulation of fatty acid biosynthetic process | 0.020585176 | 2 |
| GO:0150146 | cell junction disassembly | 0.020585176 | 2 |
| GO:0002418 | immune response to tumor cell | 0.024972325 | 2 |
| GO:0044342 | type B pancreatic cell proliferation | 0.024972325 | 2 |
| GO:0045954 | positive regulation of natural killer cell mediated cytotoxicity | 0.024972325 | 2 |
| GO:0070269 | pyroptosis | 0.024972325 | 2 |
| GO:0019886 | antigen processing and presentation of exogenous peptide antige | 0.028373032 | 2 |
| GO:0032372 | negative regulation of sterol transport | 0.028373032 | 2 |
| GO:0032375 | negative regulation of cholesterol transport | 0.028373032 | 2 |
| GO:0034368 | protein-lipid complex remodeling | 0.028373032 | 2 |
| GO:0034369 | plasma lipoprotein particle remodeling | 0.028373032 | 2 |
| GO:0045940 | positive regulation of steroid metabolic process | 0.028373032 | 2 |
| GO:0002717 | positive regulation of natural killer cell mediated immunity | 0.03070184 | 2 |
| GO:0034367 | protein-containing complex remodeling | 0.03070184 | 2 |

| | | | |
|---|---|---|---|
| GO:0002495 | antigen processing and presentation of peptide antigen via MHC | 0.032527054 | 2 |
| GO:0010984 | regulation of lipoprotein particle clearance | 0.032527054 | 2 |
| GO:0032743 | positive regulation of interleukin-2 production | 0.032527054 | 2 |
| GO:0002504 | antigen processing and presentation of peptide or polysaccharide | 0.034780986 | 2 |
| GO:0048261 | negative regulation of receptor-mediated endocytosis | 0.034780986 | 2 |
| GO:1903725 | regulation of phospholipid metabolic process | 0.036477878 | 2 |
| GO:0002478 | antigen processing and presentation of exogenous peptide antige | 0.040959044 | 2 |
| GO:0032965 | regulation of collagen biosynthetic process | 0.040959044 | 2 |
| GO:0019432 | triglyceride biosynthetic process | 0.042216096 | 2 |
| GO:0042119 | neutrophil activation | 0.042216096 | 2 |
| GO:0002714 | positive regulation of B cell mediated immunity | 0.04373164 | 2 |
| GO:0002891 | positive regulation of immunoglobulin mediated immune respons | 0.04373164 | 2 |
| GO:0002347 | response to tumor cell | 0.045255401 | 2 |
| GO:0010828 | positive regulation of glucose transmembrane transport | 0.04678688 | 2 |
| GO:0010712 | regulation of collagen metabolic process | 0.047580448 | 2 |
| GO:0031295 | T cell costimulation | 0.047580448 | 2 |
| GO:0042551 | neuron maturation | 0.047580448 | 2 |
| GO:0071827 | plasma lipoprotein particle organization | 0.047580448 | 2 |
| GO:0001774 | microglial cell activation | 0.049477236 | 2 |
| GO:0031294 | lymphocyte costimulation | 0.049477236 | 2 |
| GO:0032369 | negative regulation of lipid transport | 0.049477236 | 2 |
| GO:0036230 | granulocyte activation | 0.049477236 | 2 |
| GO:0042269 | regulation of natural killer cell mediated cytotoxicity | 0.049477236 | 2 |
| hsa05310 | Asthma | 0.031587744 | 2 |

| ID | Description | p.adjust | Count |
|---|---|---|---|
| GO:0019221 | cytokine-mediated signaling pathway | 0.000121182 | 10 |
| hsa04060 | Cytokine-cytokine receptor interaction | 3.46552E-07 | 10 |
| GO:0140375 | immune receptor activity | 3.23836E-09 | 9 |
| GO:0004896 | cytokine receptor activity | 3.23836E-09 | 8 |
| GO:0019955 | cytokine binding | 5.20138E-08 | 8 |
| GO:0002449 | lymphocyte mediated immunity | 0.000728969 | 8 |
| GO:0002443 | leukocyte mediated immunity | 0.002700678 | 8 |
| GO:0002460 | adaptive immune response based on somatic recombination of im | 0.00394177 | 7 |
| GO:0050863 | regulation of T cell activation | 0.00394177 | 7 |
| hsa05166 | Human T-cell leukemia virus 1 infection | 0.000866652 | 6 |
| HALLMARK_IL6_JAK_STAT3_SIGNALING | HALLMARK_IL6_JAK_STAT3_SIGNALING | 9.89889E-05 | 6 |
| GO:0070663 | regulation of leukocyte proliferation | 0.026284654 | 5 |
| GO:0002440 | production of molecular mediator of immune response | 0.044517837 | 5 |
| GO:0070661 | leukocyte proliferation | 0.045497846 | 5 |
| hsa04061 | Viral protein interaction with cytokine and cytokine receptor | 0.000339408 | 5 |
| hsa04144 | Endocytosis | 0.005253248 | 5 |
| HALLMARK_E2F_TARGETS | HALLMARK_E2F_TARGETS | 0.020185171 | 5 |
| HALLMARK_G2M_CHECKPOINT | HALLMARK_G2M_CHECKPOINT | 0.020185171 | 5 |
| HALLMARK_INFLAMMATORY_RESPONSE | HALLMARK_INFLAMMATORY_RESPONSE | 0.020185171 | 5 |
| GO:0019838 | growth factor binding | 0.00330302 | 4 |
| GO:0002456 | T cell mediated immunity | 0.015180866 | 4 |
| GO:0071887 | leukocyte apoptotic process | 0.015435358 | 4 |
| GO:0002687 | positive regulation of leukocyte migration | 0.026284654 | 4 |
| GO:0002706 | regulation of lymphocyte mediated immunity | 0.040832587 | 4 |
| GO:0042129 | regulation of T cell proliferation | 0.040832587 | 4 |
| GO:0002695 | negative regulation of leukocyte activation | 0.045497846 | 4 |
| hsa04640 | Hematopoietic cell lineage | 0.002467046 | 4 |
| hsa04659 | Th17 cell differentiation | 0.002467046 | 4 |
| hsa04630 | JAK-STAT signaling pathway | 0.008941169 | 4 |
| GO:0016493 | C-C chemokine receptor activity | 0.000571261 | 3 |
| GO:0019957 | C-C chemokine binding | 0.000571261 | 3 |
| GO:0001637 | G protein-coupled chemoattractant receptor activity | 0.000571261 | 3 |
| GO:0004950 | chemokine receptor activity | 0.000571261 | 3 |
| GO:0019956 | chemokine binding | 0.001037343 | 3 |
| GO:0002369 | T cell cytokine production | 0.012162125 | 3 |
| GO:0002724 | regulation of T cell cytokine production | 0.012162125 | 3 |
| GO:0070231 | T cell apoptotic process | 0.026048449 | 3 |
| GO:0071677 | positive regulation of mononuclear cell migration | 0.040832587 | 3 |
| GO:0070227 | lymphocyte apoptotic process | 0.040832587 | 3 |
| GO:0002709 | regulation of T cell mediated immunity | 0.045497846 | 3 |
| GO:0070098 | chemokine-mediated signaling pathway | 0.045497846 | 3 |
| hsa05330 | Allograft rejection | 0.002467046 | 3 |
| hsa05332 | Graft-versus-host disease | 0.002467046 | 3 |
| hsa04940 | Type I diabetes mellitus | 0.002467046 | 3 |
| hsa05320 | Autoimmune thyroid disease | 0.004083407 | 3 |
| hsa04658 | Th1 and Th2 cell differentiation | 0.015232627 | 3 |
| hsa05162 | Measles | 0.043932067 | 3 |
| GO:0005031 | tumor necrosis factor receptor activity | 0.00330302 | 2 |
| GO:0005035 | death receptor activity | 0.005129432 | 2 |
| GO:0042605 | peptide antigen binding | 0.038780751 | 2 |
| GO:2000659 | regulation of interleukin-1-mediated signaling pathway | 0.021876563 | 2 |
| GO:0002693 | positive regulation of cellular extravasation | 0.040832587 | 2 |
| GO:0002827 | positive regulation of T-helper 1 type immune response | 0.040832587 | 2 |

| ID | Description | p.adjust | Count |
|---|---|---|---|
| GO:0001819 | positive regulation of cytokine production | 7.14335E-05 | 11 |
| GO:0002696 | positive regulation of leukocyte activation | 0.000164985 | 10 |
| GO:0050867 | positive regulation of cell activation | 0.000164985 | 10 |
| GO:0006417 | regulation of translation | 0.000164985 | 10 |
| HALLMARK_TNFA_SIGNALING_VIA_NFKB | HALLMARK_TNFA_SIGNALING_VIA_NFKB | 8.71157E-05 | 10 |
| hsa05132 | Salmonella infection | 0.000112422 | 9 |
| GO:1903706 | regulation of hemopoiesis | 0.001038744 | 8 |
| GO:0007159 | leukocyte cell-cell adhesion | 0.001038744 | 8 |
| GO:0045785 | positive regulation of cell adhesion | 0.001825093 | 8 |
| hsa05130 | Pathogenic Escherichia coli infection | 0.000112422 | 8 |
| hsa05166 | Human T-cell leukemia virus 1 infection | 0.000176707 | 8 |
| GO:0140297 | DNA-binding transcription factor binding | 0.009328639 | 7 |
| GO:0002700 | regulation of production of molecular mediator of immune respon | 0.000164985 | 7 |
| GO:0001906 | cell killing | 0.000164985 | 7 |
| GO:0002699 | positive regulation of immune effector process | 0.000628054 | 7 |
| GO:1903039 | positive regulation of leukocyte cell-cell adhesion | 0.000782822 | 7 |
| GO:1903311 | regulation of mRNA metabolic process | 0.001182391 | 7 |
| GO:0022409 | positive regulation of cell-cell adhesion | 0.001270767 | 7 |
| GO:0002440 | production of molecular mediator of immune response | 0.001483038 | 7 |
| GO:0070997 | neuron death | 0.002120625 | 7 |
| GO:0050863 | regulation of T cell activation | 0.002228803 | 7 |
| GO:1903037 | regulation of leukocyte cell-cell adhesion | 0.002228803 | 7 |
| GO:0002697 | regulation of immune effector process | 0.002228803 | 7 |
| GO:0051251 | positive regulation of lymphocyte activation | 0.002941562 | 7 |
| GO:0002443 | leukocyte mediated immunity | 0.005081782 | 7 |
| GO:1903131 | mononuclear cell differentiation | 0.00546283 | 7 |
| GO:0022407 | regulation of cell-cell adhesion | 0.006185096 | 7 |
| GO:0061629 | RNA polymerase II-specific DNA-binding transcription factor bindir | 0.009328639 | 6 |
| GO:0033218 | amide binding | 0.01334403 | 6 |
| GO:0002718 | regulation of cytokine production involved in immune response | 0.000164985 | 6 |
| GO:0002367 | cytokine production involved in immune response | 0.000164985 | 6 |
| GO:0002702 | positive regulation of production of molecular mediator of immun | 0.000243371 | 6 |
| GO:0002705 | positive regulation of leukocyte mediated immunity | 0.000291162 | 6 |
| GO:0045727 | positive regulation of translation | 0.000291162 | 6 |
| GO:0034250 | positive regulation of cellular amide metabolic process | 0.000628054 | 6 |
| GO:0002274 | myeloid leukocyte activation | 0.001844173 | 6 |
| GO:0002703 | regulation of leukocyte mediated immunity | 0.001895458 | 6 |
| GO:0050870 | positive regulation of T cell activation | 0.002178135 | 6 |
| GO:0030217 | T cell differentiation | 0.00386371 | 6 |
| GO:1902105 | regulation of leukocyte differentiation | 0.004822735 | 6 |
| GO:0006959 | humoral immune response | 0.00496842 | 6 |
| GO:1901214 | regulation of neuron death | 0.00511163 | 6 |
| GO:0002237 | response to molecule of bacterial origin | 0.007449008 | 6 |
| GO:0002449 | lymphocyte mediated immunity | 0.007924538 | 6 |
| GO:0001818 | negative regulation of cytokine production | 0.008274399 | 6 |
| GO:0030098 | lymphocyte differentiation | 0.013780542 | 6 |
| GO:0002683 | negative regulation of immune system process | 0.017720936 | 6 |
| GO:0008380 | RNA splicing | 0.019816003 | 6 |
| hsa05161 | Hepatitis B | 0.001751271 | 6 |
| hsa05167 | Kaposi sarcoma-associated herpesvirus infection | 0.002838746 | 6 |
| hsa05169 | Epstein-Barr virus infection | 0.002838746 | 6 |
| hsa05203 | Viral carcinogenesis | 0.002838746 | 6 |
| hsa05170 | Human immunodeficiency virus 1 infection | 0.002949393 | 6 |
| hsa05163 | Human cytomegalovirus infection | 0.003534094 | 6 |
| hsa05131 | Shigellosis | 0.005125754 | 6 |
| HALLMARK_APOPTOSIS | HALLMARK_APOPTOSIS | 0.027045362 | 6 |
| GO:0031625 | ubiquitin protein ligase binding | 0.018393938 | 5 |
| GO:0044389 | ubiquitin-like protein ligase binding | 0.020932696 | 5 |
| GO:0042277 | peptide binding | 0.022608137 | 5 |
| GO:0045296 | cadherin binding | 0.022608137 | 5 |
| GO:0002720 | positive regulation of cytokine production involved in immune re: | 0.000291162 | 5 |
| GO:0002824 | positive regulation of adaptive immune response based on somat | 0.001047117 | 5 |
| GO:0002708 | positive regulation of lymphocyte mediated immunity | 0.001182391 | 5 |
| GO:0002821 | positive regulation of adaptive immune response | 0.001182391 | 5 |
| GO:0001909 | leukocyte mediated cytotoxicity | 0.001527169 | 5 |
| GO:0002706 | regulation of lymphocyte mediated immunity | 0.00370831 | 5 |
| GO:0045580 | regulation of T cell differentiation | 0.00370831 | 5 |

| | | | |
|---|---|---|---|
| GO:0043484 | regulation of RNA splicing | 0.003737662 | 5 |
| GO:0002822 | regulation of adaptive immune response based on somatic recoml | 0.00386371 | 5 |
| GO:1902107 | positive regulation of leukocyte differentiation | 0.00386371 | 5 |
| GO:1903708 | positive regulation of hemopoiesis | 0.00386371 | 5 |
| GO:0002819 | regulation of adaptive immune response | 0.004796612 | 5 |
| GO:0045619 | regulation of lymphocyte differentiation | 0.00546283 | 5 |
| GO:0009612 | response to mechanical stimulus | 0.005468956 | 5 |
| GO:0097193 | intrinsic apoptotic signaling pathway | 0.017720936 | 5 |
| GO:0051054 | positive regulation of DNA metabolic process | 0.017753026 | 5 |
| GO:2001020 | regulation of response to DNA damage stimulus | 0.019587237 | 5 |
| GO:0000377 | RNA splicing, via transesterification reactions with bulged adenos | 0.022390367 | 5 |
| GO:0000398 | mRNA splicing, via spliceosome | 0.022390367 | 5 |
| GO:0009895 | negative regulation of catabolic process | 0.022981826 | 5 |
| GO:0000375 | RNA splicing, via transesterification reactions | 0.023128703 | 5 |
| GO:0070482 | response to oxygen levels | 0.024044204 | 5 |
| GO:0032496 | response to lipopolysaccharide | 0.024510545 | 5 |
| GO:0010506 | regulation of autophagy | 0.025287104 | 5 |
| GO:0042176 | regulation of protein catabolic process | 0.029705223 | 5 |
| GO:0032535 | regulation of cellular component size | 0.030910556 | 5 |
| GO:2001233 | regulation of apoptotic signaling pathway | 0.032061394 | 5 |
| GO:0002460 | adaptive immune response based on somatic recombination of im | 0.032186619 | 5 |
| GO:0050727 | regulation of inflammatory response | 0.04245228 | 5 |
| GO:0045860 | positive regulation of protein kinase activity | 0.044170857 | 5 |
| GO:0043434 | response to peptide hormone | 0.044403184 | 5 |
| GO:0032102 | negative regulation of response to external stimulus | 0.049064078 | 5 |
| hsa04612 | Antigen processing and presentation | 0.000701152 | 5 |
| hsa04625 | C-type lectin receptor signaling pathway | 0.001877697 | 5 |
| hsa04380 | Osteoclast differentiation | 0.002838746 | 5 |
| hsa04210 | Apoptosis | 0.002949393 | 5 |
| hsa05135 | Yersinia infection | 0.002949393 | 5 |
| hsa04010 | MAPK signaling pathway | 0.044207428 | 5 |
| GO:0005200 | structural constituent of cytoskeleton | 0.009328639 | 4 |
| GO:0090079 | translation regulator activity, nucleic acid binding | 0.009328639 | 4 |
| GO:0001916 | positive regulation of T cell mediated cytotoxicity | 0.000243371 | 4 |
| GO:0001914 | regulation of T cell mediated cytotoxicity | 0.000548938 | 4 |
| GO:0001913 | T cell mediated cytotoxicity | 0.001033704 | 4 |
| GO:0001912 | positive regulation of leukocyte mediated cytotoxicity | 0.001236113 | 4 |
| GO:0002711 | positive regulation of T cell mediated immunity | 0.001270767 | 4 |
| GO:0034113 | heterotypic cell-cell adhesion | 0.001270767 | 4 |
| GO:0031100 | animal organ regeneration | 0.001527169 | 4 |
| GO:0031343 | positive regulation of cell killing | 0.001533888 | 4 |
| GO:0032729 | positive regulation of interferon-gamma production | 0.002228803 | 4 |
| GO:0001910 | regulation of leukocyte mediated cytotoxicity | 0.003034575 | 4 |
| GO:0002709 | regulation of T cell mediated immunity | 0.003252282 | 4 |
| GO:0008630 | intrinsic apoptotic signaling pathway in response to DNA damage | 0.004558159 | 4 |
| GO:0031341 | regulation of cell killing | 0.004558159 | 4 |
| GO:0032677 | regulation of interleukin-8 production | 0.004659368 | 4 |
| GO:0032637 | interleukin-8 production | 0.004689985 | 4 |
| GO:0042116 | macrophage activation | 0.005081782 | 4 |
| GO:0048024 | regulation of mRNA splicing, via spliceosome | 0.005081782 | 4 |
| GO:0002456 | T cell mediated immunity | 0.00546283 | 4 |
| GO:0045582 | positive regulation of T cell differentiation | 0.006086371 | 4 |
| GO:0032609 | interferon-gamma production | 0.006185096 | 4 |
| GO:0032649 | regulation of interferon-gamma production | 0.006185096 | 4 |
| GO:0002698 | negative regulation of immune effector process | 0.006644383 | 4 |
| GO:0019730 | antimicrobial humoral response | 0.007205834 | 4 |
| GO:0045621 | positive regulation of lymphocyte differentiation | 0.007924538 | 4 |
| GO:2001235 | positive regulation of apoptotic signaling pathway | 0.008770989 | 4 |
| GO:0050684 | regulation of mRNA processing | 0.00967058 | 4 |
| GO:1903313 | positive regulation of mRNA metabolic process | 0.009834756 | 4 |
| GO:0051100 | negative regulation of binding | 0.015866008 | 4 |
| GO:0021953 | central nervous system neuron differentiation | 0.017753026 | 4 |
| GO:0043488 | regulation of mRNA stability | 0.017753026 | 4 |
| GO:0051048 | negative regulation of secretion | 0.017753026 | 4 |
| GO:0043487 | regulation of RNA stability | 0.021083049 | 4 |
| GO:0061013 | regulation of mRNA catabolic process | 0.021487819 | 4 |
| GO:0031099 | regeneration | 0.022015238 | 4 |
| GO:0071222 | cellular response to lipopolysaccharide | 0.031187891 | 4 |

| | | | |
|---|---|---|---|
| GO:0071219 | cellular response to molecule of bacterial origin | 0.035202805 | 4 |
| GO:0071356 | cellular response to tumor necrosis factor | 0.035202805 | 4 |
| GO:0031330 | negative regulation of cellular catabolic process | 0.039869533 | 4 |
| GO:0006402 | mRNA catabolic process | 0.040224509 | 4 |
| GO:1903320 | regulation of protein modification by small protein conjugation or | 0.041725548 | 4 |
| GO:0034612 | response to tumor necrosis factor | 0.042277748 | 4 |
| GO:0051402 | neuron apoptotic process | 0.04245228 | 4 |
| GO:0006898 | receptor-mediated endocytosis | 0.043221293 | 4 |
| GO:0050708 | regulation of protein secretion | 0.043221293 | 4 |
| GO:0017148 | negative regulation of translation | 0.043608526 | 4 |
| GO:0071216 | cellular response to biotic stimulus | 0.044569878 | 4 |
| GO:0031348 | negative regulation of defense response | 0.049219411 | 4 |
| hsa05416 | Viral myocarditis | 0.002540654 | 4 |
| hsa04657 | IL-17 signaling pathway | 0.00556538 | 4 |
| hsa04659 | Th17 cell differentiation | 0.008030299 | 4 |
| hsa04660 | T cell receptor signaling pathway | 0.011256237 | 4 |
| hsa04145 | Phagosome | 0.02448557 | 4 |
| hsa04218 | Cellular senescence | 0.024796788 | 4 |
| hsa04514 | Cell adhesion molecules | 0.024985525 | 4 |
| hsa04530 | Tight junction | 0.031209146 | 4 |
| GO:0097718 | disordered domain specific binding | 0.009328639 | 3 |
| GO:0042605 | peptide antigen binding | 0.009328639 | 3 |
| GO:0008135 | translation factor activity, RNA binding | 0.020932696 | 3 |
| GO:0031490 | chromatin DNA binding | 0.037486973 | 3 |
| GO:0051082 | unfolded protein binding | 0.039327772 | 3 |
| GO:0019885 | antigen processing and presentation of endogenous peptide antig | 0.001047117 | 3 |
| GO:0002483 | antigen processing and presentation of endogenous peptide antig | 0.001211031 | 3 |
| GO:0001911 | negative regulation of leukocyte mediated cytotoxicity | 0.001895458 | 3 |
| GO:0046641 | positive regulation of alpha-beta T cell proliferation | 0.001895458 | 3 |
| GO:0019883 | antigen processing and presentation of endogenous antigen | 0.002085317 | 3 |
| GO:0031342 | negative regulation of cell killing | 0.002228803 | 3 |
| GO:0097421 | liver regeneration | 0.002428903 | 3 |
| GO:0002474 | antigen processing and presentation of peptide antigen via MHC | 0.002638888 | 3 |
| GO:0002369 | T cell cytokine production | 0.004340758 | 3 |
| GO:0002724 | regulation of T cell cytokine production | 0.004340758 | 3 |
| GO:0002478 | antigen processing and presentation of exogenous peptide antige | 0.004689985 | 3 |
| GO:0150077 | regulation of neuroinflammatory response | 0.004822735 | 3 |
| GO:0046640 | regulation of alpha-beta T cell proliferation | 0.00546283 | 3 |
| GO:0046633 | alpha-beta T cell proliferation | 0.006181189 | 3 |
| GO:0019884 | antigen processing and presentation of exogenous antigen | 0.006644383 | 3 |
| GO:0009409 | response to cold | 0.007924538 | 3 |
| GO:0002707 | negative regulation of lymphocyte mediated immunity | 0.008770989 | 3 |
| GO:0043331 | response to dsRNA | 0.008770989 | 3 |
| GO:0043030 | regulation of macrophage activation | 0.011250904 | 3 |
| GO:0048002 | antigen processing and presentation of peptide antigen | 0.012109655 | 3 |
| GO:0002704 | negative regulation of leukocyte mediated immunity | 0.013920119 | 3 |
| GO:0014015 | positive regulation of gliogenesis | 0.014391925 | 3 |
| GO:0046635 | positive regulation of alpha-beta T cell activation | 0.016354947 | 3 |
| GO:0150076 | neuroinflammatory response | 0.016354947 | 3 |
| GO:0042267 | natural killer cell mediated cytotoxicity | 0.016864489 | 3 |
| GO:0002228 | natural killer cell mediated immunity | 0.017753026 | 3 |
| GO:0061844 | antimicrobial humoral immune response mediated by antimicrobi | 0.019235002 | 3 |
| GO:0006446 | regulation of translational initiation | 0.019587237 | 3 |
| GO:0021954 | central nervous system neuron development | 0.021083049 | 3 |
| GO:0033077 | T cell differentiation in thymus | 0.021487819 | 3 |
| GO:0010507 | negative regulation of autophagy | 0.021584016 | 3 |
| GO:1903312 | negative regulation of mRNA metabolic process | 0.025166454 | 3 |
| GO:0032088 | negative regulation of NF-kappaB transcription factor activity | 0.025287104 | 3 |
| GO:0030101 | natural killer cell activation | 0.027714015 | 3 |
| GO:1903076 | regulation of protein localization to plasma membrane | 0.029894424 | 3 |
| GO:1901796 | regulation of signal transduction by p53 class mediator | 0.030910556 | 3 |
| GO:0042102 | positive regulation of T cell proliferation | 0.031295892 | 3 |
| GO:0014013 | regulation of gliogenesis | 0.031953524 | 3 |
| GO:0019882 | antigen processing and presentation | 0.032061394 | 3 |
| GO:0033209 | tumor necrosis factor-mediated signaling pathway | 0.032539372 | 3 |
| GO:1901222 | regulation of NIK/NF-kappaB signaling | 0.033875986 | 3 |
| GO:0046634 | regulation of alpha-beta T cell activation | 0.036284419 | 3 |
| GO:0050830 | defense response to Gram-positive bacterium | 0.038333947 | 3 |

| | | | |
|---|---|---|---|
| GO:0071346 | cellular response to interferon-gamma | 0.038333947 | 3 |
| GO:0045446 | endothelial cell differentiation | 0.039869533 | 3 |
| GO:0002224 | toll-like receptor signaling pathway | 0.040906115 | 3 |
| GO:0006413 | translational initiation | 0.040906115 | 3 |
| GO:0022612 | gland morphogenesis | 0.040906115 | 3 |
| GO:1904375 | regulation of protein localization to cell periphery | 0.042160479 | 3 |
| GO:0019079 | viral genome replication | 0.044612597 | 3 |
| hsa05330 | Allograft rejection | 0.005125754 | 3 |
| hsa05332 | Graft-versus-host disease | 0.005882078 | 3 |
| hsa04940 | Type I diabetes mellitus | 0.005989287 | 3 |
| hsa05134 | Legionellosis | 0.011256237 | 3 |
| hsa05140 | Leishmaniasis | 0.024796788 | 3 |
| hsa04933 | AGE-RAGE signaling pathway in diabetic complications | 0.044207428 | 3 |
| hsa05142 | Chagas disease | 0.045160403 | 3 |
| GO:0030957 | Tat protein binding | 0.009328639 | 2 |
| GO:0031386 | protein tag | 0.01277816 | 2 |
| GO:0043023 | ribosomal large subunit binding | 0.01277816 | 2 |
| GO:0044548 | S100 protein binding | 0.01277816 | 2 |
| GO:0003746 | translation elongation factor activity | 0.018393938 | 2 |
| GO:0023026 | MHC class II protein complex binding | 0.026784572 | 2 |
| GO:0051019 | mitogen-activated protein kinase binding | 0.027280513 | 2 |
| GO:0051059 | NF-kappaB binding | 0.031719822 | 2 |
| GO:0023023 | MHC protein complex binding | 0.038739201 | 2 |
| GO:0031492 | nucleosomal DNA binding | 0.043669529 | 2 |
| GO:0048156 | tau protein binding | 0.0482882 | 2 |
| GO:0046643 | regulation of gamma-delta T cell activation | 0.007924538 | 2 |
| GO:0042492 | gamma-delta T cell differentiation | 0.010305164 | 2 |
| GO:0060213 | positive regulation of nuclear-transcribed mRNA poly(A) tail short | 0.010305164 | 2 |
| GO:0002475 | antigen processing and presentation via MHC class Ib | 0.011767448 | 2 |
| GO:0042994 | cytoplasmic sequestering of transcription factor | 0.013168066 | 2 |
| GO:0060211 | regulation of nuclear-transcribed mRNA poly(A) tail shortening | 0.013168066 | 2 |
| GO:0061684 | chaperone-mediated autophagy | 0.014472973 | 2 |
| GO:0010958 | regulation of amino acid import across plasma membrane | 0.017282194 | 2 |
| GO:0150079 | negative regulation of neuroinflammatory response | 0.017282194 | 2 |
| GO:1903789 | regulation of amino acid transmembrane transport | 0.017282194 | 2 |
| GO:0002923 | regulation of humoral immune response mediated by circulating i | 0.017753026 | 2 |
| GO:0043031 | negative regulation of macrophage activation | 0.017753026 | 2 |
| GO:0045953 | negative regulation of natural killer cell mediated cytotoxicity | 0.017753026 | 2 |
| GO:0002716 | negative regulation of natural killer cell mediated immunity | 0.019235002 | 2 |
| GO:0032495 | response to muramyl dipeptide | 0.019235002 | 2 |
| GO:0060252 | positive regulation of glial cell proliferation | 0.020586268 | 2 |
| GO:0010888 | negative regulation of lipid storage | 0.021487819 | 2 |
| GO:0051220 | cytoplasmic sequestering of protein | 0.021487819 | 2 |
| GO:1903209 | positive regulation of oxidative stress-induced cell death | 0.021487819 | 2 |
| GO:0048025 | negative regulation of mRNA splicing, via spliceosome | 0.022673895 | 2 |
| GO:0002922 | positive regulation of humoral immune response | 0.024044204 | 2 |
| GO:0031579 | membrane raft organization | 0.025287104 | 2 |
| GO:0050686 | negative regulation of mRNA processing | 0.025287104 | 2 |
| GO:0002726 | positive regulation of T cell cytokine production | 0.026597834 | 2 |
| GO:0046629 | gamma-delta T cell activation | 0.026597834 | 2 |
| GO:0048026 | positive regulation of mRNA splicing, via spliceosome | 0.026597834 | 2 |
| GO:0002710 | negative regulation of T cell mediated immunity | 0.028143857 | 2 |
| GO:0042026 | protein refolding | 0.028143857 | 2 |
| GO:0032703 | negative regulation of interleukin-2 production | 0.029705223 | 2 |
| GO:0033119 | negative regulation of RNA splicing | 0.029705223 | 2 |
| GO:0001773 | myeloid dendritic cell activation | 0.030910556 | 2 |
| GO:0010592 | positive regulation of lamellipodium assembly | 0.030910556 | 2 |
| GO:0009651 | response to salt stress | 0.032061394 | 2 |
| GO:1903203 | regulation of oxidative stress-induced neuron death | 0.032061394 | 2 |
| GO:0035767 | endothelial cell chemotaxis | 0.033314479 | 2 |
| GO:0036475 | neuron death in response to oxidative stress | 0.036284419 | 2 |
| GO:0045943 | positive regulation of transcription by RNA polymerase I | 0.036284419 | 2 |
| GO:1902253 | regulation of intrinsic apoptotic signaling pathway by p53 class m | 0.036284419 | 2 |
| GO:0000289 | nuclear-transcribed mRNA poly(A) tail shortening | 0.038282799 | 2 |
| GO:0002719 | negative regulation of cytokine production involved in immune re | 0.039275837 | 2 |
| GO:0050685 | positive regulation of mRNA processing | 0.039275837 | 2 |
| GO:1901099 | negative regulation of signal transduction in absence of ligand | 0.039275837 | 2 |
| GO:2001240 | negative regulation of extrinsic apoptotic signaling pathway in ab | 0.039275837 | 2 |

| | | | |
|---|---|---|---|
| GO:1902745 | positive regulation of lamellipodium organization | 0.042073734 | 2 |
| GO:0060251 | regulation of glial cell proliferation | 0.044446362 | 2 |
| GO:0030866 | cortical actin cytoskeleton organization | 0.045839575 | 2 |
| GO:0033120 | positive regulation of RNA splicing | 0.045839575 | 2 |
| GO:0002714 | positive regulation of B cell mediated immunity | 0.049064078 | 2 |
| GO:0002891 | positive regulation of immunoglobulin mediated immune respons | 0.049064078 | 2 |
| GO:0010591 | regulation of lamellipodium assembly | 0.049064078 | 2 |
| GO:0014002 | astrocyte development | 0.049064078 | 2 |
| GO:0045429 | positive regulation of nitric oxide biosynthetic process | 0.049064078 | 2 |

**Table S1. [Enriched pathways for each lymphoid-derived irMP], Related to Figure 1.**
“GeneRatio” is the fraction of irMP genes that are found in the corresponding enriched pathway. “BgRatio” is the fraction of genes annotated to the enriched pathway versus the number of genes annotated to that specific pathway database within the background. “geneID” contains the irMP genes that are in the enriched pathway.

| V1 | V2 | V3 | V4 | V5 | V6 | V7 |
|---|---|---|---|---|---|---|
| MYBL1 | TRAC | EIF1AY | DUSP4 | CD1B | RRM2 | RGS1 |
| AICDA | SATB1 | TXLNGY | CSF2RB | CD1E | STMN1 | CCR2 |
| MME | CD28 | KDM5D | IL2RA | DNTT | GZMA | PTGER2 |
| PRKAR2B | ABLIM1 | CD1A | METTL7A | TXLNGY | DLGAP5 | SGK1 |
| SPRED2 | ICOS | DNTT | CCR6 | TRDV3 | TUBA1B | P2RY14 |
| ASPM | CD69 | CD1E | PMCH | CD8A | TUBB | TIGIT |
| KANK1 | CD3D | TRDV3 | LOC101928152 | CD1A | CCR1 | IL17RB |
| KANK2 | TRBC1 | CD1B | FAM129A | EIF1AY | HJURP | TMEM200A |
| DLGAP5 | LTB | HIST1H3I | TIGIT | CD1C | CDK1 | CSF2RB |
| UCHL1 | CD52 | HIST1H2BM | TRIB1 | KDM5D | BUB1B | PDE4B |
| E2F8 | LBH | HIST1H4E | RP11-28F1.2 | AEBP1 | ENO1 | CXCL8 |
| PBK | SH2D1A | HIST1H2AJ | CTLA4 | SH3TC1 | CKS2 | ADAM19 |
| PDGFD | EVI2A | AKR1C3 | PTGER2 | HMGB2 | BUB1 | TLR1 |
| CCNE2 | LCP2 | HIST1H2BH | LOC101929734 | LOC100507600 | ASPM | RTKN2 |
| CEP55 | CYTIP | ND6 | IL1R1 | PXYLP1 | SNRPA | AGPAT9 |
| CDC20 | CD247 | GJC1 | RAB11FIP1 | MND1 | UQCRFS1 | ITM2B |
| TPX2 | BACH2 | HTATSF1P2 | CXCR6 | RCAN1 | GNLY | MEOX1 |
| TJP2 | ITM2A | HIST1H2BE | RTKN2 | FABP5 | ATP5F1 | FRMD4B |
| SUGCT | ID2 | MALAT1 | AC017002.2 | MALAT1 | FTL | MB21D2 |
| PRC1 | CLEC2D | SNORD8 | LOC101928173 | RAG2 | TUBA1C | CTSH |
| KIF2C | GPR18 | H2BFS | FANK1 | ND6 | CCNB2 | HLA-DPA1 |
| HJURP | LAT | CD1C | CISH | HMGN2 | CEP55 | MYBL1 |
| BUB1B | LRRN3 | HIST1H4J | ZNF662 | RAG1 | ZWINT | TRADD |
| KIF4A | ITPKB | RRM2 | PDE4A | PTMA | CD38 | ITGAM |
| KIF20A | DENND2D | LOC100996740 | PLS3 | HNRNPA2B1 | KIR2DL4 | GZMB |
| HMMR | LCK | HIST1H2AE | S100A4 | GXYLT2 | PBK | GZMA |
| SPRR3 | PDE4D | HIST1H4L | GJB6 | TFDP2 | NUSAP1 | KAT2B |
| GINS2 | PLXDC1 | HMGB2 | LAIR2 | GLUL | H2AFV | TWIST1 |
| CDKN3 | CD53 | MCM8 | TWIST1 | NUCB2 | CLEC2B | LRRC32 |
| PALLD | PPP1R16B | TCEAL3 | TMEM62 | RRM2 | KIF15 | CXCL9 |
| CCNB2 | CHI3L2 | KCNJ4 | TNFRSF1B | HSP90AA1 | ARPC1B | TNFRSF1B |
| OIP5 | ZFP36L1 | HNRNPA2B1 | TOX2 | CPVL | ADSL | KLF2 |
| ATAD2 | CD2 | BCORP1 | MYB | MYB | KPNA2 | DDX60 |
| ECT2 | LPXN | TYMS | ATP1B1 | SMIM24 | PKM | KLF6 |
| CCNB1 | GPR171 | HMGB3 | PKD2 | BUB1B | MANF | FKBP11 |
| CTNNAL1 | HLA-E | HIST1H1C | SOCS2 | TUBB | PRF1 | TUBB2A |
| ZWINT | SLAMF1 | PTMA | RGS2 | UHRF1 | GINS2 | RAB11FIP1 |
| CCNA2 | TRAT1 | ND2 | ADAM19 | MAP1A | NDUFA13 | CECR1 |
| DMD | CRLF3 | HIST1H4H | RAB37 | TUBA1C | LGALS1 | TGFBR3 |
| SPC25 | FRY | HIST1H3G | CLDND1 | PTTG1 | CENPU | TRIM22 |
| MET | BTG1 | LRRN1 | TNFRSF9 | SATB1 | TUBB4B | HBEGF |
| TTK | ARHGEF6 | MTPN | NEBL | KCNJ4 | ALDOA | PMCH |
| KIF14 | LAPTM5 | SNHG19 | VMP1 | CCR9 | CDKN2D | OLAH |
| CENPU | ZBED5 | HIST1H2AB | IL1R2 | HMGB1 | BANF1 | BASP1 |
| FGD6 | TRAF3IP3 | RAG1 | ZDHHC23 | LDOC1L | HSP90AB1 | IL2RA |
| RGS13 | ARNTL | ATP6 | CAMK2N1 | HNRNPAB | ATP5H | CISH |
| IGF2BP3 | HERC1 | PET100 | APLP2 | CCNB1 | KIAA0101 | HMGB2 |
| FHOD3 | ST6GAL1 | UTY | DUSP6 | HSPD1 | CCNB1 | MAP3K8 |
| NEIL3 | SKAP1 | GXYLT2 | PFKFB3 | CHRNA3 | KIF11 | FAS |
| ASB13 | IER3 | HMGN2 | NACC2 | DLGAP5 | RNF167 | CCND2 |

| V8 | V9 | V10 | V11 | V12 | V13 | V14 |
|---|---|---|---|---|---|---|
| HUWE1 | ENO1 | GSTT1 | ZBED2 | DNTT | XIST | HLA-DQB1 |
| ACTB | TUBA1C | CD74 | SLC27A2 | RAG1 | TSIX | CMPK2 |
| NPM1 | PGK1 | VMP1 | RRM2 | MZB1 | NELL2 | IFI44L |
| NACA | GAPDH | ADM | TOP2A | CD1B | RGS1 | XAF1 |
| EEF1A1 | PCNA | FAM118A | KIAA0101 | SYK | CTSL | IFITM1 |
| LDHA | ALDOA | MAL | BUB1B | GNA15 | NOG | OAS1 |
| PPIA | CEP55 | CCL22 | DLGAP5 | TRDV3 | RP11-664D1.1 | PRKAR1A |
| TMSB10 | MCM6 | FABP4 | ASPM | MYB | EPHA1 | RAP2C |
| FAU | TUBB | NOTCH2NL | EGR3 | TOP2A | SLC18A2 | RP11-28F1.2 |
| GAPDH | PGAM1 | ABCG1 | KIF15 | TFDP2 | GZMA | MALAT1 |
| TPT1 | FDPS | NFXL1 | CEP55 | ADA | PDE3B | IFI44 |
| GLTSCR2 | STAT1 | ACTB | CRNDE | FABP5 | RAB33A | CHMP5 |
| B2M | TPI1 | ND6 | GZMB | NOTCH1 | GIMAP8 | ELL3 |
| EIF1 | IMPDH2 | Y16709 | EGR2 | CD1A | GPR171 | DTX3L |
| HLA-A | STMN1 | EGFR | SQRDL | GUCY1A3 | CCR7 | IFIT1 |
| ANP32B | LDHA | APOC1 | RAD51AP1 | DEFA6 | IFITM1 | EPSTI1 |
| RP5-882O7.1 | RACGAP1 | STARD7 | CCNB2 | NINL | FBLN7 | ISG15 |
| TMSB4X | KIAA0101 | ABCC3 | NDFIP2 | MAL | GPR155 | RSAD2 |
| ATP6 | HMMR | FABP5 | P2RX5 | HLA-DPA1 | PRRT3 | ND6 |
| MALAT1 | CDC2 | IL1RN | STRIP2 | GZMB | GIMAP4 | TLR7 |
| DUSP1 | MTHFD1 | CD300A | ZC3H12D | MCM2 | TCEA3 | STAT1 |
| KLRB1 | PSMB9 | PLIN2 | APOBEC3B | DTL | FGFBP2 | CSGALNACT2 |
| COX2 | CCT8 | LOC102724870 | LAG3 | IGJ | LRRN3 | DDX60 |
| LYZ | PSMA1 | AGPAT9 | CTLA4 | GAPDH | GIMAP7 | IRF7 |
| COX5A | H2AFZ | ERV3-2 | SMCO4 | TLR7 | TCF3 | CCNG2 |
| HLA-DPA1 | CCT7 | ZFYVE16 | SYNGR3 | MCM7 | PLAC8 | TSPAN13 |
| IGHM | C14ORF1 | LIPA | DTL | RAG2 | SELPLG | FNTA |
| IGLV1-44 | C16ORF75 | PAPD4 | DUSP6 | CD99 | MEOX1 | TMEM123 |
| HLA-DRA | COX1 | FTL | ZNF282 | ENPP2 | TMEM204 | USP18 |
| GZMB | ATP5B | MPRIP | TNFRSF9 | GUCY1B3 | FOS | IGHM |
| CCL5 | CCT4 | RAB11FIP1 | MST4 | SLC1A4 | FARS2 | CD69 |
| FAIM3 | FDFT1 | TMED9 | TRIB1 | KIF18B | DNAJB1 | LGALS3BP |
| RP3-486D24.1 | CLIC1 | LGALS3 | NUF2 | FXYD2 | DNAJC15 | LYRM5 |
| EEF1D | ENO2 | ND4 | ETS1 | CEP70 | SELL | KRR1 |
| HLA-DRB4 | PLP2 | HEXB | DDIT4 | ATIC | IL7R | YY1 |
| LIPA | PDIA6 | SDCBP | SCP2 | BUB1B | HLA-B | COMMD8 |
| ND6 | DARS | RP5-882O7.1 | MMADHC | GUK1 | IL12RB1 | AMFR |
| HNRNPA2B1 | RAD51AP1 | MALAT1 | IFNG | KIAA0922 | OAS1 | UQCR11 |
| NUCB2 | HNRNPA2B1 | B2M | MTHFD2 | MCM6 | RHOU | CETN3 |
| TUBA1B | EXOC3 | HADHB | EPAS1 | CPVL | IFI44L | ZNF800 |
| HSPA6 | HMGB2 | MMP12 | SLC1A4 | CBFA2T3 | FOXO1 | HIST1H2BD |
| ISG15 | CDC20 | HUWE1 | ALDOA | GAS7 | EPSTI1 | NDUFA5 |
| MX1 | ACTR3 | C1ORF122 | TOX2 | NUSAP1 | RSAD2 | MX1 |
| IFITM1 | PARK7 | NGLY1 | CD3D | SAC3D1 | RGS2 | MORC3 |
| OPHN1 | PPIA | CAP1 | PRDX3 | GTF3A | CAPN2 | DCK |
| DDX5 | C12ORF11 | ATP1B3 | RAB13 | CCNB2 | GBP2 | UBE2D1 |
| GNB2L1 | SMC4 | GALNT12 | DENND2D | CD3D | ZBTB24 | TMEM60 |
| ACTG1 | PTTG1 | NISCH | UBB | TYMS | UBA52 | TMA7 |
| PTMA | SQRDL | AKAP17A | UBE2F | PTP4A2 | ESYT1 | ZNF107 |
| CFL1 | SMS | TGFBI | HMGCR | IRF8 | ISG20 | KPNA2 |

| V15 | V16 | V17 | V18 | V19 |
|---|---|---|---|---|
| ZNF548 | CD52 | EIF1AY | LGALS3 | VIM |
| EGFR | HLA-DQB1 | APOC1 | FAM129A | EEF1A1 |
| ODC1 | UBC | HLA-DPA1 | MEOX1 | B2M |
| THBD | ETS1 | GZMH | IL1R1 | TMSB4X |
| ITGB2 | NDUFA1 | SERPINA1 | LOC101928173 | FAU |
| ALDOA | NACA | IFNG | PMCH | GAPDH |
| GAPDH | P2RX5 | HLA-DQA1 | TRIB1 | HLA-A |
| LGALS1 | HLA-DMA | CRTAM | MIAT | JUND |
| TRIB1 | FBL | NR4A2 | FANK1 | EZR |
| ACTB | CYBA | CPVL | CSF2RB | EIF4G2 |
| TUBA1C | RBM3 | TUBA4A | RP11-28F1.2 | ACTB |
| PYCARD | TMSB10 | TXLNGY | TXLNGY | IER3 |
| HLA-DRB4 | HLA-E | ERRFI1 | KDM5D | OTUD4 |
| GSTT1 | COX6A1 | RGCC | KIAA0101 | EGR3 |
| CD300A | FTL | TP53INP2 | RRM2 | RBM3 |
| FCGR2B | ALAS2 | EFNB2 | CXCR6 | EEF2 |
| CTSD | SELL | GZMK | MELK | SLC7A5 |
| SPATA24 | ADD3 | PPP1R2P9 | IL1R2 | CD97 |
| MYL6 | PABPC1 | GZMB | IL2RB | TUBA1A |
| SASH3 | DYNLL1 | ALDH2 | CCR2 | NACA |
| APEX1 | TMBIM6 | CD83 | CLDND1 | PABPC1 |
| PDIA6 | VIM | ZNF331 | DUSP4 | PGAM1 |
| FCER2 | KIF1C | TFRC | NUSAP1 | UBB |
| ADORA3 | NDUFC1 | APOC2 | RP5-882O7.1 | NPM1 |
| COX6B1 | SSR2 | S100A4 | TIGIT | UBC |
| TMSB4X | UQCRB | CCL4 | CCR6 | CRIP1 |
| FTH1 | HBD | TUBA1C | HMMR | ARPC2 |
| MGST2 | HMGB2 | CCRN4L | KLRB1 | DUSP2 |
| FAM98A | HLA-A | KLF6 | FAS | CD200 |
| DAD1 | TCAP | FAM46C | TOX | KDM6B |
| CRELD2 | SNRPG | NR4A3 | RAB11FIP1 | TNF |
| HLA-B | RAC2 | AHNAK | OAZ1 | CD2 |
| ENO1 | SEL1L3 | GNLY | HNRNPH1 | UCP2 |
| HMGN2 | EEF1D | ODC1 | KPNA2 | HLA-F |
| PPIB | PPIB | KLRD1 | TOP2A | HNRNPK |
| OSTC | CXCR4 | MAP3K8 | FRMD4B | HMGN2 |
| CD52 | S100A8 | SIK1 | SMCHD1 | EGR2 |
| BLVRB | HMGN1 | ITGAM | IL2RA | IER2 |
| MAT2A | CORO1A | ARL4A | HLA-C | TGFBR2 |
| TYROBP | SYPL1 | EIF4A3 | VMP1 | DDX21 |
| PIK3C2B | ISG20 | IDI1 | STAM | NCL |
| TMSB10 | EIF3E | GZMA | HLA-DQB1 | NFKBIA |
| TUBB6 | YWHAB | ANXA1 | RGS1 | COX4I1 |
| GAA | EIF1AX | NFKBIA | MALAT1 | AHNAK |
| VAMP8 | COX7C | BCL2A1 | YWHAZ | BTG2 |
| FCGRT | VOPP1 | EIF1 | CDCA7 | PTPRC |
| GPX1 | FAU | LYZ | TNFRSF1B | SLA |
| S100A13 | HNRNPA2B1 | NKG7 | KIF11 | NFATC1 |
| PIK3CD | ACTR2 | C3 | PLEKHB2 | HSP90AA1 |
| HHEX | PGAM1 | FOSB | RTKN2 | MCL1 |

**Table S2. [Lymphoid-derived gene sets], Related to Figure 1**.
The 19 lymphoid-derived irMPs with their constituent genes. Each column is one irMP.

| ID | Description | p.adjust | Count |
|---|---|---|---|
| HALLMARK_INTERFERON_GAMMA_RESPONS | HALLMARK_INTERFERON_GAMMA_RESPONSE | 8.48951E-13 | 17 |
| GO:0009615 | response to virus | 1.33456E-09 | 14 |
| HALLMARK_INTERFERON_ALPHA_RESPONSE | HALLMARK_INTERFERON_ALPHA_RESPONSE | 3.02366E-13 | 14 |
| GO:0051607 | defense response to virus | 4.74183E-09 | 12 |
| GO:0140546 | defense response to symbiont | 4.74183E-09 | 12 |
| GO:0016032 | viral process | 6.92013E-05 | 9 |
| hsa05164 | Influenza A | 1.08097E-06 | 9 |
| hsa05169 | Epstein-Barr virus infection | 2.31969E-06 | 9 |
| hsa05166 | Human T-cell leukemia virus 1 infection | 3.87345E-05 | 8 |
| GO:0045069 | regulation of viral genome replication | 5.91071E-07 | 7 |
| GO:0048525 | negative regulation of viral process | 6.77101E-07 | 7 |
| GO:0019882 | antigen processing and presentation | 1.50554E-06 | 7 |
| GO:0019079 | viral genome replication | 4.47535E-06 | 7 |
| GO:1903900 | regulation of viral life cycle | 6.96453E-06 | 7 |
| GO:0050792 | regulation of viral process | 1.39612E-05 | 7 |
| GO:0019058 | viral life cycle | 0.000882045 | 7 |
| GO:0001819 | positive regulation of cytokine production | 0.007983302 | 7 |
| GO:0045071 | negative regulation of viral genome replication | 1.09042E-06 | 6 |
| GO:0048002 | antigen processing and presentation of peptide ant | 1.77841E-06 | 6 |
| GO:0051251 | positive regulation of lymphocyte activation | 0.013966739 | 6 |
| GO:0002696 | positive regulation of leukocyte activation | 0.020753884 | 6 |
| GO:0050867 | positive regulation of cell activation | 0.022753521 | 6 |
| GO:0019221 | cytokine-mediated signaling pathway | 0.027770964 | 6 |
| hsa04612 | Antigen processing and presentation | 1.65202E-05 | 6 |
| hsa04145 | Phagosome | 0.000315388 | 6 |
| hsa05160 | Hepatitis C | 0.000341357 | 6 |
| hsa05163 | Human cytomegalovirus infection | 0.001843272 | 6 |
| hsa05165 | Human papillomavirus infection | 0.010832269 | 6 |
| GO:0002483 | antigen processing and presentation of endogenou | 3.29619E-07 | 5 |
| GO:0019883 | antigen processing and presentation of endogenou | 1.05146E-06 | 5 |
| GO:0002478 | antigen processing and presentation of exogenous | 5.97464E-06 | 5 |
| GO:0019884 | antigen processing and presentation of exogenous | 1.36169E-05 | 5 |
| GO:0034340 | response to type I interferon | 0.000145788 | 5 |
| GO:0050870 | positive regulation of T cell activation | 0.013230186 | 5 |
| GO:1903039 | positive regulation of leukocyte cell-cell adhesion | 0.016133813 | 5 |
| GO:0022409 | positive regulation of cell-cell adhesion | 0.024804585 | 5 |
| GO:0002440 | production of molecular mediator of immune respo | 0.027770964 | 5 |
| GO:0002237 | response to molecule of bacterial origin | 0.034284658 | 5 |
| GO:0002831 | regulation of response to biotic stimulus | 0.034284658 | 5 |
| GO:0002449 | lymphocyte mediated immunity | 0.034804049 | 5 |
| GO:0002460 | adaptive immune response based on somatic recon | 0.037059077 | 5 |
| GO:0050863 | regulation of T cell activation | 0.037059077 | 5 |
| GO:1903037 | regulation of leukocyte cell-cell adhesion | 0.037059077 | 5 |
| hsa04218 | Cellular senescence | 0.002560967 | 5 |
| hsa05170 | Human immunodeficiency virus 1 infection | 0.009724147 | 5 |
| hsa05171 | Coronavirus disease - COVID-19 | 0.012687806 | 5 |
| GO:0042605 | peptide antigen binding | 0.000570325 | 4 |
| GO:0003823 | antigen binding | 0.045749614 | 4 |
| GO:0019885 | antigen processing and presentation of endogenou | 8.29886E-06 | 4 |
| GO:0140374 | antiviral innate immune response | 2.08108E-05 | 4 |
| GO:0002474 | antigen processing and presentation of peptide ant | 6.92013E-05 | 4 |
| GO:0002711 | positive regulation of T cell mediated immunity | 0.000893483 | 4 |
| GO:0071357 | cellular response to type I interferon | 0.002188191 | 4 |
| GO:0002709 | regulation of T cell mediated immunity | 0.00347482 | 4 |
| GO:0002456 | T cell mediated immunity | 0.007732719 | 4 |
| GO:0002824 | positive regulation of adaptive immune response b | 0.007770497 | 4 |
| GO:0002708 | positive regulation of lymphocyte mediated immur | 0.008666263 | 4 |
| GO:0002821 | positive regulation of adaptive immune response | 0.008712986 | 4 |
| GO:0002705 | positive regulation of leukocyte mediated immunit | 0.013230186 | 4 |
| GO:0010821 | regulation of mitochondrion organization | 0.015144954 | 4 |
| GO:0002706 | regulation of lymphocyte mediated immunity | 0.022639007 | 4 |
| GO:0002822 | regulation of adaptive immune response based on | 0.022753521 | 4 |
| GO:0002819 | regulation of adaptive immune response | 0.028138302 | 4 |
| GO:0016064 | immunoglobulin mediated immune response | 0.034284658 | 4 |
| GO:0019724 | B cell mediated immunity | 0.034759126 | 4 |
| GO:0002377 | immunoglobulin production | 0.034759126 | 4 |
| GO:0002703 | regulation of leukocyte mediated immunity | 0.044161584 | 4 |

| | | | |
|---|---|---|---|
| hsa05330 | Allograft rejection | 0.000305799 | 4 |
| hsa05332 | Graft-versus-host disease | 0.000315388 | 4 |
| hsa04940 | Type I diabetes mellitus | 0.000315388 | 4 |
| hsa05320 | Autoimmune thyroid disease | 0.00058323 | 4 |
| hsa05416 | Viral myocarditis | 0.000866939 | 4 |
| hsa05140 | Leishmaniasis | 0.001947315 | 4 |
| hsa04514 | Cell adhesion molecules | 0.017598597 | 4 |
| hsa04217 | Necroptosis | 0.017598597 | 4 |
| hsa05203 | Viral carcinogenesis | 0.034850106 | 4 |
| hsa05417 | Lipid and atherosclerosis | 0.03767467 | 4 |
| hsa05208 | Chemical carcinogenesis - reactive oxygen species | 0.040062339 | 4 |
| GO:0023026 | MHC class II protein complex binding | 0.005159627 | 3 |
| GO:0023023 | MHC protein complex binding | 0.008256431 | 3 |
| GO:0002399 | MHC class II protein complex assembly | 0.000522422 | 3 |
| GO:0002503 | peptide antigen assembly with MHC class II protei | 0.000522422 | 3 |
| GO:0002396 | MHC protein complex assembly | 0.000893483 | 3 |
| GO:0002501 | peptide antigen assembly with MHC protein compl | 0.000893483 | 3 |
| GO:0019886 | antigen processing and presentation of exogenous | 0.002902508 | 3 |
| GO:0001916 | positive regulation of T cell mediated cytotoxicity | 0.003097152 | 3 |
| GO:0002495 | antigen processing and presentation of peptide ant | 0.003836519 | 3 |
| GO:0002504 | antigen processing and presentation of peptide or | 0.004418827 | 3 |
| GO:0001914 | regulation of T cell mediated cytotoxicity | 0.006340618 | 3 |
| GO:0001913 | T cell mediated cytotoxicity | 0.010296092 | 3 |
| GO:0009409 | response to cold | 0.011252361 | 3 |
| GO:0001912 | positive regulation of leukocyte mediated cytotoxi | 0.013434244 | 3 |
| GO:0031343 | positive regulation of cell killing | 0.016133813 | 3 |
| GO:0006949 | syncytium formation | 0.016741833 | 3 |
| GO:0002381 | immunoglobulin production involved in immunoglo | 0.020336025 | 3 |
| GO:0060337 | type I interferon signaling pathway | 0.020753884 | 3 |
| GO:0045824 | negative regulation of innate immune response | 0.021376702 | 3 |
| GO:0002720 | positive regulation of cytokine production involved | 0.022318834 | 3 |
| GO:0031397 | negative regulation of protein ubiquitination | 0.024402165 | 3 |
| GO:0001910 | regulation of leukocyte mediated cytotoxicity | 0.026261277 | 3 |
| GO:1903321 | negative regulation of protein modification by sma | 0.03260569 | 3 |
| GO:0031341 | regulation of cell killing | 0.034757298 | 3 |
| GO:0030593 | neutrophil chemotaxis | 0.03666947 | 3 |
| GO:0008637 | apoptotic mitochondrial changes | 0.037059077 | 3 |
| GO:0002832 | negative regulation of response to biotic stimulus | 0.04398399 | 3 |
| GO:0002718 | regulation of cytokine production involved in immu | 0.045241338 | 3 |
| GO:0002367 | cytokine production involved in immune response | 0.046968111 | 3 |
| GO:0006413 | translational initiation | 0.049853972 | 3 |
| hsa03250 | Viral life cycle - HIV-1 | 0.012687806 | 3 |
| hsa05321 | Inflammatory bowel disease | 0.013154744 | 3 |
| hsa04622 | RIG-I-like receptor signaling pathway | 0.016106284 | 3 |
| hsa04658 | Th1 and Th2 cell differentiation | 0.029184578 | 3 |
| hsa04640 | Hematopoietic cell lineage | 0.032579401 | 3 |
| hsa04061 | Viral protein interaction with cytokine and cytokir | 0.032579401 | 3 |
| hsa04659 | Th17 cell differentiation | 0.037433693 | 3 |
| hsa05145 | Toxoplasmosis | 0.03767467 | 3 |
| hsa04668 | TNF signaling pathway | 0.039227475 | 3 |
| GO:0042608 | T cell receptor binding | 0.016058555 | 2 |
| GO:0042118 | endothelial cell activation | 0.011410514 | 2 |
| GO:0030656 | regulation of vitamin metabolic process | 0.0130346 | 2 |
| GO:0032351 | negative regulation of hormone metabolic process | 0.0130346 | 2 |
| GO:0002524 | hypersensitivity | 0.013777535 | 2 |
| GO:0002863 | positive regulation of inflammatory response to an | 0.013777535 | 2 |
| GO:0002475 | antigen processing and presentation via MHC class | 0.015144954 | 2 |
| GO:0002468 | dendritic cell antigen processing and presentation | 0.016133813 | 2 |
| GO:0002864 | regulation of acute inflammatory response to antig | 0.016133813 | 2 |
| GO:0001732 | formation of cytoplasmic translation initiation com | 0.017456612 | 2 |
| GO:0050862 | positive regulation of T cell receptor signaling path | 0.017456612 | 2 |
| GO:0019081 | viral translation | 0.020753884 | 2 |
| GO:0032695 | negative regulation of interleukin-12 production | 0.022318834 | 2 |
| GO:0035455 | response to interferon-alpha | 0.022753521 | 2 |
| GO:0060339 | negative regulation of type I interferon-mediated s | 0.022753521 | 2 |
| GO:0071359 | cellular response to dsRNA | 0.027770964 | 2 |
| GO:0002438 | acute inflammatory response to antigenic stimulus | 0.032903763 | 2 |
| GO:0002726 | positive regulation of T cell cytokine production | 0.032903763 | 2 |

| | | | |
|---|---|---|---|
| GO:0050857 | positive regulation of antigen receptor-mediated s | 0.032903763 | 2 |
| GO:1903901 | negative regulation of viral life cycle | 0.032903763 | 2 |
| GO:0090200 | positive regulation of release of cytochrome c from | 0.034284658 | 2 |
| GO:0002675 | positive regulation of acute inflammatory response | 0.034759126 | 2 |
| GO:0034694 | response to prostaglandin | 0.0394772 | 2 |
| GO:0045070 | positive regulation of viral genome replication | 0.041612029 | 2 |
| hsa05310 | Asthma | 0.032579401 | 2 |

| ID | Description | p.adjust | Count |
|---|---|---|---|
| GO:0002237 | response to molecule of bacterial origin | 2.55791E-05 | 9 |
| GO:0050863 | regulation of T cell activation | 3.2368E-05 | 9 |
| GO:1903037 | regulation of leukocyte cell-cell adhesion | 3.2368E-05 | 9 |
| GO:0007159 | leukocyte cell-cell adhesion | 6.62488E-05 | 9 |
| GO:0022407 | regulation of cell-cell adhesion | 0.000200429 | 9 |
| HALLMARK_INTERFERON_GAMMA_RESPONS | HALLMARK_INTERFERON_GAMMA_RESPONSE | 0.000133489 | 9 |
| GO:0042277 | peptide binding | 8.70015E-05 | 8 |
| GO:0033218 | amide binding | 0.000243192 | 8 |
| hsa04145 | Phagosome | 2.98535E-05 | 8 |
| GO:0019886 | antigen processing and presentation of exogenous | 1.06348E-09 | 7 |
| GO:0002495 | antigen processing and presentation of peptide ant | 1.39499E-09 | 7 |
| GO:0002504 | antigen processing and presentation of peptide or | 1.43789E-09 | 7 |
| GO:0002478 | antigen processing and presentation of exogenous | 2.39115E-09 | 7 |
| GO:0019884 | antigen processing and presentation of exogenous | 8.67151E-09 | 7 |
| GO:0048002 | antigen processing and presentation of peptide ant | 5.08598E-08 | 7 |
| GO:0019882 | antigen processing and presentation | 1.14986E-06 | 7 |
| GO:0016064 | immunoglobulin mediated immune response | 8.91533E-05 | 7 |
| GO:0019724 | B cell mediated immunity | 9.20804E-05 | 7 |
| GO:0050870 | positive regulation of T cell activation | 0.000200429 | 7 |
| GO:1903039 | positive regulation of leukocyte cell-cell adhesion | 0.000305099 | 7 |
| GO:0044403 | biological process involved in symbiotic interaction | 0.000578881 | 7 |
| GO:0019058 | viral life cycle | 0.000665881 | 7 |
| GO:0060326 | cell chemotaxis | 0.000665881 | 7 |
| GO:0022409 | positive regulation of cell-cell adhesion | 0.000683963 | 7 |
| GO:0002440 | production of molecular mediator of immune respo | 0.000772728 | 7 |
| GO:0032496 | response to lipopolysaccharide | 0.000794844 | 7 |
| GO:0002449 | lymphocyte mediated immunity | 0.001161557 | 7 |
| GO:0051098 | regulation of binding | 0.001236241 | 7 |
| GO:0002460 | adaptive immune response based on somatic recon | 0.00127096 | 7 |
| GO:0050900 | leukocyte migration | 0.001661801 | 7 |
| GO:0051251 | positive regulation of lymphocyte activation | 0.001680346 | 7 |
| GO:0030099 | myeloid cell differentiation | 0.001837522 | 7 |
| GO:1903706 | regulation of hemopoiesis | 0.001975654 | 7 |
| GO:0016032 | viral process | 0.002186279 | 7 |
| GO:0002696 | positive regulation of leukocyte activation | 0.003135828 | 7 |
| GO:0002443 | leukocyte mediated immunity | 0.003611753 | 7 |
| GO:0050867 | positive regulation of cell activation | 0.003680445 | 7 |
| GO:0045785 | positive regulation of cell adhesion | 0.004439903 | 7 |
| GO:0019221 | cytokine-mediated signaling pathway | 0.004997903 | 7 |
| hsa05152 | Tuberculosis | 0.000285154 | 7 |
| hsa05169 | Epstein-Barr virus infection | 0.000432084 | 7 |
| HALLMARK_ALLOGRAFT_REJECTION | HALLMARK_ALLOGRAFT_REJECTION | 0.004398438 | 7 |
| GO:0023026 | MHC class II protein complex binding | 1.33254E-08 | 6 |
| GO:0023023 | MHC protein complex binding | 4.30767E-08 | 6 |
| GO:0140375 | immune receptor activity | 8.70015E-05 | 6 |
| GO:0030546 | signaling receptor activator activity | 0.021651637 | 6 |
| GO:0051701 | biological process involved in interaction with host | 0.000600898 | 6 |
| GO:0002573 | myeloid leukocyte differentiation | 0.000772728 | 6 |
| GO:0002377 | immunoglobulin production | 0.000772728 | 6 |
| GO:0071219 | cellular response to molecule of bacterial origin | 0.000982027 | 6 |
| GO:0030595 | leukocyte chemotaxis | 0.001047624 | 6 |
| GO:0071216 | cellular response to biotic stimulus | 0.001504668 | 6 |
| GO:0070663 | regulation of leukocyte proliferation | 0.001661801 | 6 |
| GO:1902105 | regulation of leukocyte differentiation | 0.003507709 | 6 |
| GO:0070661 | leukocyte proliferation | 0.005099307 | 6 |
| GO:1903131 | mononuclear cell differentiation | 0.017829078 | 6 |
| GO:0051051 | negative regulation of transport | 0.018826237 | 6 |
| GO:0050673 | epithelial cell proliferation | 0.019032768 | 6 |
| GO:0001819 | positive regulation of cytokine production | 0.019459721 | 6 |
| hsa04612 | Antigen processing and presentation | 6.39112E-05 | 6 |
| hsa05164 | Influenza A | 0.000739702 | 6 |
| hsa05166 | Human T-cell leukemia virus 1 infection | 0.002466244 | 6 |
| GO:0001664 | G protein-coupled receptor binding | 0.014292692 | 5 |
| GO:0002399 | MHC class II protein complex assembly | 6.34694E-08 | 5 |
| GO:0002503 | peptide antigen assembly with MHC class II protei | 6.34694E-08 | 5 |
| GO:0002396 | MHC protein complex assembly | 1.78926E-07 | 5 |
| GO:0002501 | peptide antigen assembly with MHC protein compl | 1.78926E-07 | 5 |

| | | | |
|---|---|---|---|
| GO:0002381 | immunoglobulin production involved in immunoglo | 9.36683E-05 | 5 |
| GO:0070098 | chemokine-mediated signaling pathway | 0.000212798 | 5 |
| GO:1990868 | response to chemokine | 0.000295913 | 5 |
| GO:1990869 | cellular response to chemokine | 0.000295913 | 5 |
| GO:0030593 | neutrophil chemotaxis | 0.000400419 | 5 |
| GO:0071621 | granulocyte chemotaxis | 0.000773559 | 5 |
| GO:1990266 | neutrophil migration | 0.000773559 | 5 |
| GO:0034341 | response to interferon-gamma | 0.001019275 | 5 |
| GO:0044409 | entry into host | 0.001504668 | 5 |
| GO:0097530 | granulocyte migration | 0.001504668 | 5 |
| GO:0051099 | positive regulation of binding | 0.002241801 | 5 |
| GO:0044000 | movement in host | 0.002576193 | 5 |
| GO:0043393 | regulation of protein binding | 0.003643764 | 5 |
| GO:0045637 | regulation of myeloid cell differentiation | 0.004676973 | 5 |
| GO:0071222 | cellular response to lipopolysaccharide | 0.005291833 | 5 |
| GO:0050670 | regulation of lymphocyte proliferation | 0.007126117 | 5 |
| GO:0097529 | myeloid leukocyte migration | 0.007429107 | 5 |
| GO:0032944 | regulation of mononuclear cell proliferation | 0.007463335 | 5 |
| GO:0006898 | receptor-mediated endocytosis | 0.008921097 | 5 |
| GO:0046651 | lymphocyte proliferation | 0.016488871 | 5 |
| GO:0032943 | mononuclear cell proliferation | 0.01765656 | 5 |
| GO:0010038 | response to metal ion | 0.027904476 | 5 |
| GO:0042176 | regulation of protein catabolic process | 0.027904476 | 5 |
| GO:0050678 | regulation of epithelial cell proliferation | 0.040006391 | 5 |
| GO:0030098 | lymphocyte differentiation | 0.041257354 | 5 |
| GO:0002683 | negative regulation of immune system process | 0.049138469 | 5 |
| GO:0048568 | embryonic organ development | 0.049138469 | 5 |
| hsa05416 | Viral myocarditis | 0.000255809 | 5 |
| hsa05323 | Rheumatoid arthritis | 0.000599001 | 5 |
| hsa05150 | Staphylococcus aureus infection | 0.000614882 | 5 |
| hsa04061 | Viral protein interaction with cytokine and cytokir | 0.000667077 | 5 |
| hsa05145 | Toxoplasmosis | 0.000903669 | 5 |
| hsa04062 | Chemokine signaling pathway | 0.007389148 | 5 |
| hsa05163 | Human cytomegalovirus infection | 0.014299373 | 5 |
| hsa05132 | Salmonella infection | 0.02059587 | 5 |
| hsa04060 | Cytokine-cytokine receptor interaction | 0.039061823 | 5 |
| HALLMARK_INTERFERON_ALPHA_RESPONSE | HALLMARK_INTERFERON_ALPHA_RESPONSE | 0.004398438 | 5 |
| GO:0042605 | peptide antigen binding | 8.70015E-05 | 4 |
| GO:0008009 | chemokine activity | 0.000233814 | 4 |
| GO:0042379 | chemokine receptor binding | 0.000799681 | 4 |
| GO:0001540 | amyloid-beta binding | 0.001460296 | 4 |
| GO:0003823 | antigen binding | 0.014292692 | 4 |
| GO:0005125 | cytokine activity | 0.038465521 | 4 |
| GO:0071347 | cellular response to interleukin-1 | 0.004404105 | 4 |
| GO:0002761 | regulation of myeloid leukocyte differentiation | 0.005758409 | 4 |
| GO:0019079 | viral genome replication | 0.007126117 | 4 |
| GO:0070555 | response to interleukin-1 | 0.008919904 | 4 |
| GO:1903900 | regulation of viral life cycle | 0.008921097 | 4 |
| GO:0001890 | placenta development | 0.010427847 | 4 |
| GO:0034614 | cellular response to reactive oxygen species | 0.010427847 | 4 |
| GO:0046718 | viral entry into host cell | 0.010560639 | 4 |
| GO:0048284 | organelle fusion | 0.010841948 | 4 |
| GO:0050792 | regulation of viral process | 0.013111136 | 4 |
| GO:0061025 | membrane fusion | 0.017049251 | 4 |
| GO:2000045 | regulation of G1/S transition of mitotic cell cycle | 0.019044465 | 4 |
| GO:0071248 | cellular response to metal ion | 0.022590941 | 4 |
| GO:0000302 | response to reactive oxygen species | 0.024970332 | 4 |
| GO:1902806 | regulation of cell cycle G1/S phase transition | 0.027904476 | 4 |
| GO:0070374 | positive regulation of ERK1 and ERK2 cascade | 0.030998285 | 4 |
| GO:0071241 | cellular response to inorganic substance | 0.033373518 | 4 |
| GO:0002274 | myeloid leukocyte activation | 0.037086261 | 4 |
| GO:0000082 | G1/S transition of mitotic cell cycle | 0.039756411 | 4 |
| GO:0010951 | negative regulation of endopeptidase activity | 0.040006391 | 4 |
| GO:0034612 | response to tumor necrosis factor | 0.040864203 | 4 |
| GO:0010466 | negative regulation of peptidase activity | 0.044393529 | 4 |
| hsa05310 | Asthma | 0.000285154 | 4 |
| hsa05330 | Allograft rejection | 0.000432084 | 4 |
| hsa05219 | Bladder cancer | 0.000474272 | 4 |

| | | | |
|---|---|---|---|
| hsa05332 | Graft-versus-host disease | 0.000474272 | 4 |
| hsa04940 | Type I diabetes mellitus | 0.000474272 | 4 |
| hsa04672 | Intestinal immune network for IgA production | 0.000614882 | 4 |
| hsa05320 | Autoimmune thyroid disease | 0.000727482 | 4 |
| hsa05321 | Inflammatory bowel disease | 0.001350855 | 4 |
| hsa05140 | Leishmaniasis | 0.00246529 | 4 |
| hsa04658 | Th1 and Th2 cell differentiation | 0.004397837 | 4 |
| hsa05215 | Prostate cancer | 0.00507397 | 4 |
| hsa01522 | Endocrine resistance | 0.00507397 | 4 |
| hsa04640 | Hematopoietic cell lineage | 0.00507397 | 4 |
| hsa04659 | Th17 cell differentiation | 0.006745451 | 4 |
| hsa05322 | Systemic lupus erythematosus | 0.014438924 | 4 |
| hsa04514 | Cell adhesion molecules | 0.022486973 | 4 |
| hsa05202 | Transcriptional misregulation in cancer | 0.040682165 | 4 |
| hsa05167 | Kaposi sarcoma-associated herpesvirus infection | 0.040682165 | 4 |
| GO:0032395 | MHC class II receptor activity | 8.70015E-05 | 3 |
| GO:0001618 | virus receptor activity | 0.014292692 | 3 |
| GO:0140272 | exogenous protein binding | 0.014292692 | 3 |
| GO:1900120 | regulation of receptor binding | 0.001029271 | 3 |
| GO:1904645 | response to amyloid-beta | 0.007820806 | 3 |
| GO:0043388 | positive regulation of DNA binding | 0.008573867 | 3 |
| GO:0046686 | response to cadmium ion | 0.009457982 | 3 |
| GO:0032092 | positive regulation of protein binding | 0.020838658 | 3 |
| GO:0033077 | T cell differentiation in thymus | 0.020838658 | 3 |
| GO:0001892 | embryonic placenta development | 0.021123818 | 3 |
| GO:0045069 | regulation of viral genome replication | 0.021123818 | 3 |
| GO:0032091 | negative regulation of protein binding | 0.026732737 | 3 |
| GO:0070664 | negative regulation of leukocyte proliferation | 0.026732737 | 3 |
| GO:0030316 | osteoclast differentiation | 0.028453881 | 3 |
| GO:1902106 | negative regulation of leukocyte differentiation | 0.033373518 | 3 |
| GO:1905818 | regulation of chromosome separation | 0.035778264 | 3 |
| GO:0002526 | acute inflammatory response | 0.037656094 | 3 |
| GO:1903707 | negative regulation of hemopoiesis | 0.037656094 | 3 |
| GO:0000079 | regulation of cyclin-dependent protein serine/thre | 0.03960012 | 3 |
| GO:0043200 | response to amino acid | 0.03960012 | 3 |
| GO:0071346 | cellular response to interferon-gamma | 0.03960012 | 3 |
| GO:0006906 | vesicle fusion | 0.039756411 | 3 |
| GO:1904029 | regulation of cyclin-dependent protein kinase activ | 0.039756411 | 3 |
| GO:0031623 | receptor internalization | 0.039756411 | 3 |
| GO:0051101 | regulation of DNA binding | 0.039756411 | 3 |
| GO:0090174 | organelle membrane fusion | 0.039756411 | 3 |
| GO:0030218 | erythrocyte differentiation | 0.044038339 | 3 |
| GO:0050868 | negative regulation of T cell activation | 0.044038339 | 3 |
| GO:0048565 | digestive tract development | 0.047046034 | 3 |
| GO:0051983 | regulation of chromosome segregation | 0.047046034 | 3 |
| GO:0001101 | response to acid chemical | 0.047732171 | 3 |
| GO:0051304 | chromosome separation | 0.048796881 | 3 |
| GO:0034101 | erythrocyte homeostasis | 0.049138469 | 3 |
| hsa04657 | IL-17 signaling pathway | 0.036096735 | 3 |
| GO:0050998 | nitric-oxide synthase binding | 0.009616875 | 2 |
| GO:0045236 | CXCR chemokine receptor binding | 0.014292692 | 2 |
| GO:0016814 | hydrolase activity, acting on carbon-nitrogen (but | 0.043285869 | 2 |
| GO:0002604 | regulation of dendritic cell antigen processing and | 0.007696067 | 2 |
| GO:2001198 | regulation of dendritic cell differentiation | 0.009904155 | 2 |
| GO:0070141 | response to UV-A | 0.010841948 | 2 |
| GO:0002468 | dendritic cell antigen processing and presentation | 0.012341865 | 2 |
| GO:0009263 | deoxyribonucleotide biosynthetic process | 0.013442999 | 2 |
| GO:0009265 | 2'-deoxyribonucleotide biosynthetic process | 0.013442999 | 2 |
| GO:0046385 | deoxyribose phosphate biosynthetic process | 0.013442999 | 2 |
| GO:0046653 | tetrahydrofolate metabolic process | 0.016707079 | 2 |
| GO:0002577 | regulation of antigen processing and presentation | 0.01919057 | 2 |
| GO:0050765 | negative regulation of phagocytosis | 0.021904613 | 2 |
| GO:0090023 | positive regulation of neutrophil chemotaxis | 0.027004826 | 2 |
| GO:0019883 | antigen processing and presentation of endogenou | 0.028166774 | 2 |
| GO:0006760 | folic acid-containing compound metabolic process | 0.029325734 | 2 |
| GO:0010460 | positive regulation of heart rate | 0.029325734 | 2 |
| GO:1905820 | positive regulation of chromosome separation | 0.029325734 | 2 |
| GO:0071624 | positive regulation of granulocyte chemotaxis | 0.030998285 | 2 |

| | | | |
|---|---|---|---|
| GO:0006221 | pyrimidine nucleotide biosynthetic process | 0.038160408 | 2 |
| GO:0002431 | Fc receptor mediated stimulatory signaling pathwa | 0.03960012 | 2 |
| GO:1902253 | regulation of intrinsic apoptotic signaling pathway | 0.03960012 | 2 |
| GO:0042558 | pteridine-containing compound metabolic process | 0.039756411 | 2 |
| GO:0090022 | regulation of neutrophil chemotaxis | 0.039756411 | 2 |
| GO:1902624 | positive regulation of neutrophil migration | 0.039756411 | 2 |
| GO:0071276 | cellular response to cadmium ion | 0.043832052 | 2 |
| GO:0072528 | pyrimidine-containing compound biosynthetic proc | 0.043832052 | 2 |
| GO:0045823 | positive regulation of heart contraction | 0.048425774 | 2 |
| GO:0140467 | integrated stress response signaling | 0.048425774 | 2 |
| GO:0001914 | regulation of T cell mediated cytotoxicity | 0.049138469 | 2 |
| GO:0042119 | neutrophil activation | 0.049138469 | 2 |
| GO:1901031 | regulation of response to reactive oxygen species | 0.049138469 | 2 |
| GO:1903524 | positive regulation of blood circulation | 0.049138469 | 2 |

| ID | Description | p.adjust | Count |
|---|---|---|---|
| HALLMARK_INTERFERON_GAMMA_RESPONS | HALLMARK_INTERFERON_GAMMA_RESPONSE | 1.42297E-16 | 17 |
| GO:0009615 | response to virus | 3.43379E-10 | 14 |
| GO:0051607 | defense response to virus | 3.50113E-08 | 11 |
| GO:0140546 | defense response to symbiont | 3.50113E-08 | 11 |
| GO:0005125 | cytokine activity | 2.86946E-08 | 10 |
| GO:0048018 | receptor ligand activity | 7.75071E-06 | 10 |
| GO:0030546 | signaling receptor activator activity | 7.75071E-06 | 10 |
| hsa04060 | Cytokine-cytokine receptor interaction | 1.20012E-05 | 10 |
| GO:0001819 | positive regulation of cytokine production | 0.000329996 | 9 |
| GO:0019221 | cytokine-mediated signaling pathway | 0.00033456 | 9 |
| HALLMARK_INTERFERON_ALPHA_RESPONSE | HALLMARK_INTERFERON_ALPHA_RESPONSE | 1.46937E-08 | 9 |
| GO:0005126 | cytokine receptor binding | 7.75071E-06 | 8 |
| GO:0032103 | positive regulation of response to external stimulu | 0.001150158 | 8 |
| GO:0071346 | cellular response to interferon-gamma | 3.33139E-06 | 7 |
| GO:0034341 | response to interferon-gamma | 8.74155E-06 | 7 |
| GO:1903131 | mononuclear cell differentiation | 0.004920559 | 7 |
| GO:0030003 | cellular cation homeostasis | 0.005541171 | 7 |
| HALLMARK_INFLAMMATORY_RESPONSE | HALLMARK_INFLAMMATORY_RESPONSE | 0.000586598 | 7 |
| GO:0003924 | GTPase activity | 0.002641112 | 6 |
| GO:0005525 | GTP binding | 0.004887005 | 6 |
| GO:0019001 | guanyl nucleotide binding | 0.004956063 | 6 |
| GO:0032561 | guanyl ribonucleotide binding | 0.004956063 | 6 |
| GO:0070663 | regulation of leukocyte proliferation | 0.002401872 | 6 |
| GO:0031349 | positive regulation of defense response | 0.004003679 | 6 |
| GO:0006959 | humoral immune response | 0.004837323 | 6 |
| GO:0072503 | cellular divalent inorganic cation homeostasis | 0.00490307 | 6 |
| GO:0070661 | leukocyte proliferation | 0.005541171 | 6 |
| GO:0072507 | divalent inorganic cation homeostasis | 0.005825101 | 6 |
| GO:0002460 | adaptive immune response based on somatic recon | 0.006254728 | 6 |
| GO:0002253 | activation of immune response | 0.007037407 | 6 |
| GO:0006875 | cellular metal ion homeostasis | 0.009176697 | 6 |
| GO:0002696 | positive regulation of leukocyte activation | 0.011733822 | 6 |
| GO:0050867 | positive regulation of cell activation | 0.012197013 | 6 |
| hsa04061 | Viral protein interaction with cytokine and cytokir | 0.000100231 | 6 |
| GO:0008009 | chemokine activity | 7.75071E-06 | 5 |
| GO:0042379 | chemokine receptor binding | 1.96499E-05 | 5 |
| GO:0001664 | G protein-coupled receptor binding | 0.007695318 | 5 |
| GO:0070098 | chemokine-mediated signaling pathway | 0.000349735 | 5 |
| GO:1990868 | response to chemokine | 0.000426421 | 5 |
| GO:1990869 | cellular response to chemokine | 0.000426421 | 5 |
| GO:0030593 | neutrophil chemotaxis | 0.000570577 | 5 |
| GO:0071621 | granulocyte chemotaxis | 0.001215068 | 5 |
| GO:1990266 | neutrophil migration | 0.001215068 | 5 |
| GO:0070555 | response to interleukin-1 | 0.001575564 | 5 |
| GO:0050729 | positive regulation of inflammatory response | 0.002267563 | 5 |
| GO:0097530 | granulocyte migration | 0.002376155 | 5 |
| GO:0071356 | cellular response to tumor necrosis factor | 0.006254728 | 5 |
| GO:0050670 | regulation of lymphocyte proliferation | 0.006254728 | 5 |
| GO:0030595 | leukocyte chemotaxis | 0.006254728 | 5 |
| GO:0097529 | myeloid leukocyte migration | 0.006254728 | 5 |
| GO:0032944 | regulation of mononuclear cell proliferation | 0.006254728 | 5 |
| GO:0034612 | response to tumor necrosis factor | 0.007037407 | 5 |
| GO:0006874 | cellular calcium ion homeostasis | 0.010589688 | 5 |
| GO:0046651 | lymphocyte proliferation | 0.012158505 | 5 |
| GO:0055074 | calcium ion homeostasis | 0.012410154 | 5 |
| GO:0032943 | mononuclear cell proliferation | 0.012801532 | 5 |
| GO:0060326 | cell chemotaxis | 0.012871404 | 5 |
| GO:0032496 | response to lipopolysaccharide | 0.015656295 | 5 |
| GO:0009306 | protein secretion | 0.017994871 | 5 |
| GO:0002237 | response to molecule of bacterial origin | 0.017994871 | 5 |
| GO:0035592 | establishment of protein localization to extracellul | 0.017994871 | 5 |
| GO:0002831 | regulation of response to biotic stimulus | 0.018050718 | 5 |
| GO:0071692 | protein localization to extracellular region | 0.018953944 | 5 |
| GO:0050863 | regulation of T cell activation | 0.02012401 | 5 |
| GO:0002697 | regulation of immune effector process | 0.020489389 | 5 |
| GO:0050900 | leukocyte migration | 0.022710235 | 5 |
| GO:0051251 | positive regulation of lymphocyte activation | 0.02303788 | 5 |

| | | | |
|---|---|---|---|
| GO:0050727 | regulation of inflammatory response | 0.025780605 | 5 |
| GO:0030098 | lymphocyte differentiation | 0.026767264 | 5 |
| GO:0052548 | regulation of endopeptidase activity | 0.028857835 | 5 |
| GO:0006816 | calcium ion transport | 0.030637799 | 5 |
| GO:0052547 | regulation of peptidase activity | 0.03551093 | 5 |
| hsa04620 | Toll-like receptor signaling pathway | 0.001500455 | 5 |
| hsa05164 | Influenza A | 0.009303637 | 5 |
| hsa04621 | NOD-like receptor signaling pathway | 0.011247979 | 5 |
| hsa04062 | Chemokine signaling pathway | 0.011247979 | 5 |
| HALLMARK_IL6_JAK_STAT3_SIGNALING | HALLMARK_IL6_JAK_STAT3_SIGNALING | 0.000586598 | 5 |
| HALLMARK_IL2_STAT5_SIGNALING | HALLMARK_IL2_STAT5_SIGNALING | 0.020390094 | 5 |
| GO:0005539 | glycosaminoglycan binding | 0.025077189 | 4 |
| GO:0048247 | lymphocyte chemotaxis | 0.001417258 | 4 |
| GO:0043367 | CD4-positive, alpha-beta T cell differentiation | 0.002934468 | 4 |
| GO:0071347 | cellular response to interleukin-1 | 0.005436953 | 4 |
| GO:0035710 | CD4-positive, alpha-beta T cell activation | 0.005541171 | 4 |
| GO:0046632 | alpha-beta T cell differentiation | 0.005762122 | 4 |
| GO:0051209 | release of sequestered calcium ion into cytosol | 0.006254728 | 4 |
| GO:0072676 | lymphocyte migration | 0.006254728 | 4 |
| GO:0051283 | negative regulation of sequestering of calcium ion | 0.006254728 | 4 |
| GO:0051282 | regulation of sequestering of calcium ion | 0.006254728 | 4 |
| GO:0019079 | viral genome replication | 0.006254728 | 4 |
| GO:0051208 | sequestering of calcium ion | 0.006254728 | 4 |
| GO:0046631 | alpha-beta T cell activation | 0.012197013 | 4 |
| GO:0002822 | regulation of adaptive immune response based on | 0.012871404 | 4 |
| GO:0042129 | regulation of T cell proliferation | 0.012871404 | 4 |
| GO:0097553 | calcium ion transmembrane import into cytosol | 0.012871404 | 4 |
| GO:0002819 | regulation of adaptive immune response | 0.016063649 | 4 |
| GO:1903828 | negative regulation of protein localization | 0.017257353 | 4 |
| GO:0071674 | mononuclear cell migration | 0.017508603 | 4 |
| GO:0045619 | regulation of lymphocyte differentiation | 0.017994871 | 4 |
| GO:0042098 | T cell proliferation | 0.018055692 | 4 |
| GO:0002573 | myeloid leukocyte differentiation | 0.019305297 | 4 |
| GO:0071222 | cellular response to lipopolysaccharide | 0.02012401 | 4 |
| GO:0051651 | maintenance of location in cell | 0.02089914 | 4 |
| GO:0071219 | cellular response to molecule of bacterial origin | 0.022456017 | 4 |
| GO:0002274 | myeloid leukocyte activation | 0.022664322 | 4 |
| GO:0045088 | regulation of innate immune response | 0.023167684 | 4 |
| GO:0071216 | cellular response to biotic stimulus | 0.029196552 | 4 |
| GO:0002366 | leukocyte activation involved in immune response | 0.039199602 | 4 |
| GO:0030217 | T cell differentiation | 0.039922675 | 4 |
| GO:0002263 | cell activation involved in immune response | 0.039922675 | 4 |
| GO:1902105 | regulation of leukocyte differentiation | 0.048171294 | 4 |
| GO:0019058 | viral life cycle | 0.048960801 | 4 |
| hsa04630 | JAK-STAT signaling pathway | 0.044858828 | 4 |
| HALLMARK_APOPTOSIS | HALLMARK_APOPTOSIS | 0.044505719 | 4 |
| GO:0045236 | CXCR chemokine receptor binding | 0.000239525 | 3 |
| GO:0042832 | defense response to protozoan | 0.002376155 | 3 |
| GO:0001562 | response to protozoan | 0.002403926 | 3 |
| GO:0002825 | regulation of T-helper 1 type immune response | 0.002834575 | 3 |
| GO:0002230 | positive regulation of defense response to virus by | 0.002934468 | 3 |
| GO:0051281 | positive regulation of release of sequestered calciu | 0.005436953 | 3 |
| GO:0032735 | positive regulation of interleukin-12 production | 0.005436953 | 3 |
| GO:0050691 | regulation of defense response to virus by host | 0.005436953 | 3 |
| GO:0045622 | regulation of T-helper cell differentiation | 0.005541171 | 3 |
| GO:0042088 | T-helper 1 type immune response | 0.005825101 | 3 |
| GO:0097028 | dendritic cell differentiation | 0.006056869 | 3 |
| GO:0043370 | regulation of CD4-positive, alpha-beta T cell differe | 0.007037407 | 3 |
| GO:0032615 | interleukin-12 production | 0.009176697 | 3 |
| GO:0032655 | regulation of interleukin-12 production | 0.009176697 | 3 |
| GO:0032731 | positive regulation of interleukin-1 beta productio | 0.009176697 | 3 |
| GO:0042093 | T-helper cell differentiation | 0.011134181 | 3 |
| GO:0002294 | CD4-positive, alpha-beta T cell differentiation invol | 0.011733822 | 3 |
| GO:0002287 | alpha-beta T cell activation involved in immune re: | 0.011733822 | 3 |
| GO:0002293 | alpha-beta T cell differentiation involved in immun | 0.011733822 | 3 |
| GO:0002548 | monocyte chemotaxis | 0.011733822 | 3 |
| GO:0019362 | pyridine nucleotide metabolic process | 0.012039441 | 3 |
| GO:0046496 | nicotinamide nucleotide metabolic process | 0.012039441 | 3 |

| | | | |
|---|---|---|---|
| GO:0046637 | regulation of alpha-beta T cell differentiation | 0.012039441 | 3 |
| GO:0032732 | positive regulation of interleukin-1 production | 0.012158505 | 3 |
| GO:0050688 | regulation of defense response to virus | 0.012158505 | 3 |
| GO:0002292 | T cell differentiation involved in immune response | 0.012871404 | 3 |
| GO:0072524 | pyridine-containing compound metabolic process | 0.012871404 | 3 |
| GO:2000514 | regulation of CD4-positive, alpha-beta T cell activa | 0.012871404 | 3 |
| GO:0061844 | antimicrobial humoral immune response mediated | 0.013404638 | 3 |
| GO:0051279 | regulation of release of sequestered calcium ion int | 0.013738509 | 3 |
| GO:0034103 | regulation of tissue remodeling | 0.014920234 | 3 |
| GO:1904427 | positive regulation of calcium ion transmembrane t | 0.017508603 | 3 |
| GO:0032760 | positive regulation of tumor necrosis factor produc | 0.02089914 | 3 |
| GO:1903557 | positive regulation of tumor necrosis factor superfa | 0.022338312 | 3 |
| GO:0032611 | interleukin-1 beta production | 0.022664322 | 3 |
| GO:0032651 | regulation of interleukin-1 beta production | 0.022664322 | 3 |
| GO:0046634 | regulation of alpha-beta T cell activation | 0.023944795 | 3 |
| GO:0002286 | T cell activation involved in immune response | 0.025574983 | 3 |
| GO:0019730 | antimicrobial humoral response | 0.028857835 | 3 |
| GO:0051224 | negative regulation of protein transport | 0.030637799 | 3 |
| GO:0051928 | positive regulation of calcium ion transport | 0.030637799 | 3 |
| GO:0032612 | interleukin-1 production | 0.030637799 | 3 |
| GO:0032652 | regulation of interleukin-1 production | 0.030637799 | 3 |
| GO:0045621 | positive regulation of lymphocyte differentiation | 0.030637799 | 3 |
| GO:1904950 | negative regulation of establishment of protein loc | 0.032147895 | 3 |
| GO:0010977 | negative regulation of neuron projection developm | 0.037178554 | 3 |
| GO:0050921 | positive regulation of chemotaxis | 0.038419731 | 3 |
| GO:0045089 | positive regulation of innate immune response | 0.038927925 | 3 |
| GO:0050671 | positive regulation of lymphocyte proliferation | 0.039199602 | 3 |
| GO:0002687 | positive regulation of leukocyte migration | 0.039922675 | 3 |
| GO:0032946 | positive regulation of mononuclear cell proliferatic | 0.040262314 | 3 |
| GO:0001890 | placenta development | 0.040725785 | 3 |
| GO:0007189 | adenylate cyclase-activating G protein-coupled rec | 0.045476978 | 3 |
| hsa05143 | African trypanosomiasis | 0.009069054 | 3 |
| GO:0008603 | cAMP-dependent protein kinase regulator activity | 0.004956063 | 2 |
| GO:0016702 | oxidoreductase activity, acting on single donors wi | 0.017627269 | 2 |
| GO:0016701 | oxidoreductase activity, acting on single donors wi | 0.017855658 | 2 |
| GO:0045625 | regulation of T-helper 1 cell differentiation | 0.006254728 | 2 |
| GO:0006568 | tryptophan metabolic process | 0.007037407 | 2 |
| GO:0070189 | kynurenine metabolic process | 0.007037407 | 2 |
| GO:0006586 | indolalkylamine metabolic process | 0.009176697 | 2 |
| GO:0009074 | aromatic amino acid family catabolic process | 0.011956792 | 2 |
| GO:0002830 | positive regulation of type 2 immune response | 0.012871404 | 2 |
| GO:0043011 | myeloid dendritic cell differentiation | 0.01452406 | 2 |
| GO:0045063 | T-helper 1 cell differentiation | 0.015523006 | 2 |
| GO:1901739 | regulation of myoblast fusion | 0.015523006 | 2 |
| GO:0044546 | NLRP3 inflammasome complex assembly | 0.01652814 | 2 |
| GO:0044827 | modulation by host of viral genome replication | 0.017508603 | 2 |
| GO:0045624 | positive regulation of T-helper cell differentiation | 0.017508603 | 2 |
| GO:0042430 | indole-containing compound metabolic process | 0.017994871 | 2 |
| GO:0140632 | inflammasome complex assembly | 0.017994871 | 2 |
| GO:0032693 | negative regulation of interleukin-10 production | 0.018533409 | 2 |
| GO:0032816 | positive regulation of natural killer cell activation | 0.018533409 | 2 |
| GO:0009435 | NAD biosynthetic process | 0.019529729 | 2 |
| GO:0042537 | benzene-containing compound metabolic process | 0.020489389 | 2 |
| GO:0010818 | T cell chemotaxis | 0.021016944 | 2 |
| GO:0019359 | nicotinamide nucleotide biosynthetic process | 0.021016944 | 2 |
| GO:0019363 | pyridine nucleotide biosynthetic process | 0.021016944 | 2 |
| GO:0001773 | myeloid dendritic cell activation | 0.022338312 | 2 |
| GO:0009072 | aromatic amino acid family metabolic process | 0.022664322 | 2 |
| GO:0060142 | regulation of syncytium formation by plasma meml | 0.022664322 | 2 |
| GO:0072525 | pyridine-containing compound biosynthetic process | 0.023485311 | 2 |
| GO:0002828 | regulation of type 2 immune response | 0.025834297 | 2 |
| GO:0043372 | positive regulation of CD4-positive, alpha-beta T ce | 0.028620753 | 2 |
| GO:0033280 | response to vitamin D | 0.02943626 | 2 |
| GO:0042092 | type 2 immune response | 0.03126172 | 2 |
| GO:0071634 | regulation of transforming growth factor beta proc | 0.032481154 | 2 |
| GO:0045124 | regulation of bone resorption | 0.03551093 | 2 |
| GO:0032814 | regulation of natural killer cell activation | 0.036635347 | 2 |
| GO:0071604 | transforming growth factor beta production | 0.036635347 | 2 |

| | | | |
|---|---|---|---|
| GO:2000516 | positive regulation of CD4-positive, alpha-beta T ce | 0.039922675 | 2 |
| GO:0044788 | modulation by host of viral process | 0.040725785 | 2 |
| GO:0046850 | regulation of bone remodeling | 0.043998852 | 2 |
| GO:0007520 | myoblast fusion | 0.047097467 | 2 |

| ID | Description | p.adjust | Count |
|---|---|---|---|
| GO:0007159 | leukocyte cell-cell adhesion | 0.000369824 | 9 |
| HALLMARK_TNFA_SIGNALING_VIA_NFKB | HALLMARK_TNFA_SIGNALING_VIA_NFKB | 0.000119807 | 8 |
| GO:0001228 | DNA-binding transcription activator activity, RNA | | 0.010663589 | 7 |
| GO:0001216 | DNA-binding transcription activator activity | 0.010663589 | 7 |
| GO:1903037 | regulation of leukocyte cell-cell adhesion | 0.012290484 | 7 |
| GO:0022407 | regulation of cell-cell adhesion | 0.018433473 | 7 |
| GO:0140297 | DNA-binding transcription factor binding | 0.026822252 | 6 |
| GO:1903039 | positive regulation of leukocyte cell-cell adhesion | 0.012719585 | 6 |
| GO:0022409 | positive regulation of cell-cell adhesion | 0.0183969 | 6 |
| GO:0071496 | cellular response to external stimulus | 0.0183969 | 6 |
| GO:0030098 | lymphocyte differentiation | 0.035783171 | 6 |
| GO:1903131 | mononuclear cell differentiation | 0.049287807 | 6 |
| GO:0061629 | RNA polymerase II-specific DNA-binding transcriptio | 0.031039925 | 5 |
| GO:0050870 | positive regulation of T cell activation | 0.033520407 | 5 |
| GO:0030217 | T cell differentiation | 0.045861691 | 5 |
| hsa04514 | Cell adhesion molecules | 0.010961317 | 5 |
| HALLMARK_ALLOGRAFT_REJECTION | HALLMARK_ALLOGRAFT_REJECTION | 0.04359997 | 5 |
| GO:0001975 | response to amphetamine | 0.012719585 | 3 |
| GO:0014075 | response to amine | 0.020290762 | 3 |
| GO:0061756 | leukocyte adhesion to vascular endothelial cell | 0.035131942 | 3 |
| GO:0034113 | heterotypic cell-cell adhesion | 0.035783171 | 3 |
| GO:0045123 | cellular extravasation | 0.049287807 | 3 |
| GO:0035259 | nuclear glucocorticoid receptor binding | 0.020562537 | 2 |
| GO:0045028 | G protein-coupled purinergic nucleotide receptor a | 0.020562537 | 2 |
| GO:0001614 | purinergic nucleotide receptor activity | 0.026822252 | 2 |
| GO:0016502 | nucleotide receptor activity | 0.026822252 | 2 |
| GO:0035589 | G protein-coupled purinergic nucleotide receptor si | 0.041484093 | 2 |

| ID | Description | p.adjust | Count |
|---|---|---|---|
| HALLMARK_TNFA_SIGNALING_VIA_NFKB | HALLMARK_TNFA_SIGNALING_VIA_NFKB | 2.63286E-08 | 12 |
| HALLMARK_INFLAMMATORY_RESPONSE | HALLMARK_INFLAMMATORY_RESPONSE | 0.000215414 | 8 |
| HALLMARK_INTERFERON_GAMMA_RESPONS | HALLMARK_INTERFERON_GAMMA_RESPONSE | 0.000215414 | 8 |
| GO:0051607 | defense response to virus | 0.011891155 | 6 |
| GO:0140546 | defense response to symbiont | 0.011891155 | 6 |
| GO:0001933 | negative regulation of protein phosphorylation | 0.015728925 | 6 |
| GO:0042326 | negative regulation of phosphorylation | 0.015728925 | 6 |
| GO:0050900 | leukocyte migration | 0.016299341 | 6 |
| GO:0009615 | response to virus | 0.016299341 | 6 |
| GO:0045936 | negative regulation of phosphate metabolic process | 0.018498522 | 6 |
| GO:0010563 | negative regulation of phosphorus metabolic process | 0.018498522 | 6 |
| GO:0001659 | temperature homeostasis | 0.011891155 | 5 |
| GO:0030595 | leukocyte chemotaxis | 0.015728925 | 5 |
| GO:0045444 | fat cell differentiation | 0.015728925 | 5 |
| GO:0060326 | cell chemotaxis | 0.024691581 | 5 |
| GO:0062012 | regulation of small molecule metabolic process | 0.028796564 | 5 |
| GO:0071900 | regulation of protein serine/threonine kinase activity | 0.041614667 | 5 |
| GO:0045786 | negative regulation of cell cycle | 0.041614667 | 5 |
| HALLMARK_INTERFERON_ALPHA_RESPONSE | HALLMARK_INTERFERON_ALPHA_RESPONSE | 0.002199126 | 5 |
| HALLMARK_HYPOXIA | HALLMARK_HYPOXIA | 0.043891068 | 5 |
| GO:1990748 | cellular detoxification | 0.015728925 | 4 |
| GO:0071901 | negative regulation of protein serine/threonine kinase | 0.015728925 | 4 |
| GO:0097237 | cellular response to toxic substance | 0.015728925 | 4 |
| GO:0034341 | response to interferon-gamma | 0.016299341 | 4 |
| GO:0106106 | cold-induced thermogenesis | 0.016299341 | 4 |
| GO:0120161 | regulation of cold-induced thermogenesis | 0.016299341 | 4 |
| GO:0098754 | detoxification | 0.018098314 | 4 |
| GO:1990845 | adaptive thermogenesis | 0.018498522 | 4 |
| GO:0006937 | regulation of muscle contraction | 0.022465904 | 4 |
| GO:0006469 | negative regulation of protein kinase activity | 0.034916516 | 4 |
| GO:0072593 | reactive oxygen species metabolic process | 0.041614667 | 4 |
| GO:0033673 | negative regulation of kinase activity | 0.041614667 | 4 |
| GO:0097529 | myeloid leukocyte migration | 0.041614667 | 4 |
| GO:0045926 | negative regulation of growth | 0.043126165 | 4 |
| GO:0009636 | response to toxic substance | 0.043126165 | 4 |
| GO:0090257 | regulation of muscle system process | 0.045715926 | 4 |
| GO:1903522 | regulation of blood circulation | 0.047811008 | 4 |
| GO:0035455 | response to interferon-alpha | 0.010330399 | 3 |
| GO:0046597 | negative regulation of viral entry into host cell | 0.010330399 | 3 |
| GO:1903901 | negative regulation of viral life cycle | 0.011551908 | 3 |
| GO:0035456 | response to interferon-beta | 0.011891155 | 3 |
| GO:0046596 | regulation of viral entry into host cell | 0.015728925 | 3 |
| GO:0045933 | positive regulation of muscle contraction | 0.015728925 | 3 |
| GO:0050873 | brown fat cell differentiation | 0.015728925 | 3 |
| GO:0052372 | modulation by symbiont of entry into host | 0.015728925 | 3 |
| GO:0045071 | negative regulation of viral genome replication | 0.016299341 | 3 |
| GO:0043903 | regulation of biological process involved in symbiotic in | 0.016299341 | 3 |
| GO:0070542 | response to fatty acid | 0.017801715 | 3 |
| GO:0006940 | regulation of smooth muscle contraction | 0.018216374 | 3 |
| GO:0060337 | type I interferon signaling pathway | 0.022465904 | 3 |
| GO:0071357 | cellular response to type I interferon | 0.023620248 | 3 |
| GO:0034340 | response to type I interferon | 0.026791706 | 3 |
| GO:0045069 | regulation of viral genome replication | 0.028796564 | 3 |
| GO:0048525 | negative regulation of viral process | 0.029739576 | 3 |
| GO:0098869 | cellular oxidant detoxification | 0.035399768 | 3 |
| GO:0030593 | neutrophil chemotaxis | 0.039737239 | 3 |
| GO:0062014 | negative regulation of small molecule metabolic proces | 0.041077159 | 3 |
| GO:0006939 | smooth muscle contraction | 0.041614667 | 3 |
| GO:0002526 | acute inflammatory response | 0.041614667 | 3 |
| GO:0045446 | endothelial cell differentiation | 0.044470783 | 3 |
| GO:0019079 | viral genome replication | 0.049137375 | 3 |
| GO:0071621 | granulocyte chemotaxis | 0.049137375 | 3 |
| GO:1990266 | neutrophil migration | 0.049137375 | 3 |
| GO:0050786 | RAGE receptor binding | 0.031496647 | 2 |
| GO:0004861 | cyclin-dependent protein serine/threonine kinase inhib | 0.031496647 | 2 |
| GO:2000271 | positive regulation of fibroblast apoptotic process | 0.015728925 | 2 |
| GO:0042416 | dopamine biosynthetic process | 0.016299341 | 2 |

| | | | |
|---|---|---|---|
| GO:0045986 | negative regulation of smooth muscle contraction | 0.021527176 | 2 |
| GO:0055119 | relaxation of cardiac muscle | 0.022465904 | 2 |
| GO:0086103 | G protein-coupled receptor signaling pathway involvec | 0.023857592 | 2 |
| GO:1903579 | negative regulation of ATP metabolic process | 0.026791706 | 2 |
| GO:2000269 | regulation of fibroblast apoptotic process | 0.026791706 | 2 |
| GO:0009713 | catechol-containing compound biosynthetic process | 0.029093889 | 2 |
| GO:0030728 | ovulation | 0.029093889 | 2 |
| GO:0042423 | catecholamine biosynthetic process | 0.029093889 | 2 |
| GO:1900543 | negative regulation of purine nucleotide metabolic pro | 0.030544962 | 2 |
| GO:0045932 | negative regulation of muscle contraction | 0.03472349 | 2 |
| GO:0045980 | negative regulation of nucleotide metabolic process | 0.03472349 | 2 |
| GO:0044346 | fibroblast apoptotic process | 0.035512763 | 2 |
| GO:0002675 | positive regulation of acute inflammatory response | 0.039737239 | 2 |
| GO:0010575 | positive regulation of vascular endothelial growth fact | 0.041173082 | 2 |
| GO:0001975 | response to amphetamine | 0.041614667 | 2 |
| GO:0045987 | positive regulation of smooth muscle contraction | 0.041614667 | 2 |
| GO:0045736 | negative regulation of cyclin-dependent protein serine, | 0.043126165 | 2 |
| GO:0071875 | adrenergic receptor signaling pathway | 0.043126165 | 2 |
| GO:1904030 | negative regulation of cyclin-dependent protein kinase | 0.043126165 | 2 |
| GO:0071398 | cellular response to fatty acid | 0.045826777 | 2 |
| GO:0090075 | relaxation of muscle | 0.045826777 | 2 |
| GO:0033280 | response to vitamin D | 0.047811008 | 2 |

| ID | Description | p.adjust | Count |
|---|---|---|---|
| GO:0050900 | leukocyte migration | 5.3005E-06 | 10 |
| GO:0007159 | leukocyte cell-cell adhesion | 7.29475E-06 | 10 |
| GO:0006875 | cellular metal ion homeostasis | 7.96201E-06 | 10 |
| GO:0032103 | positive regulation of response to external stimulu | 1.57421E-05 | 10 |
| GO:0030003 | cellular cation homeostasis | 2.68478E-05 | 10 |
| GO:0046916 | cellular transition metal ion homeostasis | 2.64204E-08 | 9 |
| GO:0055076 | transition metal ion homeostasis | 8.77011E-08 | 9 |
| GO:0030595 | leukocyte chemotaxis | 1.53085E-06 | 9 |
| GO:0097529 | myeloid leukocyte migration | 1.53085E-06 | 9 |
| GO:0060326 | cell chemotaxis | 8.58672E-06 | 9 |
| GO:0051235 | maintenance of location | 1.37036E-05 | 9 |
| GO:0001819 | positive regulation of cytokine production | 0.000164869 | 9 |
| GO:0033218 | amide binding | 0.001896902 | 8 |
| GO:0030593 | neutrophil chemotaxis | 1.62975E-07 | 8 |
| GO:0071621 | granulocyte chemotaxis | 3.14203E-07 | 8 |
| GO:1990266 | neutrophil migration | 3.14203E-07 | 8 |
| GO:0097530 | granulocyte migration | 1.17935E-06 | 8 |
| GO:0050863 | regulation of T cell activation | 0.000209271 | 8 |
| GO:1903037 | regulation of leukocyte cell-cell adhesion | 0.000209271 | 8 |
| GO:0051251 | positive regulation of lymphocyte activation | 0.000308425 | 8 |
| GO:0002696 | positive regulation of leukocyte activation | 0.00060299 | 8 |
| GO:0050867 | positive regulation of cell activation | 0.0007585 | 8 |
| GO:0045785 | positive regulation of cell adhesion | 0.000929073 | 8 |
| GO:0022407 | regulation of cell-cell adhesion | 0.000959444 | 8 |
| GO:0019221 | cytokine-mediated signaling pathway | 0.000992784 | 8 |
| HALLMARK_INTERFERON_GAMMA_RESPONS | HALLMARK_INTERFERON_GAMMA_RESPONSE | 0.000735304 | 8 |
| GO:0042277 | peptide binding | 0.002155429 | 7 |
| GO:0051651 | maintenance of location in cell | 8.45556E-05 | 7 |
| GO:0050870 | positive regulation of T cell activation | 0.000164869 | 7 |
| GO:1903039 | positive regulation of leukocyte cell-cell adhesion | 0.000259542 | 7 |
| GO:0044403 | biological process involved in symbiotic interaction | 0.000467387 | 7 |
| GO:0006959 | humoral immune response | 0.000598886 | 7 |
| GO:0022409 | positive regulation of cell-cell adhesion | 0.00060299 | 7 |
| GO:0072503 | cellular divalent inorganic cation homeostasis | 0.00060299 | 7 |
| GO:0072507 | divalent inorganic cation homeostasis | 0.000959444 | 7 |
| GO:0010038 | response to metal ion | 0.000992784 | 7 |
| GO:0002443 | leukocyte mediated immunity | 0.003769158 | 7 |
| hsa04978 | Mineral absorption | 6.93783E-07 | 7 |
| GO:0048018 | receptor ligand activity | 0.015121545 | 6 |
| GO:0030546 | signaling receptor activator activity | 0.015501763 | 6 |
| GO:0002478 | antigen processing and presentation of exogenous | 2.78289E-07 | 6 |
| GO:0006882 | cellular zinc ion homeostasis | 2.78289E-07 | 6 |
| GO:0055069 | zinc ion homeostasis | 3.14203E-07 | 6 |
| GO:0019884 | antigen processing and presentation of exogenous | 5.53437E-07 | 6 |
| GO:0048002 | antigen processing and presentation of peptide ant | 1.85246E-06 | 6 |
| GO:0019882 | antigen processing and presentation | 2.09452E-05 | 6 |
| GO:0019730 | antimicrobial humoral response | 4.11513E-05 | 6 |
| GO:0034341 | response to interferon-gamma | 7.77099E-05 | 6 |
| GO:0001906 | cell killing | 0.00034419 | 6 |
| GO:0071248 | cellular response to metal ion | 0.000421251 | 6 |
| GO:0071241 | cellular response to inorganic substance | 0.000824353 | 6 |
| GO:0002685 | regulation of leukocyte migration | 0.000850692 | 6 |
| GO:0050670 | regulation of lymphocyte proliferation | 0.000959444 | 6 |
| GO:0032944 | regulation of mononuclear cell proliferation | 0.000992784 | 6 |
| GO:0009636 | response to toxic substance | 0.001129054 | 6 |
| GO:0002699 | positive regulation of immune effector process | 0.001373881 | 6 |
| GO:0070663 | regulation of leukocyte proliferation | 0.001558156 | 6 |
| GO:0007249 | I-kappaB kinase/NF-kappaB signaling | 0.002239342 | 6 |
| GO:0046651 | lymphocyte proliferation | 0.002947023 | 6 |
| GO:0031349 | positive regulation of defense response | 0.00301527 | 6 |
| GO:0032943 | mononuclear cell proliferation | 0.00325134 | 6 |
| GO:0070372 | regulation of ERK1 and ERK2 cascade | 0.003382664 | 6 |
| GO:0002440 | production of molecular mediator of immune respo | 0.004354974 | 6 |
| GO:0070371 | ERK1 and ERK2 cascade | 0.004705417 | 6 |
| GO:0070661 | leukocyte proliferation | 0.0050408 | 6 |
| GO:0045765 | regulation of angiogenesis | 0.0050408 | 6 |
| GO:1901342 | regulation of vasculature development | 0.00538143 | 6 |

| | | | |
|---|---|---|---|
| GO:0070997 | neuron death | 0.006324294 | 6 |
| GO:0051098 | regulation of binding | 0.00663224 | 6 |
| GO:0002697 | regulation of immune effector process | 0.007043006 | 6 |
| GO:0042742 | defense response to bacterium | 0.007274073 | 6 |
| GO:1903706 | regulation of hemopoiesis | 0.00905709 | 6 |
| GO:0050727 | regulation of inflammatory response | 0.00905709 | 6 |
| GO:0030098 | lymphocyte differentiation | 0.009354403 | 6 |
| GO:0016032 | viral process | 0.009518612 | 6 |
| GO:0052548 | regulation of endopeptidase activity | 0.010001548 | 6 |
| GO:0002683 | negative regulation of immune system process | 0.012124271 | 6 |
| GO:1903829 | positive regulation of protein localization | 0.012239846 | 6 |
| GO:0052547 | regulation of peptidase activity | 0.012801284 | 6 |
| GO:1903131 | mononuclear cell differentiation | 0.013739846 | 6 |
| GO:0043410 | positive regulation of MAPK cascade | 0.016397865 | 6 |
| hsa04612 | Antigen processing and presentation | 6.00247E-05 | 6 |
| hsa04657 | IL-17 signaling pathway | 0.000120688 | 6 |
| hsa05164 | Influenza A | 0.001389441 | 6 |
| HALLMARK_TNFA_SIGNALING_VIA_NFKB | HALLMARK_TNFA_SIGNALING_VIA_NFKB | 0.019774263 | 6 |
| GO:0005125 | cytokine activity | 0.006371845 | 5 |
| GO:0019886 | antigen processing and presentation of exogenous | 1.73747E-06 | 5 |
| GO:0002495 | antigen processing and presentation of peptide ant | 2.91748E-06 | 5 |
| GO:0002504 | antigen processing and presentation of peptide or | 3.69131E-06 | 5 |
| GO:0071276 | cellular response to cadmium ion | 4.00922E-06 | 5 |
| GO:0010043 | response to zinc ion | 1.46189E-05 | 5 |
| GO:0046686 | response to cadmium ion | 2.63443E-05 | 5 |
| GO:0002688 | regulation of leukocyte chemotaxis | 0.000606944 | 5 |
| GO:0050729 | positive regulation of inflammatory response | 0.001203346 | 5 |
| GO:0098754 | detoxification | 0.001203346 | 5 |
| GO:0051100 | negative regulation of binding | 0.001556381 | 5 |
| GO:0043123 | positive regulation of I-kappaB kinase/NF-kappaB s | 0.002881123 | 5 |
| GO:0070374 | positive regulation of ERK1 and ERK2 cascade | 0.0050408 | 5 |
| GO:0050920 | regulation of chemotaxis | 0.005634122 | 5 |
| GO:0002703 | regulation of leukocyte mediated immunity | 0.006706916 | 5 |
| GO:0045926 | negative regulation of growth | 0.007421847 | 5 |
| GO:0043122 | regulation of I-kappaB kinase/NF-kappaB signaling | 0.007767667 | 5 |
| GO:0002366 | leukocyte activation involved in immune response | 0.011576592 | 5 |
| GO:0030217 | T cell differentiation | 0.011866337 | 5 |
| GO:0042886 | amide transport | 0.011866337 | 5 |
| GO:0002263 | cell activation involved in immune response | 0.011964264 | 5 |
| GO:0051047 | positive regulation of secretion | 0.012586917 | 5 |
| GO:1902105 | regulation of leukocyte differentiation | 0.013739846 | 5 |
| GO:0019058 | viral life cycle | 0.014036752 | 5 |
| GO:1901214 | regulation of neuron death | 0.014896702 | 5 |
| GO:0016042 | lipid catabolic process | 0.016492057 | 5 |
| GO:0032496 | response to lipopolysaccharide | 0.016912475 | 5 |
| GO:0002237 | response to molecule of bacterial origin | 0.020180986 | 5 |
| GO:0042176 | regulation of protein catabolic process | 0.02023529 | 5 |
| GO:0002449 | lymphocyte mediated immunity | 0.021245148 | 5 |
| GO:0045862 | positive regulation of proteolysis | 0.022171605 | 5 |
| GO:2001233 | regulation of apoptotic signaling pathway | 0.022478519 | 5 |
| GO:0002460 | adaptive immune response based on somatic recon | 0.022633075 | 5 |
| GO:0009615 | response to virus | 0.029142579 | 5 |
| GO:0051090 | regulation of DNA-binding transcription factor activ | 0.041550611 | 5 |
| GO:0031667 | response to nutrient levels | 0.046807059 | 5 |
| hsa05140 | Leishmaniasis | 0.000618226 | 5 |
| hsa04620 | Toll-like receptor signaling pathway | 0.001389441 | 5 |
| hsa04142 | Lysosome | 0.003261528 | 5 |
| hsa04145 | Phagosome | 0.005223198 | 5 |
| hsa05169 | Epstein-Barr virus infection | 0.012557456 | 5 |
| hsa05132 | Salmonella infection | 0.024389527 | 5 |
| HALLMARK_ALLOGRAFT_REJECTION | HALLMARK_ALLOGRAFT_REJECTION | 0.036508959 | 5 |
| HALLMARK_INFLAMMATORY_RESPONSE | HALLMARK_INFLAMMATORY_RESPONSE | 0.036508959 | 5 |
| HALLMARK_KRAS_SIGNALING_UP | HALLMARK_KRAS_SIGNALING_UP | 0.036508959 | 5 |
| GO:0140375 | immune receptor activity | 0.007892213 | 4 |
| GO:0031406 | carboxylic acid binding | 0.011138029 | 4 |
| GO:0005126 | cytokine receptor binding | 0.034097931 | 4 |
| GO:0010273 | detoxification of copper ion | 4.91597E-06 | 4 |
| GO:1990169 | stress response to copper ion | 4.91597E-06 | 4 |

| | | | |
|---|---|---|---|
| GO:0061687 | detoxification of inorganic compound | 8.67476E-06 | 4 |
| GO:0097501 | stress response to metal ion | 1.05284E-05 | 4 |
| GO:0071294 | cellular response to zinc ion | 2.38249E-05 | 4 |
| GO:0051238 | sequestering of metal ion | 3.03567E-05 | 4 |
| GO:0071280 | cellular response to copper ion | 3.45222E-05 | 4 |
| GO:0046688 | response to copper ion | 0.000164869 | 4 |
| GO:0002548 | monocyte chemotaxis | 0.000959444 | 4 |
| GO:0061844 | antimicrobial humoral immune response mediated | 0.0013945 | 4 |
| GO:0002690 | positive regulation of leukocyte chemotaxis | 0.002668875 | 4 |
| GO:0045445 | myoblast differentiation | 0.004206825 | 4 |
| GO:0002718 | regulation of cytokine production involved in immu | 0.004743603 | 4 |
| GO:0050830 | defense response to Gram-positive bacterium | 0.004743603 | 4 |
| GO:0071346 | cellular response to interferon-gamma | 0.004743603 | 4 |
| GO:0002367 | cytokine production involved in immune response | 0.005003633 | 4 |
| GO:0006022 | aminoglycan metabolic process | 0.006803044 | 4 |
| GO:0002705 | positive regulation of leukocyte mediated immunit | 0.007654031 | 4 |
| GO:0070555 | response to interleukin-1 | 0.007654031 | 4 |
| GO:0050921 | positive regulation of chemotaxis | 0.007998516 | 4 |
| GO:0050671 | positive regulation of lymphocyte proliferation | 0.008352483 | 4 |
| GO:0002687 | positive regulation of leukocyte migration | 0.008583214 | 4 |
| GO:0032946 | positive regulation of mononuclear cell proliferatic | 0.008644037 | 4 |
| GO:0046718 | viral entry into host cell | 0.009013499 | 4 |
| GO:0044409 | entry into host | 0.009581836 | 4 |
| GO:0070665 | positive regulation of leukocyte proliferation | 0.011283199 | 4 |
| GO:1901136 | carbohydrate derivative catabolic process | 0.012586917 | 4 |
| GO:0048771 | tissue remodeling | 0.013115943 | 4 |
| GO:0002791 | regulation of peptide secretion | 0.013435767 | 4 |
| GO:0042129 | regulation of T cell proliferation | 0.013435767 | 4 |
| GO:0044000 | movement in host | 0.013626936 | 4 |
| GO:0090087 | regulation of peptide transport | 0.013655136 | 4 |
| GO:1902107 | positive regulation of leukocyte differentiation | 0.013655136 | 4 |
| GO:1903708 | positive regulation of hemopoiesis | 0.013655136 | 4 |
| GO:0002700 | regulation of production of molecular mediator of i | 0.014036752 | 4 |
| GO:0002695 | negative regulation of leukocyte activation | 0.016407823 | 4 |
| GO:0051701 | biological process involved in interaction with host | 0.01749395 | 4 |
| GO:0071674 | mononuclear cell migration | 0.01816444 | 4 |
| GO:0009612 | response to mechanical stimulus | 0.019022176 | 4 |
| GO:0042098 | T cell proliferation | 0.019254607 | 4 |
| GO:0045637 | regulation of myeloid cell differentiation | 0.01972418 | 4 |
| GO:1901215 | negative regulation of neuron death | 0.020582252 | 4 |
| GO:0050866 | negative regulation of cell activation | 0.021245148 | 4 |
| GO:0002790 | peptide secretion | 0.023706886 | 4 |
| GO:0071356 | cellular response to tumor necrosis factor | 0.02445881 | 4 |
| GO:2000116 | regulation of cysteine-type endopeptidase activity | 0.024701569 | 4 |
| GO:0015833 | peptide transport | 0.02993791 | 4 |
| GO:0034612 | response to tumor necrosis factor | 0.02993791 | 4 |
| GO:0051402 | neuron apoptotic process | 0.030234767 | 4 |
| GO:0001894 | tissue homeostasis | 0.037914857 | 4 |
| GO:1903532 | positive regulation of secretion by cell | 0.039230029 | 4 |
| GO:0043270 | positive regulation of ion transport | 0.043078119 | 4 |
| GO:0051222 | positive regulation of protein transport | 0.045546814 | 4 |
| GO:0097193 | intrinsic apoptotic signaling pathway | 0.046807059 | 4 |
| GO:0051607 | defense response to virus | 0.048275632 | 4 |
| GO:0140546 | defense response to symbiont | 0.048655061 | 4 |
| hsa05332 | Graft-versus-host disease | 0.000742385 | 4 |
| hsa04940 | Type I diabetes mellitus | 0.000742385 | 4 |
| hsa05416 | Viral myocarditis | 0.001855898 | 4 |
| hsa04623 | Cytosolic DNA-sensing pathway | 0.003614136 | 4 |
| hsa05323 | Rheumatoid arthritis | 0.00645419 | 4 |
| hsa04659 | Th17 cell differentiation | 0.009898835 | 4 |
| hsa04380 | Osteoclast differentiation | 0.014825258 | 4 |
| hsa05418 | Fluid shear stress and atherosclerosis | 0.018189252 | 4 |
| hsa04217 | Necroptosis | 0.026997843 | 4 |
| hsa05152 | Tuberculosis | 0.038555346 | 4 |
| hsa04062 | Chemokine signaling pathway | 0.041785626 | 4 |
| hsa05130 | Pathogenic Escherichia coli infection | 0.04222212 | 4 |
| HALLMARK_INTERFERON_ALPHA_RESPONSE | HALLMARK_INTERFERON_ALPHA_RESPONSE | 0.028013935 | 4 |
| GO:0023026 | MHC class II protein complex binding | 0.002847258 | 3 |

| | | | |
|---|---|---|---|
| GO:0023023 | MHC protein complex binding | 0.004454046 | 3 |
| GO:0042605 | peptide antigen binding | 0.004454046 | 3 |
| GO:0008009 | chemokine activity | 0.005907773 | 3 |
| GO:0042379 | chemokine receptor binding | 0.009609177 | 3 |
| GO:0033293 | monocarboxylic acid binding | 0.011856628 | 3 |
| GO:0048306 | calcium-dependent protein binding | 0.011856628 | 3 |
| GO:0001540 | amyloid-beta binding | 0.012942727 | 3 |
| GO:0043177 | organic acid binding | 0.034097931 | 3 |
| GO:0005178 | integrin binding | 0.043206328 | 3 |
| GO:0070486 | leukocyte aggregation | 0.000196649 | 3 |
| GO:0017014 | protein nitrosylation | 0.000396466 | 3 |
| GO:0018119 | peptidyl-cysteine S-nitrosylation | 0.000396466 | 3 |
| GO:0019883 | antigen processing and presentation of endogenou | 0.001068331 | 3 |
| GO:0014002 | astrocyte development | 0.003743441 | 3 |
| GO:0031640 | killing of cells of another organism | 0.00463161 | 3 |
| GO:0046596 | regulation of viral entry into host cell | 0.00463161 | 3 |
| GO:0018198 | peptidyl-cysteine modification | 0.005380242 | 3 |
| GO:0052372 | modulation by symbiont of entry into host | 0.006201672 | 3 |
| GO:0050832 | defense response to fungus | 0.006947606 | 3 |
| GO:0001912 | positive regulation of leukocyte mediated cytotoxi | 0.007654031 | 3 |
| GO:0043903 | regulation of biological process involved in symbiot | 0.007654031 | 3 |
| GO:0002711 | positive regulation of T cell mediated immunity | 0.007767667 | 3 |
| GO:0032757 | positive regulation of interleukin-8 production | 0.008583214 | 3 |
| GO:0048247 | lymphocyte chemotaxis | 0.00905709 | 3 |
| GO:0006879 | cellular iron ion homeostasis | 0.00905709 | 3 |
| GO:0031343 | positive regulation of cell killing | 0.00905709 | 3 |
| GO:0009620 | response to fungus | 0.009202748 | 3 |
| GO:0031638 | zymogen activation | 0.009202748 | 3 |
| GO:0032722 | positive regulation of chemokine production | 0.010001548 | 3 |
| GO:0071677 | positive regulation of mononuclear cell migration | 0.011147995 | 3 |
| GO:0045661 | regulation of myoblast differentiation | 0.011808038 | 3 |
| GO:0032729 | positive regulation of interferon-gamma productior | 0.012239846 | 3 |
| GO:0002720 | positive regulation of cytokine production involvec | 0.012672359 | 3 |
| GO:0048708 | astrocyte differentiation | 0.013435767 | 3 |
| GO:0034103 | regulation of tissue remodeling | 0.014036752 | 3 |
| GO:0001910 | regulation of leukocyte mediated cytotoxicity | 0.014896702 | 3 |
| GO:0032370 | positive regulation of lipid transport | 0.015161977 | 3 |
| GO:0055072 | iron ion homeostasis | 0.015161977 | 3 |
| GO:0002709 | regulation of T cell mediated immunity | 0.015574943 | 3 |
| GO:0070098 | chemokine-mediated signaling pathway | 0.015905097 | 3 |
| GO:0045185 | maintenance of protein location | 0.016912475 | 3 |
| GO:0032091 | negative regulation of protein binding | 0.01816444 | 3 |
| GO:0032642 | regulation of chemokine production | 0.01874185 | 3 |
| GO:1990868 | response to chemokine | 0.01874185 | 3 |
| GO:1990869 | cellular response to chemokine | 0.01874185 | 3 |
| GO:0032602 | chemokine production | 0.019022176 | 3 |
| GO:0032755 | positive regulation of interleukin-6 production | 0.019022176 | 3 |
| GO:0031341 | regulation of cell killing | 0.020180986 | 3 |
| GO:0032677 | regulation of interleukin-8 production | 0.02054734 | 3 |
| GO:0032637 | interleukin-8 production | 0.020916287 | 3 |
| GO:0042102 | positive regulation of T cell proliferation | 0.021661887 | 3 |
| GO:0071347 | cellular response to interleukin-1 | 0.023892909 | 3 |
| GO:0002456 | T cell mediated immunity | 0.02445881 | 3 |
| GO:1905954 | positive regulation of lipid localization | 0.02445881 | 3 |
| GO:0002824 | positive regulation of adaptive immune response b | 0.024853507 | 3 |
| GO:0051702 | biological process involved in interaction with sym | 0.025987583 | 3 |
| GO:0021782 | glial cell development | 0.026391735 | 3 |
| GO:0045582 | positive regulation of T cell differentiation | 0.026391735 | 3 |
| GO:0002708 | positive regulation of lymphocyte mediated immur | 0.027091267 | 3 |
| GO:0032609 | interferon-gamma production | 0.027091267 | 3 |
| GO:0032649 | regulation of interferon-gamma production | 0.027091267 | 3 |
| GO:0002821 | positive regulation of adaptive immune response | 0.027501631 | 3 |
| GO:0002761 | regulation of myeloid leukocyte differentiation | 0.028650175 | 3 |
| GO:0030203 | glycosaminoglycan metabolic process | 0.028865205 | 3 |
| GO:0071675 | regulation of mononuclear cell migration | 0.028865205 | 3 |
| GO:0072676 | lymphocyte migration | 0.02928539 | 3 |
| GO:0045621 | positive regulation of lymphocyte differentiation | 0.032487185 | 3 |
| GO:0002702 | positive regulation of production of molecular med | 0.034345066 | 3 |

| | | | |
|---|---|---|---|
| GO:0001909 | leukocyte mediated cytotoxicity | 0.034808949 | 3 |
| GO:0038061 | NIK/NF-kappaB signaling | 0.039922939 | 3 |
| GO:1903900 | regulation of viral life cycle | 0.039922939 | 3 |
| GO:2001056 | positive regulation of cysteine-type endopeptidase | 0.040479424 | 3 |
| GO:0050768 | negative regulation of neurogenesis | 0.042295396 | 3 |
| GO:0051961 | negative regulation of nervous system developmer | 0.045546814 | 3 |
| GO:0007584 | response to nutrient | 0.046807059 | 3 |
| GO:0051092 | positive regulation of NF-kappaB transcription fact | 0.046807059 | 3 |
| GO:0032368 | regulation of lipid transport | 0.047192913 | 3 |
| GO:0090316 | positive regulation of intracellular protein transpo | 0.048193003 | 3 |
| hsa05330 | Allograft rejection | 0.006004007 | 3 |
| hsa04216 | Ferroptosis | 0.006517025 | 3 |
| hsa05320 | Autoimmune thyroid disease | 0.012195173 | 3 |
| hsa05134 | Legionellosis | 0.012797365 | 3 |
| hsa05321 | Inflammatory bowel disease | 0.017765563 | 3 |
| hsa04658 | Th1 and Th2 cell differentiation | 0.038555346 | 3 |
| hsa04640 | Hematopoietic cell lineage | 0.041785626 | 3 |
| hsa04061 | Viral protein interaction with cytokine and cytokir | 0.041785626 | 3 |
| hsa04064 | NF-kappa B signaling pathway | 0.04222212 | 3 |
| hsa04625 | C-type lectin receptor signaling pathway | 0.04222212 | 3 |
| hsa05145 | Toxoplasmosis | 0.048653566 | 3 |
| GO:0032395 | MHC class II receptor activity | 0.005907773 | 2 |
| GO:0042608 | T cell receptor binding | 0.005907773 | 2 |
| GO:0050786 | RAGE receptor binding | 0.005907773 | 2 |
| GO:0008199 | ferric iron binding | 0.006371845 | 2 |
| GO:0035325 | Toll-like receptor binding | 0.006476504 | 2 |
| GO:0050998 | nitric-oxide synthase binding | 0.007053286 | 2 |
| GO:0036041 | long-chain fatty acid binding | 0.008200933 | 2 |
| GO:0008198 | ferrous iron binding | 0.017518832 | 2 |
| GO:0001968 | fibronectin binding | 0.022091502 | 2 |
| GO:0042056 | chemoattractant activity | 0.02739528 | 2 |
| GO:0043394 | proteoglycan binding | 0.02739528 | 2 |
| GO:0031492 | nucleosomal DNA binding | 0.032441561 | 2 |
| GO:0048156 | tau protein binding | 0.034097931 | 2 |
| GO:0048020 | CCR chemokine receptor binding | 0.038698937 | 2 |
| GO:0005504 | fatty acid binding | 0.040646782 | 2 |
| GO:0050661 | NADP binding | 0.044525446 | 2 |
| GO:0050840 | extracellular matrix binding | 0.048094332 | 2 |
| GO:0031392 | regulation of prostaglandin biosynthetic process | 0.007767667 | 2 |
| GO:0046598 | positive regulation of viral entry into host cell | 0.008583214 | 2 |
| GO:0075294 | positive regulation by symbiont of entry into host | 0.008583214 | 2 |
| GO:0002468 | dendritic cell antigen processing and presentation | 0.00905709 | 2 |
| GO:0097396 | response to interleukin-17 | 0.00905709 | 2 |
| GO:0097398 | cellular response to interleukin-17 | 0.00905709 | 2 |
| GO:2001279 | regulation of unsaturated fatty acid biosynthetic p | 0.00905709 | 2 |
| GO:0002399 | MHC class II protein complex assembly | 0.009809658 | 2 |
| GO:0002503 | peptide antigen assembly with MHC class II proteir | 0.009809658 | 2 |
| GO:0002483 | antigen processing and presentation of endogenou | 0.012545853 | 2 |
| GO:0002523 | leukocyte migration involved in inflammatory resp | 0.012545853 | 2 |
| GO:0002544 | chronic inflammatory response | 0.012545853 | 2 |
| GO:0002396 | MHC protein complex assembly | 0.013225276 | 2 |
| GO:0002501 | peptide antigen assembly with MHC protein compl | 0.013225276 | 2 |
| GO:0035455 | response to interferon-alpha | 0.013225276 | 2 |
| GO:0090026 | positive regulation of monocyte chemotaxis | 0.013739846 | 2 |
| GO:0045723 | positive regulation of fatty acid biosynthetic proce | 0.015673657 | 2 |
| GO:1903902 | positive regulation of viral life cycle | 0.016608931 | 2 |
| GO:0045662 | negative regulation of myoblast differentiation | 0.017641922 | 2 |
| GO:0002726 | positive regulation of T cell cytokine production | 0.018685071 | 2 |
| GO:0050995 | negative regulation of lipid catabolic process | 0.018685071 | 2 |
| GO:1901623 | regulation of lymphocyte chemotaxis | 0.019344706 | 2 |
| GO:0090025 | regulation of monocyte chemotaxis | 0.021245148 | 2 |
| GO:0060142 | regulation of syncytium formation by plasma meml | 0.02240652 | 2 |
| GO:0001916 | positive regulation of T cell mediated cytotoxicity | 0.023581695 | 2 |
| GO:0001516 | prostaglandin biosynthetic process | 0.02445881 | 2 |
| GO:0006516 | glycoprotein catabolic process | 0.02445881 | 2 |
| GO:0046457 | prostanoid biosynthetic process | 0.02445881 | 2 |
| GO:0034383 | low-density lipoprotein particle clearance | 0.02647927 | 2 |
| GO:0090022 | regulation of neutrophil chemotaxis | 0.02647927 | 2 |

| | | | |
|---|---|---|---|
| GO:0030212 | hyaluronan metabolic process | 0.027501631 | 2 |
| GO:0006026 | aminoglycan catabolic process | 0.028650175 | 2 |
| GO:0033280 | response to vitamin D | 0.028650175 | 2 |
| GO:0002369 | T cell cytokine production | 0.030590081 | 2 |
| GO:0002724 | regulation of T cell cytokine production | 0.030590081 | 2 |
| GO:0043368 | positive T cell selection | 0.030590081 | 2 |
| GO:0045923 | positive regulation of fatty acid metabolic process | 0.033560405 | 2 |
| GO:0001914 | regulation of T cell mediated cytotoxicity | 0.034808949 | 2 |
| GO:2000403 | positive regulation of lymphocyte migration | 0.036346672 | 2 |
| GO:0021762 | substantia nigra development | 0.039342473 | 2 |
| GO:0030574 | collagen catabolic process | 0.040479424 | 2 |
| GO:1900271 | regulation of long-term synaptic potentiation | 0.04193089 | 2 |
| GO:0140353 | lipid export from cell | 0.043078119 | 2 |
| GO:1902622 | regulation of neutrophil migration | 0.043078119 | 2 |
| GO:0035094 | response to nicotine | 0.046188401 | 2 |
| GO:0006692 | prostanoid metabolic process | 0.046807059 | 2 |
| GO:0006693 | prostaglandin metabolic process | 0.046807059 | 2 |
| GO:0042304 | regulation of fatty acid biosynthetic process | 0.046807059 | 2 |
| GO:0061082 | myeloid leukocyte cytokine production | 0.046807059 | 2 |
| GO:0001913 | T cell mediated cytotoxicity | 0.047803195 | 2 |
| GO:0019083 | viral transcription | 0.047803195 | 2 |
| GO:0045058 | T cell selection | 0.047803195 | 2 |
| GO:0006636 | unsaturated fatty acid biosynthetic process | 0.048774882 | 2 |
| GO:0043392 | negative regulation of DNA binding | 0.048774882 | 2 |
| hsa05310 | Asthma | 0.040049703 | 2 |

| ID | Description | p.adjust | Count |
|---|---|---|---|
| GO:0050900 | leukocyte migration | 0.003188285 | 7 |
| HALLMARK_KRAS_SIGNALING_UP | HALLMARK_KRAS_SIGNALING_UP | 0.000810609 | 7 |
| GO:0033218 | amide binding | 0.006178415 | 6 |
| GO:0030593 | neutrophil chemotaxis | 0.000245915 | 6 |
| GO:0071621 | granulocyte chemotaxis | 0.000274024 | 6 |
| GO:1990266 | neutrophil migration | 0.000274024 | 6 |
| GO:0034341 | response to interferon-gamma | 0.000317356 | 6 |
| GO:0097530 | granulocyte migration | 0.000513668 | 6 |
| GO:0071674 | mononuclear cell migration | 0.001567978 | 6 |
| GO:0030595 | leukocyte chemotaxis | 0.002109873 | 6 |
| GO:0097529 | myeloid leukocyte migration | 0.002109873 | 6 |
| GO:0044403 | biological process involved in symbiotic interaction | 0.004770759 | 6 |
| GO:0019058 | viral life cycle | 0.00541231 | 6 |
| GO:0060326 | cell chemotaxis | 0.00541231 | 6 |
| GO:0016032 | viral process | 0.01758809 | 6 |
| GO:0002683 | negative regulation of immune system process | 0.022029619 | 6 |
| GO:0032103 | positive regulation of response to external stimulu | 0.025449944 | 6 |
| GO:0050673 | epithelial cell proliferation | 0.028444913 | 6 |
| GO:0019221 | cytokine-mediated signaling pathway | 0.030393979 | 6 |
| GO:0043410 | positive regulation of MAPK cascade | 0.030393979 | 6 |
| GO:0038024 | cargo receptor activity | 0.000237049 | 5 |
| GO:0001664 | G protein-coupled receptor binding | 0.008397572 | 5 |
| GO:0072676 | lymphocyte migration | 0.001567978 | 5 |
| GO:0046718 | viral entry into host cell | 0.002431991 | 5 |
| GO:0044409 | entry into host | 0.002831432 | 5 |
| GO:0044000 | movement in host | 0.004547737 | 5 |
| GO:0051701 | biological process involved in interaction with host | 0.005641801 | 5 |
| GO:0034612 | response to tumor necrosis factor | 0.012799818 | 5 |
| GO:0043547 | positive regulation of GTPase activity | 0.019481823 | 5 |
| GO:0043087 | regulation of GTPase activity | 0.046128028 | 5 |
| hsa04610 | Complement and coagulation cascades | 0.000734317 | 5 |
| GO:0005044 | scavenger receptor activity | 0.000237049 | 4 |
| GO:0048020 | CCR chemokine receptor binding | 0.000237049 | 4 |
| GO:0008009 | chemokine activity | 0.000237049 | 4 |
| GO:0042379 | chemokine receptor binding | 0.000579655 | 4 |
| GO:0005125 | cytokine activity | 0.026602407 | 4 |
| GO:0005126 | cytokine receptor binding | 0.039778262 | 4 |
| GO:0048247 | lymphocyte chemotaxis | 0.002080812 | 4 |
| GO:0002548 | monocyte chemotaxis | 0.002109873 | 4 |
| GO:0070098 | chemokine-mediated signaling pathway | 0.004169049 | 4 |
| GO:1990868 | response to chemokine | 0.004752635 | 4 |
| GO:1990869 | cellular response to chemokine | 0.004752635 | 4 |
| GO:1902106 | negative regulation of leukocyte differentiation | 0.005641801 | 4 |
| GO:0071347 | cellular response to interleukin-1 | 0.006046744 | 4 |
| GO:1903707 | negative regulation of hemopoiesis | 0.006469565 | 4 |
| GO:0071346 | cellular response to interferon-gamma | 0.007140004 | 4 |
| GO:0070555 | response to interleukin-1 | 0.012799818 | 4 |
| GO:0007009 | plasma membrane organization | 0.021097043 | 4 |
| GO:0070374 | positive regulation of ERK1 and ERK2 cascade | 0.045385016 | 4 |
| hsa05150 | Staphylococcus aureus infection | 0.009130334 | 4 |
| hsa04061 | Viral protein interaction with cytokine and cytokir | 0.009130334 | 4 |
| hsa04145 | Phagosome | 0.033060746 | 4 |
| GO:0045028 | G protein-coupled purinergic nucleotide receptor a | 0.000237049 | 3 |
| GO:0030169 | low-density lipoprotein particle binding | 0.000281664 | 3 |
| GO:0001614 | purinergic nucleotide receptor activity | 0.000410228 | 3 |
| GO:0016502 | nucleotide receptor activity | 0.000410228 | 3 |
| GO:0038187 | pattern recognition receptor activity | 0.000714047 | 3 |
| GO:0071813 | lipoprotein particle binding | 0.000821514 | 3 |
| GO:0071814 | protein-lipid complex binding | 0.000821514 | 3 |
| GO:0035589 | G protein-coupled purinergic nucleotide receptor si | 0.000877909 | 3 |
| GO:0048245 | eosinophil chemotaxis | 0.001567978 | 3 |
| GO:0072677 | eosinophil migration | 0.002109873 | 3 |
| GO:0035590 | purinergic nucleotide receptor signaling pathway | 0.004401308 | 3 |
| GO:0045638 | negative regulation of myeloid cell differentiation | 0.036583519 | 3 |
| GO:0070664 | negative regulation of leukocyte proliferation | 0.036583519 | 3 |
| GO:0008035 | high-density lipoprotein particle binding | 0.005213254 | 2 |
| GO:0031994 | insulin-like growth factor I binding | 0.005711741 | 2 |

| | | | |
|---|---|---|---|
| GO:0005520 | insulin-like growth factor binding | 0.010212061 | 2 |
| GO:0001848 | complement binding | 0.018123446 | 2 |
| GO:0001530 | lipopolysaccharide binding | 0.03283437 | 2 |
| GO:0097278 | complement-dependent cytotoxicity | 0.010837105 | 2 |
| GO:0034375 | high-density lipoprotein particle remodeling | 0.020199833 | 2 |
| GO:0035455 | response to interferon-alpha | 0.027905264 | 2 |
| GO:0035461 | vitamin transmembrane transport | 0.027905264 | 2 |
| GO:0030449 | regulation of complement activation | 0.030393979 | 2 |
| GO:0046597 | negative regulation of viral entry into host cell | 0.030393979 | 2 |
| GO:0043567 | regulation of insulin-like growth factor receptor sig | 0.036583519 | 2 |
| GO:1903901 | negative regulation of viral life cycle | 0.038770744 | 2 |
| GO:0034368 | protein-lipid complex remodeling | 0.047746121 | 2 |
| GO:0034369 | plasma lipoprotein particle remodeling | 0.047746121 | 2 |

| ID | Description | p.adjust | Count |
|---|---|---|---|
| GO:0002449 | lymphocyte mediated immunity | 1.91851E-05 | 9 |
| GO:0002443 | leukocyte mediated immunity | 0.000113028 | 9 |
| GO:0002460 | adaptive immune response based on somatic recon | 0.000207702 | 8 |
| GO:0050863 | regulation of T cell activation | 0.000207702 | 8 |
| GO:0002253 | activation of immune response | 0.000290738 | 8 |
| hsa05150 | Staphylococcus aureus infection | 3.13691E-08 | 8 |
| GO:0002440 | production of molecular mediator of immune respo | 0.000784117 | 7 |
| GO:0007159 | leukocyte cell-cell adhesion | 0.002716868 | 7 |
| GO:0002696 | positive regulation of leukocyte activation | 0.004207795 | 7 |
| GO:0050867 | positive regulation of cell activation | 0.00478509 | 7 |
| hsa04640 | Hematopoietic cell lineage | 7.43899E-07 | 7 |
| HALLMARK_ALLOGRAFT_REJECTION | HALLMARK_ALLOGRAFT_REJECTION | 0.002562359 | 7 |
| GO:0019886 | antigen processing and presentation of exogenous | 8.54418E-08 | 6 |
| GO:0002495 | antigen processing and presentation of peptide ant | 9.24138E-08 | 6 |
| GO:0002504 | antigen processing and presentation of peptide or | 9.24138E-08 | 6 |
| GO:0002478 | antigen processing and presentation of exogenous | 1.35596E-07 | 6 |
| GO:0019884 | antigen processing and presentation of exogenous | 3.8879E-07 | 6 |
| GO:0048002 | antigen processing and presentation of peptide ant | 1.69039E-06 | 6 |
| GO:0019882 | antigen processing and presentation | 1.99141E-05 | 6 |
| GO:0016064 | immunoglobulin mediated immune response | 0.000773233 | 6 |
| GO:0019724 | B cell mediated immunity | 0.000784117 | 6 |
| GO:0002429 | immune response-activating cell surface receptor s | 0.004207795 | 6 |
| GO:0002757 | immune response-activating signal transduction | 0.004207795 | 6 |
| GO:0002768 | immune response-regulating cell surface receptor s | 0.005693205 | 6 |
| GO:1903037 | regulation of leukocyte cell-cell adhesion | 0.008319119 | 6 |
| GO:0050900 | leukocyte migration | 0.009326727 | 6 |
| GO:0051251 | positive regulation of lymphocyte activation | 0.009355651 | 6 |
| GO:1903706 | regulation of hemopoiesis | 0.010360263 | 6 |
| GO:0032103 | positive regulation of response to external stimulu | 0.016894051 | 6 |
| GO:0022407 | regulation of cell-cell adhesion | 0.020388757 | 6 |
| hsa04612 | Antigen processing and presentation | 3.62468E-06 | 6 |
| hsa04145 | Phagosome | 9.02721E-05 | 6 |
| hsa04514 | Cell adhesion molecules | 9.02721E-05 | 6 |
| hsa05152 | Tuberculosis | 0.000134678 | 6 |
| hsa05166 | Human T-cell leukemia virus 1 infection | 0.000323073 | 6 |
| HALLMARK_COMPLEMENT | HALLMARK_COMPLEMENT | 0.006918472 | 6 |
| HALLMARK_INFLAMMATORY_RESPONSE | HALLMARK_INFLAMMATORY_RESPONSE | 0.006918472 | 6 |
| GO:0042277 | peptide binding | 0.032304765 | 5 |
| GO:0071674 | mononuclear cell migration | 0.005064488 | 5 |
| GO:0002377 | immunoglobulin production | 0.006085396 | 5 |
| GO:0030595 | leukocyte chemotaxis | 0.008319119 | 5 |
| GO:0097529 | myeloid leukocyte migration | 0.008319119 | 5 |
| GO:0050870 | positive regulation of T cell activation | 0.009326727 | 5 |
| GO:0050851 | antigen receptor-mediated signaling pathway | 0.009326727 | 5 |
| GO:1903039 | positive regulation of leukocyte cell-cell adhesion | 0.010989996 | 5 |
| GO:1902105 | regulation of leukocyte differentiation | 0.019223871 | 5 |
| GO:0060326 | cell chemotaxis | 0.019319132 | 5 |
| GO:0006959 | humoral immune response | 0.019319132 | 5 |
| GO:0022409 | positive regulation of cell-cell adhesion | 0.019507897 | 5 |
| GO:0010038 | response to metal ion | 0.028835319 | 5 |
| GO:0002697 | regulation of immune effector process | 0.035514078 | 5 |
| hsa05164 | Influenza A | 0.000908388 | 5 |
| hsa05169 | Epstein-Barr virus infection | 0.001548524 | 5 |
| HALLMARK_APOPTOSIS | HALLMARK_APOPTOSIS | 0.013325946 | 5 |
| HALLMARK_KRAS_SIGNALING_UP | HALLMARK_KRAS_SIGNALING_UP | 0.027460588 | 5 |
| GO:0023026 | MHC class II protein complex binding | 9.54298E-05 | 4 |
| GO:0023023 | MHC protein complex binding | 0.000157483 | 4 |
| GO:0140375 | immune receptor activity | 0.016658016 | 4 |
| GO:0002399 | MHC class II protein complex assembly | 6.9424E-06 | 4 |
| GO:0002503 | peptide antigen assembly with MHC class II protei | 6.9424E-06 | 4 |
| GO:0002396 | MHC protein complex assembly | 1.46802E-05 | 4 |
| GO:0002501 | peptide antigen assembly with MHC protein compl | 1.46802E-05 | 4 |
| GO:0002548 | monocyte chemotaxis | 0.001344461 | 4 |
| GO:0002381 | immunoglobulin production involved in immunoglo | 0.001508261 | 4 |
| GO:0030593 | neutrophil chemotaxis | 0.00478509 | 4 |
| GO:0071347 | cellular response to interleukin-1 | 0.005323595 | 4 |
| GO:0071621 | granulocyte chemotaxis | 0.008319119 | 4 |

| | | | |
|---|---|---|---|
| GO:1990266 | neutrophil migration | 0.008319119 | 4 |
| GO:0050852 | T cell receptor signaling pathway | 0.009326727 | 4 |
| GO:0070555 | response to interleukin-1 | 0.009326727 | 4 |
| GO:0050921 | positive regulation of chemotaxis | 0.009492715 | 4 |
| GO:0097530 | granulocyte migration | 0.011855327 | 4 |
| GO:0050920 | regulation of chemotaxis | 0.036492184 | 4 |
| GO:0002274 | myeloid leukocyte activation | 0.039712749 | 4 |
| GO:0034612 | response to tumor necrosis factor | 0.047749316 | 4 |
| hsa05310 | Asthma | 5.31386E-05 | 4 |
| hsa05330 | Allograft rejection | 9.02721E-05 | 4 |
| hsa05332 | Graft-versus-host disease | 9.02721E-05 | 4 |
| hsa04940 | Type I diabetes mellitus | 9.02721E-05 | 4 |
| hsa04672 | Intestinal immune network for IgA production | 0.000134678 | 4 |
| hsa05320 | Autoimmune thyroid disease | 0.000157232 | 4 |
| hsa05416 | Viral myocarditis | 0.000238086 | 4 |
| hsa05321 | Inflammatory bowel disease | 0.00030367 | 4 |
| hsa05140 | Leishmaniasis | 0.000517587 | 4 |
| hsa04658 | Th1 and Th2 cell differentiation | 0.000908604 | 4 |
| hsa05323 | Rheumatoid arthritis | 0.000908604 | 4 |
| hsa04659 | Th17 cell differentiation | 0.001534077 | 4 |
| hsa05145 | Toxoplasmosis | 0.001548524 | 4 |
| hsa05322 | Systemic lupus erythematosus | 0.003284578 | 4 |
| hsa04060 | Cytokine-cytokine receptor interaction | 0.046612498 | 4 |
| HALLMARK_COAGULATION | HALLMARK_COAGULATION | 0.034381909 | 4 |
| GO:0048020 | CCR chemokine receptor binding | 0.010085424 | 3 |
| GO:0008009 | chemokine activity | 0.010085424 | 3 |
| GO:0042379 | chemokine receptor binding | 0.020055098 | 3 |
| GO:0048247 | lymphocyte chemotaxis | 0.010989996 | 3 |
| GO:0002709 | regulation of T cell mediated immunity | 0.023175012 | 3 |
| GO:0070098 | chemokine-mediated signaling pathway | 0.023527851 | 3 |
| GO:1990868 | response to chemokine | 0.028835319 | 3 |
| GO:1990869 | cellular response to chemokine | 0.028835319 | 3 |
| GO:1902106 | negative regulation of leukocyte differentiation | 0.036492184 | 3 |
| GO:0002456 | T cell mediated immunity | 0.039712749 | 3 |
| GO:1903707 | negative regulation of hemopoiesis | 0.040118267 | 3 |
| GO:0002286 | T cell activation involved in immune response | 0.04296709 | 3 |
| GO:0071346 | cellular response to interferon-gamma | 0.04296709 | 3 |
| GO:0072676 | lymphocyte migration | 0.048045373 | 3 |
| hsa04610 | Complement and coagulation cascades | 0.009165581 | 3 |
| hsa04061 | Viral protein interaction with cytokine and cytokir | 0.013501455 | 3 |
| GO:0002674 | negative regulation of acute inflammatory respons | 0.009326727 | 2 |
| GO:2001198 | regulation of dendritic cell differentiation | 0.010029831 | 2 |
| GO:0071236 | cellular response to antibiotic | 0.010989996 | 2 |
| GO:0006957 | complement activation, alternative pathway | 0.016894051 | 2 |

| ID | Description | p.adjust | Count |
|---|---|---|---|
| HALLMARK_INTERFERON_GAMMA_RESPONS | HALLMARK_INTERFERON_GAMMA_RESPONSE | 4.47919E-12 | 15 |
| GO:0060326 | cell chemotaxis | 2.11209E-07 | 10 |
| GO:0050900 | leukocyte migration | 6.51181E-07 | 10 |
| GO:1990266 | neutrophil migration | 8.58535E-09 | 9 |
| GO:0097530 | granulocyte migration | 2.50114E-08 | 9 |
| GO:0030595 | leukocyte chemotaxis | 2.20422E-07 | 9 |
| GO:0097529 | myeloid leukocyte migration | 2.20422E-07 | 9 |
| GO:0019221 | cytokine-mediated signaling pathway | 3.27385E-05 | 9 |
| GO:0034341 | response to interferon-gamma | 1.93742E-07 | 8 |
| GO:0002685 | regulation of leukocyte migration | 1.55289E-06 | 8 |
| GO:0050863 | regulation of T cell activation | 3.88666E-05 | 8 |
| GO:1903037 | regulation of leukocyte cell-cell adhesion | 3.88666E-05 | 8 |
| GO:0007159 | leukocyte cell-cell adhesion | 7.37309E-05 | 8 |
| GO:0022407 | regulation of cell-cell adhesion | 0.000223525 | 8 |
| HALLMARK_INFLAMMATORY_RESPONSE | HALLMARK_INFLAMMATORY_RESPONSE | 0.00023217 | 8 |
| GO:0042379 | chemokine receptor binding | 2.49921E-08 | 7 |
| GO:0005126 | cytokine receptor binding | 3.97634E-05 | 7 |
| GO:0001664 | G protein-coupled receptor binding | 5.06865E-05 | 7 |
| GO:0070098 | chemokine-mediated signaling pathway | 1.93742E-07 | 7 |
| GO:1990868 | response to chemokine | 2.11209E-07 | 7 |
| GO:1990869 | cellular response to chemokine | 2.11209E-07 | 7 |
| GO:0030593 | neutrophil chemotaxis | 2.6709E-07 | 7 |
| GO:0071346 | cellular response to interferon-gamma | 4.45235E-07 | 7 |
| GO:0071621 | granulocyte chemotaxis | 6.80448E-07 | 7 |
| GO:0071674 | mononuclear cell migration | 1.06922E-05 | 7 |
| GO:0050920 | regulation of chemotaxis | 2.02334E-05 | 7 |
| GO:0050870 | positive regulation of T cell activation | 3.30227E-05 | 7 |
| GO:1903039 | positive regulation of leukocyte cell-cell adhesion | 5.55736E-05 | 7 |
| GO:0022409 | positive regulation of cell-cell adhesion | 0.000141379 | 7 |
| GO:0051251 | positive regulation of lymphocyte activation | 0.000469834 | 7 |
| GO:0016032 | viral process | 0.000638215 | 7 |
| GO:0002696 | positive regulation of leukocyte activation | 0.000954321 | 7 |
| GO:0032103 | positive regulation of response to external stimulu | 0.001093151 | 7 |
| GO:0050867 | positive regulation of cell activation | 0.001109839 | 7 |
| GO:0045785 | positive regulation of cell adhesion | 0.001349266 | 7 |
| hsa04145 | Phagosome | 4.64101E-05 | 7 |
| HALLMARK_ALLOGRAFT_REJECTION | HALLMARK_ALLOGRAFT_REJECTION | 0.001080071 | 7 |
| HALLMARK_TNFA_SIGNALING_VIA_NFKB | HALLMARK_TNFA_SIGNALING_VIA_NFKB | 0.001080071 | 7 |
| GO:0008009 | chemokine activity | 6.96101E-08 | 6 |
| GO:0140375 | immune receptor activity | 2.22143E-05 | 6 |
| GO:0005125 | cytokine activity | 0.000186309 | 6 |
| GO:0048018 | receptor ligand activity | 0.006989705 | 6 |
| GO:0030546 | signaling receptor activator activity | 0.006989705 | 6 |
| GO:0019884 | antigen processing and presentation of exogenous | 1.93742E-07 | 6 |
| GO:0048247 | lymphocyte chemotaxis | 4.28783E-07 | 6 |
| GO:0002548 | monocyte chemotaxis | 6.29607E-07 | 6 |
| GO:0019882 | antigen processing and presentation | 5.23325E-06 | 6 |
| GO:0072676 | lymphocyte migration | 1.05241E-05 | 6 |
| GO:0002688 | regulation of leukocyte chemotaxis | 1.10095E-05 | 6 |
| GO:0044403 | biological process involved in symbiotic interaction | 0.000966643 | 6 |
| GO:0019058 | viral life cycle | 0.001109839 | 6 |
| GO:0032496 | response to lipopolysaccharide | 0.001497798 | 6 |
| GO:0002237 | response to molecule of bacterial origin | 0.001885807 | 6 |
| GO:0002449 | lymphocyte mediated immunity | 0.002062096 | 6 |
| GO:0002460 | adaptive immune response based on somatic recon | 0.002216101 | 6 |
| GO:0002443 | leukocyte mediated immunity | 0.005543191 | 6 |
| GO:1903131 | mononuclear cell differentiation | 0.006058929 | 6 |
| hsa04061 | Viral protein interaction with cytokine and cytokir | 4.64101E-05 | 6 |
| hsa04062 | Chemokine signaling pathway | 0.000794213 | 6 |
| hsa04060 | Cytokine-cytokine receptor interaction | 0.003064319 | 6 |
| HALLMARK_INTERFERON_ALPHA_RESPONSE | HALLMARK_INTERFERON_ALPHA_RESPONSE | 0.00023217 | 6 |
| GO:0048020 | CCR chemokine receptor binding | 2.53968E-06 | 5 |
| GO:0042277 | peptide binding | 0.006989705 | 5 |
| GO:0033218 | amide binding | 0.015600124 | 5 |
| GO:0019886 | antigen processing and presentation of exogenous | 4.28783E-07 | 5 |
| GO:0002495 | antigen processing and presentation of peptide ant | 6.29607E-07 | 5 |
| GO:0002504 | antigen processing and presentation of peptide or | 7.15987E-07 | 5 |

| | | | |
|---|---|---|---|
| GO:0002478 | antigen processing and presentation of exogenous | 1.17948E-06 | 5 |
| GO:0048002 | antigen processing and presentation of peptide ant | 1.05134E-05 | 5 |
| GO:0071675 | regulation of mononuclear cell migration | 0.000164988 | 5 |
| GO:0050921 | positive regulation of chemotaxis | 0.000319568 | 5 |
| GO:0002687 | positive regulation of leukocyte migration | 0.000353428 | 5 |
| GO:0071222 | cellular response to lipopolysaccharide | 0.001854209 | 5 |
| GO:0070374 | positive regulation of ERK1 and ERK2 cascade | 0.001854209 | 5 |
| GO:0071219 | cellular response to molecule of bacterial origin | 0.002183299 | 5 |
| GO:0071216 | cellular response to biotic stimulus | 0.003371276 | 5 |
| GO:0070663 | regulation of leukocyte proliferation | 0.003801644 | 5 |
| GO:0043547 | positive regulation of GTPase activity | 0.004456094 | 5 |
| GO:0070372 | regulation of ERK1 and ERK2 cascade | 0.00651078 | 5 |
| GO:0070371 | ERK1 and ERK2 cascade | 0.008593165 | 5 |
| GO:0070661 | leukocyte proliferation | 0.009239407 | 5 |
| GO:0043087 | regulation of GTPase activity | 0.011808513 | 5 |
| GO:0030098 | lymphocyte differentiation | 0.016370457 | 5 |
| GO:0006816 | calcium ion transport | 0.019441624 | 5 |
| GO:0002683 | negative regulation of immune system process | 0.020300991 | 5 |
| GO:0001819 | positive regulation of cytokine production | 0.025905989 | 5 |
| GO:0043410 | positive regulation of MAPK cascade | 0.027668455 | 5 |
| GO:0030003 | cellular cation homeostasis | 0.027915911 | 5 |
| hsa04612 | Antigen processing and presentation | 0.000190132 | 5 |
| hsa05323 | Rheumatoid arthritis | 0.00033795 | 5 |
| hsa05164 | Influenza A | 0.002721644 | 5 |
| hsa05152 | Tuberculosis | 0.002721644 | 5 |
| GO:0023026 | MHC class II protein complex binding | 1.0916E-05 | 4 |
| GO:0023023 | MHC protein complex binding | 2.40656E-05 | 4 |
| GO:0048245 | eosinophil chemotaxis | 3.24669E-06 | 4 |
| GO:0072677 | eosinophil migration | 6.75556E-06 | 4 |
| GO:1901623 | regulation of lymphocyte chemotaxis | 1.10095E-05 | 4 |
| GO:2000401 | regulation of lymphocyte migration | 0.000319568 | 4 |
| GO:0002690 | positive regulation of leukocyte chemotaxis | 0.001062732 | 4 |
| GO:0071347 | cellular response to interleukin-1 | 0.001613449 | 4 |
| GO:0071887 | leukocyte apoptotic process | 0.001854209 | 4 |
| GO:0051928 | positive regulation of calcium ion transport | 0.002428674 | 4 |
| GO:0070555 | response to interleukin-1 | 0.003365998 | 4 |
| GO:0046718 | viral entry into host cell | 0.004136375 | 4 |
| GO:0044409 | entry into host | 0.004604823 | 4 |
| GO:0044000 | movement in host | 0.007106188 | 4 |
| GO:0051701 | biological process involved in interaction with host | 0.009612152 | 4 |
| GO:0045619 | regulation of lymphocyte differentiation | 0.01057264 | 4 |
| GO:0016064 | immunoglobulin mediated immune response | 0.011198597 | 4 |
| GO:0019724 | B cell mediated immunity | 0.011672474 | 4 |
| GO:0071356 | cellular response to tumor necrosis factor | 0.014186591 | 4 |
| GO:0050670 | regulation of lymphocyte proliferation | 0.015055008 | 4 |
| GO:0032944 | regulation of mononuclear cell proliferation | 0.01545041 | 4 |
| GO:0034612 | response to tumor necrosis factor | 0.017023143 | 4 |
| GO:0006898 | receptor-mediated endocytosis | 0.017449661 | 4 |
| GO:0051924 | regulation of calcium ion transport | 0.019829719 | 4 |
| GO:0006874 | cellular calcium ion homeostasis | 0.02283731 | 4 |
| GO:0043270 | positive regulation of ion transport | 0.02362765 | 4 |
| GO:0030217 | T cell differentiation | 0.025538073 | 4 |
| GO:0046651 | lymphocyte proliferation | 0.027458881 | 4 |
| GO:0055074 | calcium ion homeostasis | 0.027915911 | 4 |
| GO:0032943 | mononuclear cell proliferation | 0.028602902 | 4 |
| GO:1902105 | regulation of leukocyte differentiation | 0.030055347 | 4 |
| GO:0072503 | cellular divalent inorganic cation homeostasis | 0.031895863 | 4 |
| GO:0002440 | production of molecular mediator of immune respo | 0.033772882 | 4 |
| GO:0072507 | divalent inorganic cation homeostasis | 0.038641744 | 4 |
| GO:0002697 | regulation of immune effector process | 0.044702141 | 4 |
| hsa05150 | Staphylococcus aureus infection | 0.002721644 | 4 |
| hsa04640 | Hematopoietic cell lineage | 0.002721644 | 4 |
| hsa04514 | Cell adhesion molecules | 0.009212094 | 4 |
| hsa05169 | Epstein-Barr virus infection | 0.018354633 | 4 |
| GO:0004896 | cytokine receptor activity | 0.010342891 | 3 |
| GO:0008528 | G protein-coupled peptide receptor activity | 0.028022696 | 3 |
| GO:0001653 | peptide receptor activity | 0.029675908 | 3 |
| GO:0003823 | antigen binding | 0.036515146 | 3 |

| | | | |
|---|---|---|---|
| GO:0008201 | heparin binding | 0.036515146 | 3 |
| GO:0035747 | natural killer cell chemotaxis | 3.30227E-05 | 3 |
| GO:0002399 | MHC class II protein complex assembly | 0.000128749 | 3 |
| GO:0002503 | peptide antigen assembly with MHC class II protei | 0.000128749 | 3 |
| GO:0002396 | MHC protein complex assembly | 0.000230879 | 3 |
| GO:0002501 | peptide antigen assembly with MHC protein compl | 0.000230879 | 3 |
| GO:0019883 | antigen processing and presentation of endogenou | 0.000469834 | 3 |
| GO:2000403 | positive regulation of lymphocyte migration | 0.001647975 | 3 |
| GO:1902622 | regulation of neutrophil migration | 0.002099802 | 3 |
| GO:0007520 | myoblast fusion | 0.002416574 | 3 |
| GO:0070228 | regulation of lymphocyte apoptotic process | 0.004155811 | 3 |
| GO:0000768 | syncytium formation by plasma membrane fusion | 0.00444707 | 3 |
| GO:0140253 | cell-cell fusion | 0.00444707 | 3 |
| GO:0006949 | syncytium formation | 0.005119398 | 3 |
| GO:0071677 | positive regulation of mononuclear cell migration | 0.006007891 | 3 |
| GO:0002381 | immunoglobulin production involved in immunoglo | 0.006108371 | 3 |
| GO:0070227 | lymphocyte apoptotic process | 0.007253778 | 3 |
| GO:2000106 | regulation of leukocyte apoptotic process | 0.010435637 | 3 |
| GO:0070664 | negative regulation of leukocyte proliferation | 0.011100901 | 3 |
| GO:0002824 | positive regulation of adaptive immune response b | 0.015576179 | 3 |
| GO:0035710 | CD4-positive, alpha-beta T cell activation | 0.015817318 | 3 |
| GO:0045582 | positive regulation of T cell differentiation | 0.016227055 | 3 |
| GO:0002821 | positive regulation of adaptive immune response | 0.017023143 | 3 |
| GO:0050868 | negative regulation of T cell activation | 0.019810778 | 3 |
| GO:0014902 | myotube differentiation | 0.019944488 | 3 |
| GO:0045621 | positive regulation of lymphocyte differentiation | 0.020231013 | 3 |
| GO:0001909 | leukocyte mediated cytotoxicity | 0.021237788 | 3 |
| GO:0050671 | positive regulation of lymphocyte proliferation | 0.025538073 | 3 |
| GO:1903038 | negative regulation of leukocyte cell-cell adhesion | 0.025853069 | 3 |
| GO:0032946 | positive regulation of mononuclear cell proliferatic | 0.026487297 | 3 |
| GO:0008360 | regulation of cell shape | 0.027915911 | 3 |
| GO:0051250 | negative regulation of lymphocyte activation | 0.032889664 | 3 |
| GO:0070665 | positive regulation of leukocyte proliferation | 0.033234872 | 3 |
| GO:1904064 | positive regulation of cation transmembrane trans | 0.034627967 | 3 |
| GO:0046631 | alpha-beta T cell activation | 0.035685442 | 3 |
| GO:0061025 | membrane fusion | 0.036912723 | 3 |
| GO:0045580 | regulation of T cell differentiation | 0.037686717 | 3 |
| GO:0002822 | regulation of adaptive immune response based on | 0.038556653 | 3 |
| GO:0042129 | regulation of T cell proliferation | 0.038556653 | 3 |
| GO:1902107 | positive regulation of leukocyte differentiation | 0.038641744 | 3 |
| GO:1903708 | positive regulation of hemopoiesis | 0.038641744 | 3 |
| GO:0034767 | positive regulation of ion transmembrane transpor | 0.039584304 | 3 |
| GO:0001906 | cell killing | 0.040906589 | 3 |
| GO:0002695 | negative regulation of leukocyte activation | 0.044509542 | 3 |
| GO:0002819 | regulation of adaptive immune response | 0.044509542 | 3 |
| GO:0022408 | negative regulation of cell-cell adhesion | 0.046272516 | 3 |
| hsa05310 | Asthma | 0.002211814 | 3 |
| hsa05330 | Allograft rejection | 0.002721644 | 3 |
| hsa05332 | Graft-versus-host disease | 0.002721644 | 3 |
| hsa04940 | Type I diabetes mellitus | 0.002721644 | 3 |
| hsa04672 | Intestinal immune network for IgA production | 0.003470152 | 3 |
| hsa05320 | Autoimmune thyroid disease | 0.004096282 | 3 |
| hsa05416 | Viral myocarditis | 0.00553629 | 3 |
| hsa05321 | Inflammatory bowel disease | 0.006593221 | 3 |
| hsa05140 | Leishmaniasis | 0.009650723 | 3 |
| hsa04658 | Th1 and Th2 cell differentiation | 0.015212632 | 3 |
| hsa04657 | IL-17 signaling pathway | 0.01542457 | 3 |
| hsa04620 | Toll-like receptor signaling pathway | 0.018746836 | 3 |
| hsa04659 | Th17 cell differentiation | 0.019980989 | 3 |
| hsa05145 | Toxoplasmosis | 0.020722621 | 3 |
| hsa04142 | Lysosome | 0.032029508 | 3 |
| hsa05322 | Systemic lupus erythematosus | 0.034142312 | 3 |
| GO:0032395 | MHC class II receptor activity | 0.002502759 | 2 |
| GO:0045236 | CXCR chemokine receptor binding | 0.006989705 | 2 |
| GO:0042605 | peptide antigen binding | 0.020955765 | 2 |
| GO:0002645 | positive regulation of tolerance induction | 0.003801644 | 2 |
| GO:0002468 | dendritic cell antigen processing and presentation | 0.005369499 | 2 |
| GO:0070234 | positive regulation of T cell apoptotic process | 0.005369499 | 2 |

| | | | |
|---|---|---|---|
| GO:0002830 | positive regulation of type 2 immune response | 0.007106188 | 2 |
| GO:0070230 | positive regulation of lymphocyte apoptotic proces | 0.007106188 | 2 |
| GO:0002643 | regulation of tolerance induction | 0.008593165 | 2 |
| GO:0140131 | positive regulation of lymphocyte chemotaxis | 0.009206605 | 2 |
| GO:1901739 | regulation of myoblast fusion | 0.009206605 | 2 |
| GO:0002689 | negative regulation of leukocyte chemotaxis | 0.01057264 | 2 |
| GO:0071676 | negative regulation of mononuclear cell migration | 0.011198597 | 2 |
| GO:0032693 | negative regulation of interleukin-10 production | 0.011808513 | 2 |
| GO:0042832 | defense response to protozoan | 0.013563343 | 2 |
| GO:0001562 | response to protozoan | 0.014186591 | 2 |
| GO:0030194 | positive regulation of blood coagulation | 0.014186591 | 2 |
| GO:1900048 | positive regulation of hemostasis | 0.014186591 | 2 |
| GO:0090025 | regulation of monocyte chemotaxis | 0.014952333 | 2 |
| GO:0050820 | positive regulation of coagulation | 0.01545041 | 2 |
| GO:0060142 | regulation of syncytium formation by plasma meml | 0.01545041 | 2 |
| GO:2000108 | positive regulation of leukocyte apoptotic process | 0.01545041 | 2 |
| GO:0001916 | positive regulation of T cell mediated cytotoxicity | 0.015817318 | 2 |
| GO:0002507 | tolerance induction | 0.015817318 | 2 |
| GO:0002828 | regulation of type 2 immune response | 0.017204759 | 2 |
| GO:0070229 | negative regulation of lymphocyte apoptotic proce | 0.020300991 | 2 |
| GO:0042092 | type 2 immune response | 0.021086346 | 2 |
| GO:0043368 | positive T cell selection | 0.021086346 | 2 |
| GO:0048246 | macrophage chemotaxis | 0.02283731 | 2 |
| GO:0001914 | regulation of T cell mediated cytotoxicity | 0.023464664 | 2 |
| GO:0051281 | positive regulation of release of sequestered calciu | 0.023464664 | 2 |
| GO:0070232 | regulation of T cell apoptotic process | 0.023464664 | 2 |
| GO:0046596 | regulation of viral entry into host cell | 0.027668455 | 2 |
| GO:0001913 | T cell mediated cytotoxicity | 0.031954073 | 2 |
| GO:0043277 | apoptotic cell clearance | 0.031954073 | 2 |
| GO:0045058 | T cell selection | 0.031954073 | 2 |
| GO:0034381 | plasma lipoprotein particle clearance | 0.033603871 | 2 |
| GO:0052372 | modulation by symbiont of entry into host | 0.033603871 | 2 |
| GO:0002686 | negative regulation of leukocyte migration | 0.035257072 | 2 |
| GO:2000107 | negative regulation of leukocyte apoptotic process | 0.035257072 | 2 |
| GO:0010332 | response to gamma radiation | 0.036912723 | 2 |
| GO:0070231 | T cell apoptotic process | 0.036912723 | 2 |
| GO:0001912 | positive regulation of leukocyte mediated cytotoxi | 0.038556653 | 2 |
| GO:0043903 | regulation of biological process involved in symbiot | 0.038556653 | 2 |
| GO:0002711 | positive regulation of T cell mediated immunity | 0.038641744 | 2 |
| GO:1905517 | macrophage migration | 0.038641744 | 2 |
| GO:0090303 | positive regulation of wound healing | 0.040613239 | 2 |
| GO:0032757 | positive regulation of interleukin-8 production | 0.041438157 | 2 |
| GO:0050918 | positive chemotaxis | 0.042487828 | 2 |
| GO:0032613 | interleukin-10 production | 0.043314542 | 2 |
| GO:0032653 | regulation of interleukin-10 production | 0.043314542 | 2 |
| GO:0031343 | positive regulation of cell killing | 0.044141012 | 2 |
| GO:0034605 | cellular response to heat | 0.044141012 | 2 |
| GO:0031638 | zymogen activation | 0.044702141 | 2 |
| GO:0050922 | negative regulation of chemotaxis | 0.04661828 | 2 |
| GO:0030193 | regulation of blood coagulation | 0.04744311 | 2 |
| GO:0042130 | negative regulation of T cell proliferation | 0.04744311 | 2 |
| GO:1900046 | regulation of hemostasis | 0.049823728 | 2 |

**Table S3. [Enriched pathways for each myeloid-derived irMP], Related to Figure 1.**
"GeneRatio" is the fraction of irMP genes that are found in the corresponding enriched pathway. "BgRatio" is the fraction of genes annotated to the enriched pathway versus the number of genes annotated to that specific pathway database within the background. "geneID" contains the irMP genes that are in the enriched pathway.

| V1 | V2 | V3 | V4 | V5 | V6 | V7 |
|---|---|---|---|---|---|---|
| CXCL10 | HLA-DQA1 | CXCL9 | LGALS2 | S100A12 | ACTB | CD14 |
| HLA-DRB4 | CXCL8 | CXCL11 | CD1D | VCAN | FTL | SEPP1 |
| PLAC8 | CCL18 | CXCL10 | FOSB | VNN2 | PSAP | DHRS2 |
| IFIT1 | PPBP | MT1M | SELL | SIK1 | GAPDH | FOLR2 |
| ATF4 | IRF2BPL | HLA-DQB1 | BRE-AS1 | PLAC8 | FTH1 | OLFML2B |
| USP18 | MMP9 | RP1-93H18.6 | NR4A2 | BRE-AS1 | TMSB10 | CCL26 |
| CD38 | PLSCR1 | IL12B | IMPA2 | FOSB | ND4 | SIGLEC1 |
| HERC5 | EGFR | SERPINB7 | RGS2 | PTGS2 | CST3 | IFITM3 |
| EIF3F | MMP12 | IDO2 | EGR3 | NR4A2 | RP5-882O7.1 | CFH |
| NMB | CDKN1B | ANKRD22 | BTG1 | PDE4B | HMOX1 | CCL24 |
| ODC1 | Y16709 | CD38 | RBP7 | RNASE2 | LGALS1 | IGFBP2 |
| CCR7 | CSTA | SERPING1 | F13A1 | DDIT4 | OAZ1 | MAFB |
| EIF3L | CD74 | GBP5 | NAPSB | RBP7 | HLA-DRA | HLA-DRB4 |
| IFITM1 | ADM | RARRES3 | ICAM3 | OLIG1 | LYZ | AP2A2 |
| NACA | GSAP | IFI27 | MS4A6A | SELL | FUCA1 | RNASE1 |
| MYOF | CCR5 | IL32 | P2RY13 | PROK2 | TYROBP | FGF13 |
| HLA-C | ANXA2 | FAM26F | TMEM154 | BTG1 | HLA-A | COLEC12 |
| SP110 | HLA-DQB1 | CCL19 | ITGA4 | ID1 | S100A8 | MS4A4A |
| HLA-DRA | CHSY1 | SUCNR1 | GPR160 | HLA-DQB1 | S100A9 | CCL23 |
| RSAD2 | IFIT1 | GBP4 | SLC18B1 | FCGR1B | PGD | GPR34 |
| SQSTM1 | SEC22B | VAMP5 | SIK1 | ALDH1A1 | NFKBIA | CD163L1 |
| KARS | MAFB | ETV7 | C10ORF54 | PID1 | CD74 | GFRA2 |
| HLA-E | TUBA1A | BATF2 | MXI1 | CDKN2D | EEF1A1 | SLC40A1 |
| ACOT9 | CSRP2 | CFB | SMARCD3 | DUSP1 | CTSL | MAP1LC3C |
| MLLT11 | FTH1P5 | GBP3 | AMICA1 | CD244 | LIPA | CD59 |
| TAP1 | B2M | GBP1P1 | MIR-223 | HPSE | HLA-DPA1 | C3AR1 |
| SLC25A6 | MTHFD2 | TNFSF10 | ASGR1 | LINC00936 | GLUL | PLTP |
| NCF2 | RB1 | CCL8 | DPEP2 | FPR1 | CCL4 | IFITM2 |
| SRSF4 | RGL1 | SYNPO2 | P2RY8 | STK17B | BASP1 | SERPINF1 |
| DNTTIP2 | GCH1 | LRRK2 | CDA | RGS2 | IL1B | CNRIP1 |
| MX2 | FCGR2B | HAPLN3 | PGM1 | GCH1 | TNFAIP6 | CLEC4G |
| CMPK2 | EVI2A | RAB12 | ADD3 | MXD1 | RNASE1 | MYO1G |
| IFI44L | UBE2F | ITGB8 | SLC24A4 | IFI44L | CHI3L1 | PDCD1LG2 |
| PPIA | ACTB | LINC00158 | LY86 | SERPINB2 | CXCL10 | TMEM176A |
| ISG15 | CCL8 | NEURL3 | EGR1 | CACNA2D3 | IFIT2 | AKR1B1 |
| TNFSF10 | UBB | IL27 | NR4A1 | CXCL3 | CCL17 | P2RY13 |
| CSRP2 | TXN | SLC6A12 | ARL4C | RPH3A | S100A10 | GPR171 |
| NT5C3A | HLA-DPA1 | RSAD2 | PECAM1 | G0S2 | LILRA3 | MAP7 |
| ISG20 | CMPK2 | IDO1 | CACNA2D3 | C9ORF72 | GPNMB | MSRB2 |
| SRXN1 | ZFAND5 | GBP1 | CLEC2B | FFAR2 | SPP1 | S100A9 |
| CCNG1 | HUWE1 | PRKAR2B | BCL10 | HAL | HCAR3 | IGFBP4 |
| CIRBP | PGAM1 | IFIT3 | KLF4 | IFITM1 | IFITM3 | LGALS3BP |
| MX1 | MRFAP1 | AIM2 | PROK2 | CDKN1C | SORL1 | CCL8 |
| SAMD9L | HLA-DPB1 | PSMB9 | ZNF395 | MAP3K7CL | XPO6 | MAGED1 |
| TFDP2 | DCK | CCND2 | TESC | IFITM3 | LGMN | C1QC |
| B2M | CEBPB | IL15RA | SYTL1 | IFITM2 | HMGN2 | CREM |
| RRAGC | RAB13 | ISG20 | CD4 | RNF103 | MT2A | LINC01094 |
| RAB29 | TNFAIP8 | APOL4 | METTL7A | ZBTB24 | MT1X | PLAU |
| NFKB1 | ITM2B | E2F7 | EIF4G3 | HP | MT1H | SCARB1 |
| AKR1C3 | LAP3 | SPP1 | RNASE2 | CTC-510F12.4 | MT1HL1 | SIPA1L1 |

| V8 | V9 |
|---|---|
| NLRP3 | CCL8 |
| SEPP1 | CCL2 |
| HLA-DRB4 | C15ORF48 |
| CCL23 | CXCL10 |
| TMEM71 | MRC1 |
| FCGR2B | CCRL2 |
| MS4A6A | CCL4 |
| GIMAP7 | ANKRD22 |
| CFH | CH25H |
| GSAP | CD74 |
| MAFB | CCL7 |
| LGMN | C12ORF4 |
| C3AR1 | FPR3 |
| SATB1 | BATF2 |
| CD2 | HLA-DRA |
| TRAC | CXCL9 |
| MS4A4A | GPR84 |
| GPR171 | SLAMF7 |
| TRBC1 | IDO1 |
| HLA-DQB1 | P2RY14 |
| CD14 | VAMP5 |
| CLC | SLAMF8 |
| SELL | RP1-93H18.6 |
| IRF2BPL | CD274 |
| KCTD12 | PLAU |
| CCL18 | MSR1 |
| CLEC12A | SMCO4 |
| CCL26 | CTSL |
| TIMP1 | MYOF |
| RNASE1 | PSTPIP2 |
| CRIM1 | M6PR |
| NR1D2 | LIPA |
| TMEM176B | TIFA |
| HSPA5 | FCGR1B |
| LCP2 | F3 |
| ABCA6 | LAP3 |
| HLA-DMB | HLA-DMA |
| CFD | RHOH |
| MS4A7 | GBP4 |
| EVI2A | AC079767.4 |
| SUCNR1 | RALGDS |
| NUCB2 | TIMM10 |
| HLA-DMA | RAB12 |
| ETS2 | TFEC |
| GZMK | HLA-DPA1 |
| GLUL | CREM |
| IL1R2 | CD1B |
| EIF1B | FAM26F |
| CDKN1B | LINC00996 |
| CPVL | CD69 |

**Table S4. [Myeloid-derived gene sets], Related to Figure 1.**
The 9 myeloid-derived irMPs with their constituent genes. Each column is one irMP.

### Supplementary Notes

The table below summarizes the rationales behind each gene set annotation. We especially thank **Matthew Gubin Ph.D.; Hind Rafei M.D.; Rafet Basar M.D.; Katy Rezvani M.D/Ph.D.; Weiyi Peng M.D/Ph.D.** for their help on commenting on the first-effort annotations and refining the annotations.

| | **Lymphoid-derived Gene sets Final Annotation** | **Datasets for validation and rationale** | **Comments and Annotations from MDACC Immunologists** |
|---|---|---|---|
| 1 | Cell Cycle 1 | COVID-19 single cell atlas<br>https://www.ncbi.nlm.nih.gov/geo/query/acc.cgi?acc=GSE150728<br><br>The authors have pre-identified a cluster of highly proliferative T cells with effector functions, and we expect this cell cycle pathway to have high activity in this cluster of cells in comparison to other T cells. (Fig 2c) | Agree |
| 2 | TCR Anchoring | T cell stimulation<br>https://www.ncbi.nlm.nih.gov/geo/query/acc.cgi?acc=GSE126030<br><br>The authors have profiled T cells in lung, blood, lymph nodes and bone marrow before and after anti-CD28/CD3 stimulations. TCR signaling are to be upregulated when T cells are engaged and activated, thus by examining the gene set activities in resting vs activated T cells, we can be certain of their function. (Fig 3c) | • This gene set is highly overlapped with KEGG-TCR signaling pathway (Figure S9c) and is highly expressed in regions enriched for CD4 and CD8 T cells in spatial image (Fig 6d).<br>• This gene set contains T cell stem like signature [possibly memory or recently activation (*CD69*, *CD52*)].<br>• *ID2*, one from this gene set, is usually upregulated in activated CD8 and memory CD8 T cells. |
| 3 | Histone associated lipid antigen presentation | pTMB TCGA<br>https://gdc.cancer.gov/about-data/publications/panimmune<br><br>Since samples with high persistent tumor mutational burden are more likely to render themselves visible to the immune system by providing a constant flow of antigen presentation, we expect this gene set to have high activities in TCGA samples with high pTMB (Fig 3d, Fig S9d) | • Enriched in Histone genes (*HIST* genes).<br>• *CD1A*, *CD1B*, *CD1C* (non-classical MHC proteins, related to the class I MHC proteins, present lipid antigens to NK T cells). |
| 4 | Interleukin induced Treg activation | COVID PBMC CITE-seq<br>https://www.ncbi.nlm.nih.gov/geo/query/acc.cgi?acc=GSE155224<br><br>CITE-seq data has profiled CD25, which is an important IL2 receptor on Tregs, once bind to IL2, will sustain the survival of Tregs. This CITE-seq data has also profiled protein markers informative of effector Tregs, such as TIGIT and TNFRSF9. (Fig S10d) | • While *CTLA4*, *IL2RA (CD25)*, *TIGIT*, *TNFRSF9 (4-1BB/CD137)*, *IL1R2*, are often expressed on activated conventional T cells to some degree, these are often highly expressed on suppressive CD4+ T regulatory cells (Tregs). Further, *DUSP4* and *SOCS2* are often highly expressed in Tregs.<br>• *IL2RA (CD25)* is one of the top 3 genes in term of NMF loading (importance) in this gene set, and the binding between IL2 and IL2RA is essential for Treg survival. |

| | | | |
|---|---|---|---|
| 5 | Antigen Presentation | pTMB TCGA<br>https://gdc.cancer.gov/about-data/publications/panimmune<br><br>Since samples with high persistent tumor mutational burden are more likely to render themselves visible to the immune system by providing a constant flow of antigen presentation, we expect this gene set to have high activities in TCGA samples with high pTMB (Fig 3d, Fig S9d) | • *CD1B*, *CD1C*, *CD1E* (non-classical MHC proteins, related to the class I MHC proteins, present lipid antigens to NK T cells. Also has *RAG1*, *RAG2* (related to VDJ recombination, B and T cell (especially thymocyte)) and *CD8A*.<br>• Enriched in cell cycle/DNA synthesis/proliferation genes (*TUBB*, *BUB1B*, *DLGAP5*, *HMGB2*, *RRM2*, *PTTG1*). But these genes have relatively low loadings (importance) in this gene set. It is possible these proliferation genes are downstream of the antigen presentation. |
| 6 | Cell Cycle 2 | COVID-19 single cell atlas<br>https://www.ncbi.nlm.nih.gov/geo/query/acc.cgi?acc=GSE150728<br><br>The authors have pre-identified a cluster of highly proliferative T cells with effector functions, and we expect this cell cycle pathway to have high activity in this cluster of cells in comparison to other T cells. (Fig 2c) | Agree |
| 7 | Chemokine Mediated T cell activation | COVID-19 cytokine storm<br>https://www.ncbi.nlm.nih.gov/geo/query/acc.cgi?acc=GSE150861<br><br>The authors performed scRNA-seq analysis on two severe COVID-19 patients diagnosed with cytokine storms (CS). As patients with CS express more cytokine/chemokines than healthy controls, we expect this gene set to be upregulated in CS patients. By comparing the activity of this gene set between these two patients and healthy controls, we can confirm that this gene set is associated with cytokine/chemokine release. (Fig 2a) | • Enriched in Cytokine signaling (*CCR2*, *CXCL8*, *CXCL9*)<br>• Contains T cell activation transcripts. Cytotoxic transcripts (*GZMB*, *GZMA*) |
| 8 | MHC mediated immunity | COVID PBMC CITE-seq<br>https://www.ncbi.nlm.nih.gov/geo/query/acc.cgi?acc=GSE155224<br><br>CITE-seq data has profiled the proteins related to MHC, APC activation, such as CD80 and CD86, and downstream immune cell activations, such as HLA proteins, TCR proteins and proteins related to T cell activation such as CD28, CD69. A typical immune activation would first involve danger recognition, APC activation, APC presentation and immune cell activation, and we would expect different transcripts/proteins to be upregulated at different stage. Therefore, by comparing abundance in these protein markers, we can refine the annotation by narrowing down whether this gene set is more related to p-MHC interface or pMHC-TCR interface. (Fig S10b) | • Has features of MHC-I and MHC-II antigen presentation. Has features of B cells (which also can be robust antigen presenting cells, and B cells also express MHC-II genes)<br>• *GRZB* and *KLRB1*, associated with Cytotoxic T cells/Natural Killer (NK) cells, suggesting activation through MHC complex |
| 9 | Metabolic Process | CD8+ T cell stimulation<br>https://www.ncbi.nlm.nih.gov/geo/query/acc.cgi?acc=GSE211602 | Agree. Also enriched in a few proliferation or cell cycle transcripts. |

| | | | |
|---|---|---|---|
| | | Authors have stimulated naïve CD8+ T cells in vitro and used scRNAseq to profile the metabolic profiles in different T cell activation stages. They observed an increase in both glycolysis and oxidative phosphorylation as T cells moves from naïve to late activation stages. We can calculate the activity of our gene set and see if it increases with T cell activation. (Fig 2d) | |
| 10 | Lipid Localization TCR synapse | T cell stimulation<br>https://www.ncbi.nlm.nih.gov/geo/query/acc.cgi?acc=GSE126030<br><br>The authors have profiled T cells in lung, blood, lymph nodes and bone marrow before and after anti-CD28/CD3 stimulations. TCR signaling are to be upregulated when T cells are engaged and activated, thus by examining the gene set activities in resting vs activated T cells, we can be certain of their function. (Fig 3c) | |
| 11 | Cell Cycle Immune Response | COVID-19 single cell atlas<br>https://www.ncbi.nlm.nih.gov/geo/query/acc.cgi?acc=GSE150728<br><br>The authors have pre-identified a cluster of highly proliferative T cells with effector functions, and we expect this cell cycle pathway to have high activity in this cluster of cells in comparison to other T cells. (Fig 2c) | Cell cycle T cell activation/effector function (*IFNG, TNFRSF9 (4-1BB/CD137), GZMB, LAG3*) |
| 12 | Lymphocyte Activation | COVID PBMC CITE-seq<br>https://www.ncbi.nlm.nih.gov/geo/query/acc.cgi?acc=GSE155224<br><br>CITE-seq data has profiled the proteins related to lymphocyte activations, such as CD16, CD69, CD8, CD3, along with some checkpoint transcripts that will also be upregulated during lymphocyte activation to maintain homeostasis, such as TIGIT, CTLA-4, and PDCD1. By comparing abundance in these protein markers, we can be certain that this gene set is related to lymphocyte activation, or even be able to narrow down the type of cells that are activated. (Fig 3a) | Enriched in T cells markers [*CD3D*, (*RAG1*, *RAG2* (B and T cell, especially thymocyte))] |
| 13 | Interferon induced Antiviral Defense | Cytokine Dictionary<br>https://www.immune-dictionary.org/app/home<br><br>The authors have profiled scRNA-seq on 17 different immune cell types extracted from mouse lymph nodes when treated with 86 different cytokines *in vivo* or PBS as control. We expect this gene set to be upregulated when cells are treated with interferons. (Fig 2b) | Enriched in GO term (defense response to virus, defense response to symbiont, viral process, viral life cycle regulation response to virus, and viral genome replication), which are driven by the following genes*: IFITM1*, *IL12RB1*, *IFI44L*, *ISG20* |
| 14 | Type-I Interferon | Cytokine Dictionary<br>https://www.immune-dictionary.org/app/home | Agree |

| | | | |
|---|---|---|---|
| | | The authors have profiled scRNA-seq on 17 different immune cell types extracted from mouse lymph nodes when treated with 86 different cytokines *in vivo* or PBS as control. We expect this gene set to be upregulated when cells are treated with interferons. (Fig 2b) | |
| 15 | Cell Adhesion 1 | PBMC CITE-seq<br>https://www.ncbi.nlm.nih.gov/geo/query/acc.cgi?acc=GSE213282<br><br>CITE-seq data has profiled the proteins related to cell adhesions, such as CD11a, CD11b, CD11c, integrin, CD49, CD62. By comparing abundance in these protein markers, we can be certain that this gene set is related to cell adhesion, or even be able to narrow down the stage of cell adhesion (rolling, weak, or extravasation) (Fig S9b) | Cell Adhesion 1 |
| 16 | Cell Adhesion 2 (CD52+) | PBMC CITE-seq<br>https://www.ncbi.nlm.nih.gov/geo/query/acc.cgi?acc=GSE213282<br><br>CITE-seq data has profiled the proteins related to cell adhesions, such as CD11a, CD11b, CD11c, integrin, CD49, CD62. By comparing abundance in these protein markers, we can be certain that this gene set is related to cell adhesion, or even be able to narrow down the stage of cell adhesion (rolling, weak, or extravasation) (Fig 9b) | • Cell Adhesion 2 (CD52+)<br>• HLA/MHC antigen presenting *HLA-DQB1*, *HLA-DMA*, *HLA-E, HLA-A* |
| 17 | IFN-gamma-induced cytotoxicity | COVID PBMC CITE-seq<br>https://www.ncbi.nlm.nih.gov/geo/query/acc.cgi?acc=GSE155224<br><br>CITE-seq data has profiled the proteins related to cytotoxicity, such as CD16, CD69, CD8, CD3, along with some checkpoint transcripts that will also be upregulated during activation to maintain homeostasis, such as TIGIT, CTLA-4, and PDCD1. (Fig 3b)<br><br>COVID-19 single cell atlas<br>https://www.ncbi.nlm.nih.gov/geo/query/acc.cgi?acc=GSE150728<br><br>The author has defined cytotoxicity score, and if our gene set activity aligns with this score, we can be more certain that this gene set is related to cytotoxicity. (Fig S9c) | • Enriched in granzyme genes, suggesting cytotoxicity.<br>• This gene set has *LYZ* (macrophage marker) and MHC-II genes (APC markers)<br>• In the spatial image (Fig s13.17), this gene set light up the most among macrophages and second most among T cells. The rationale is that activated T cells will secrete IFNG, then the IFNG will activate macrophages to eat/kill the target cells. |
| 18 | Cytokine Receptor Signaling | COVID-19 cytokine storm<br>https://www.ncbi.nlm.nih.gov/geo/query/acc.cgi?acc=GSE150861<br><br>The authors performed scRNA-seq analysis on two severe COVID-19 patients diagnosed with cytokine storms (CS). As patients with CS express more cytokine/chemokines than healthy controls, we expect this gene set | Since this gene set has more cytokine receptors than cytokine, it is more specific to name it "cytokine receptor"<br>*IL1R1/CSF2RB/CXCR6/IL1R2/IL2RB/CCR2/CCR6 /IL2RA/TNFRSF1B* |

| | | | |
|---|---|---|---|
| | | to be upregulated in CS patients. By comparing the activity of this gene set between these two patients and healthy controls, we can confirm that this gene set is associated with inflammatory cytokine release. (Fig 2a) | |
| 19 | T cell activation | COVID PBMC CITE-seq<br>https://www.ncbi.nlm.nih.gov/geo/query/acc.cgi?acc=GSE155224<br><br>CITE-seq data has profiled the proteins related to lymphocyte activations, such as CD16, CD69, CD8, CD3, along with some checkpoint transcripts that will also be upregulated during lymphocyte activation to maintain homeostasis, such as TIGIT, CTLA-4, and PDCD1. By comparing abundance in these protein markers, we can be certain that this gene set is related to lymphocyte activation, or even be able to narrow down the type of cells that are activated. (Fig 3a) | • This gene set has NF-KB signaling associated genes.<br>• MHC-I antigen presentation (*B2M*, *HLA-F*, *HLA-A*)<br>• This gene set has negative association with CD16, CD56, and KLRG1 protein abundance, positive association with CD8, CD3 and CD4. (Fig 3a) |

| | **Myeloid-derived Gene sets Final Annotation** | **Datasets for validation and rationale** | **Comments and Annotations from MDACC Immunologists** |
|---|---|---|---|
| 1 | Antiviral Defense Network | Cytokine Dictionary<br>https://www.immune-dictionary.org/app/home<br><br>The authors have profiled scRNA-seq on 17 different immune cell types extracted from mouse lymph nodes when treated with 86 different cytokines *in vivo* or PBS as control. We expect this gene set to be upregulated when cells are treated with interferons. (Fig 2b) | This gene set is enriched in interferon stimulated genes: *ISG15*, *ISG20*, *IFITM1*, *IFIT1*, *IFI44L*. These genes are important in antiviral response. |
| 2 | Antigen Processing | COVID PBMC CITE-seq<br>https://www.ncbi.nlm.nih.gov/geo/query/acc.cgi?acc=GSE155224<br><br>CITE-seq data has profiled proteins related to MHC, APC activation (CD80/CD86), and downstream immune cell activations (HLA proteins, TCR proteins and proteins related to T cell activation such as CD28, CD69). A typical immune activation would first involve danger recognition, APC activation, APC presentation and immune cell activation, and we would expect different transcripts/proteins to be upregulated at different stage. Therefore, by comparing abundance in these protein markers, we can refine the annotation by narrowing down whether this gene set is more related to p-MHC interface or pMHC-TCR interface. | • The PPI network of this gene set forms a module of MHC-II molecules (*HLA-DPA1*, *HLA-DQA1*, *HLA-DPB1*, *HLA-DQB1*), suggesting APC activation and presentation |

| | | | |
|---|---|---|---|
| | | (Fig S10a) | |
| 3 | Cytokine Production | COVID-19 cytokine storm<br>https://www.ncbi.nlm.nih.gov/geo/query/acc.cgi?acc=GSE150861<br><br>The authors performed scRNA-seq analysis on two severe COVID-19 patients diagnosed with cytokine storms (CS). As patients with CS express more cytokine/chemokines than healthy controls, we expect this gene set to be upregulated in CS patients. By comparing the activity of this gene set between these two patients and healthy controls, we can confirm that this gene set is associated with inflammatory cytokine release. (Fig 2a) | This gene set is enriched in cytokine and chemokines: *CXCL9/CXCL11/CXCL10/CD38/ SERPING1/IFI27/GBP4/VAMP5/ BATF2/CFB/TNFSF10/RSAD2/IDO1 /IFIT3/PSMB9/IL15RA/ISG20* |
| 4 | Cell Adhesion 1 | PBMC CITE-seq<br>https://www.ncbi.nlm.nih.gov/geo/query/acc.cgi?acc=GSE213282<br><br>CITE-seq data has profiled the proteins related to cell adhesions, such as CD11a, CD11b, CD11c, integrin, CD49, CD62. By comparing abundance in these protein markers, we can be certain that this gene set is related to cell adhesion, or even be able to narrow down the stage of cell adhesion (rolling, weak, or extravasation) (Fig S9a) | • *SELL*, *ITGA4*, *PECAM1* are cell adhesion genes.<br>• Enriched in leukocyte cell-cell adhesion pathway driven by *CD1D/SELL/EGR3/ITGA4/SMARCD3/ PECAM1/BCL10/KLF4/CD4* |
| 5 | Modulation of Cell Migration | PBMC CITE-seq<br>https://www.ncbi.nlm.nih.gov/geo/query/acc.cgi?acc=GSE213282<br><br>CITE-seq data has profiled the proteins related to cell adhesions, such as CD11a, CD11b, CD11c, integrin, CD49, CD62. If we don't see association between these protein abundance with the activity level of this gene set, then we can say that this gene set is not related to cell adhesion. (Fig S9a) | • *SIK1* regulate cytoskeletal dynamics and cell polarity. It may affect the activity of proteins involved in actin dynamics and focal adhesions, which are essential for cell migration.<br>• *FOSB* can modulate cell migration by regulating the expression of genes involved in cell motility, such as integrins.<br>• *PTGS2* influence cell migration through its involvement in the production of prostaglandins. Prostaglandins play a role in inflammation and can affect cell migration and invasion.<br>• *NR4A2* is a transcription factor that can influence cell migration by regulating the expression of genes involved in cell adhesion and motility. *NR4A2* inhibits mast cell migration by modulating production of the chemotactic cytokine. (doi: 10.1080/09553002.2019.1642535) |
| 6 | Cell Adhesion 2 | PBMC CITE-seq<br>https://www.ncbi.nlm.nih.gov/geo/query/acc.cgi?acc=GSE213282<br><br>CITE-seq data has profiled the proteins related to cell adhesions, such as CD11a, CD11b, CD11c, integrin, CD49, CD62. By comparing abundance in these protein markers, we can be certain that this gene set is related to | • *ACTB*: key component of the cytoskeleton and is essential for cell structure and movement. Actin filaments play a crucial role in cell adhesion and migration.<br>• *LGALS1* (Galectin-1): can interact with glycoproteins on the cell surface |

| | | | |
|---|---|---|---|
| | | cell adhesion, or even be able to narrow down the stage of cell adhesion (rolling, weak, or extravasation) (Fig S9a) | • The HLA genes (*HLA-DRA*, *HLA-A*, *HLA-DPA1*) are involved in cell adhesion during immune cell interactions and recognition.<br>• *S100A8* and *S100A9* can bind to cell surface receptors and contribute to cell adhesion and migration in the context of inflammation and immune responses. |
| 7 | Eosinophil chemotaxis | PBMC CITE-seq<br>https://www.ncbi.nlm.nih.gov/geo/query/acc.cgi?acc=GSE213282<br><br>CITE-seq data has profiled the proteins related to cell adhesions, such as CD11a, CD11b, CD11c, integrin, CD49, CD62. If we don't see association between these protein abundance with the activity level of this gene set, then we can say that this gene set is not related to cell adhesion. (Fig S9a) | Genes related to eosinophil chemotaxis: *CCL26/CCL24/CCL8* |
| 8 | MHC-II mediated Lymphocyte Activation | COVID PBMC CITE-seq<br>https://www.ncbi.nlm.nih.gov/geo/query/acc.cgi?acc=GSE155224<br><br>CITE-seq data has profiled proteins related to MHC-II genes, macrophages and CD4 activations. Since the function of this gene set is related to MHC-II dependent lymphocyte activation, which typically happen between a macrophage and a CD4 T cell, thus we would expect cells with high activity in this gene set show higher protein abundance in CD4 activation, MHC-II molecules, and macrophage markers. (Fig S10c) | • MHC-II genes (*HLA-DRB4/ TRBC1/HLA-DQB1 /HLA-DMB/HLA-DMA*)<br>• Lymphocyte activation genes (*TRAC*, *TRBC1*, *GZMK*, *MS4A4A*, *MS4A6A* for B cell activation) |
| 9 | Chemokine activity | COVID-19 cytokine storm<br>https://www.ncbi.nlm.nih.gov/geo/query/acc.cgi?acc=GSE150861<br><br>The authors performed scRNA-seq analysis on two severe COVID-19 patients diagnosed with cytokine storms (CS). As patients with CS express more cytokine/chemokines than healthy controls, we expect this gene set to be upregulated in CS patients. By comparing the activity of this gene set between these two patients and healthy controls, we can confirm that this gene set is associated with inflammatory cytokine release. (Fig 2a) | This gene set is enriched in chemokine genes (*CCL8/CCL2/CXCL10/CCRL2/CCL4/ CH25H/CD74/CCL7/CXCL9*), connecting with *CD69* (T cell activation gene) in the PPI network |
